# Sterically Controlled Conformational Gating of Rhodamine Spirolactam Switches for Super-Resolution Imaging

**DOI:** 10.64898/2026.11.16.767006

**Authors:** Shaowei Wu, Yalin Huang, Guanyu Jiang, Shurui Hu, Wenhao Jia, Xiaogang Liu, Qinglong Qiao, Zhaochao Xu

## Abstract

The reversible interconversion of rhodamines between nonfluorescent spirocyclic and fluorescent ring-open states underpins fluorogenic probes and spontaneously blinking dyes for super-resolution imaging. Existing strategies mainly tune the equilibrium between these states through electronic modification, whereas the role of conformational return in ring closure remains less explored. Here, we establish steric regulation of rhodamine spirolactams through amide N-substituents. Bulky substituents generate intramolecular repulsion with the xanthene scaffold, favoring ring opening. Crystallographic observations reveal substantial amide reorientation upon opening, suggesting increased separation between the nitrogen and meso-carbon reaction centers. We propose that substituent-dependent conformational accessibility modulates C–N bond reformation and the lifetime of the emissive state. The tert-butyl-substituted dye **1e** exhibits an open-state residence time of 145 ms, despite its lower p*K*_cycl_. Calculations also indicate a lower ring-opening barrier for **1e** than for less hindered **1a** (1.12 vs 1.37 eV). Incorporating this design into HaloTag probes enables SIM, STED, and SMLM imaging. Importantly, **N-halo-RBRB** enables dual-modal SIM/SMLM imaging with a single fluorophore, integrating the rapid reconstruction of overall cellular architecture by SIM with single-molecule localization information from SMLM. This work identifies sterically modulated conformational accessibility as a design principle for engineering reversible rhodamine switches with tailored imaging capabilities.

## Introduction

Fluorescence super-resolution microscopy has expanded the observation of subcellular structures beyond the diffraction limit.^[1–5]^ Although structured illumination microscopy (SIM), stimulated emission depletion (STED) microscopy and single-molecule localization microscopy (SMLM) operate through different optical principles, each places stringent demands on the molecular behavior of fluorescent labels. ^[6–10]^ SIM and STED benefit from bright, photostable probes with minimal nonspecific background, while SMLM additionally requires temporally sparse emission events that can be individually localized. ^[11–14]^ These requirements make the regulation of fluorescent and dark states a central challenge in molecular probe design. ^[15–16]^

Rhodamines are particularly attractive because the same chromophore can access a bright, conjugated ring-open state and a dark, typically more membrane-permeable spirocyclic state (Scheme 1a). Target-induced opening provides fluorogenic contrast, whereas reversible opening and closure can produce spontaneous blinking without external chemical switching additives. ^[16–20]^ The intramolecular equilibrium is sensitive to pH, solvent polarity and molecular structure, permitting adaptation to labeling environments and imaging modes. The feasibility of spontaneous spirocyclization-based blinking was established by Urano and coworkers, while later work, including systematic substitution of rhodamine carboxyl derivatives by Lardon and coworkers, demonstrated broad tuning of spirocyclization equilibria and the resulting microscopy performance. Much of the molecular engineering of rhodamine switches has focused on tuning their spirocyclization equilibria through electronic effects, including modulation of the electrophilicity of the *meso*-carbon, the nucleophilicity of the ring-closing group, and the relative stabilization of the open and closed states (Scheme 1b). ^[21–38]^ Such approaches effectively change equilibrium populations, but equilibrium alone does not describe the temporal characteristics of fluorescence switching. Two compounds can exhibit comparable or even oppositely ordered equilibrium preferences while displaying substantially different ON-state residence times. ^[25,36]^ A strategy that controls the structural pathway by which a ring-open molecule returns to its bond-forming geometry could therefore complement electronic tuning.

Steric interactions offer a route to this objective. For a rhodamine spirolactam, opening breaks the bond between the amide nitrogen and the *meso*-carbon, allowing reorientation of the amide-containing substituent. If this rearrangement moves nitrogen away from the carbon reaction center, subsequent ring closure requires a conformational return before C–N bond formation. We hypothesized that steric crowding near the nitrogen could regulate both the preference for the ring-open structure and the accessibility of this return pathway. A bulky substituent may impede return to a bond-forming geometry and prolong the emissive state, whereas a less hindered substituent may permit repeated closure and reopening. This is a mechanistic hypothesis to be evaluated against structural, energetic and switching data, rather than a directly measured elementary reaction sequence.

We envisioned that steric interactions could provide a means to regulate both the thermodynamic and kinetic aspects of rhodamine spirocyclization. In a rhodamine spirolactam, cleavage of the C–N bond upon ring opening allows conformational rearrangement of the amide-containing substituent. Such rearrangement may displace the anionic nitrogen from the cationic xanthene center, requiring a conformational return before C–N bond reformation can occur. We therefore hypothesized that steric crowding near the amide nitrogen could influence the accessibility of this bond-forming geometry. Specifically, a bulky N-substituent could destabilize the closed conformation through steric repulsion with the xanthene plane while promoting an alternative open-state conformation that impedes the re-approach of the reactive centers. This dual effect could favor the fluorescent open state and prolong its residence time. Conversely, less sterically demanding substituents may allow more facile conformational return and ring closure. This proposed mechanism provides a testable framework for understanding how steric structure influences rhodamine switching behavior.

Herein, we investigate this concept by introducing an intramolecular steric engineering strategy to regulate rhodamine switching behavior through modulation of the conformational dynamics associated with ring opening and closure. Specifically, we converted the *ortho*-carboxyl group into an *N*-substituted amide bearing a tert-butyl group, a bulky substituent commonly employed to introduce steric constraints (Scheme 1c). In the spirocyclic state, steric repulsion between the tert-butyl group and the xanthene plane is expected to destabilize the closed conformation, thereby favoring ring opening. Upon C–N bond cleavage, the resulting spirolactam moiety can undergo conformational rearrangement involving rotation of the amide group, potentially relieving steric strain while displacing the anionic nitrogen from the cationic xanthene center. This rearrangement may increase the conformational barrier to C–N bond reformation, thereby suppressing ring closure and prolonging the lifetime of the fluorescent open state. In contrast, less sterically demanding amide substituents are expected to allow the reactive centers to remain in closer proximity following ring opening, facilitating conformational return and subsequent C–N bond reformation. Thus, steric engineering may exert a dual effect on rhodamine switching by destabilizing the closed state and restricting access to the bond-forming geometry in open state, providing a means to modulate both spirocyclization equilibrium and switching kinetics. The resulting shift toward the fluorescent open state also offers opportunities for fluorogenic labeling and SMLM imaging. To explore these possibilities, we further incorporated HaloTag ligands into the molecular framework and developed fluorogenic probes for wash-free super-resolution imaging in living cells. Moreover, the prolonged ON-state lifetime enabled dual-modal super-resolution imaging using a single fluorophore.

**Scheme 1.**
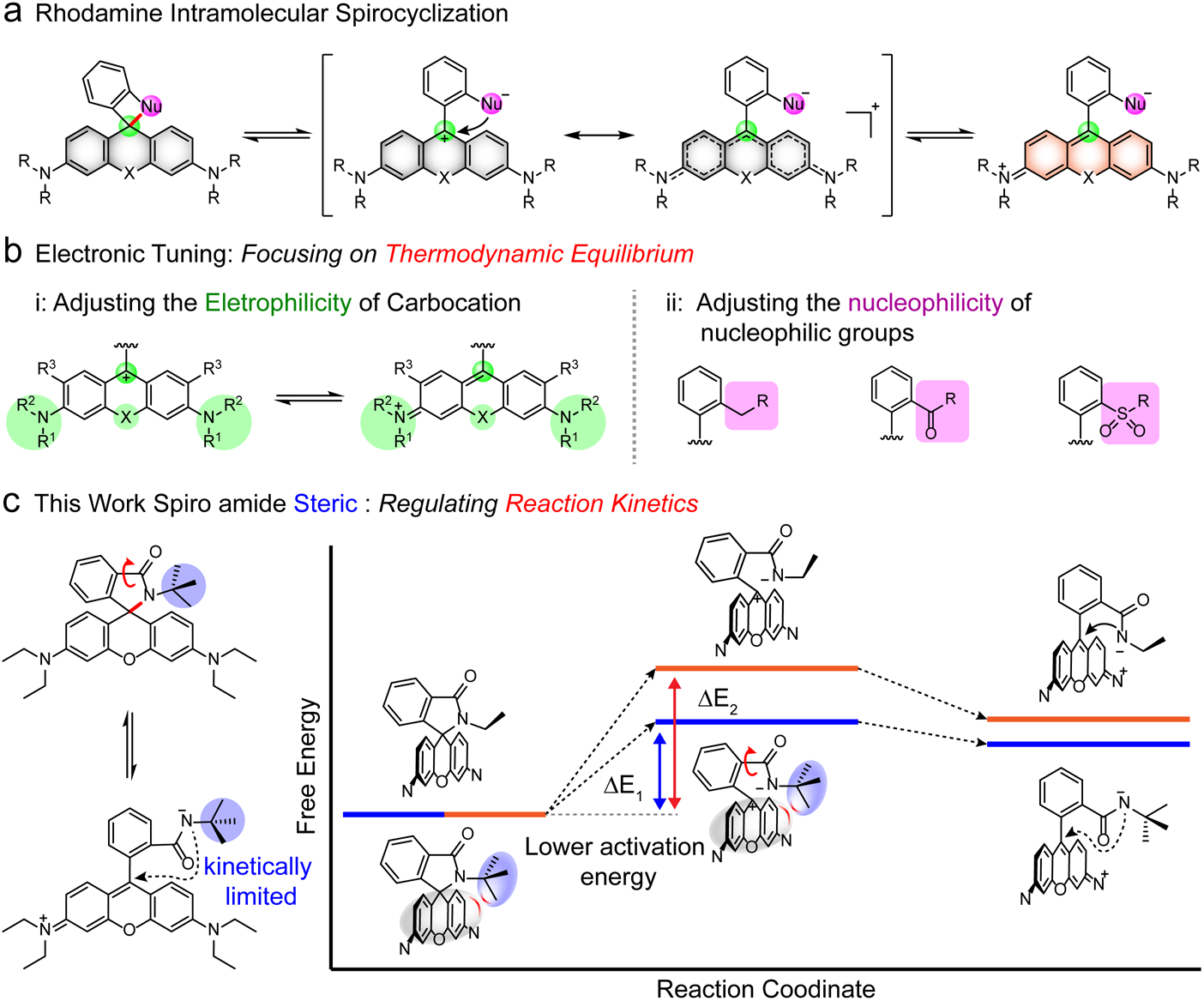
(a) Intramolecular reversible spirolactamization of rhodamine derivatives. (b) Current strategies for regulating the thermodynamic equilibrium. (c) Kinetic regulation through steric hindrance near the spirocyclic center.

## Results and discussion

### Molecular Design and Synthesis

To perturb the local environment of the reversible C–N bond without extensively changing the xanthene chromophore, we transformed the ortho-carboxyl functionality of rhodamine derivatives into N-substituted spirolactams. The initial series **1a**–**1f** spans substituents with different spatial profiles, including a tert-butyl-derived motif. Our working model predicts that a substituent sufficiently close to the xanthene plane creates repulsion in the closed state, while the corresponding open form can relax through amide reorientation. The magnitude of this effect should depend on the actual three-dimensional placement of the substituent rather than substituent size alone.

### Steric regulation of the rhodamine spirocyclization equilibrium

We first evaluated the p*K*_cycl_ values of compounds **1a**–**1e** in PBS buffer and their corresponding D_50_ values in a water–dioxane system to investigate the influence of steric hindrance on the rhodamine spirocyclization equilibrium. Here, p*K*_cycl_ and D_50_ represent the pH and solvent dielectric constant, respectively, at which 50% of the fluorophore population exists in the ring-open form, providing quantitative measures of the equilibrium between the open and closed states under different environmental conditions. Compounds **1a**-**1d** exhibited no apparent changes in p*K*_cycl_ as the steric substituents were varied from ethyl to cyclohexyl groups. In contrast, 1e showed a pronounced shift toward the ring-open form, with a p*K*_cycl_ value of 4.9 (Figure 1a, b and Figure S1a-f). Consistent with this observation, D_50_ measurements revealed a similar trend (Figure 1a, c and Figure S2a–f). Whereas compounds **1a**–**1d** showed no substantial changes in D_50_, compound 1e exhibited a marked shift, reaching a D_50_ value of 50.

**Figure 1.**
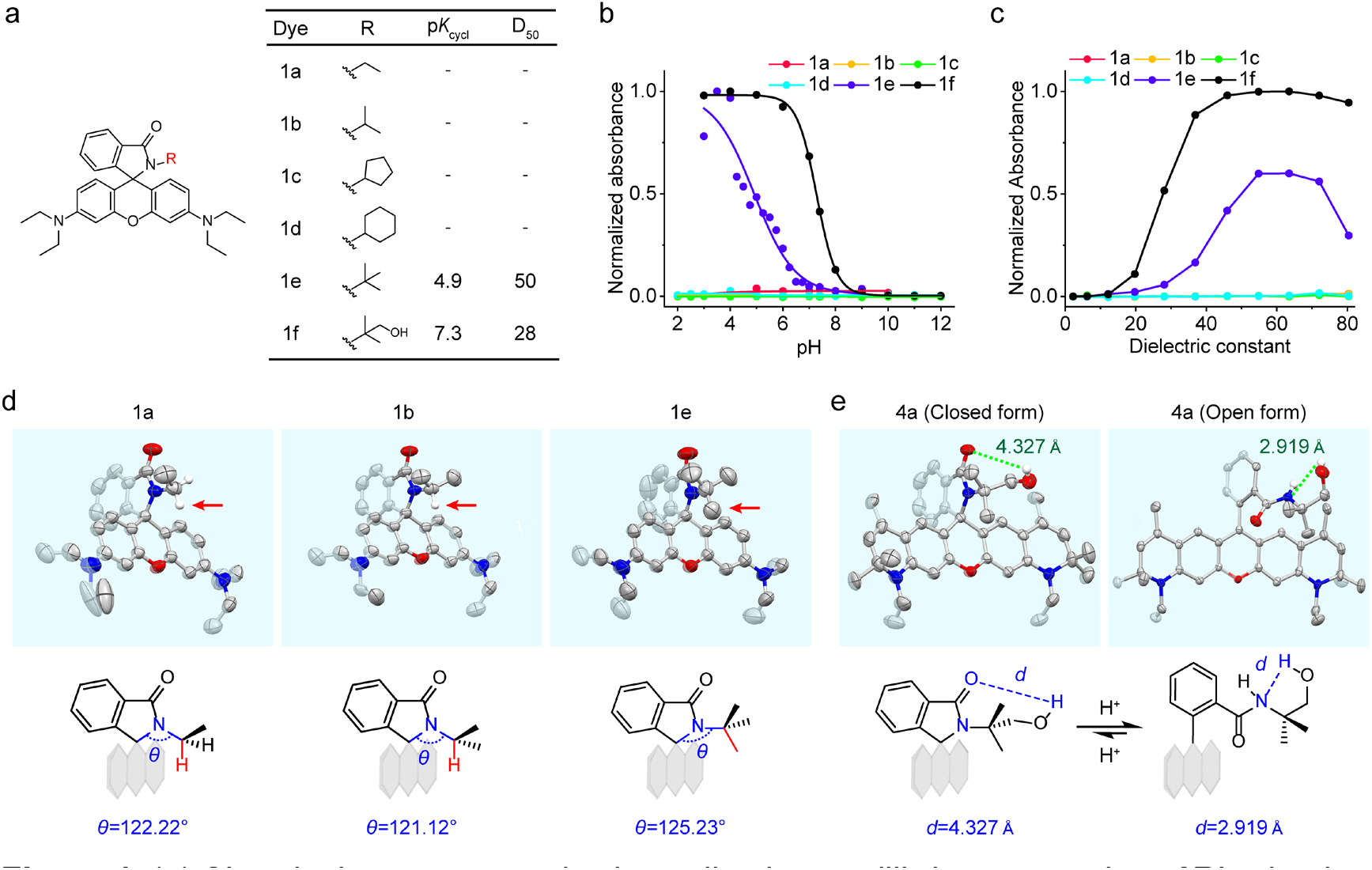
(a) Chemical structures and spirocyclization equilibrium properties of Rhodamine B derivatives **1a-1f**. (b) Normalized maximum absorbance of **1a-1f** at room temperature as a function of pH. (c) Normalized maximum absorbance of **1a-1f** at room temperature as a function of the dielectric constant. (d) Closed-form crystal structures of **1a**, **1b, 1e.** (e) Closed- and open-form crystal structures of **4a.**

Notably, increasing the electron-donating character of N-substituent would generally be expected to enhance the electron density and nucleophilicity of the amide nitrogen, potentially favoring intramolecular nucleophilic addition and ring closure. The pronounced preference for the ring-open state observed for **1e** therefore cannot be readily explained by electronic effects alone. Instead, these results suggest that the steric repulsion introduced by the tert-butyl group plays a dominant role in destabilizing the closed conformation and shifting the spirocyclization equilibrium toward the fluorescent open state. Interestingly, compound **1f** exhibited an even stronger preference for the ring-open state than **1e**. We speculate that the additional hydroxyl group on its tert-butyl substituent may form an intramolecular hydrogen bond with the anionic nitrogen generated upon ring opening, providing additional stabilization of the open conformation and further shifting the equilibrium toward the fluorescent state.

Interestingly, the steric effect was distinctly non-monotonic, with no progressive shift toward the ring-open state as substituent bulk increased. This suggests that steric regulation depends not simply on the nominal volume of the N-substituent but also on its spatial orientation relative to the xanthene framework, which determines whether effective steric crowding can be established. To further investigate this effect and determine whether the observed trend was specific to the rhodamine scaffold, we synthesized an analogous series of Rho 6G derivatives (**2a**–**2f**). A similar trend was observed (Figures S1g–l, S2g–l, and S3a,b; Scheme S1): compounds **2a**–**2d** exhibited comparable p*K*_cycl_ values of approximately 3.3, with D_50_ values beyond the experimentally accessible range, whereas **2e** showed a pronounced shift toward the ring-open state (p*K*_cycl_ = 7.4, D_50_ > 60). The consistent behavior across both dye scaffolds supports the existence of a steric threshold, highlighting the importance of substituent spatial arrangement rather than size alone.

We next explored the applicability of this strategy to other rhodamine derivatives by synthesizing compounds 3a and 4a (Scheme S1). Their p*K*_cycl_ values, together with those of **1f** and **2f** (6.7, 5.8, 7.0, and 8.2, respectively), revealed considerable differences in the ring-opening equilibrium among these derivatives (Figures S1m, n and S3c; Scheme S1), indicating that the molecular scaffold and its electronic structure also contribute to the equilibrium position. Their D_50_ values also consistent with the above trend (Figures S2m, n, and S3d; Scheme S1). To further examine the role of scaffold structure, we introduced a tert-butyl group into the Si-rhodamine scaffold to obtain compound 21 (Figure S1o). However, neither its absorption nor fluorescence spectra showed appreciable pH-dependent changes, indicating that tert-butyl group alone was insufficient to induce effective spirocyclization switching in this scaffold. These results indicate that steric engineering can modulate the spirocyclization equilibrium across different rhodamine derivatives, but its effectiveness is strongly dependent on the molecular scaffold and electronic environment.

### Crystallographic evidence for steric repulsion and post-opening rearrangement

To elucidate the structural origin of steric threshold, we analyzed the single-crystal structures of **1a**, **1b**, and **1e** in their closed forms (Figure 1d and Table S1, deposition numbers 2603541, 2603545 and 2603546). Although the substituents differ in overall size, their spatial arrangements revealed a key distinction that a methyl group in 1e, but not in 1a or 1b, is positioned above the xanthene framework. This local steric interaction is accompanied by an increase in the corresponding dihedral angle (θ) from 122.22° and 121.12° in 1a and 1b, respectively, to 125.23° in 1e (Figure 1d). This structural distortion is consistent with enhanced steric crowding between the substituent and xanthene plane, providing a structural explanation for the pronounced shift in the spirocyclization equilibrium observed for 1e.

To further investigate the conformational changes associated with ring opening, we compared the closed- and open-form crystal structures of **4a** (Figure 1e and Table S1, deposition numbers 2603448 and 2603548). Ring opening is accompanied by torsional reorientation of the amide-containing group, resulting in increased spatial separation between the nitrogen and meso-carbon reaction centers. This observation suggests a conformational-gating mechanism in which ring closure requires the molecule to return to a geometry favorable for C–N bond formation. Steric crowding may hinder this conformational return and reduce access to the ring-closing geometry, whereas less bulky substituents may allow more facile access to the reactive conformation. Such differences in conformational accessibility could contribute to variations in the ON-state lifetime. Although the crystal structures capture distinct conformations rather than their interconversion dynamics, they provide a structural basis for this proposed mechanism.

The crystal structures of **4a** further provide a possible explanation for the enhanced ring-opening tendency of hydroxyl-containing compound **1f**. In the closed form, the hydroxymethyl group points away from the carbonyl group, with a distance of 4.237 Å between the hydroxyl hydrogen and carbonyl oxygen. Upon ring opening, conformational rearrangement brings the hydroxyl group closer to the amide nitrogen, reducing the H···N distance to 2.919 Å (Figure 1e). This geometry is compatible with a possible intramolecular hydrogen bond, which may stabilize the anionic nitrogen in the open form and further favor ring opening. These observations suggest that introducing additional functional groups onto the steric substituent provides a means to fine-tune the spirocyclization equilibrium through secondary intramolecular interactions.

### Theoretical Calculations Corroborate the Role of Steric Hindrance in Promoting Rhodamine Ring Opening

To gain further insight into the effect of steric hindrance on the ring-opening process, we performed theoretical calculations on compounds **1a** and **1e**, representing systems with relatively low and high steric hindrance, respectively. The ring-opening pathways were investigated by modeling three key states, including closed lactam, transition state, and zwitterionic intermediate. During ring opening, the distance between the carbon and nitrogen atoms involved in spirocyclization progressively increases (Figure 2a). Notably, 1e exhibited a lower calculated activation energy for ring opening than 1a (1.12 *vs* 1.37 eV). Moreover, the zwitterionic intermediate of 1e exhibited a lower relative energy than that of 1a, with the energy of each closed lactam set as the reference. These results indicate that the bulky substituent in 1e facilitates ring opening by both lowering the activation barrier and energetically favoring the open form relative to the closed form.

**Figure 2.**
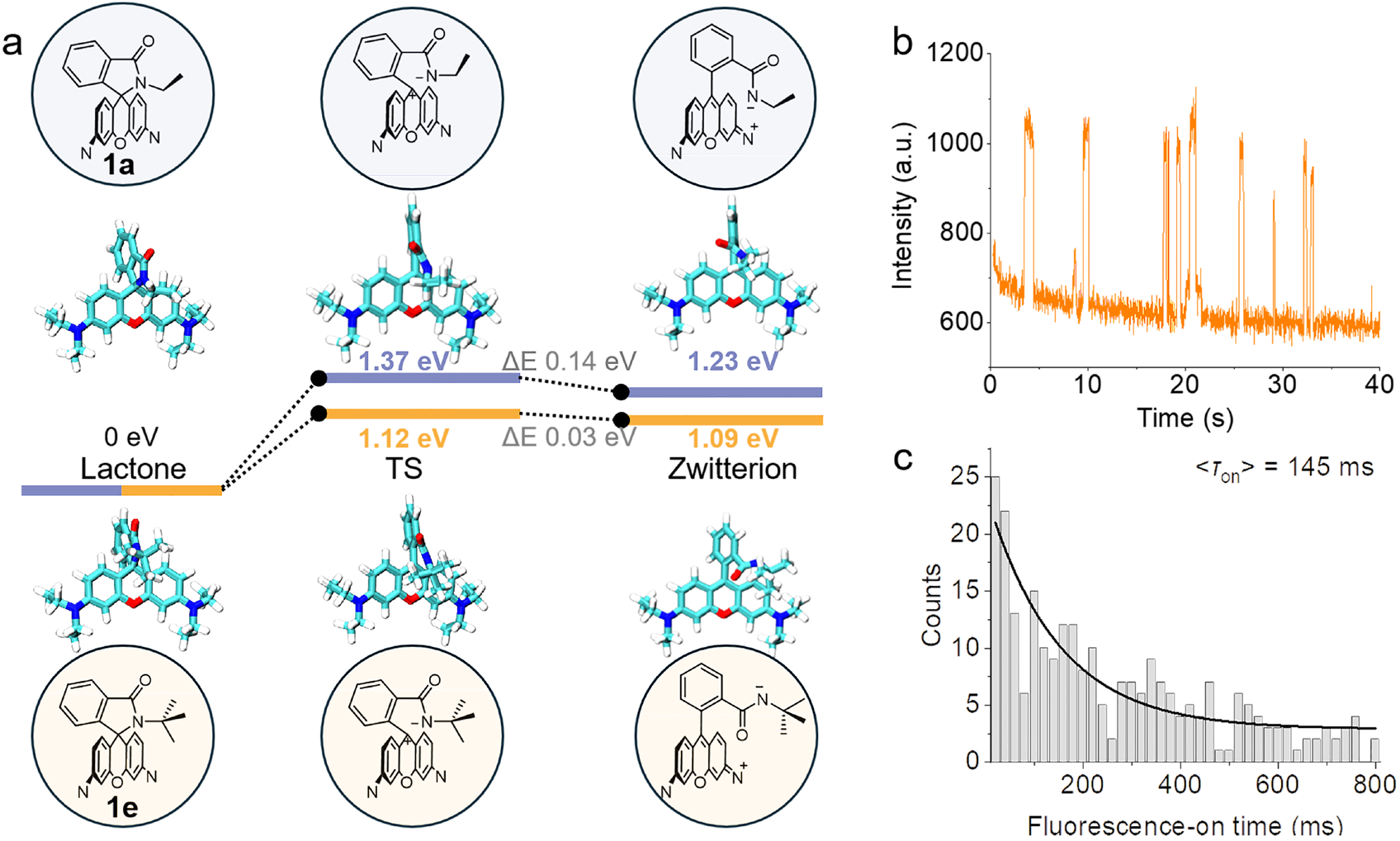
(a) Theoretical energy profiles for the ring-opening processes of **1a** and **1e** in water. (b) Representative single-molecule fluorescence trace of **1e**. (c) Distribution of fluorescence-on time of **1e**.

The calculated energy profiles can be rationalized by considering the steric interactions in the closed and open forms. In the closed lactam conformation, the bulky -R group experiences steric repulsion with the xanthene scaffold, which destabilizes the closed state. Upon ring opening, the increased conformational freedom around the amide-containing substituent may allow partial relief of this steric repulsion. This relief could contribute to the relative stabilization of the transition state and zwitterionic intermediate. Thus, these calculations suggest that steric hindrance promotes ring opening by lowering the activation barrier and reducing the relative energy difference between the closed and open forms. The calculated energetic preference for the open form is also consistent with the higher p*K*_cycl_ value of **1e**, given that a higher p*K*_cycl_ corresponds to a greater population of ring-open form under the same conditions.

These experimental and computational results indicate that the tert-butyl group effectively modulates the spirocyclization equilibrium of rhodamine switches by introducing steric repulsion in the closed conformation and facilitating ring opening. We also reasoned that the steric hindrance might impede the conformational return required for ring closure, potentially prolonging the lifetime of open state while preserving spontaneous switching behavior. To examine this possibility, we next characterized the single-molecule switching behavior of **1e** by depositing nanomolar solutions onto hydroxylated coverslips to achieve molecular dispersion. Compound **1e** exhibited reversible ON/OFF switching at pH 7.4 (Figure 2b), demonstrating that the introduction of steric hindrance preserves the spontaneous blinking behavior of rhodamine fluorophores. We further quantified its ON-state lifetime (*τ*_on_) and compared it with that of our previously reported compound HM-DS531.^[39]^ Compound **1e** exhibited a p*K*_cycl_ value of 4.9 and a *τ*_on_ of 145 ms (Figure 2c), whereas HM-DS531 exhibited a p*K*_cycl_ value of 5.3 and a shorter *τ*_on_ of 50 ms. The lower p*K*cycl value of **1e** indicates a less favorable thermodynamic equilibrium for the ring-open form than that of **HM-DS531**. Nevertheless, **1e** exhibited a substantially prolonged *τ*_on_. This apparent discrepancy highlights that the thermodynamic preference for the ring-open state does not necessarily correlate with its kinetic persistence. The prolonged *τ*_on_ of **1e** is consistent with steric hindrance impeding the conformational return required for ring closure, thereby kinetically extending the lifetime of the fluorescent open state.

### Conversion into HaloTag-compatible fluorogenic and blinking probes

HaloTag is a widely used protein-labeling system in which fluorophores functionalized with a chloroalkane ligand covalently bind to HaloTag-fused target proteins, enabling selective labeling of proteins of interest.^[40]^ For wash-free imaging, fluorogenic probes are particularly desirable because they exhibit low fluorescence before binding to the target protein, thereby minimizing background signals.^[41]^ In addition, Halo-Tag-based probes with suitable photophysical properties are highly valuable for super-resolution imaging, particularly SMLM imaging.^[28]^

We investigated whether our steric regulation strategy could be applied to the development of HaloTag-based probes for fluorogenic and super-resolution imaging. For fluorogenic imaging, probes should remain predominantly nonfluorescent before binding to HaloTag. For SMLM imaging, based on our preceding results, we selected compounds with p*K*_cycl_ values below 6 as candidates, aiming to retain a sufficiently large population of the nonfluorescent closed form under physiological conditions while enabling spontaneous fluorescence switching. Accordingly, compounds **1e**, **1f**, **3a**, and **4a** were selected as starting scaffolds for probe development. We first investigated whether the linker attachment position and bonding mode would affect the spirocyclization equilibrium. To this end, we synthesized compounds **5,6-halo-RhB**, **O-halo-RhB**, and **N-halo-RhB** (Figure 3) and measured their p*K*_cycl_ values (Figure S4). Compounds **5,6-halo-RhB** and **O-halo-RhB** exhibited similar p*K*_cycl_ values of 3.7 and 3.6, respectively, whereas **N-halo-RB** showed a higher p*K*_cycl_ value of 5.9. These results suggest that the attachment position has a relatively minor influence on the spirocyclization equilibrium, whereas the amide linkage can shift the equilibrium toward the ring-open form. We speculate that this effect may be associated with intramolecular hydrogen bonding, analogous to the interaction suggested by the crystal structure of **4a** (Scheme S2). Based on these findings, we introduced the HaloTag linker through an ester bond into **3a** and an amide bond into **4a** (Figure 3), yielding **O-halo-JF549** (p*K*_cycl_ = 4.0) and **N-halo-JA21** (p*K*_cycl_ = 2.9), respectively. Both compounds met the p*K*_cycl_ criterion established for our SMLM probe design (Figure S4).

**Figure 3.**
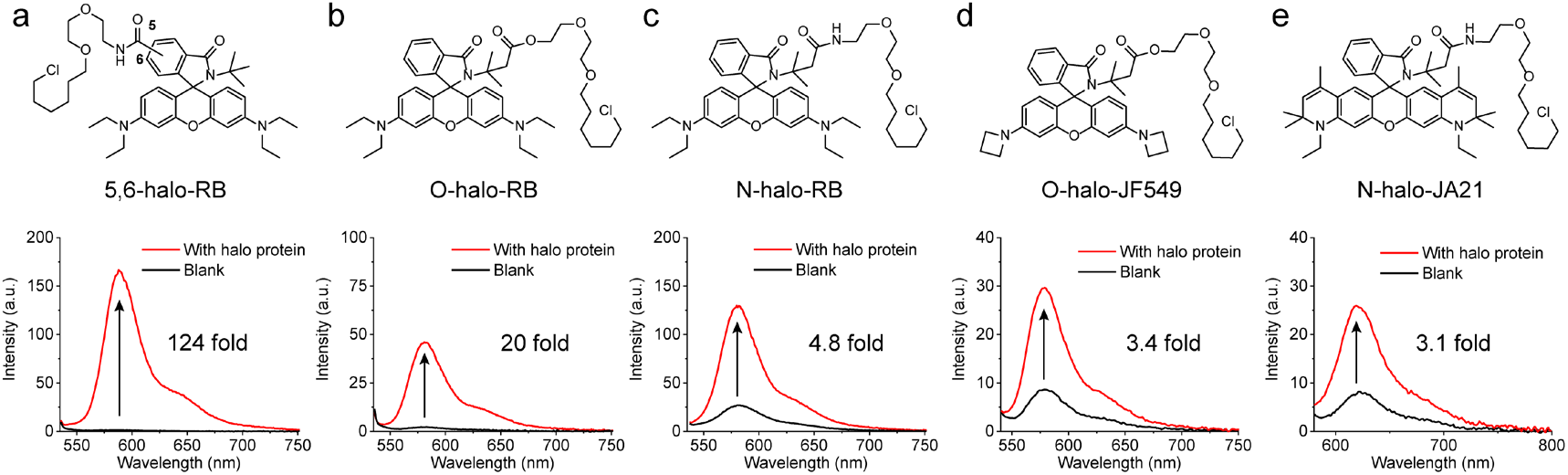
Chemical structures of **5,6-halo-RB** (a), **O-halo-RB** (b), **N-halo-RB** (c), **O-halo-JF549** (d) **and N-halo-JA21** (e) and their fluorescence spectra before and after reaction with HaloTag protein.

We next evaluated the fluorogenic responses of these probes upon reaction with HaloTag protein (Figure 3). After 2 h of incubation, **5,6-halo-RB** exhibited a 124-fold increase in fluorescence intensity, while **O-halo-RB** showed a 20-fold enhancement. In contrast, **N-halo-RB**, **O-halo-JF549**, and **N-halo-JA21** exhibited substantially smaller fluorescence enhancements of 4.8-, 3.4-, and 3.1-fold, respectively. These results demonstrate that both the linker attachment position and bonding mode influence the fluorogenic response upon HaloTag labeling. In particular, introducing the HaloTag linker at the 5- or 6-position of the rhodamine scaffold appears more favorable for developing fluorogenic probes. We further investigated the labeling kinetics by monitoring the fluorescence enhancement over time (Figure S5). The reactions of **N-halo-RB** and **O-halo-JF549** were completed within 30 min, whereas the other probes showed continued increases in fluorescence over the measurement period. Considering both the fluorescence enhancement and labeling kinetics, we selected **5,6-halo-RB**, **O-halo-RB**, and **N-halo-RB** for further fluorogenic imaging studies.

In parallel, we investigated whether the HaloTag linker affected the spontaneous switching behavior required for SMLM imaging. Single-molecule fluorescence measurements of all five probes (**5,6-halo-RB**, **O-halo-RB**, and **N-halo-RB**, **O-halo-JF549**, and **N-halo-JA21**) on PMMA films revealed repetitive ON/OFF switching (Figure S6), indicating that linker functionalization preserved their spontaneous blinking behavior. These results support the potential of all five probes for SMLM imaging. Prior to live-cell super-resolution imaging, we further evaluated their cytotoxicity in HeLa cells. Following 24 h of incubation at 5 μM, cell viability remained above 90% for all five probes, indicating good cytocompatibility under the tested conditions (Figure S7).

### Wash-free fluorogenic imaging and long-term live-cell SIM and STED imaging

We further evaluated the fluorogenic properties of **5,6-halo-RB**, **O-halo-RB**, and **N-halo-RB** through live-cell, wash-free imaging of HeLa cells expressing an H2B-Halo fusion protein localized to the nucleus (Figure 4a and Figure S8). The signal-to-background ratios were assessed by comparing the nuclear fluorescence intensity (F_nuc_) with the cytosolic background intensity (F_cyt_) (Figure 4b). **5,6-halo-RB** and **N-halo-RB** exhibited comparable nuclear-to-cytosolic fluorescence ratios, with F_nuc_ */* F_cyt_ values of 4.1 and 4.8, respectively. However, **N-halo-RB** also produced substantial background fluorescence in lysosomes, potentially limiting its performance in fluorogenic imaging. In contrast, **O-halo-RB** showed no detectable fluorescence under the same imaging conditions (Figure S8. To determine whether this result was caused by unsuccessful transfection, we further examined these probes in HeLa cells expressing a Halo-actin-EGFP fusion protein (Figure S9). Successful transfection could be verified by detecting EGFP fluorescence in the 488 nm channel. Both **5,6-halo-RB** and **N-halo-RB** selectively labeled transfected cells and revealed the actin structures. However, **O-halo-RB** again showed no detectable fluorescence in any of the observed cells, despite the presence of EGFP fluorescence confirming successful transfection. We therefore speculate that the fraction of the fluorescent open form of **O-halo-RB** following HaloTag labeling may be insufficient to generate a detectable signal under confocal imaging conditions. Such a low fluorescence background could potentially be advantageous for SMLM, where sparse fluorescence emission is desirable.

**Figure 4.**
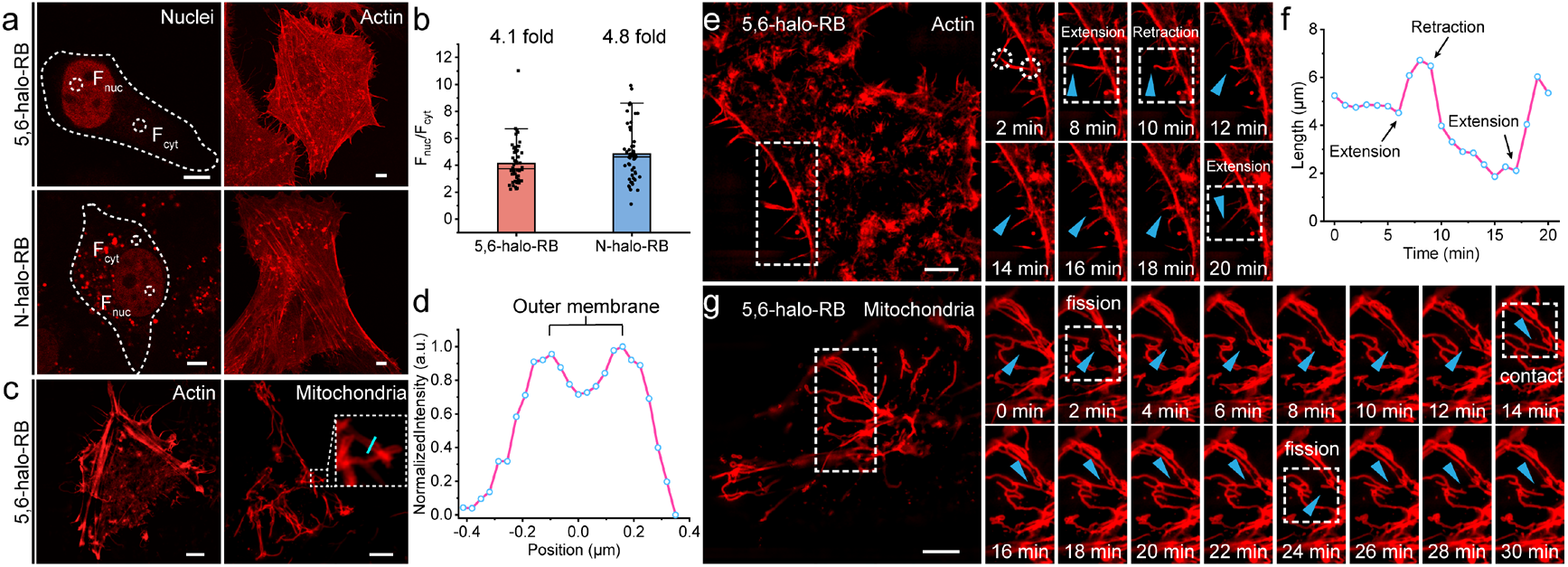
Live-cell fluorogenic imaging. (a) Live-cell, no-wash confocal images of HeLa cells expressing H2B-Halo or Halo-Actin-EGFP fusion proteins labeled with **5,6-halo-RB** and **N-halo-RB** (400 nM). (b) Fluorescence ratio of **5,6-halo-RB** and **N-halo-RB** in live cell, no wash confocal microscopy. Bar plot shows the nuclear signal (F_nuc_) and cytosolic signal (F_cyt_) ratio in transfected cells. Error bars show ± SD. (c) Live cell, no wash SIM images of HeLa cells expressing Halo-LifeAct fusion protein and HeLa cells expressing tom20-halo-EGFP fusion protein labeled with **5,6-halo-RB** (400 nM). (d) Normalized intensity profiles across the cyan line in (c). (e) Long-term SIM imaging of pseudopodial dynamics. (f) Length of the selected pseudopodium in (e) as a function of time. (g) Long-term SIM imaging of mitochondrial dynamics. Scale bar: 5 μm.

Encouraged by the fluorogenic performance of **5,6-halo-RB**, we next applied it to SIM imaging of live HeLa cells expressing H2B-Halo fusion proteins localized to the nucleus (Figure S10a). Intensity analysis along the white line revealed an approximately eightfold enhancement in nuclear fluorescence (Figure S10b). We further explored its capability for resolving finer cellular structures by performing SIM imaging of pseudopodia and mitochondrial outer membranes in living cells. The reconstructed images clearly resolved actin-rich pseudopodial structures and mitochondrial membranes (Figure 4c and Figure S10c). Notably, the fluorescence intensity profile across an individual mitochondrion displayed two distinct peaks, consistent with signals arising from the two sides of its outer membrane. This result indicates that **5,6-halo-RB** enables the visualization of mitochondrial membrane structures at the subcellular level (Figure 4d and Figure S10d–h).

We further investigated the long-term imaging performance of **5,6-halo-RB** by monitoring the dynamic changes in pseudopodia and mitochondria (Figure 4e-g). Images were acquired with an exposure time of 200 ms. The length of the selected pseudopodium remained nearly unchanged during the first 6 min. It then increased abruptly, followed by gradual shortening over the next 9 min and subsequent recovery over 3 min (Figure 4e, f). During the 30 min observation of mitochondrial dynamics, the mitochondrion within the white dashed box in Figure 4g underwent successive morphological changes involving fission, contact with another mitochondrial segment, and a subsequent fission event. These results demonstrate that steric regulation of the spirocyclization equilibrium enables the development of fluorogenic HaloTag probes for live-cell imaging. In particular, introducing the HaloTag linker at the 5- or 6-position of the rhodamine scaffold appears favorable for achieving fluorogenicity, while **5,6-halo-RB** also supports SIM imaging of fine cellular structures and their dynamic changes.

Following the successful SIM imaging, we further explored the applicability of **5,6-halo-RB** to STED microscopy. We first used this probe to visualize the outer mitochondrial membrane in HeLa cells expressing Halo-Tom20, obtaining images that clearly resolved mitochondrial membrane structures (Figure 5a). A resolution of 162 nm was achieved for the mitochondrial outer membrane. We also successfully applied **5,6-halo-RB** to image actin filaments in cells expressing Halo-LifeAct. Two intersecting actin filaments that could not be clearly distinguished by confocal microscopy were readily resolved by STED microscopy. The achieved spatial resolution was 132 nm, enabling the discrimination of two actin filaments separated by 274 nm (Figure 5b). The dynamic remodeling of actin structures was also captured under STED microscopy (Figure 5c). Three filopodia located on the same lamellipodium, indicated by cyan arrows, gradually approached one another after imaging commenced and eventually merged into a single filopodium. Meanwhile, the filopodium indicated by the green arrow gradually detached from the upper lamellipodium and moved toward the three filopodia marked by cyan arrows, eventually merging with them. In addition, two groups of lamellipodia observed at the beginning of imaging gradually contracted and merged, ultimately giving rise to the two filopodia indicated by the magenta and cyan arrows (Figure 5c). Collectively, these results demonstrate that steric engineering of rhodamine spirocyclization provides an effective strategy for developing fluorogenic HaloTag probes for live-cell super-resolution imaging. In particular, introducing the HaloTag linker at the 5- or 6-position of the rhodamine scaffold appears favorable for achieving a pronounced fluorogenic response. The resulting probe, **5,6-halo-RB**, supports both SIM and STED imaging of fine cellular structures and enables the visualization of their dynamic remodeling over extended periods in living cells.

**Figure 5.**
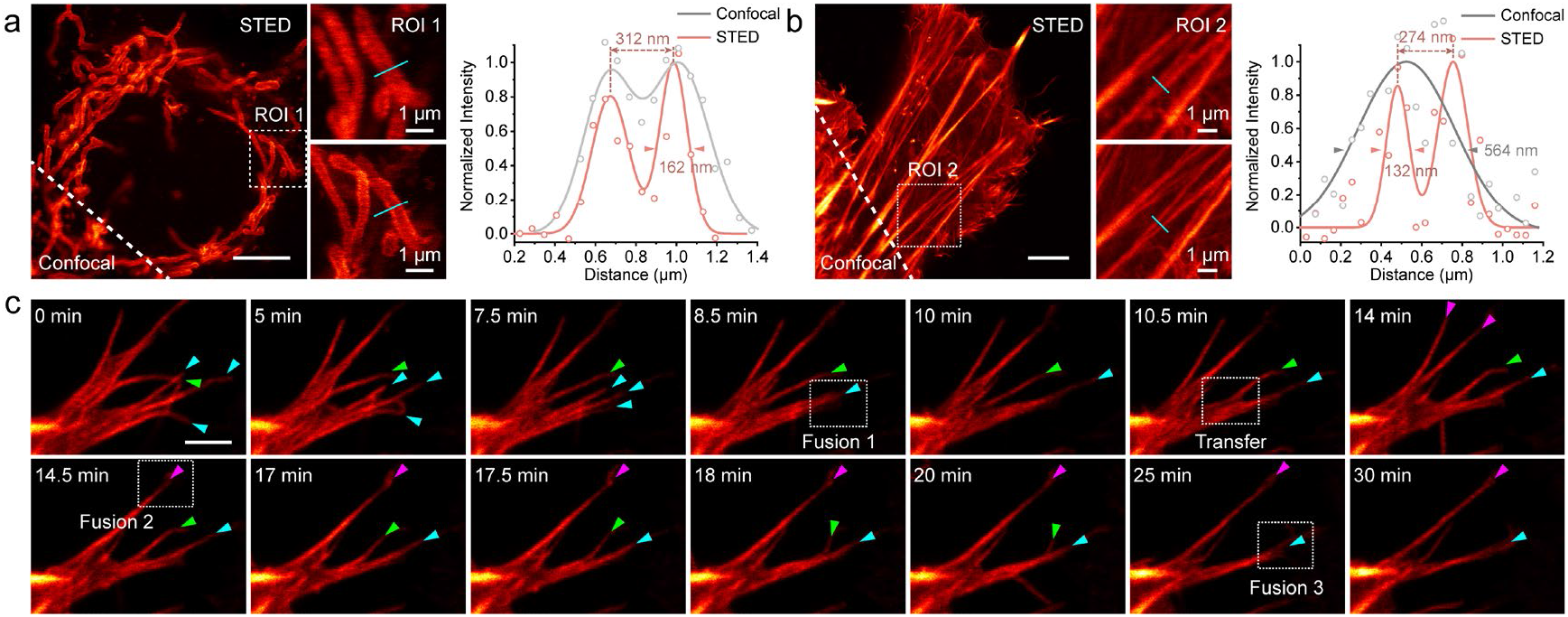
Confocal and STED imaging of the mitochondrial outer membrane (a) and actin filaments (b) in live HeLa cells using **5,6-halo-RB** (400 nM). Right panels: Fluorescence intensity profiles along the cyan lines within the indicated ROIs. (c) STED imaging of contraction and fusion of lamellipodia during 30 min labeled with **5,6-halo-RB** (400 nM). The HeLa cells were transfected with halo-tom20 and Halo-LifeAct fusion protein respectively. Scale bar: 5 μm.

### Optimizing the steric regulated switches for SMLM

Having demonstrated the spontaneous blinking behavior of **5,6-halo-RB**, **O-halo-RB**, **N-halo-RB**, **O-halo-JF549**, and **N-halo-JA21** on PMMA films, we next evaluated their performance in live-cell SMLM. HeLa cells expressing an H2B-Halo fusion protein localized to the nucleus were labeled with these probes and imaged under no-wash conditions. Although **5,6-halo-RB** exhibited pronounced fluorogenicity in conventional live-cell imaging, no detectable blinking was observed under SMLM imaging conditions employed. In contrast, **O-halo-RB**, which showed no detectable fluorescence in conventional fluorogenic imaging, enabled successful SMLM imaging, with an average localization precision of 24.6 nm and an average photon count of 856 (Figure 6a–c). Compound **N-halo-RB** also exhibited clear blinking behavior, with an average localization precision of 20.4 nm and an average photon count of 1,046 (Figure 6d–f). Similarly, **O-halo-JF549** displayed spontaneous blinking properties, yielding an average localization precision of 19.2 nm and an average photon count of 1,008 (Figure 6g–i). However, **N-halo-JA21** failed to produce detectable fluorescence from HaloTag-labeled proteins in live cells. We speculate that its relatively slow reaction kinetics, together with its high lipophilicity, may reduce its effective intracellular availability for productive HaloTag labeling under the conditions employed.

**Figure 6.**
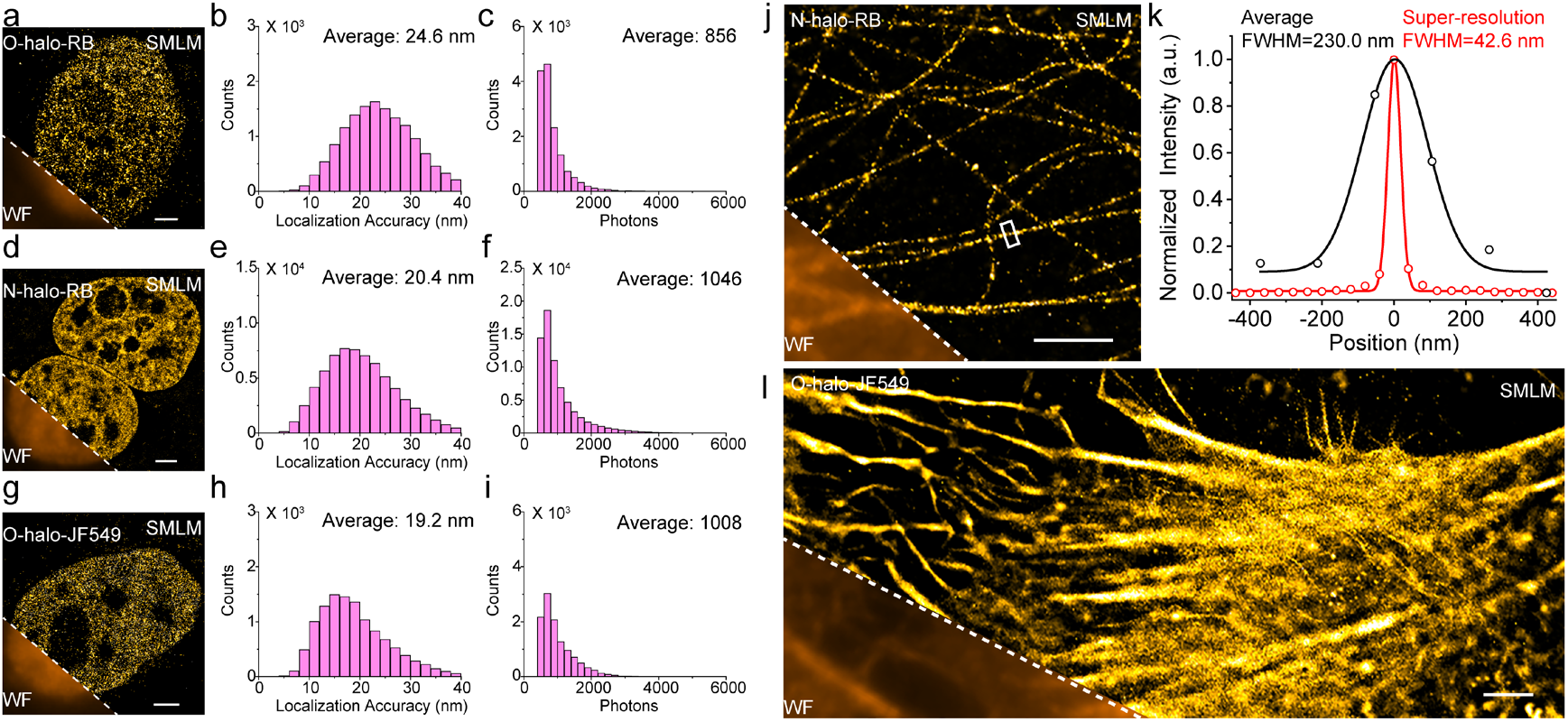
Live cell SMLM imaging of HeLa cells expressing H2B-halo fusion protein labeled with **O-halo-RB** (a), **N-halo-RB** (d) and **O-halo-JF549** (g) for 1 h (500 nM) respectively. Lower left: Wide-field imaging. (b), (e) and (h) Histograms of localization precision. (c), (f) and (i) Histograms of photon counts. (j) SMLM imaging of fixed HeLa cells expressing halo-tubulin fusion protein labeled with **N-halo-RB** (500 nM) for 1 h. Lower left: Wide-field imaging. (k) Wide-field and SMLM imaging intensity distributions in the white box in (j). (l) Live cell SMLM imaging of HeLa cells expressing LifeAct-halo fusion protein labeled with **O-halo-JF549** for 1 h. Lower left: Wide-field imaging. The cut off value was 200 photons. Scale bar: 2 μm.

We further evaluated the performance of **N-halo-RB** for microtubule imaging in fixed HeLa cells expressing a Halo-tubulin fusion protein (Figure 6j). The resulting SMLM images resolved fine microtubule structures, with the narrowest measured full width at half maximum (FWHM) reaching 42.6 nm, comparable to previously reported values (Figure 6k).^[21]^ We also used **O-halo-JF549** to visualize actin structures in living cells (Figure 6l). Two adjacent actin structures that could not be clearly distinguished by conventional imaging were readily resolved by SMLM (Figure S11).

Given the previously demonstrated brightness and photostability of nitrocyclobutane-substituted rhodamine derivatives, together with the superior localization precision of **O-halo-JF549** compared with **O-halo-RB** and **N-halo-RB**, we selected **O-halo-JF549** for time-lapse SMLM imaging in living cells. Actin structures were imaged with an exposure time of 20 ms, and super-resolution images were reconstructed from every 1,000 consecutive frames (frames 1–1000, 1001–2000, 2001–3000, and so forth). This approach enabled us to track the movement and retraction of cell pseudopodia over time. The retraction of a selected pseudopodium was fitted with an exponential decay function, Len = 165.52e^(-t/50.49)^ + 5.31 (Figure 7a,b and Figure S12a,b). Importantly, the localization precision, photon counts, and number of detected molecules remained relatively stable throughout the imaging process, indicating that **O-halo-JF549** maintained reliable imaging performance during time-lapse SMLM (Figure S12c-e). In addition to pseudopodial dynamics, we also monitored changes in the cell contour (Figure 7c). The area within the magenta box gradually decreased over time, suggesting a progressive reduction in the contact region between the target cell and its neighboring cell (Figure 7d). These results establish **O-halo-JF549** as a promising probe for time-lapse SMLM of dynamic cellular structures in living cells and further demonstrate the potential of sterically engineered rhodamine switches for live-cell super-resolution imaging.

**Figure 7.**
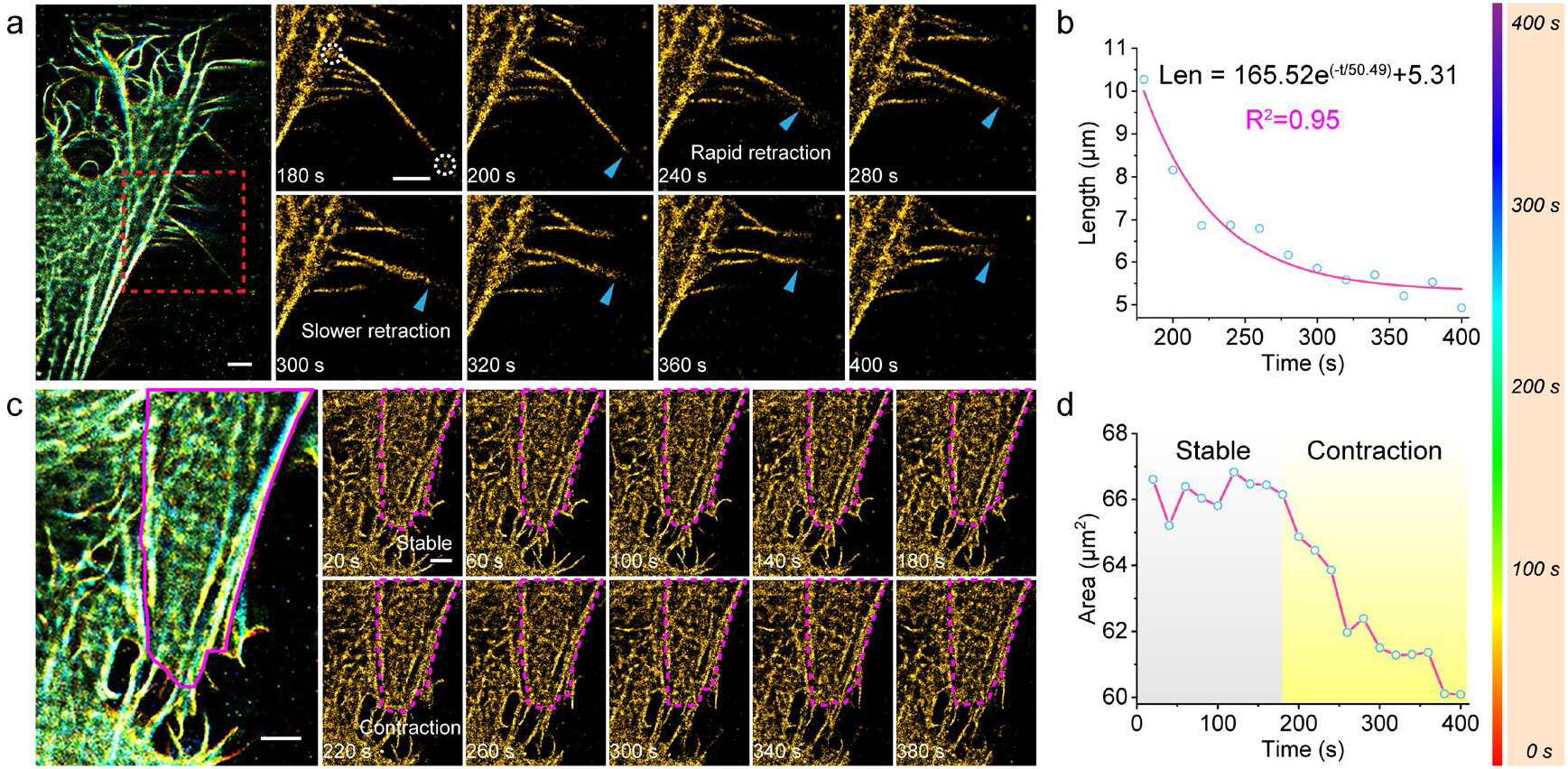
Time-lapse SMLM imaging of actin dynamics in living cells labeled with **O-halo-JF549**. (a) Overlaid time-lapse images and long-term SMLM imaging of the region of interest of a selected pseudopodium region. (b) Length of the selected pseudopodium in (a) as a function of time. (c) Overlaid time-lapse images and long-term SMLM imaging of cellular movement. The magenta box shows the cell contours at different times. (d) Area of the region of interest in (c) as a function of time. Scale bar: 2 μm.

### Optimizing sterically regulated switches for dual-modal imaging with a single probe

Although SMLM provides single-molecule-level spatial information, its reliance on the sequential localization of individual fluorophores generally requires prolonged acquisition to reconstruct complete structural features. Consequently, when imaging is restricted to a short time window, particularly for rapidly evolving biological processes, the limited number of localization events may be insufficient to recover the overall structural information (Figure 8a). In contrast, SIM enables rapid reconstruction of entire cellular structures, although it does not provide single-molecule-level information. Thus, combining these complementary imaging modalities offers the opportunity to simultaneously visualize overall structural organization and the underlying distribution of single-molecule events. Importantly, implementing both modalities with a single fluorophore would further eliminate the need for separate fluorescent labels and imaging channels, preserving additional spectral channels for multicolor imaging of other cellular components. However, this strategy presents a considerable photophysical challenge because the two techniques impose distinct, and potentially conflicting, requirements on fluorophore emission behavior. SIM requires a sufficiently large population of simultaneously emitting fluorophores to generate high-contrast structured-illumination patterns, whereas SMLM relies on sparse and temporally separated emission events to enable precise single-molecule localization. Accordingly, continuously emitting fluorophores are generally unsuitable for SMLM, while spontaneously blinking dyes may provide insufficient fluorescence for SIM because of a low population of simultaneously emitting molecules and/or excessively short *τ*_on_ times.

**Figure 8.**
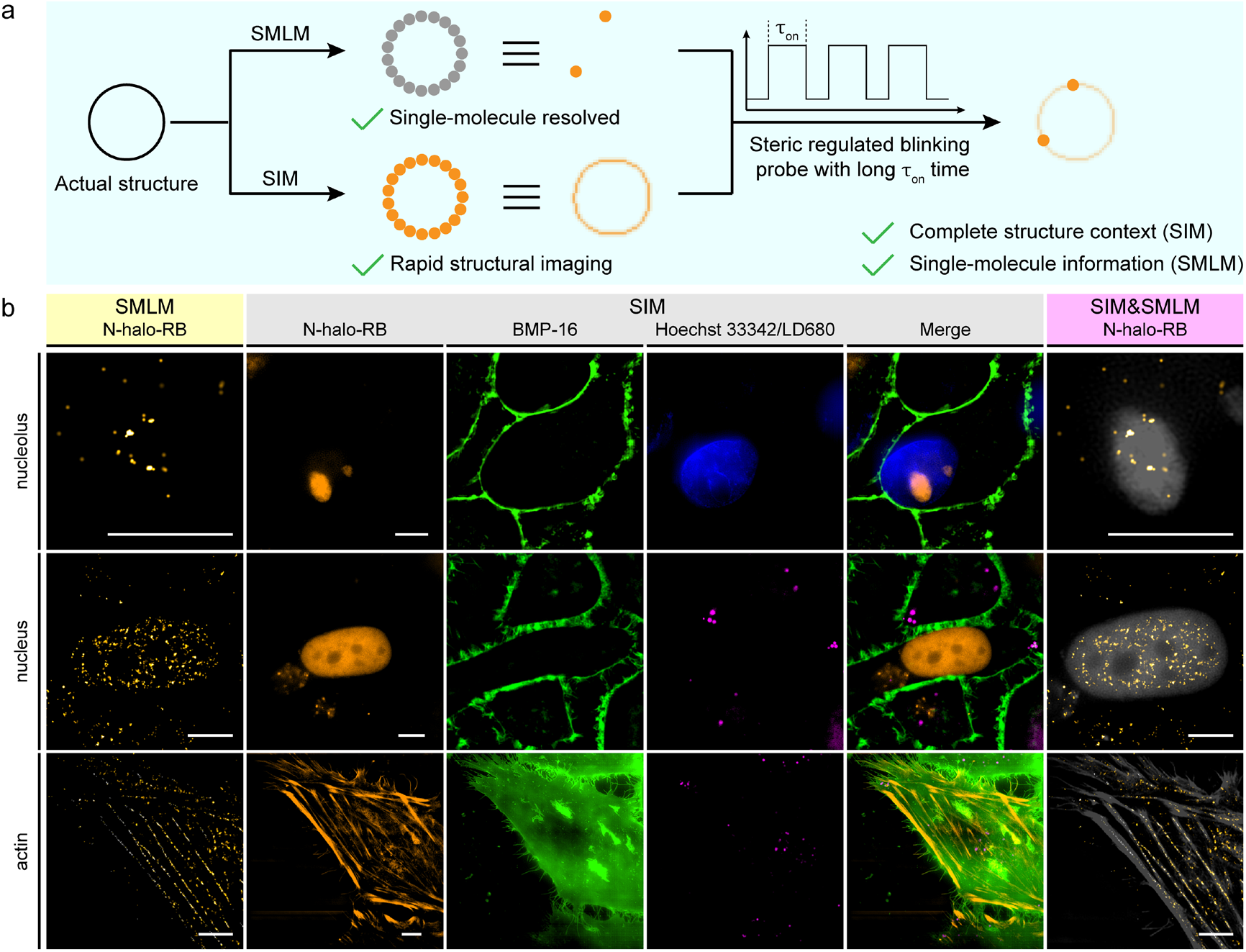
Dual-modal SIM/SMLM imaging with a single fluorophore. (a) Schematic illustration of the complementary imaging capabilities of SIM and SMLM. (b) In the top row, nucleoli were stained with **N-halo-RB** (400 nM, orange channel), the plasma membrane was labeled with **BMP-16** (1 μM, green channel), and nuclei were stained with **Hoechst 33342** (4 μM, blue channel). In the middle and bottom rows, nuclei and actin were labeled with **N-halo-RB** (100 nM, orange channel), the cell membrane was stained with **BMP-16** (1 μM, green channel), and lipid droplets were labeled with **LD680** (magenta channel). The right panel shows the mapping of single-molecule events obtained by SMLM onto the complete structural image obtained by SIM. Scale bar: 5 μm.

Our previous results identified **N-halo-RB** as a promising candidate for overcoming this challenge. In conventional fluorogenic imaging, **N-halo-RB** showed exhibited a pronounced fluorescence turn-on response upon HaloTag labeling, indicating that a sufficient population of fluorescent molecules could be generated for ensemble imaging (Figure 4a,b). Meanwhile, it displayed distinct spontaneous blinking behavior and good localization precision under SMLM conditions (Figure 6d–f). This combination distinguishes **N-halo-RB** from the other probes investigated. Specifically, **O-halo-RB** produced no detectable fluorescence under conventional imaging conditions, potentially owing to an insufficient population of emissive molecules, whereas **5,6-halo-RB**, despite its pronounced fluorogenic response, showed no detectable blinking under our SMLM imaging conditions. Notably, **N-halo-RB** has a higher p*K*_cycl_ value than **5,6-halo-RB** and **O-halo-RB**, indicating a greater propensity to populate the ring-open state under comparable conditions. Combined with the prolonged *τ*_on_ enabled by our steric regulation strategy, this equilibrium preference may provide a favorable balance between the population of emissive molecules and the duration of individual emission events. Such a balance is expected to support the generation of sufficient fluorescence for SIM while retaining the temporal intermittency required for SMLM. On this basis, we selected **N-halo-RB** to explore dual-modal super-resolution imaging using a single probe.

To demonstrate this capability, we employed **N-halo-RB** to label the nucleoli, nuclei, and actin in HeLa cells and performed SIM and SMLM imaging. We first examined the information obtainable from SMLM under a deliberately restricted acquisition period of 2 s, mimicking the challenges encountered when imaging dynamic biological processes. Under these conditions, the limited number of localization events was insufficient to reconstruct the complete structures of the labeled cellular components. In contrast, SIM recovered their overall structural features within only 200 ms. Importantly, overlaying the two imaging modalities enabled the localization events resolved by SMLM to be mapped onto the complete structural context provided by SIM, thereby combining single-molecule-level information with rapid visualization of cellular architecture (Figure 8b). Furthermore, because both modalities relied on the same probe, **N-halo-RB**, dual-modal imaging required only a single excitation/detection channel, leaving other spectral channels available for additional fluorescent probes. To demonstrate this potential for multicolor imaging, we further introduced **Hoechst 33342**, **BMP-16**, and **LD680** to label nuclei, plasma membranes, and lipid droplets, respectively. This strategy establishes a single-probe approach for integrating SIM and SMLM, while retaining the spectral flexibility needed to investigate the spatial organization and interactions of multiple subcellular structures.

### Conclusions

In conclusion, we established N-substituent steric regulation as a molecular strategy for tuning the spirocyclization equilibrium and switching kinetics of rhodamine fluorophores. Introducing bulky substituents, particularly tert-butyl groups, onto the amide nitrogen modulates both ring-opening thermodynamics and switching kinetics. Single-crystal analysis revealed substantial amide reorientation upon ring opening, increasing the separation between the nitrogen and meso-carbon reaction centers. Together with theoretical calculations, these findings suggest a conformational-gating mechanism in which sterically regulated accessibility of the reverse conformation controls C–N bond reformation. Notably, the tert-butyl-substituted dye **1e** exhibited an open-state residence time of 145 ms, despite its lower p*K*_cycl_, and a lower ring-opening barrier than **1a** (1.12 vs 1.37 eV). We further extended this strategy to different rhodamine scaffolds to develop fluorogenic HaloTag probes with fluorescence enhancements of up to 124-fold. The resulting probes support multiple super-resolution imaging modalities, including SIM, STED, and SMLM. Importantly, **N-halo-RB** enabled dual-modal SIM/SMLM imaging with a single fluorophore and fluorescence channel, combining rapid structural reconstruction with single-molecule-level information. Overall, this work establishes sterically modulated conformational accessibility as a complementary design principle for engineering reversible fluorescent switches and expanding their applications in super-resolution imaging.

## ACKNOWLEDGMENTS

This work is supported by the National Natural Science Foundation of China (Nos. 22225806, 22522816, 22578443, 22378385, 22278394, 22507121, 22508383, 22541705, U25A20620); The Chinese Academy of Sciences Project for Young Scientists in Basic Research (No. YSBR-104); The Energy Revolution S&T Program of Yulin Innovation Institute of Clean Energy (No. E412050705); Liaoning Binhai Laboratory (No. LBLD-2024-07); Dalian Institute of Chemical Physics (Nos. DICP I202436, DICP I202522, DICP I202512, DICP&SIA UN202502, DMU-1&DICP UN202301, DMU-1&DICP UN202302, DMU-2&DICP UN202502); Natural Science Foundation of Liaoning (Nos. 2025-MS-061, 2025-BS-0147); Dalian Science and Technology Innovation Fund Program (No. 2022JJ11CG007).

## 1 Computational methods

### 1.1 Computational methods

The ground-state geometries of compounds were optimized using density functional theory (DFT) at the M06-2X^[1]^/TZVP level using the Gaussian 16 A.03 program. We have performed frequency calculations to ensure that we find stable/transition states. The solvent effect in water was accounted for using the SMD model.

## 2 Experimental details

### 2.1 Materials and instrument

Unless explicitly stated, all reagents were purchased from commercial suppliers (Sigma-Aldrich, J&K, Innochem and Aladdin) and used without further purification. Solvents [dimethyl sulfoxide (DMSO), dimethylformamide (DMF), methanol (MeOH)] were purchased from J&K and used without further treatment or distillation. Silica gel (200-300 mesh) and aluminium oxide basic (200-300 mesh) were purchased from Innochem. Halo-Tag standard protein was purchased from Promega. Plasmids used in this paper were all purchased from Addgene.

All ^1^H-NMR and ^13^C-NMR spectra were recorded on a Bruker 400 M spectrometer or 700 M spectrometer with TMS as an internal standard. Chemical shifts were given in ppm and coupling constants (*J*) in Hz. High resolution mass spectrometry data were obtained with a HP1100LC/MSD mass spectrometer and a LC/Q-TOF MS spectrometer. UV-vis absorption spectra were collected on an Agilent Cary 60 UV-Vis Spectrophotometer. The blank correction was performed using the respective pure solvent as a reference. Fluorescence measurements were performed on an Agilent CARY Eclipse fluorescence spectrophotometer and Horiba Fluoromax-4 spectrophotometer. The X ray crystal diffraction patterns were recorded by Agilent GeminiUltra X-ray diffractometer with Atlas CCD at 20 °C.

Confocal images were performed on Olympus FV1000MPE with a microscope IX 71, a 100 × / NA 1.40 oil objective lens, LU-NV series laser unit (laser combination: 405 nm; 543 nm; 488 nm; 635 nm). STED images were performed on Abberior STED super-resolution scanning unit and detection system (STEDYCON). 775 nm depletion laser was used for imaging.

SMLM images were performed by Nikon *N*-STORM/SIM 5.0 Super-Resolution Microscope System with a motorized inverted microscope ECLIPSE Ti2-E, a 100 × / NA 1.49 oil immersion TIRF objective lens (CFI HP), LU-NV series laser unit (405 nm, 488 nm, 561 nm, 647 nm), and an ORCA-Flash 4.0 SCMOS camera (Hamamatsu Photonics K.K.). Confocal images were performed on Olympus FV1000MPE with a microscope IX 71, a 100 × / NA 1.40 oil objective lens, LU-NV series laser unit (laser combination: 543 nm).

### 2.2 Spectroscopic studies

Probes were firstly prepared as 2 mM stock solution in HPLC DMSO. Samples for spectroscopic analysis were prepared through diluting the stock solution in respective solvents. The concentration of samples was usually prepared at 5 μM, unless stated otherwise. UV-vis and fluorescence spectra were recorded using the Agilent Cary 60 UV-Vis Spectrophotometer and Agilent CARY Eclipse fluorescence spectrophotometer. In order to record UV-vis and fluorescence spectra of probes at different pH, different buffer solutions were prepared to stabilize pH. Phosphate buffer (20 mM) was selected to provided stable pH conditions at pH 6 - 8. Citrate buffer solution (20 mM) was selected to provided stable pH conditions at pH 3 - 6. Water-dioxane mixtures containing 0%, 10%, 20%, 30%, 40%, 50%, 60%, 70%, 80%, 90%, 100% anhydrous dioxane were prepared for measurements in Water-Dioxane Mixtures. All samples were stabilized in dark at room temperature for at least 1 h before testing.

### 2.3 Cell culture and transfection

#### 2.3.1 Cell culture

HeLa (helacyton gartleri) cells were purchased from Cell Bank of Type Culture Collection of Chinese Academy of Sciences. HeLa cells were maintained in Dulbecco’s modified Eagle’s medium (DMEM, Gibco) supplemented with 10% fetal bovine serum (FBS, Hyclone) which were cultured in a humidified atmosphere of 5% CO_2_/95% air at 37 °C. Before the imaging experiments, HeLa cells were seeded on glass bottom cell culture dish (Nest, polystyrene, Φ 15 mm) for 1 - 2 days to reach 40–90% confluency. The cells were then used for further experiments.

#### 2.3.2 Cell transfection

Halo-H2B, Halo-tubulin, Halo-actin-EGFP, Halo-LifeAct-halo-tom20 and halo-tom20-EGFP were purchased from Addgene. Transfection experiment was performed according to the PE-1 manufacturer’s protocol.

Briefly, 2 μL PE-1 and appropriate plasmid were firstly diluted in 20 μL DMEM (dulbecco’s modified eagle medium), respectively. 5 min later, the diluted plasmid in 20 μL DMEM was added to the diluted PE-1 with homogeneous mixing. Another 10 min later, the mixture was added to the cell-culture dish in 1 mL DMEM. The final concentration of plasmid was controlled at 500-1000 ng/mL. After incubated 4 h in 37 °C, the culture medium was changed from DMEM to DMEM with 10% FBS. 24 - 48 h later, the transfected cells were used for confocal imaging and super-resolution imaging.

#### 2.3.3 Cell viability

The cell viability was tested via MTT assay. HeLa cells were incubated in 96-well plate for 24 h before used in cell viability experiment. The DMEM was changed with fresh DMEM containing different concentrations of porbes (0.1, 0.5, 1.0, 2.0, 5.0,10 μM). The cells were incubated for 24 h at 37 °C with a 5% CO2 atmosphere. Then, the medium was replaced with 100 μL fresh medium and 10 μL of 5 mg/mL MTT solution per well. 4 h later, the medium was removed out carefully, and 200 μL DMSO was added to each well to dissolve blue formazan. After sufficient shaking for 10 min, the absorption was recorded at 490 nm using a UV-Vis microplate reader and used for analysis of cell viability.

### 2.4 Single molecule studies

#### 2.4.1 Coverslip cleaning

The coverslips were placed in a clean beaker and sonicated first in ultrapure water for 10 minutes. Then the ultrapure water was replaced with a 10% Alconox solution and sonicated for 10 minutes. The coverslips were rinsed with ultrapure water until no bubbles were visible and then sonicated in ethanol and acetone for 10 minutes respectively. After drying under a stream of nitrogen, the coverslips were placed into a piranha solution (H_2_SO_4_/30% H_2_O_2_ = 7:3, v/v) and heated to 90-100 °C for 2 hours. The cleaned coverslips were rinsed with ultrapure water and acetone and dried under a stream of nitrogen.

The molecules were dropped onto a cleaned coverslip and form a PMMA film at a concentration controlled at 10^-9^ – 10^-11^ M. Then, the sample was used for single-molecule analysis recorded with the microscopy described above. Single-molecule signals were acquired with an exposure time of 20 ms under selected laser intensity for each sample (Excitation: 561 nm; 0.12 kW/cm^2^, 0.60 kW/cm^2^, 0.12 kW/cm^2^, 0.60 kW/cm^2^, 0.12 kW/cm^2^).

### 2.5 Imaging

#### 2.5.1 confocal imaging for fluorogenic characterization

Imaging of different organelles labeled with **5,6-halo-RB**, **O-halo-RB** and **N-halo-RB** respectively were performed through confocal imaging. Ex: 543 nm; collected: 560-660 nm.

#### 2.5.2 confocal imaging for validating the transfection effectiveness

Imaging between EGFP and **5,6-halo-RB**, **O-halo-RB**, **N-halo-RB** were performed through confocal imaging. Ex: 488 nm; collected: 500-530 nm for EGFP. Ex: 543 nm; collected: 560-660 nm for **5,6-halo-RB**, **O-halo-RB**, **N-halo-RB**.

#### 2.5.3 SIM imaging for fluorogenic characterization

The fluorogenic ability was further demonstrated through Structured Illumination Microscopy (SIM; Nikon *N-*SIM Ti-2E). The cells which were transfected with Halo-H2B, Halo-tubulin, lifeact-Halo and halo-tom20-EGFP were incubated with **5,6-halo-RB**. Ex:561nm, collected: 570-640 nm.

#### 2.5.4 STED imaging for fluorogenic characterization

The fluorogenic ability was further demonstrated through Stimulated Emission Depletion Microscopy (STED; Abberior STEDYCON). 775 nm depletion laser was used for imaging.). The cells which were transfected with lifeact-Halo and halo-tom20-EGFP were incubated with **5,6-halo-RB**. Ex:561nm, collected: 570-640 nm.

#### 2.5.5 SMLM imaging for nuclei in living cells

Generally, HeLa cells were incubated with probes (500 nM) for 1 h in a humidified atmosphere of 5% CO_2_/95% air at 37 °C. Then the DMEM was removed and changed with 1 mL PBS (pH = 7.4). The super-resolution imaging was performed on Nikon *N*-STORM/SIM 5.0 Super-Resolution Microscope System with an exposure time of 20 ms.

#### 2.5.6 SMLM imaging for actin in living cells

Generally, HeLa cells were incubated with **O-halo-JF549** (400 nM) for 1 h in a humidified atmosphere of 5% CO_2_/95% air at 37 °C. The super-resolution imaging was performed on Nikon *N*-STORM/SIM 5.0 Super-Resolution Microscope System with an exposure time of 20 ms.

#### 2.5.7 SMLM imaging for tubulin in fixed cells

Generally, HeLa cells were incubated with **N-halo-RB** (500 nM) for 1 h in a humidified atmosphere of 5% CO_2_/95% air at 37 °C. Then the cells are fixed according to the procedure reported in the literature.^[2]^ The super-resolution imaging was performed on Nikon *N*-STORM/SIM 5.0 Super-Resolution Microscope System with an exposure time of 20 ms.

## 3 Synthesis

### 3.1 Synthesis of **1a**

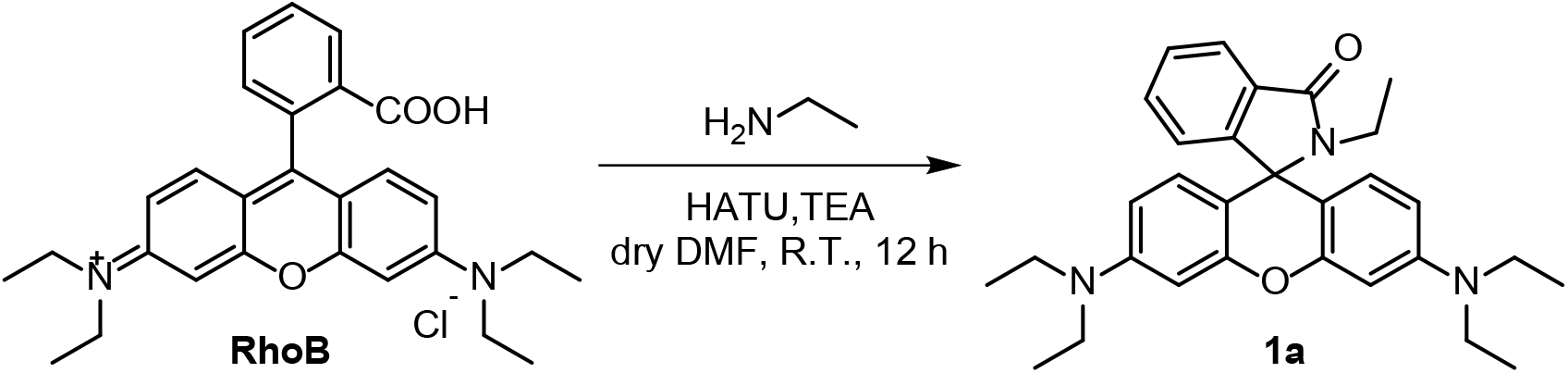

To a solution of rhodamine B (0.2111 g, 0.44 mmol, 1 eq) and 2-(7-Azabenzotriazol-1-yl)-N,N,N’,N’-tetramethyluronium hexafluorophosphate (HATU, 0.1662 g, 0.49 mmol, 1.1 eq) in dry N,N-dimethylformamide (DMF, 10 mL) was added triethylamine (TEA, 90 μL, 1.5 eq). The reaction was stirred at room temperature for 20 min. Following the addition of 2 mol/L ethylamine in THF (245 μL, 0.49 mmol, 1.1 eq), the reaction was stirred under room temperature for 12 h. After removing the solvent under reduced pressure, the residue was purified by alkaline alumina column chromatography (petroleum ether/ethyl acetate=3:1, v/v) to afford a white solid (20.4 mg, 9.3%). ^1^H NMR (400 MHz, CDCl_3_) δ 7.90 (s, 1H), 7.41 (s, 2H), 7.06 (s, 1H), 6.46 (d, *J* = 8.6 Hz, 2H), 6.39 (s, 2H), 6.27 (d, *J* = 8.4 Hz, 2H), 3.33 (d, *J* = 6.7 Hz, 8H), 3.20 (d, *J* = 6.9 Hz, 2H), 1.16 (t, *J* = 6.4 Hz, 12H), 0.82 (t, *J* = 6.7 Hz, 3H).

### 3.2 Synthesis of **1b**

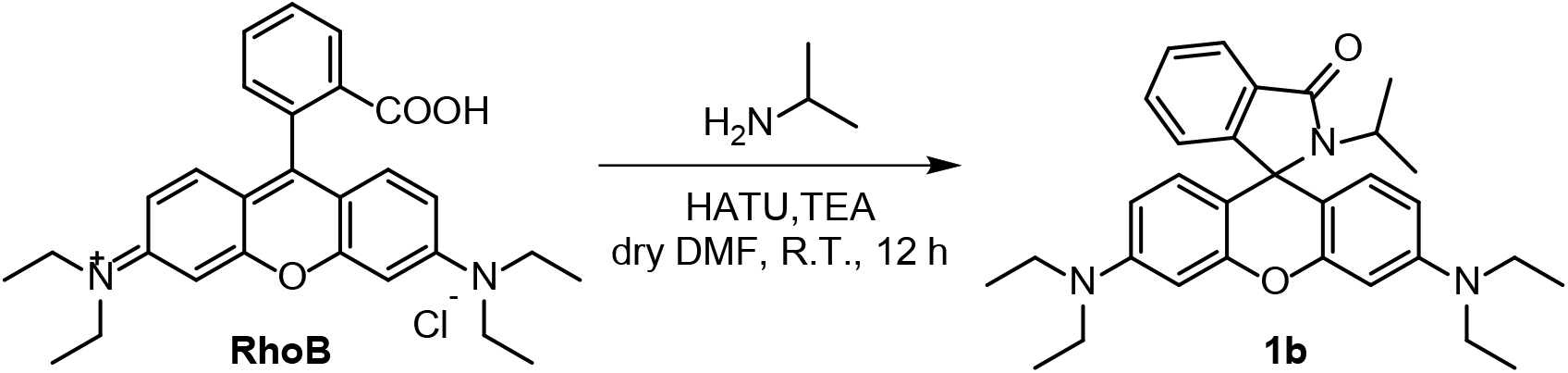

The synthesis of **1b** was similar with **1a**. Rhodamine B (0.0535 g, 0.11 mmol, 1 eq), HATU (0.0619 g, 0.16 mmol, 1.5 eq), TEA (30 μL, 2 eq) and isopropyl amine (20 μL, 0.23 mmol, 2 eq). **1b** was obtained as a white solid (20.0 mg, 37.6%). ^1^H NMR (400 MHz, CDCl_3_) δ 7.91 – 7.82 (m, 1H), 7.43 – 7.36 (m, 2H), 7.03 – 6.98 (m, 1H), 6.50 (d, *J* = 8.8 Hz, 2H), 6.37 (d, *J* = 2.2 Hz, 2H), 6.27 (dd, *J* = 8.8, 2.3 Hz, 2H), 3.39 – 3.26 (m, 9H), 1.20 – 1.15 (m, 18H). ^13^C NMR (101 MHz, CDCl_3_) δ 167.60, 153.56, 151.71, 147.26, 132.55, 132.05, 132.05, 129.11, 127.86, 123.64, 122.41, 117.59, 106.12, 96.48, 77.35, 77.03, 76.71, 45.74, 38.40, 20.05, 20.05, 16.65, 14.77, 14.77. HRMS (ESI) calcd for C_31_H_38_N_3_O_2_ [M+H]^+^ 484.2964, found 484.2968.

### 3.3 Synthesis of **1c**

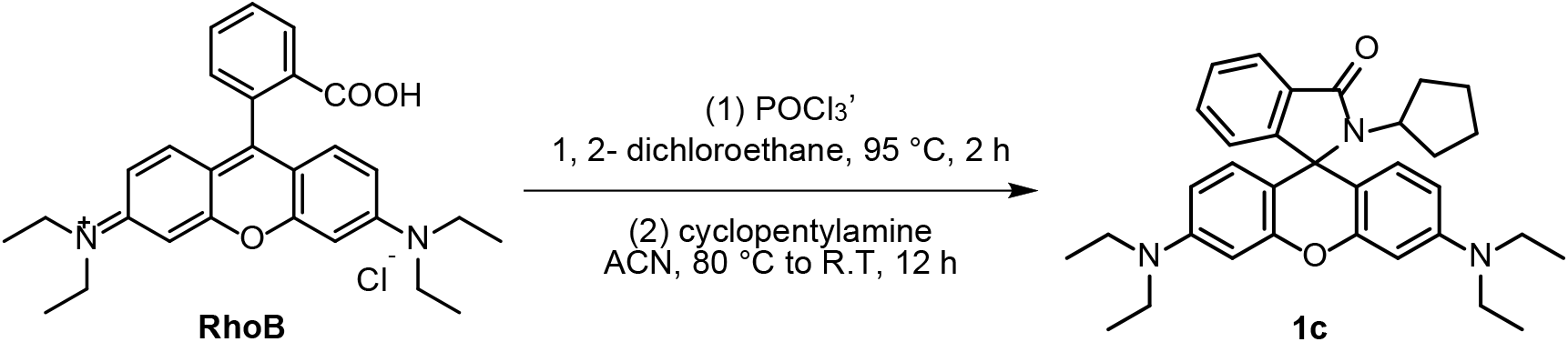

To the solution of rhodamine B (0.8500 g, 1.77 mmol, 1 eq) in 1,2-dichloroethane (50 mL) was added phosphorus oxychloride (1 mL, 10.73 mmol, 6 eq). The reaction was refluxed at 95 °C for 2 h. After cooling to room temperature, the solvent was removed under reduced pressure. To the residue dissolved in acetonitrile (30 mL) was added TEA (0.5 mL,3.6 eq) dropwise. After cyclopentylamine (349 μL, 3.54 mmol, 2 eq) was added, the reaction was refluxed at 80 °C for 1h and then cooled to room temperature and stirred for 12 h. After removing the solvent under reduced pressure, the residue was purified by silica gel column chromatography (petroleum ether/ethyl acetate=3:1, v/v) to afford a white solid (600.0 mg, 66.5%). ^1^H NMR (400 MHz, CDCl_3_) δ 7.83 (dd, *J* = 5.6, 2.5 Hz, 1H), 7.35 (p, *J* = 7.7 Hz, 2H), 6.97 (dd, *J* = 5.6, 2.4 Hz, 1H), 6.56 (d, *J* = 8.8 Hz, 2H), 6.38 (d, *J* = 2.3 Hz, 2H), 6.28 (dd, *J* = 8.9, 2.3 Hz, 2H), 3.42 – 3.29 (m, 9H), 2.20 (dt, *J* = 15.1, 7.5 Hz, 2H), 1.82 – 1.73 (m, 2H), 1.44 – 1.29 (m, 4H), 1.17 (t, *J* = 7.0 Hz, 12H). ^13^C NMR (101 MHz, CDCl_3_) δ 167.61, 153.98, 153.09, 148.64, 131.91, 131.82, 128.93, 127.69, 123.47, 122.36, 107.87, 105.99, 97.79, 65.74, 54.73, 44.33, 29.02, 24.79, 12.64.

### 3.4 Synthesis of **1d**

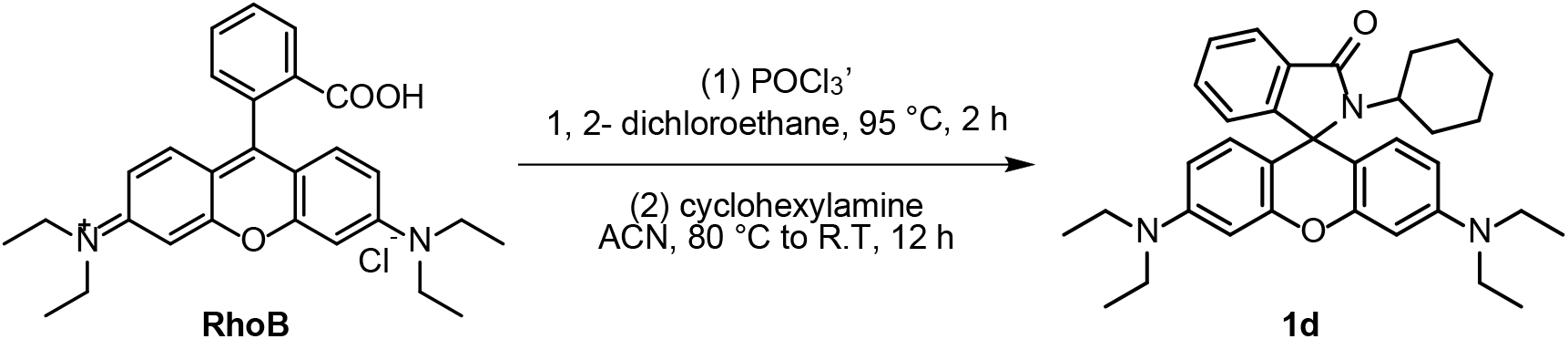

The synthesis of **1d** was similar with **1c**. Rhodamine B (0.8500 g, 1.77 mmol, 1 eq), phosphorus oxychloride (1 mL, 10.73 mmol, 6 eq), TEA (0.5 mL,3.6 eq) and cyclopentylamine (405 μL, 3.54 mmol, 2 eq). **1d** was obtained as a white solid (500.0 mg, 53.9%). ^1^H NMR (400 MHz, CDCl_3_) δ 7.85 (d, *J* = 5.4 Hz, 1H), 7.37 (s, 2H), 6.98 (d, *J* = 5.6 Hz, 1H), 6.47 (t, *J* = 16.6 Hz, 2H), 6.37 (s, 2H), 6.26 (d, *J* = 8.1 Hz, 2H), 3.34 (d, *J* = 5.5 Hz, 8H), 2.93 (s, 1H), 2.17 (dd, *J* = 22.5, 11.1 Hz, 2H), 1.57 (d, *J* = 11.4 Hz, 2H), 1.44 (d, *J* = 12.1 Hz, 1H), 1.33 – 1.06 (m, 16H), 0.95 (d, *J* = 12.2 Hz, 1H). ^13^C NMR (101 MHz, CDCl_3_) δ 167.42, 153.60, 153.25, 148.69, 132.50, 131.90, 129.48, 127.75, 123.60, 122.38, 107.85, 105.86, 97.71, 65.53, 54.03, 44.33, 29.48, 26.36, 25.28, 12.62.

### 3.5 Synthesis of **1e**

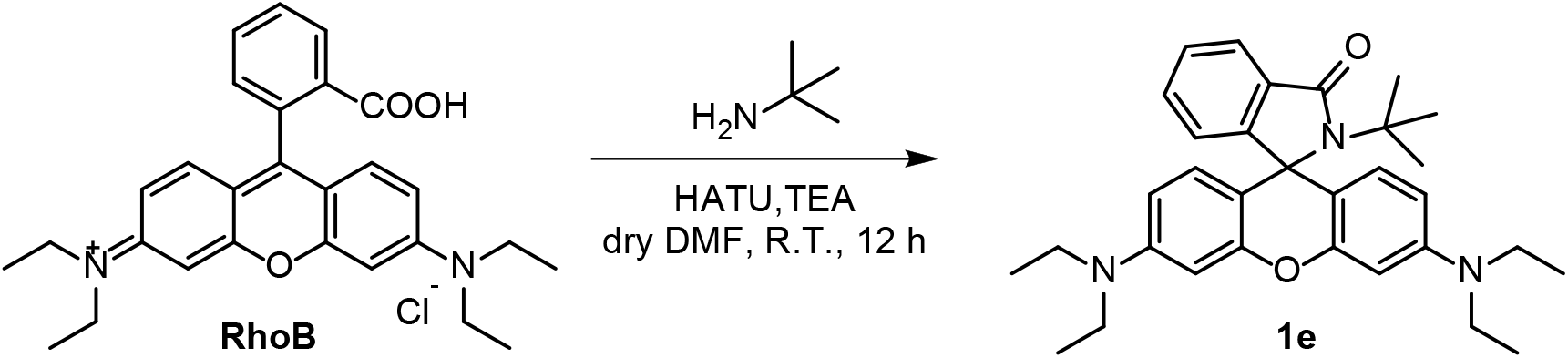

The synthesis of **1e** was similar with **1a**. Rhodamine B (0.0613 g, 0.13 mmol, 1 eq), HATU (0.0780 g, 0.20 mmol, 1.5 eq), TEA (36 μL, 2 eq) and tert-Butylamine (100 μL, 0.94 mmol, 7.2 eq). **1e** was obtained as a white solid (33.9 mg, 52.4%). ^1^H NMR (700 MHz, DMSO-*d6*) δ 7.67 (d, *J* = 6.9 Hz, 1H), 7.42 – 7.34 (m, 2H), 6.78 (d, *J* = 6.9 Hz, 1H), 6.46 (d, *J* = 8.7 Hz, 2H), 6.39 – 6.30 (m, 4H), 3.33 – 3.25 (m, 8H), 1.20 (s, 8H), 1.09 (t, *J* = 6.9 Hz, 12H).

### 3.6 Synthesis of **1f**

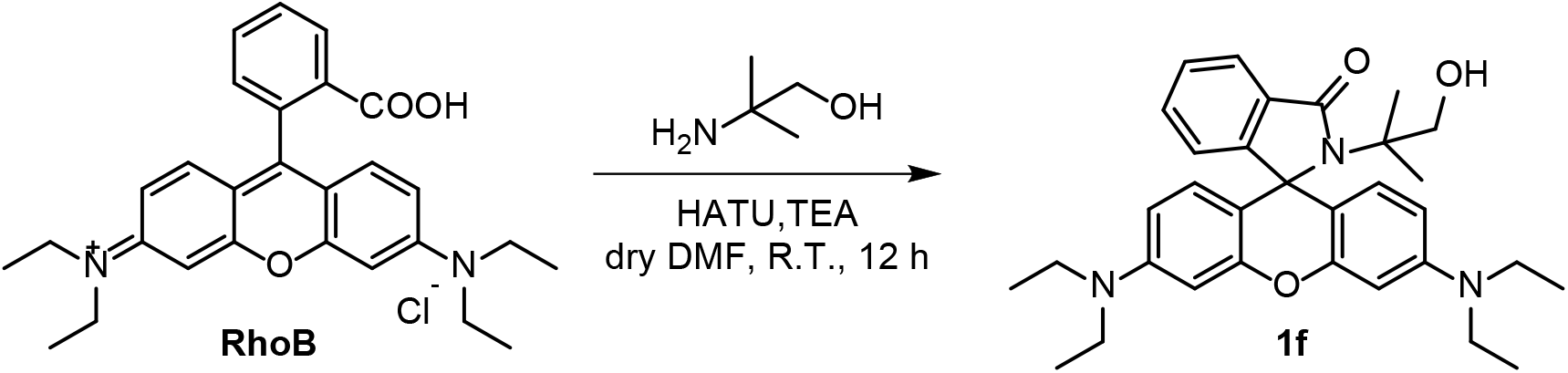

The synthesis of **1f** was similar with **1a**. Rhodamine B (0.4790 g, 1.00 mmol, 1 eq), HATU (0.5700 g, 1.68 mmol, 1.7 eq), TEA (300 μL, 2.2 eq) and 2-amino-2-methyl-1-propanol (200 μL, 2.09 mmol, 2.1 eq). **1f** was obtained as a pink solid (330.0 mg, 64.2%). ^1^H NMR (400 MHz, CDCl_3_) δ 7.85 – 7.76 (m, 1H), 7.41 – 7.30 (m, 2H), 6.92 – 6.78 (m, 1H), 6.56 (d, *J* = 8.8 Hz, 2H), 6.35 (d, *J* = 2.5 Hz, 2H), 6.28 (dd, *J* = 8.9, 2.6 Hz, 2H), 5.46 (t, *J* = 6.2 Hz, 1H), 3.61 (d, *J* = 6.2 Hz, 2H), 3.33 (q, *J* = 7.1 Hz, 8H), 1.23 – 1.10 (m, 18H). ^13^C NMR (101 MHz, CDCl_3_) δ 171.74, 155.75, 151.91, 148.78, 132.76, 128.98, 128.73, 127.64, 123.28, 122.43, 107.76, 107.11, 97.89, 73.89, 66.69, 61.90, 44.32, 23.48, 12.60. HRMS (ESI) calcd for C_32_H_40_N_3_O_3_ [M+H]^+^ 514.3070, found 514.3078.

### 3.7 Synthesis of **O-halo-RB**

The reported 2-(2-((6-chlorohexyl)oxy)ethoxy)ethan-1-ol was synthesized according to literature.^[3] 1^H NMR (400 MHz, CDCl_3_) δ 3.76 – 3.72 (m, 2H), 3.68 (dd, *J* = 5.7, 3.5 Hz, 2H), 3.65 – 3.57 (m, 4H), 3.54 (t, *J* = 6.7 Hz, 2H), 3.48 (t, *J* = 6.6 Hz, 2H), 1.82 – 1.74 (m, 2H), 1.66 – 1.57 (m, 2H), 1.51 – 1.33 (m, 4H).

The reported 2-(2-((6-chlorohexyl)oxy)ethoxy)ethan-1-amine was synthesized according to literature.^[4] 1^H NMR (400 MHz, CDCl_3_) δ 3.72 (s, 2H), 3.68 – 3.61 (m, 2H), 3.61 – 3.55 (m, 2H), 3.52 (t, *J* = 6.6 Hz, 2H), 3.46 (t, *J* = 6.8 Hz, 2H), 3.18 (s, 2H), 1.83 – 1.70 (m, 2H), 1.64 – 1.52 (m, 2H), 1.50 – 1.38 (m, 2H), 1.39 – 1.26 (m, 2H).

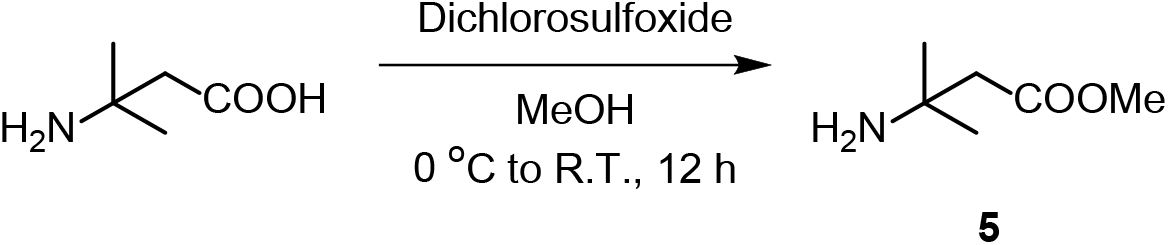

The solution of 3-amino-3-methylbutanoic acid (5.3 g, 45 mmol, 1 eq) in methanol (150 mL) was stirred at 0 °C for 0.5 h. After dichlorosulfoxide (6.0 mL, 83 mmol, 1.8 eq) was added dropwise, the reaction was warmed to room temperature and stirred for 12 h. The solvent was evaporated, and the residue was washed with water (100 mL, 3×). The residue was extracted by dichloromethane and dried over anhydrous sodium sulfate. The organic phase was concentrated to afford a white waxy solid (5.5g, 93.2%).

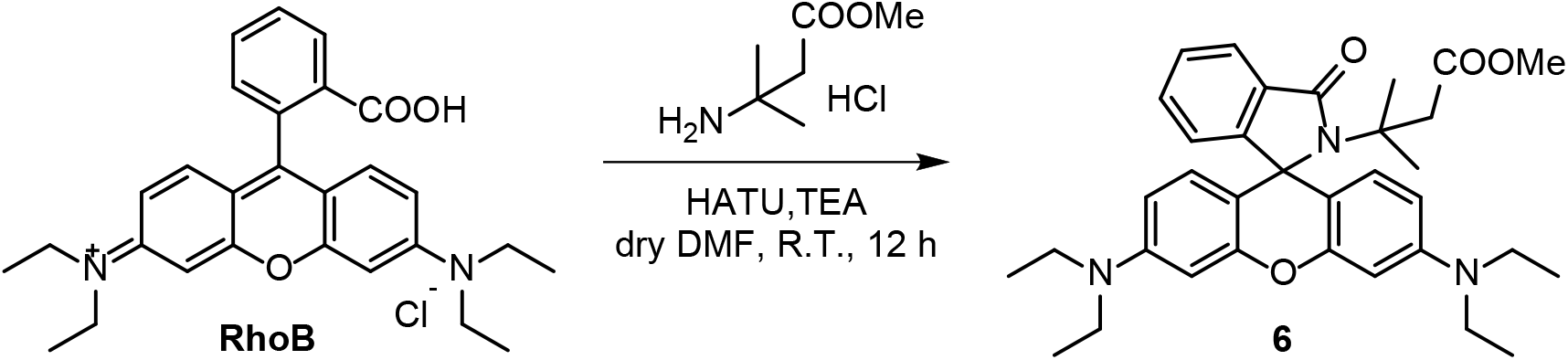

To a solution of rhodamine B (0.4790 g, 1.00 mmol, 1 eq) and HATU (0.5894 g, 1.55 mmol, 1.6 eq) in dry DMF (10 mL) was added TEA (300 μL, 2.2 eq). Following the addition of methyl 3-amino-3-methylbutanoate hydrochloride (0.3436 g, 2.05 mmol, 2.05 eq), the reaction was stirred under room temperature for 12 h. After removing the solvent under reduced pressure, the residue was purified by silica gel column chromatography (petroleum ether/ethyl acetate=5:1, v/v) to afford a white solid (345.9 mg, 61.0%). ^1^H NMR (400 MHz, CDCl_3_) δ 7.84 – 7.78 (m, 1H), 7.37 – 7.30 (m, 2H), 6.87 – 6.82 (m, 1H), 6.54 (d, *J* = 8.8 Hz, 2H), 6.34 (d, *J* = 2.6 Hz, 2H), 6.27 (dd, *J* 8.9, 2.6 Hz, 2H), 3.56 (s, 3H), 3.33 (q, *J* = 7.1 Hz, 8H), 3.15 (s, 2H), 1.31 (s, 6H), 1.17 (t, *J* = 7.0 Hz, 12H). ^13^C NMR (101 MHz, CDCl_3_) δ 171.74, 169.88, 155.40, 151.97, 148.68, 132.42, 129.80, 129.03, 127.46, 123.21, 122.28, 107.83, 107.72, 97.89, 65.94, 58.16, 51.22, 44.31, 44.07, 25.43, 12.62. HRMS (ESI) calcd for C_34_H_42_N_3_O_4_ [M+H]^+^ 556.3175, found 556.3243.

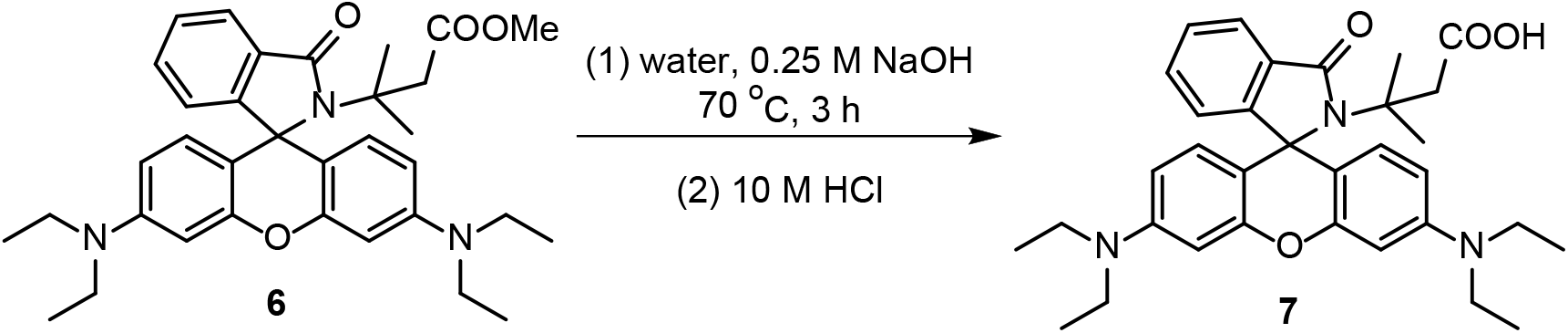

To the suspension of compound **6** (300 mg, 0.53 mmol) in water (20 mL) was added 0.25 M sodium hydroxide aqueous solution (10 mL). The mixture was stirred at 70 °C for 3 h. After cooling to room temperature, the mixture was extracted with dichloromethane (50 mL, 3x). The organic layer was collected and dried over anhydrous sodium sulfate. After removing the solvent under reduced pressure, the residue was collected as a red solid and used for next step without further purification. ^1^H NMR (400 MHz, CDCl_3_) δ 7.84 – 7.80 (m, 1H), 7.40 – 7.33 (m, 2H), 6.90 – 6.85 (m, 1H), 6.51 (d, *J* = 8.8 Hz, 2H), 6.34 (d, *J* = 2.3 Hz, 2H), 6.26 (dd, *J* = 8.8, 2.3 Hz, 2H), 3.33 (q, *J* = 7.0 Hz, 8H), 3.06 (s, 2H), 1.29 (s, 6H), 1.17 (t, *J* = 7.0 Hz, 12H). ^13^C NMR (101 MHz, CDCl_3_) δ 173.37, 170.62, 152.01, 148.85, 132.92, 129.19, 128.80, 127.72, 123.32, 122.40, 107.82, 106.85, 97.89, 66.93, 57.42, 47.55, 44.32, 26.61, 12.60. HRMS (ESI) calcd for C_33_H_40_N_3_O_4_ [M+H]^+^ 542.3019, found 542.3025.

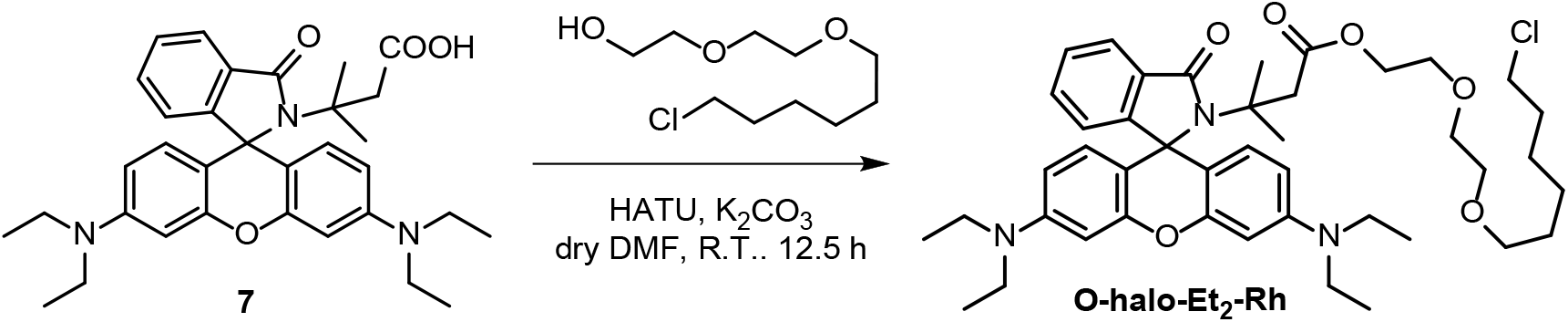

To a suspension of compound **7** (0.0391 g, 0.07 mmol, 1 eq), HATU (0.0276 g, 0.08 mmol, 1.1 eq) and anhydrous potassium carbonate (0.0261 g, 0.19 mmol, 2.7 eq) in dry DMF (5 mL) was added 2-(2-((6-chlorohexyl)oxy)ethoxy)ethan-1-ol (0.0158 g, 0.07 mmol, 1 eq). The reaction was stirred at room temperature for 12.5 h. After removing the solvent under reduced pressure, the residue was purified by alkaline alumina column chromatography (petroleum ether/ethyl acetate=3:1, v/v) to afford a red viscous liquid (11.5 mg, 21.9%). ^1^H NMR (400 MHz, CDCl_3_) δ 7.84 – 7.77 (m, 1H), 7.36 – 7.30 (m, 2H), 6.87 – 6.80 (m, 1H), 6.55 (d, *J* = 8.8 Hz, 2H), 6.34 (d, *J* = 2.6 Hz, 2H), 6.27 (dd, *J* = 8.9, 2.6 Hz, 2H), 4.16 – 4.09 (m, 2H), 3.63 – 3.60 (m, 2H), 3.58 – 3.48 (m, 6H), 3.42 (t, *J* = 6.6 Hz, 2H), 3.33 (q, *J* = 7.1 Hz, 8H), 3.15 (s, 2H), 1.81 – 1.73 (m, 2H), 1.61 – 1.53 (m, 2H), 1.45 – 1.36 (m, 4H), 1.33 (s, 6H), 1.17 (t, *J* = 7.0 Hz, 12H). HRMS (ESI) calcd for C_43_H_59_ClN_3_O_6_ [M+H]^+^ 748.4092, found 748.4115.

### 3.8 Synthesis of **N-halo-RB**

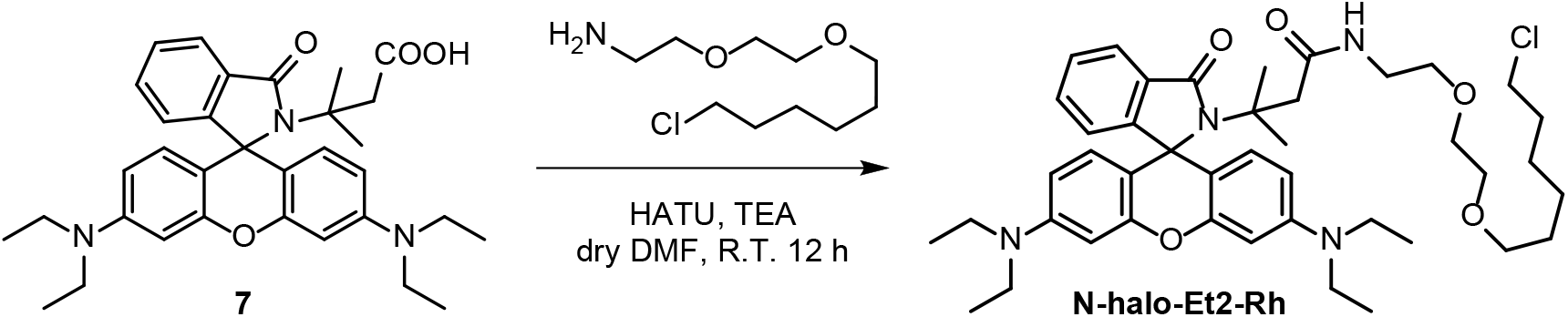

The synthesis of **N-halo-RB** was similar with **1a**. Compound **7** (0.1013 g, 0.19 mmol, 1 eq), HATU (0.1102 g, 0.29 mmol, 1.5 eq), TEA (53 μL, 2 eq) and 2-(2-((6-chlorohexyl)oxy)ethoxy)ethan-1-amine (0.0850 g, 0.38 mmol, 1.2 eq). **N-halo-RB** was obtained as a pink solid (28.4 mg, 20.1%). ^1^H NMR (400 MHz, CDCl_3_) δ 7.83 – 7.73 (m, 1H), 7.39 – 7.31 (m, 2H), 6.91 – 6.81 (m, 1H), 6.69 – 6.60 (m, 1H), 6.54 – 6.42 (m, 2H), 6.38 – 6.28 (m, 2H), 6.21 (ddd, 2H), 3.64 – 3.20 (m, 20H), 3.04 – 2.78 (m, 2H), 1.82 – 1.72 (m, 2H), 1.68 – 1.56 (m, 2H), 1.52 – 1.34 (m, 4H), 1.34 – 1.25 (m, 6H), 1.17 (t, *J* = 7.0 Hz, 12H). ^13^C NMR (101 MHz, CDCl_3_) δ 170.56, 151.98, 148.71, 129.72, 129.71, 128.84, 127.56, 127.49, 123.34, 122.07, 122.03, 107.66, 107.44, 97.90, 71.31, 70.33, 70.05, 69.90, 58.56, 47.07, 45.07, 44.29, 39.16, 32.55, 29.70, 29.46, 26.73, 25.96, 25.44, 12.63. HRMS (ESI) calcd for C_43_H_60_ClN_4_O_5_ [M+H]^+^747.4252, found 747.4229.

### 3.9 Synthesis of **5,6-halo-RB**

Compound **8** was synthesized according to literature *Chem. Eur. J*. **2014**, 20, 13162 – 13173.

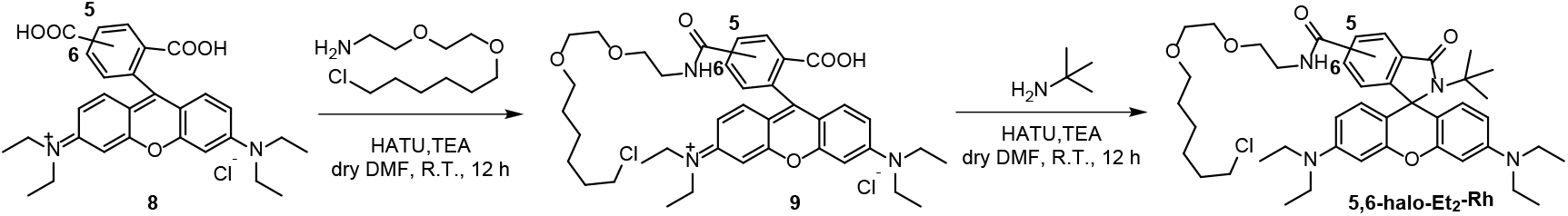

To a solution of **compound 8** (0.0522 g, 0.10 mmol, 1 eq) and HATU (0.0399 g, 0.10 mmol, 1 eq) in dry DMF (5 mL) was added TEA (60 μL, 3.9 eq). The reaction was stirred at room temperature for 30 min. Following the addition of 2-(2-((6-chlorohexyl)oxy)ethoxy)ethan-1-amine (0.0382 g, 0.11 mmol, 1 eq), the reaction was stirred under room temperature for 15 h. After removing the solvent under reduced pressure, the residue was purified by silica gel column chromatography (dichloromethane/methanol=10:1, v/v) to afford **compound 9** a red solid. ^1^H NMR (400 MHz, CDCl_3_) δ 8.10 (d, *J* = 7.9 Hz, 1H), 8.02 (d, *J* = 7.6 Hz, 1H), 7.62 (s, 1H), 6.71 (d, *J* = 8.7 Hz, 2H), 6.51 (s, 2H), 6.45 (d, *J* = 9.0 Hz, 2H), 3.67 – 3.59 (m, 6H), 3.57 – 3.47 (m, 6H), 3.41 (dt, *J* = 9.0, 6.8 Hz, 8H), 1.78 – 1.70 (m, 2H), 1.57 – 1.49 (m, 2H), 1.46 – 1.37 (m, 2H), 1.33 (dd, *J* = 13.9, 7.2 Hz, 2H), 1.21 (t, *J* = 7.0 Hz, 12H). **Compound 9** was used for the synthesis of **5,6-halo-RB** which was similar with **1a**. HATU (0.0406 g, 0.11 mmol, 1 eq), tert-butylamine (30 μL, 0.11 mmol, 1 eq) and TEA (30 μL, 3.9 eq). **5,6-halo-RB** was obtained as a white solid (2.1 mg, 2.8%). ^1^H NMR (400 MHz, CDCl_3_) δ 8.04 (d, *J* = 1.2 Hz, 1H), 7.80 (dd, *J* = 8.0, 1.6 Hz, 1H), 7.77 (d, *J* = 7.9 Hz, 1H), 7.65 (dd, *J* = 7.9, 1.4 Hz, 1H), 6.86 (d, *J* = 8.0 Hz, 1H), 6.47 (dd, *J* = 8.8, 1.0 Hz, 4H), 6.28 (d, *J* = 2.5 Hz, 4H), 6.19 (dt, *J* = 8.9, 2.4 Hz, 4H), 3.62 – 3.40 (m, 23H), 3.32 (t, *J* = 6.6 Hz, 3H), 3.27 (q, *J* = 7.2 Hz, 16H), 1.67 (dq, *J* = 13.1, 6.6 Hz, 4H), 1.53 – 1.42 (m, 4H), 1.40 – 1.29 (m, 8H), 1.23 (d, *J* = 2.6 Hz, 18H), 1.10 (td, *J* = 7.0, 1.7 Hz, 24H). HRMS (ESI) calcd for C_43_H_60_ClN_4_O_5_ [M+H]^+^ 747.4252, found 747.4263.

### 3.10 Synthesis of **2a**

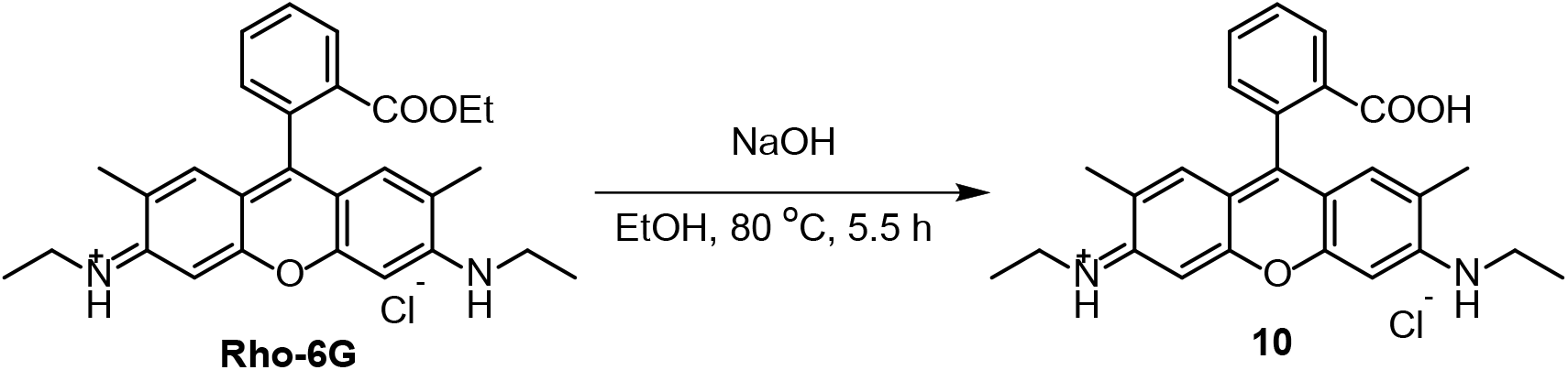

The solution of rhodamine 6G (1.0462 g, 2.18 mmol, 1eq) and NaOH (0.4457 g, 11.14 mmol, 5.1 eq) in ethanol (30 ml) was refluxed for 5.5 h. The reaction was cooled to room temperature and neutralized with concentrated HCl. The mixture was concentrated and purified by silica gel column chromatography (dichloromethane/methanol=8:1, v/v) to afford a red solid (490.4 mg, 49.9%).

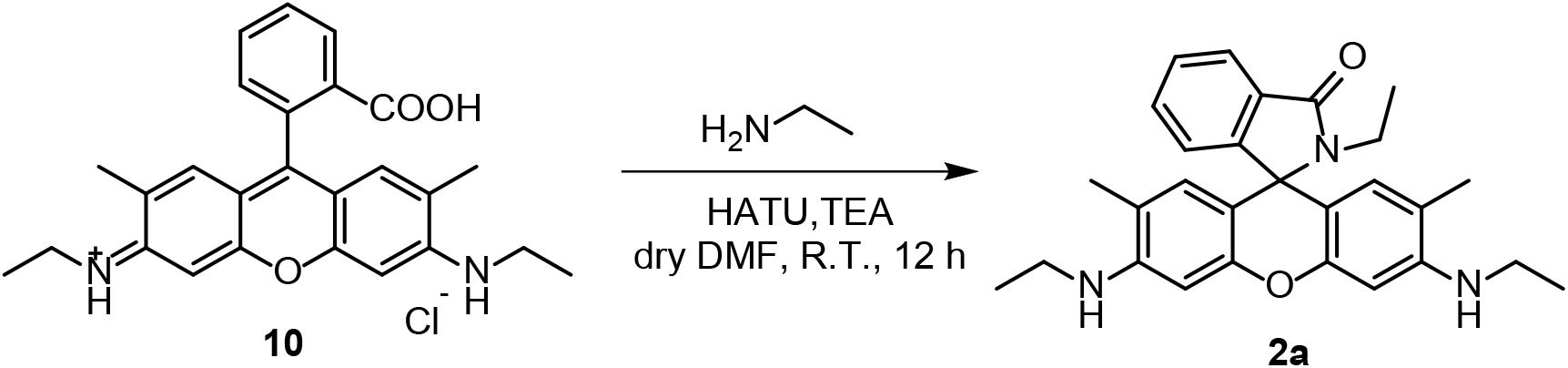

To a solution of **compound 10** (0.1173 g, 0.28 mmol, 1 eq) and HATU (0.1142 g, 0.34 mmol, 1.2 eq) in dry DMF (5 mL) was added TEA (50 μL, 1.3 eq). The reaction was stirred at room temperature for 20 min. Following the addition of 2 mol/L ethylamine in THF (500 μL, 1.0 mmol, 3.6 eq), the reaction was stirred under room temperature for 12 h. After removing the solvent under reduced pressure, the residue was purified by alkaline alumina column chromatography (petroleum ether/ethyl acetate=3:1, v/v) to afford a white solid (61.7 mg, 50.0%). ^1^H NMR (400 MHz, CDCl_3_) δ 7.92 (d, *J* = 6.5 Hz, 1H), 7.42 (p, *J* = 7.5 Hz, 2H), 7.02 (d, *J* = 6.6 Hz, 1H), 6.35 (s, 2H), 6.26 (s, 2H), 3.26 – 3.13 (m, 6H), 1.90 (s, 6H), 1.32 (t, *J* = 7.0 Hz, 6H), 0.78 (t, *J* = 7.1 Hz, 3H).

### 3.11 Synthesis of **2b**

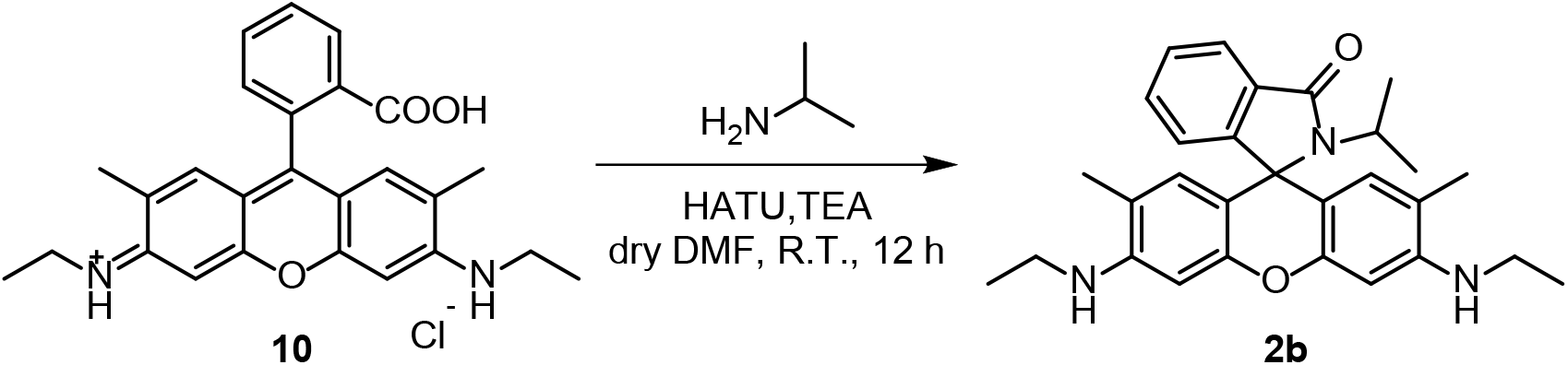

To a solution of **compound 10** (0.0789 g, 0.17 mmol, 1 eq) and HATU (0.0955 g, 0.25 mmol, 1.5 eq) in dry DMF (5 mL) was added TEA (47 μL, 2 eq). The reaction was stirred at room temperature for 20 min. Following the addition of Isopropyl amine (30 μL, 0.35 mmol, 2 eq), the reaction was stirred under room temperature for 12 h. After removing the solvent under reduced pressure, the residue was purified by silica gel column chromatography (petroleum ether/ethyl acetate=3:1, v/v) to afford a white solid (4.6 mg, 5.93%). ^1^H NMR (400 MHz, CDCl_3_) δ 7.88 (d, *J* = 6.8 Hz, 1H), 7.45 – 7.35 (m, 2H), 6.97 (d, *J* = 6.9 Hz, 1H), 6.34 (s, 2H), 6.31 (s, 2H), 3.61 (s, 2H), 3.29 (dt, *J* = 13.9, 7.0 Hz, 1H), 3.21 (q, *J* = 7.0 Hz, 4H), 1.91 (s, 5H), 1.33 (t, *J* = 7.1 Hz, 6H), 1.13 (d, *J* = 6.8 Hz, 6H). ^13^C NMR (101 MHz, CDCl_3_) δ 167.60, 153.55, 151.71, 147.26, 132.54, 132.05, 129.11, 127.86, 123.64, 122.41, 117.59, 106.12, 96.48, 65.65, 45.74, 38.40, 20.04, 16.65, 14.76. HRMS (ESI) calcd for C_29_H_34_N_3_O_2_ [M+H]^+^ 456.2651, found 456..2652.

### 3.12 Synthesis of **2c**

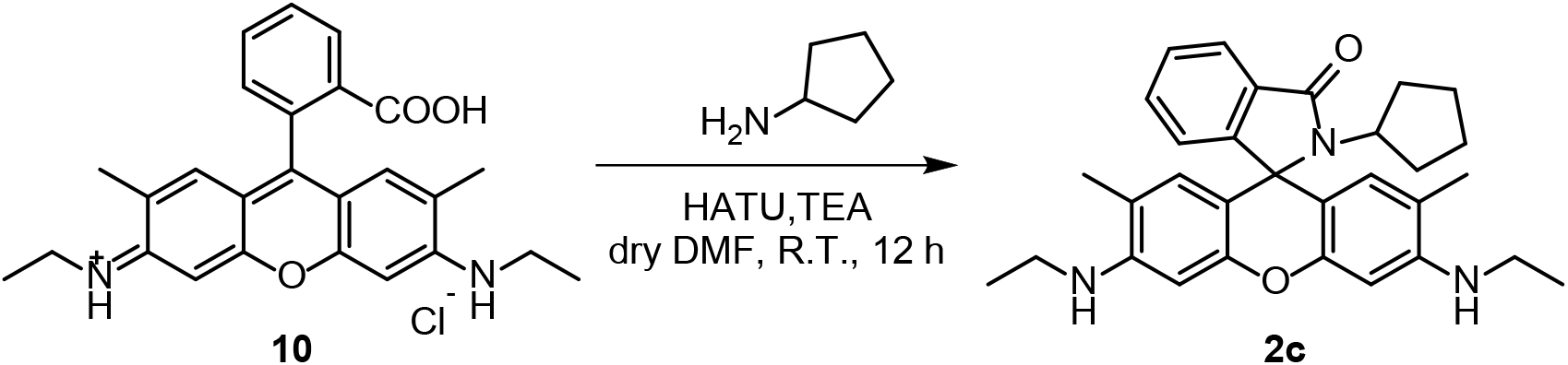

The synthesis of **2c** was similar with **2a**. **Compound 10** (0.1121 g, 0.25 mmol, 1 eq), HATU (0.1490 g, 0.39 mmol, 1.6 eq), TEA (52 μL, 1.5 eq) and cyclopentylamine (36.8 μL, 0.38 mmol, 1.5 eq). **2c** was obtained as a faint yellow solid (64.4 mg, 53.4%). ^1^H NMR (400 MHz, CDCl_3_) δ 7.85 (d, *J* = 7.1 Hz, 1H), 7.42 – 7.31 (m, 2H), 6.94 (d, *J* = 7.2 Hz, 1H), 6.40 – 6.32 (m, 4H), 3.49 (s, 2H), 3.38 – 3.27 (m, 1H), 3.21 (q, *J* = 7.0 Hz, 4H), 2.25 – 2.10 (m, 2H), 1.91 (s, 6H), 1.72 (d, *J* = 22.5 Hz, 4H), 1.32 (t, *J* = 7.1 Hz, 8H).

### 3.13 Synthesis of **2d**

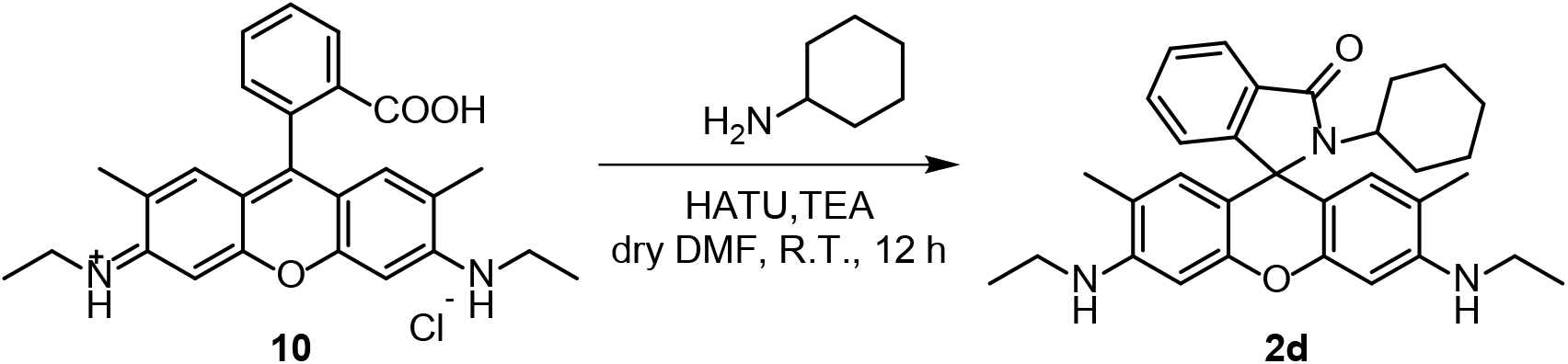

The synthesis of **2d** was similar with **2a**. **Compound 10** (0.1015 g, 0.23 mmol, 1 eq), HATU (0.1293 g, 0.34 mmol, 1.5 eq), TEA (64 μL, 2 eq) and cyclohexylamine (52.4 μL, 0.61 mmol, 2.6 eq). **2d** was obtained as a pink solid (64.4 mg, 53.4%). ^1^H NMR (400 MHz, CDCl_3_) δ 7.88 (d, *J* = 7.5 Hz, 1H), 7.50 – 7.34 (m, 2H), 6.95 (d, *J* = 6.9 Hz, 1H), 6.34 (s, 2H), 6.29 (s, 2H), 3.56 (s, 2H), 3.22 (q, *J* = 6.9 Hz, 4H), 2.90 (ddd, *J* = 12.1, 8.5, 3.7 Hz, 1H), 2.20 – 2.01 (m, 2H), 1.90 (s, 6H), 1.58 – 1.51 (m, 2H), 1.44 (d, *J* = 7.5 Hz, 1H), 1.33 (t, *J* = 7.1 Hz, 6H), 1.17 (d, *J* = 12.8 Hz, 2H), 1.12 – 1.03 (m, 1H), 0.99 – 0.89 (m, 2H).

### 3.14 Synthesis of **2e**

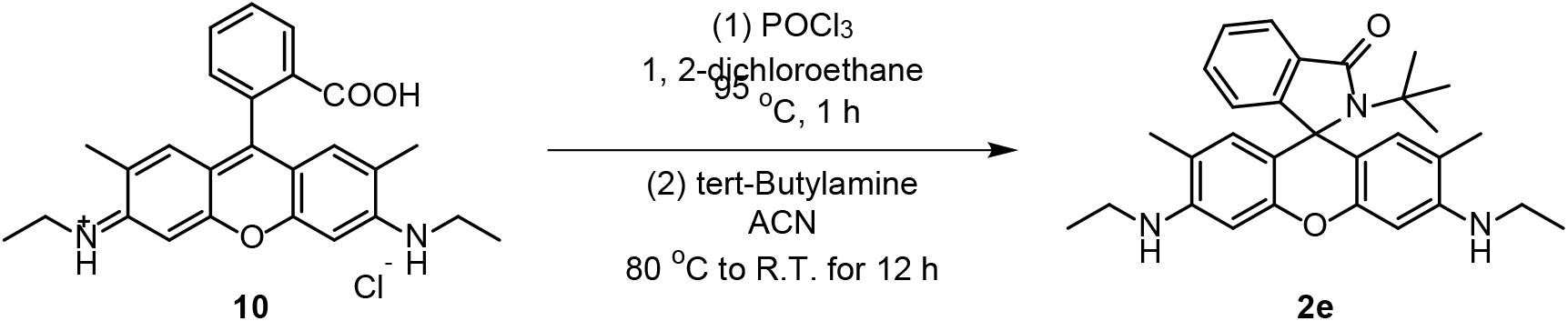

To the suspension of **compound 10** (0.0516 g, 0.11 mmol, 1 eq) in 1,2-dichloroethane (5 mL) was added phosphorus oxychloride (0.5 mL, 5.36 mmol, 48.7 eq). The reaction was refluxed at 95 °C for 1 h. After cooling to room temperature, the solvent was removed under reduced pressure. To the residue dissolved in acetonitrile (5 mL), tert-butylamine (1 mL, 9.43 mmol, 85.7 eq) was added dropwise. The reaction was refluxed at 80 °C for 1h and then cooled to room temperature and stirred for 12 h. After removing the solvent under reduced pressure, the residue was purified by alkaline alumina column chromatography (petroleum ether/ethyl acetate=5:1, v/v) to afford a white solid (3.7 mg, 7.1%). ^1^H NMR (700 MHz, DMSO-*d6*) δ 7.67 (d, *J* = 7.6 Hz, 1H), 7.43 – 7.32 (m, 2H), 6.74 (d, *J* = 6.9 Hz, 1H), 6.23 (s, 4H), 5.01 (t, *J* = 5.1 Hz, 2H), 3.17 – 3.06 (m, 4H), 1.89 (s, 6H), 1.22 (t, *J* = 7.1 Hz, 6H), 1.18 (s, 9H). ^13^C NMR (101 MHz, CDCl_3_) δ 169.80, 155.49, 150.13, 147.09, 132.21, 130.01, 128.55, 127.38, 123.02, 122.22, 117.44, 108.82, 96.78, 65.75, 56.93, 38.43, 28.24, 16.68, 14.78. HRMS (ESI) calcd for C_30_H_36_N_3_O_2_ [M+H]^+^ 470.2808, found 470.2811.

### 3.15 Synthesis of **2f**

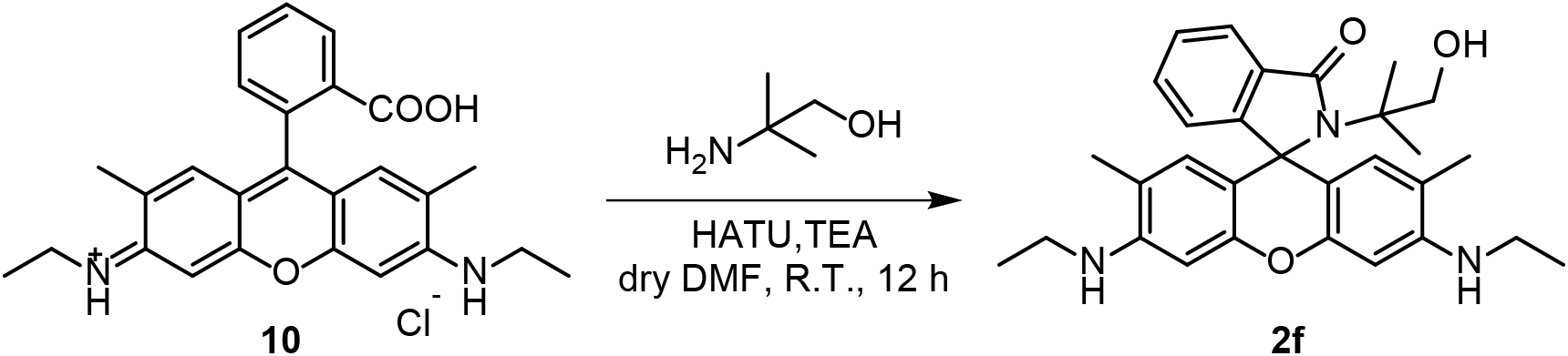

The synthesis of **2f** was similar with **2b**. **Compound 10** (0.1093 g, 0.24 mmol, 1 eq), HATU (0.1539 g, 0.40 mmol, 1.7 eq), TEA (72 μL, 2.2 eq) and 2-amino-2-methyl-1-propanol (50 μL, 0.52 mmol, 2.2 eq). **2f** was obtained as a white solid (55.5 mg, 43.9%). ^1^H NMR (400 MHz, DMSO-*d6*) δ 7.69 – 7.62 (m, 1H), 7.41 – 7.33 (m, 2H), 6.76 – 6.69 (m, 1H), 6.26 (s, 2H), 6.22 (s, 2H), 4.99 (t, *J* = 5.3 Hz, 2H), 4.66 (t, *J* = 6.1 Hz, 1H), 3.60 (d, *J* = 6.1 Hz, 2H), 3.17 – 3.06 (m, 4H), 1.88 (s, 6H), 1.21 (t, *J* = 7.1 Hz, 6H), 1.04 (s, 6H). ^13^C NMR (101 MHz, DMSO-*d6*) δ 169.09, 155.79, 149.98, 147.85, 132.89, 129.79, 128.51, 128.04, 123.49, 122.29, 118.09, 107.65, 96.18, 65.92, 65.62, 60.97, 37.99, 22.64, 17.42, 14.66. HRMS (ESI) calcd for C_30_H_36_N_3_O_3_ [M+H]^+^ 486.2757, found 486.2748.

### 3.16 Synthesis of **3a**

**Compound 11** was synthesized according to literature.^[5] 1^H NMR (400 MHz, CDCl_3_) δ 7.99 (d, *J* = 7.4 Hz, 1H), 7.63 (td, *J* = 7.4, 1.3 Hz, 1H), 7.58 (td, *J* = 7.4, 3.7 Hz, 1H), 7.17 (d, *J* = 7.5 Hz, 1H), 6.55 (d, *J* = 8.6 Hz, 2H), 6.20 (d, *J* = 2.3 Hz, 2H), 6.08 (dd, *J* = 8.6, 2.3 Hz, 2H), 3.90 (t, *J* = 7.3 Hz, 8H), 2.45 – 2.29 (m, 4H).

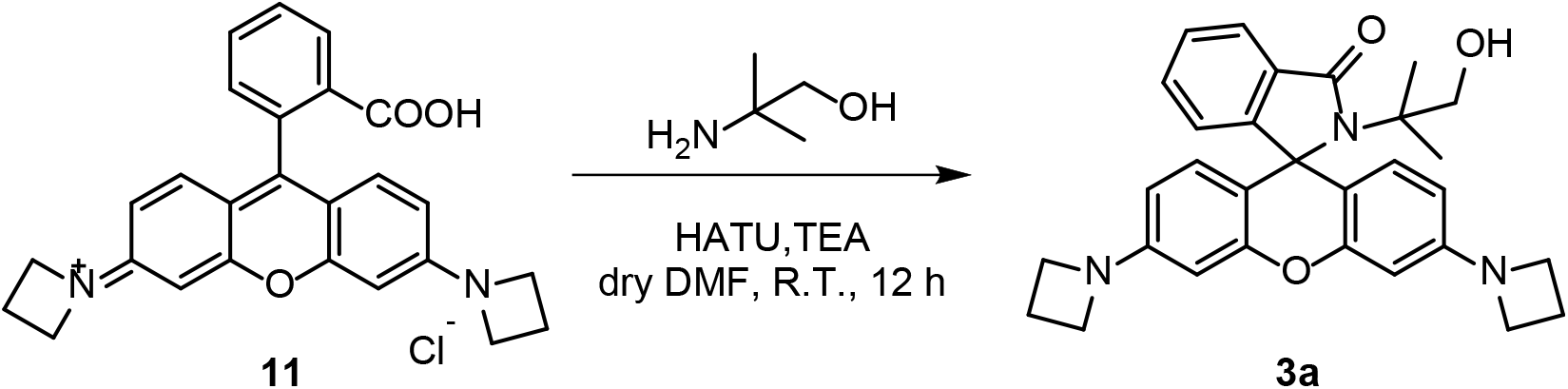

A vial was charged with **compound 11** (0.0208 g, 0.051 mmol, 1 eq) and HATU (0.0296 g, 0.078 mmol, 1.5 eq). The vial was sealed and evacuated/backfilled with nitrogen (3×). Dry DMF (5 mL) and TEA (100 μL, 14 eq) was added. The reaction was stirred at room temperature for 15 min. Following the addition of 2-amino-2-methyl-1-propanol (10 μL, 0.105 mmol, 2 eq), the reaction was stirred under room temperature for 12 h. After removing the solvent under reduced pressure, the residue was purified by silica gel column chromatography (petroleum ether/ethyl acetate=1:1, v/v) to afford a white solid (7.2 mg, 30.4%). ^1^H NMR (400 MHz, CDCl_3_) δ 7.81 (dd, *J* = 5.7, 3.0 Hz, 1H), 7.36 (dd, *J* = 5.7, 3.1 Hz, 2H), 6.85 – 6.74 (m, 1H), 6.57 (d, *J* = 8.5 Hz, 1H), 6.12 (d, *J* = 2.2 Hz, 1H), 6.05 (dd, *J* = 8.5, 2.2 Hz, 1H), 5.35 (t, *J* = 6.2 Hz, 1H), 3.99 – 3.80 (m, 8H), 3.58 (d, *J* = 6.2 Hz, 2H), 2.45 – 2.25 (m, 4H), 1.10 (s, 6H). ^13^C NMR (101 MHz, CDCl_3_) δ 154.70, 151.69, 150.34, 131.84, 127.67, 126.76, 122.12, 121.53, 107.87, 106.31, 96.76, 72.79, 65.56, 60.93, 51.01, 28.68, 22.41, 15.68. HRMS (ESI) calcd for C_30_H_32_N_3_O_3_ [M+H]^+^ 482.2444, found 482.2443.

### 3.17 Synthesis of **O-halo-JF549**

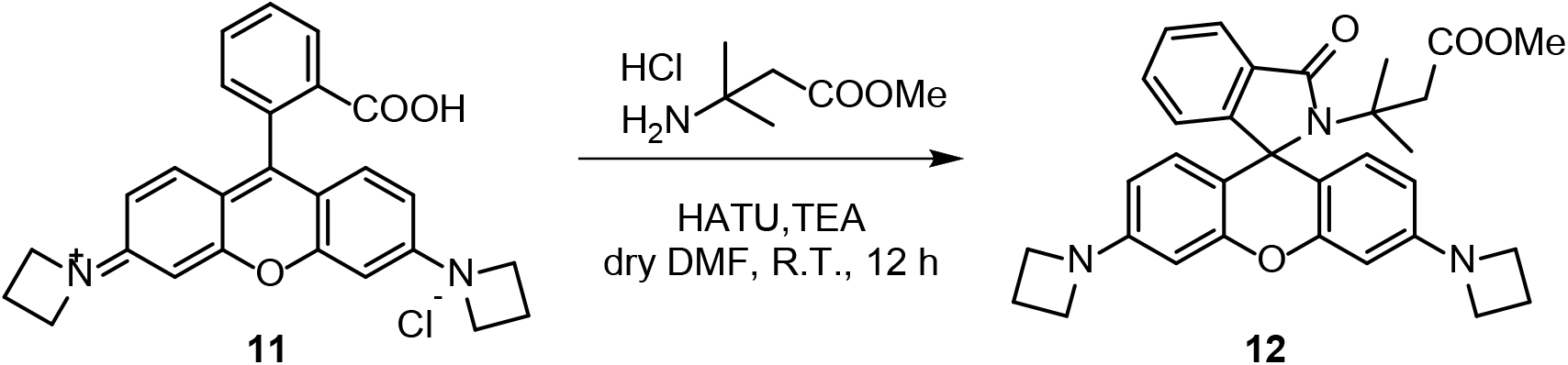

To a solution of **compound 11** (0.0946 g, 0.21 mmol, 1 eq) and HATU (0.1184 g, 0.31 mmol, 1.5 eq) in dry DMF (10 mL) was added TEA (100 μL, 3.4 eq). Following the addition of methyl 3-amino-3-methylbutanoate hydrochloride (0.0374 g, 0.28 mmol, 1.3 eq), the reaction was stirred under room temperature for 12 h. After removing the solvent under reduced pressure, the residue was purified by silica gel column chromatography (petroleum ether/ethyl acetate =3:1, v/v) to afford a white solid (20.6 mg, 18%). ^1^H NMR (700 MHz, CDCl_3_) δ 7.81 (dd, *J* = 5.9, 2.6 Hz, 1H), 7.33 (dd, *J* = 5.5, 3.0 Hz, 2H), 6.81 – 6.76 (m, 1H), 6.55 (d, *J* = 8.5 Hz, 2H), 6.11 (d, *J* = 2.1 Hz, 2H), 6.04 (dd, *J* = 8.5, 2.1 Hz, 2H), 3.96 – 3.82 (m, 8H), 3.55 (s, 3H), 3.12 (s, 2H), 2.42 – 2.31 (m, 4H), 1.28 (s, 6H). HRMS (ESI) calcd for C_32_H_34_N_3_O_4_ [M+H]^+^ 524.2549, found 524.2574.

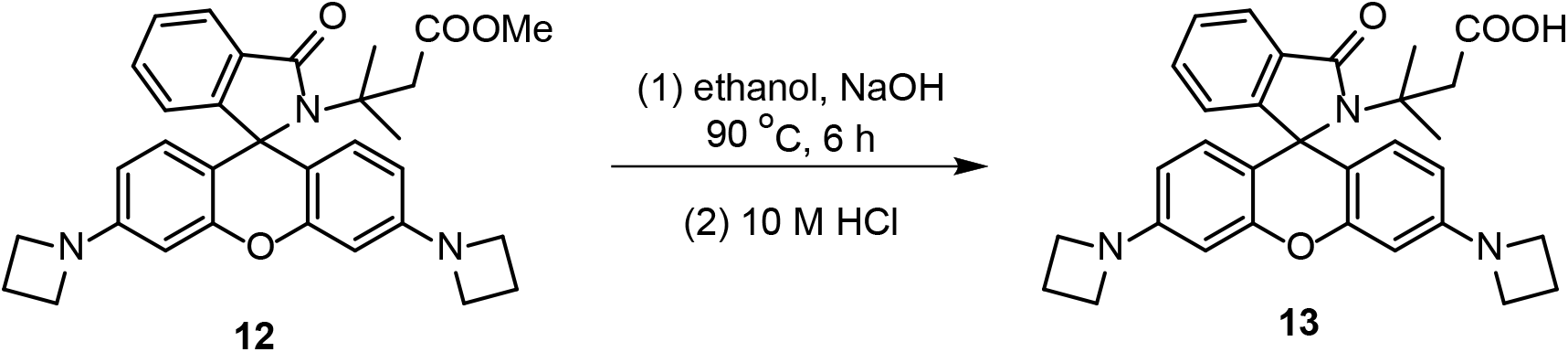

To the solution of **compound 12** (0.011 g, 0.021 mmol, 1 eq) in ethanol (2 mL) was added NaOH (0.1130 g, 2.8 mmol). The mixture was refluxed for 6 h. After cooling to room temperature, the mixture was neutralized with concentrated hydrochloric acid. After removing the solvent under reduced pressure, the residue was purified by silica gel column chromatography dichloromethane/methanol=20:1, v/v) to afford a white solid (9.6 mg, 90%). HRMS (ESI) calcd for C_31_H_32_N_3_O_4_ [M+H]^+^ 510.2393, found 510.2430.

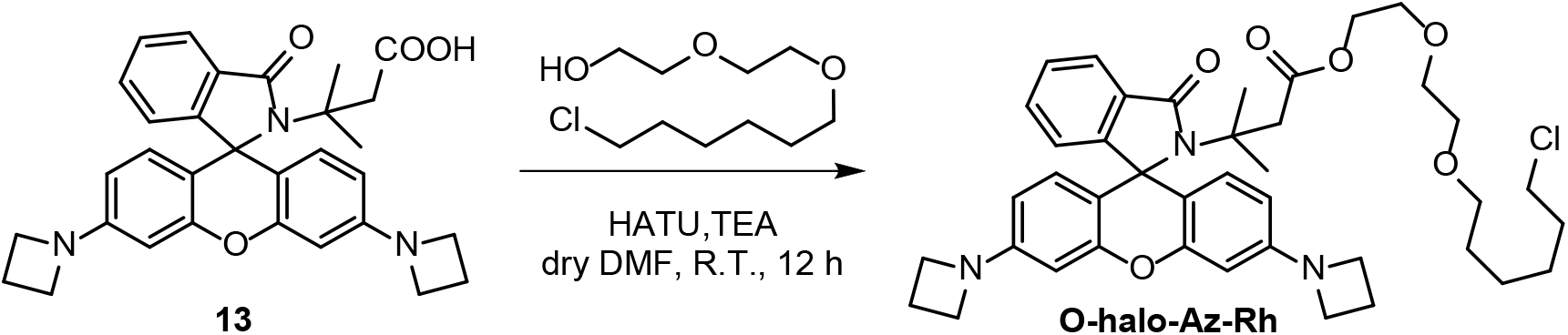

To a suspension of **compound 13** (0.0400 g, 0.078 mmol, 1.0 eq), HATU (0.0358 g, 0.094 mmol, 1.0 eq) and TEA (50 μL, 4.6 eq) in dry DMF (5 mL) was added 2-(2-((6-chlorohexyl)oxy)ethoxy)ethan-1-ol (0.0158 g, 0.07 mmol, 1 eq). The reaction was stirred at room temperature for 12 h. After removing the solvent under reduced pressure, the residue was purified by alkaline alumina column chromatography (petroleum ether/ethyl acetate=5:1, v/v) to afford a white solid (3.5 mg, 6.2%). ^1^H NMR (700 MHz, CDCl_3_) δ 7.80 (dd, *J* = 6.0, 2.6 Hz, 1H), 7.33 (d, *J* = 8.6 Hz, 2H), 6.79 (dd, *J* = 5.9, 2.5 Hz, 1H), 6.56 (d, *J* = 8.5 Hz, 2H), 6.11 (d, *J* = 2.2 Hz, 2H), 6.04 (dd, *J* = 8.5, 2.2 Hz, 2H), 4.14 – 4.09 (m, 2H), 3.94 – 3.82 (m, 8H), 3.70 – 3.36 (m, 10H), 3.12 (s, 2H), 2.37 (p, *J* = 7.2 Hz, 4H), 1.80 – 1.74 (m, 2H), 1.62 – 1.53 (m, 2H), 1.44 (dt, *J* = 15.1, 7.4 Hz, 2H), 1.39 – 1.32 (m, 2H), 1.29 (s, 6H). HRMS (ESI) calcd for C_41_H_51_ClN_3_O_6_ [M+H]^+^ 716.3466, found 716.3462.

### 3.18 Synthesis of **4a**

Compound **14** was synthesized according to literature.^[6]^

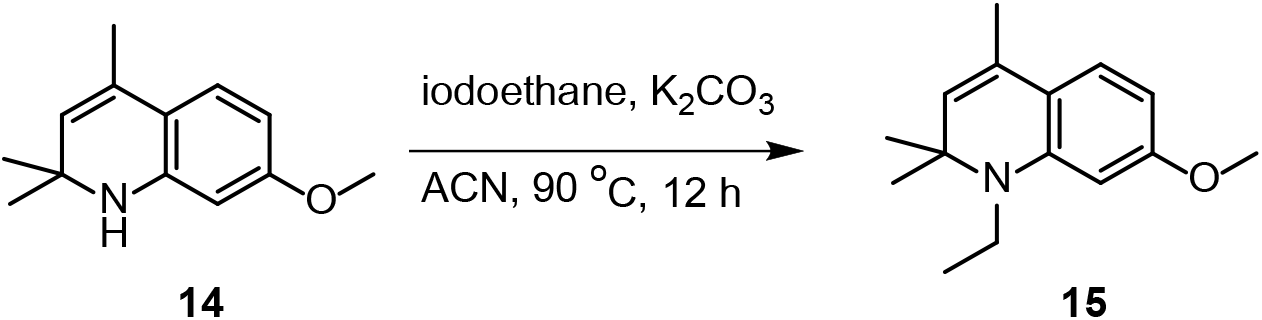

Iodoethane (2.00 mL) was added to a solution of **14** (2.00 g, 10.0 mmol) and K_2_CO_3_ (2.12 g, 20 mmol) in 30 mL ACN. The mixture was heated to 90°C and stirred for 12 h. The solvent was removed under reduced pressure and the residue was further purified by silica gel chromatography (PE) to afford **15** as colorless liquid 2.20 g, yield 95%. ^1^H NMR (400 MHz, CDCl_3_) δ 6.96 (d, *J* = 8.3 Hz, 1H), 6.21 – 6.12 (m, 1H), 6.07 (d, *J* = 2.3 Hz, 1H), 5.09 (d, *J* = 1.1 Hz, 1H), 3.78 (s, 3H), 3.29 (q, *J* = 7.0 Hz, 2H), 1.94 (d, *J* = 1.2 Hz, 3H), 1.30 (s, 6H), 1.20 (t, *J* = 7.0 Hz, 3H).

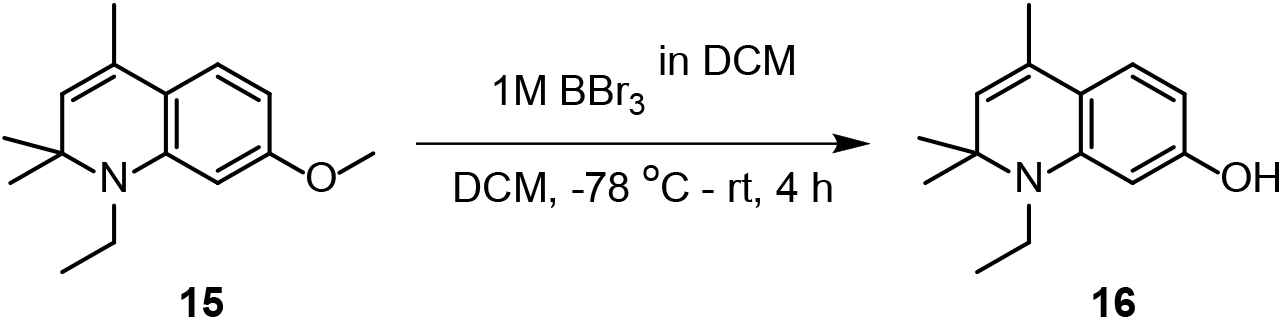

Compound **15** (2.0 g, 8.66 mmol) was dissolved in 20 mL DCM and the solution was cooled to -78°C under an nitrogen atmosphere. Then 9 mL BBr_3_ (1M in DCM) was slowly added to the mixture over 20 min. The reaction mixture was next removed to room temperature and stirred for 4 h. The reaction was quenched by 2 mL deionized water and washed with ammonium chloride solution (50 mL × 3). The organic phase was collected and dried by Na_2_SO_4_. The solvent was removed under reduced pressure to obtain **16** as brown solid 1.32 g, yield 70%.^1^H NMR (400 MHz, CDCl_3_) δ 6.88 (d, *J* = 7.9 Hz, 1H), 6.05 (dt, *J* = 6.6, 2.3 Hz, 2H), 5.07 (d, *J* = 1.3 Hz, 1H), 3.27 (q, *J* = 7.1 Hz, 2H), 1.92 (d, *J* = 1.3 Hz, 3H), 1.29 (s, 6H), 1.17 (t, *J* = 7.1 Hz, 3H). ^13^C NMR (101 MHz, CDCl_3_) δ 156.60, 145.27, 127.42, 126.61, 124.73, 116.15, 101.65, 97.75, 56.90, 38.21, 28.67, 18.80, 14.26.

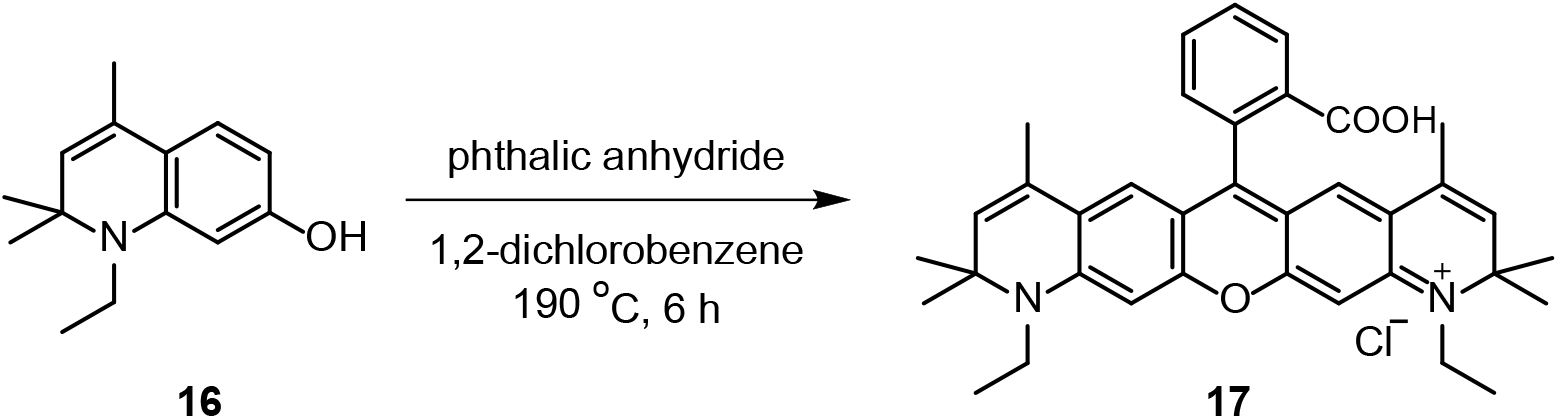

Compound **16** (650 mg, 3.00 mmol) and phthalic anhydride (222 mg, 1.50 mmol) were dissolved in 10 mL 1,2-dichlorobenzene and the solution was heated to 130°C which turned to red liquid. 30 min later, the reaction was heated to 190°C and stirred for 6 h. After the mixture cooled to room temperature, 200 μL concentrated hydrochloric acid was added. Next, the mixture was further purified by silica gel chromatography (MeOH/DCM = 50:1-5:1; V/V) to afforded **17** as a dark blue solid 510 mg, yield 58%.^1^H NMR (400 MHz, MeOD) δ 8.03 (dd, *J* = 6.3, 2.6 Hz, 1H), 7.57 (pd, *J* = 7.5, 3.8 Hz, 2H), 7.23 – 7.11 (m, 1H), 6.82 (s, 2H), 6.68 (s, 2H), 5.48 (s, 2H), 3.61 (q, *J* = 7.0 Hz, 4H), 1.67 (d, *J* = 0.9 Hz, 6H), 1.39 (s, 12H), 1.27 (t, *J* = 7.0 Hz, 6H). ^13^C NMR (101 MHz, MeOD) δ 158.09, 157.29, 152.20, 133.24, 132.62, 131.87, 130.72, 130.58, 130.50, 129.98, 129.94, 125.06, 123.47, 121.80, 113.73, 95.40, 60.08, 39.78, 28.20, 28.17, 16.81, 12.25. HRMS (ESI) m/z Found 547.2969 (M)^+^, calculated 547.2955 for C_36_H_39_N_2_O_3_.

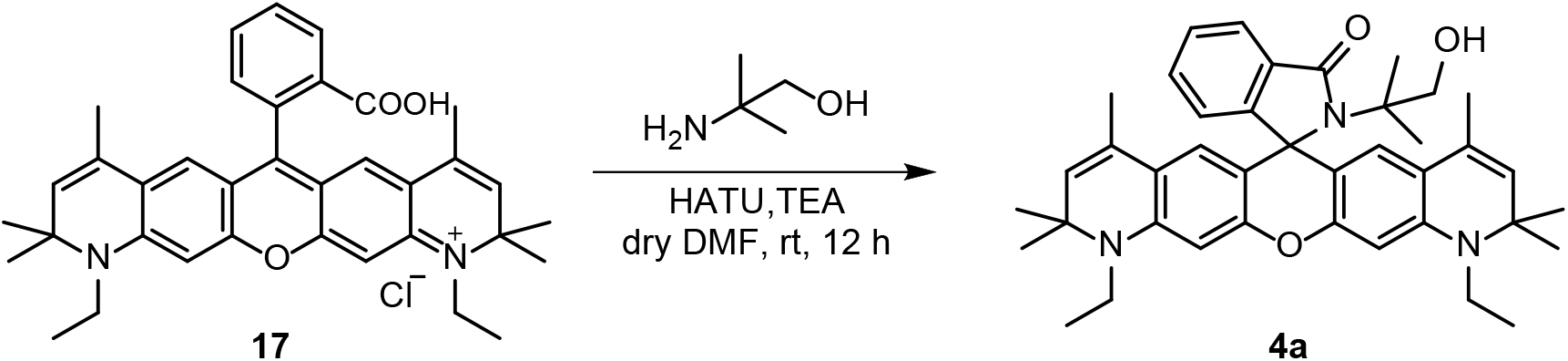

A vial was charged with compound **17** (0.2348 g, 0.40 mmol, 1 eq) and HATU (0.2442 g, 0.64 mmol, 1.6 eq). The vial was sealed and evacuated/backfilled with nitrogen (3×). Dry DMF (5 mL) and TEA (200 μL, 3.6 eq) was added. Following the addition of 2-amino-2-methyl-1-propanol (82 μL, 0.86 mmol, 2.1 eq), the reaction was stirred under room temperature for 12 h. After removing the solvent under reduced pressure, the residue was purified by silica gel column chromatography (petroleum ether/ethyl acetate=5:1, v/v) to afford a white solid (237.6 mg, 96.1%). ^1^H NMR (400 MHz, CDCl_3_) δ 7.81 (dd, *J* = 5.5, 2.8 Hz, 1H), 7.39 – 7.31 (m, 2H), 6.89 (dd, *J* = 5.5, 2.6 Hz, 1H), 6.34 (s, 2H), 6.19 (s, 2H), 5.42 (t, *J* = 6.2 Hz, 1H), 5.16 – 5.06 (m, *J* = 1.0 Hz, 2H), 3.64 (d, *J* = 6.3 Hz, 2H), 3.40 – 3.28 (m, 4H), 1.68 (d, *J* = 1.0 Hz, 6H), 1.35 (s, 6H), 1.29 – 1.23 (m, 12H), 1.18 (s, 6H). ^13^C NMR (101 MHz, CDCl_3_) δ 171.98, 155.57, 151.19, 144.79, 132.75, 128.58, 128.45, 127.60, 126.86, 123.26, 122.46, 119.18, 106.56, 97.27, 73.95, 66.85, 62.00, 57.00, 38.53, 34.67, 28.99, 28.45, 26.92, 25.28, 23.62, 18.61, 14.03. HRMS (ESI) calcd for C_40_H_48_N_3_O_3_ [M+H]^+^ 618.3696, found 618.3685.

### 3.19 Synthesis of **N-halo-JA21**

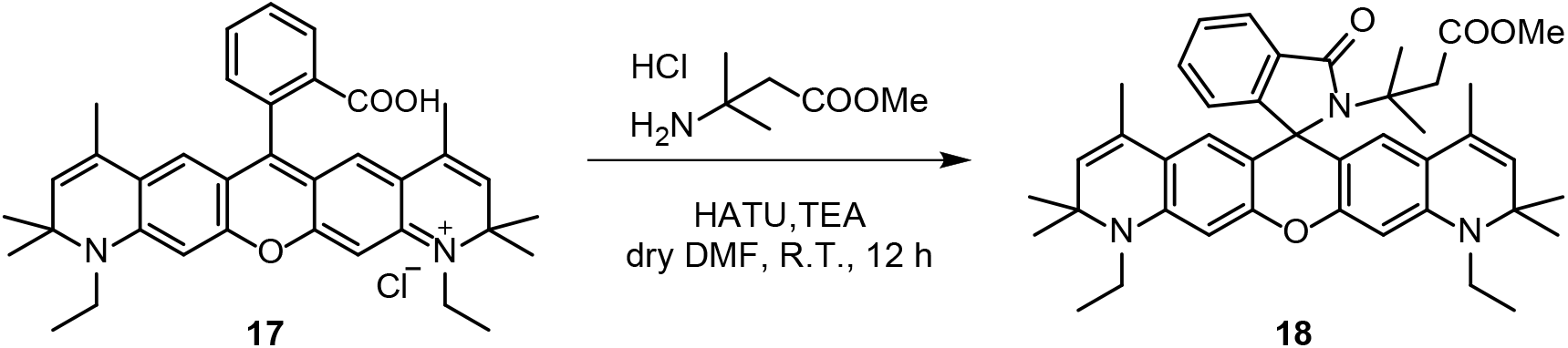

To a solution of compound **17** (0.0994 g, 0.18 mmol, 1 eq) and HATU (0.1204 g, 0.32 mmol, 1.8 eq) in dry DMF (10 mL) was added TEA (100 μL, 4.0 eq). Following the addition of methyl 3-amino-3-methylbutanoate hydrochloride (0.0549 g, 0.42 mmol, 2.3 eq), the reaction was stirred under room temperature for 12 h. After removing the solvent under reduced pressure, the residue was purified by silica gel column chromatography (dichloromethane/methanol=150:1, v/v) to afford a white solid (89.1 mg, 75.0%). ^1^H NMR (400 MHz, CDCl_3_) δ 7.90 (d, *J* = 7.0 Hz, 1H), 7.76 – 7.65 (m, 2H), 7.33 (d, *J* = 7.8 Hz, 1H), 6.83 (s, 2H), 6.71 (s, 2H), 5.47 (s, 2H), 3.70 – 3.57 (m, 7H), 2.49 (s, 2H), 1.81 (s, 6H), 1.49 (s, 6H), 1.45 (s, 6H), 1.39 (t, *J* = 7.0 Hz, 6H), 1.18 (s, 6H). HRMS (ESI) calcd for C_42_H_50_N_3_O_4_ [M]^+^ 660.3801, found 660.3794.

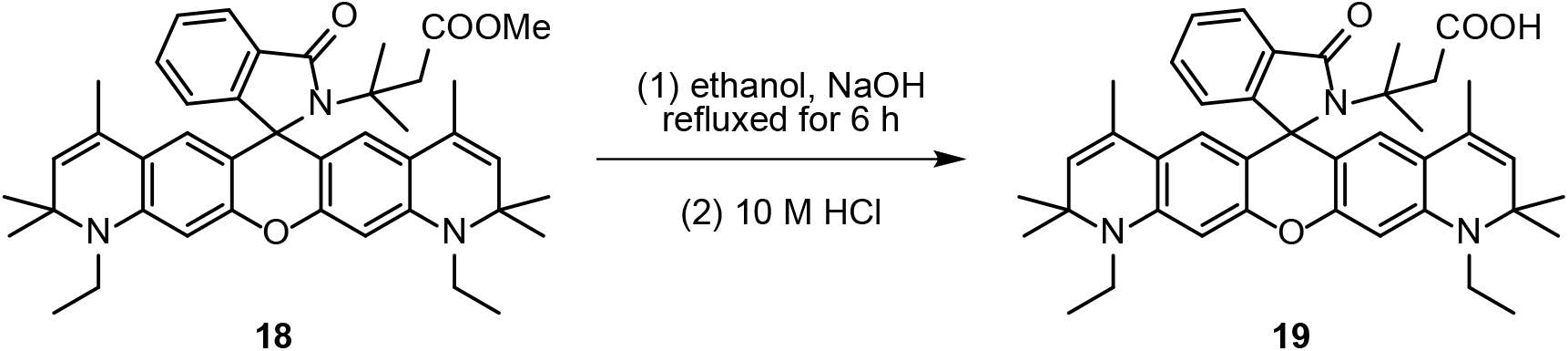

To the solution of **compound 18** (0.0793 g, 0.12 mmol) in ethanol (20 mL) was added NaOH (0.2097 g, 5.2 mmol). The mixture was refluxed for 6 h. After cooling to room temperature, the mixture was neutralized with concentrated hydrochloric acid. After removing the solvent under reduced pressure, the residue was purified by silica gel column chromatography (dichloromethane/methanol=20:1, v/v) to afford a red solid (63.3 mg, 81.6%). ^1^H NMR (400 MHz, CDCl_3_) δ 7.96 (d, *J* = 7.4 Hz, 1H), 7.73 (t, *J* = 7.6 Hz, 1H), 7.67 (t, *J* = 7.3 Hz, 1H), 7.30 (d, *J* = 7.3 Hz, 1H), 6.86 (s, 2H), 6.68 (s, 2H), 5.48 (s, 2H), 3.68 – 3.59 (m, 4H), 2.51 (s, 2H), 1.81 (s, 6H), 1.49 (s, 6H), 1.46 (s, 6H), 1.39 (t, *J* = 6.9 Hz, 6H), 1.22 (s, 6H). HRMS (ESI) calcd for C_42_H_50_N_3_O_4_ [M+H]^+^ 646.3645, found 646.3654.

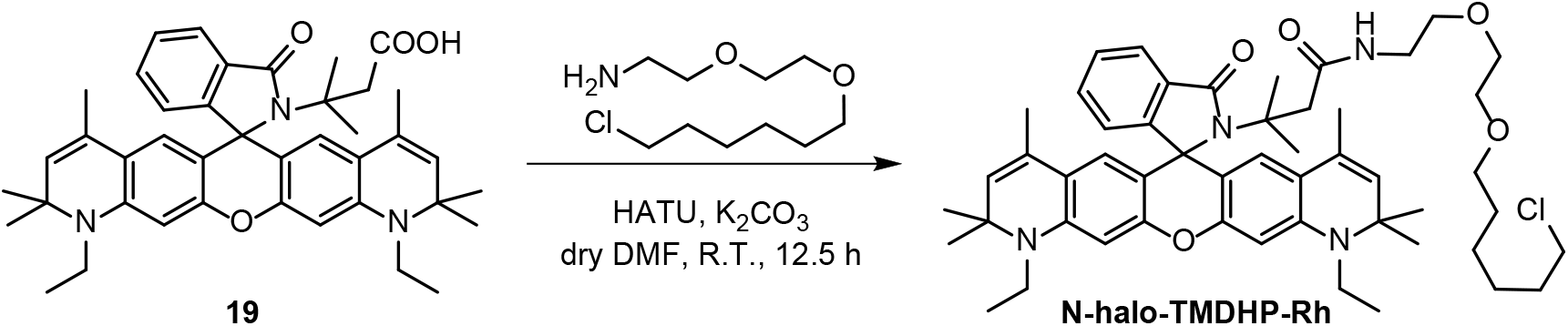

To a suspension of **compound 19** (0.0633 g, 0.098 mmol, 1.0 eq), HATU (0.0224 g, 0.100 mmol, 1.0 eq) and anhydrous potassium carbonate (0.0272 g, 0.197 mmol, 2.0 eq) in dry DMF (5 mL) was added 2-(2-((6-chlorohexyl)oxy)ethoxy)ethan-1-amine (0.0158 g, 0.07 mmol, 1 eq). The reaction was stirred at room temperature for 12.5 h. After removing the solvent under reduced pressure, the residue was purified by alkaline alumina column chromatography (petroleum ether/ethyl acetate=1:1, v/v) to afford a blue solid (2.9 mg, 3.5%). ^1^H NMR (400 MHz, CDCl_3_) δ 8.99 (s, 1H), 8.67 (s, 1H), 7.98 (d, *J* = 7.6 Hz, 1H), 7.69 (t, *J* = 7.0 Hz, 1H), 7.62 (t, *J* = 7.4 Hz, 1H), 7.28 (s, 1H), 6.91 (s, 2H), 6.59 (s, 2H), 5.46 (s, 2H), 3.66 – 3.50 (m, 12H), 3.44 (dd, *J* = 13.6, 6.8 Hz, 4H), 2.41 (s, 2H), 1.86 – 1.72 (m, 8H), 1.57 (d, *J* = 7.3 Hz, 2H), 1.49 (s, 6H), 1.46 (s, 8H), 1.41 – 1.34 (m, 8H), 1.18 (s, 6H). HRMS (ESI) calcd for C_51_H_68_ClN_4_O_5_ [M+H]^+^ 851.4878, found 851.4801.

### 3.20 Synthesis of **21**

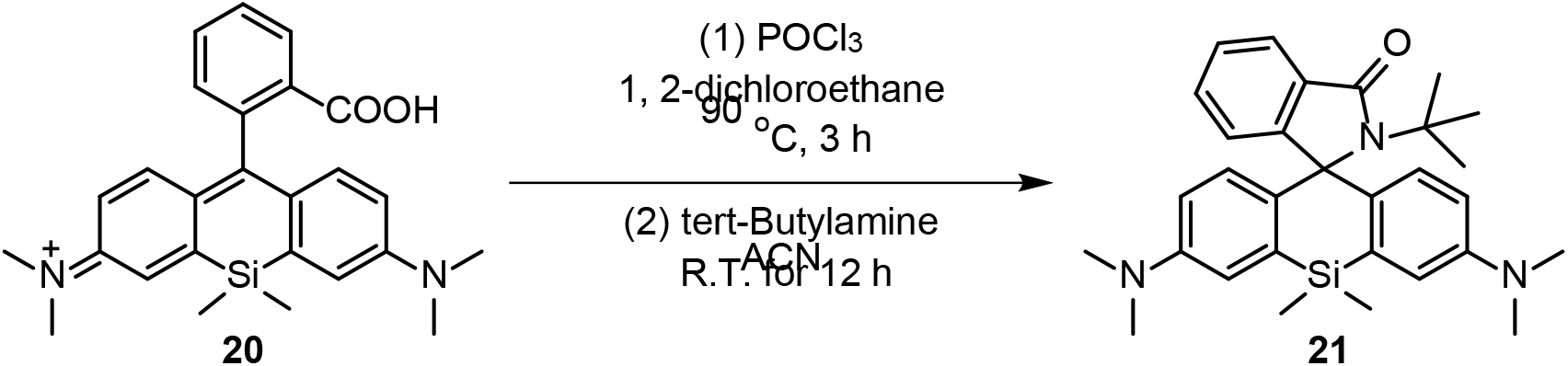

To the suspension of **compound 20** (0.0020 g, 0.047 mmol, 1 eq) in 1,2-dichloroethane (5 mL) was added phosphorus oxychloride (0.7 mL, 7.504 mmol, 160 eq). The reaction was refluxed at 95 °C for 3 h. After cooling to room temperature, the solvent was removed under reduced pressure. To the residue dissolved in acetonitrile (5 mL), tert-butylamine (3 mL, 28.29 mmol, 602 eq) was added dropwise. The reaction was stirred at room temperature for 12 h. After removing the solvent under reduced pressure, the residue was purified by silica gel column chromatography (dichloromethane/methanol=50:1, v/v) to afford a light blue solid (5.2 mg, 22.9%). ^1^H NMR (700 MHz, CDCl_3_) δ 7.79 (d, *J* = 7.5 Hz, 1H), 7.21 (t, *J* = 7.3 Hz, 1H), 7.18 (t, *J* = 7.3 Hz, 1H), 6.88 (d, *J* = 9.0 Hz, 2H), 6.84 (d, *J* = 2.4 Hz, 2H), 6.62 (dd, *J* = 9.0, 2.6 Hz, 2H), 6.55 (d, *J* = 7.5 Hz, 1H), 2.96 (s, 12H), 1.28 (s, 9H), 0.57 (s, 3H), 0.48 (s, 3H). HRMS (ESI) calcd for C_30_H_38_N_3_OSi [M+H]^+^ 484.2779, found 484.2773.

## 4 UV-vis spectra and fluorescence spectra of 1a-1f, 2a-2f, 3a and 4a

### 4.1 UV-vis spectra and fluorescence spectra at different pH

**Figure S1.**
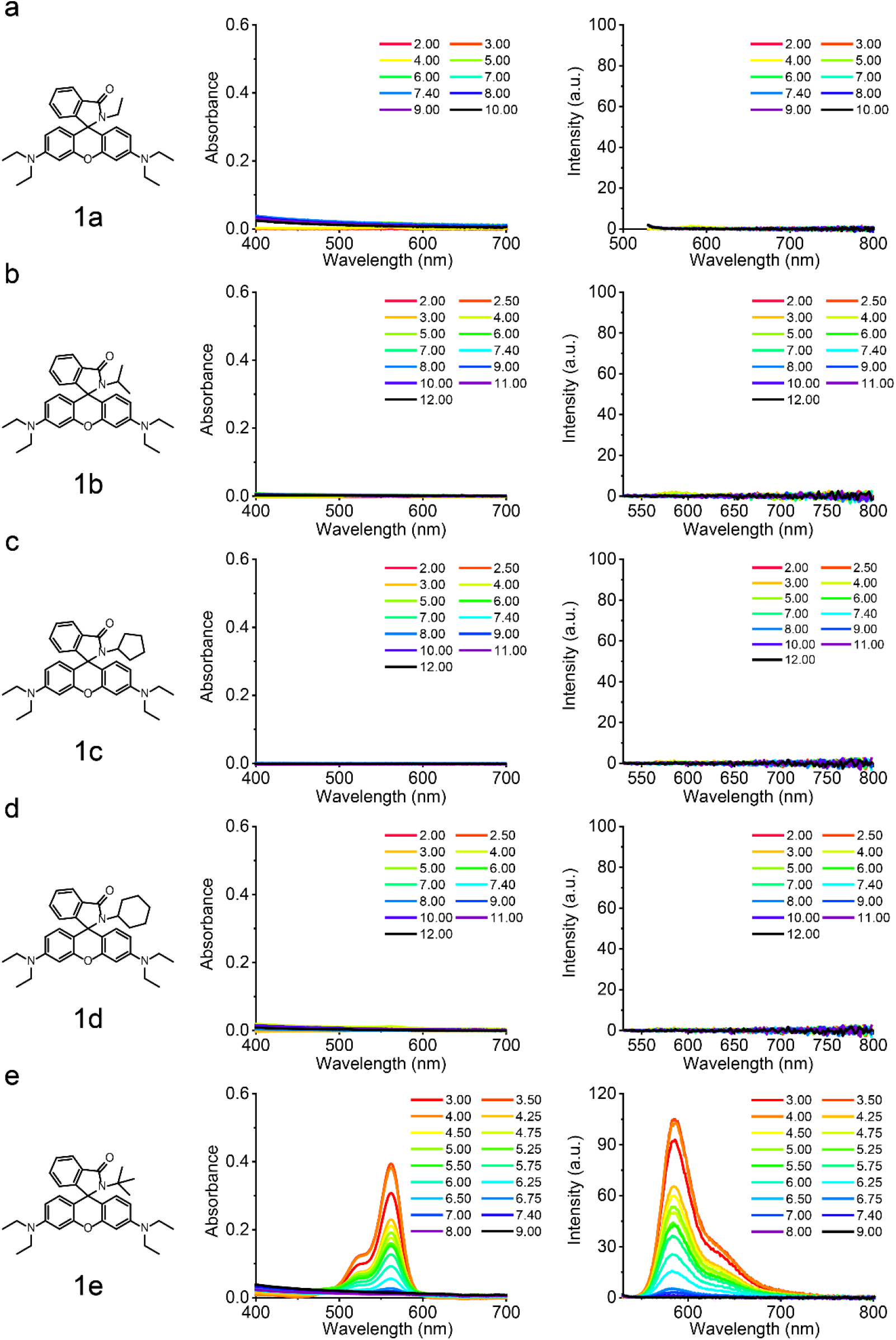
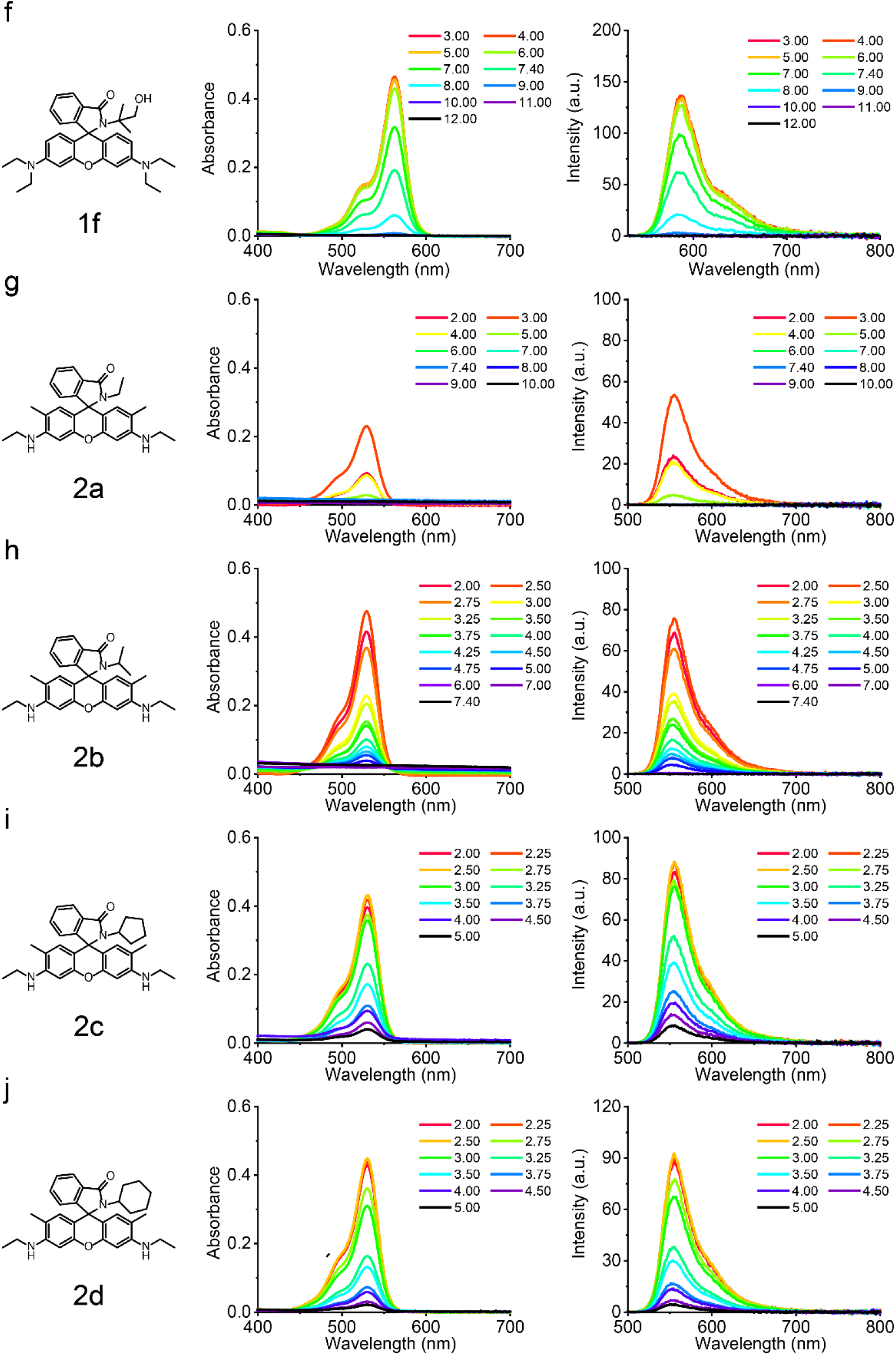
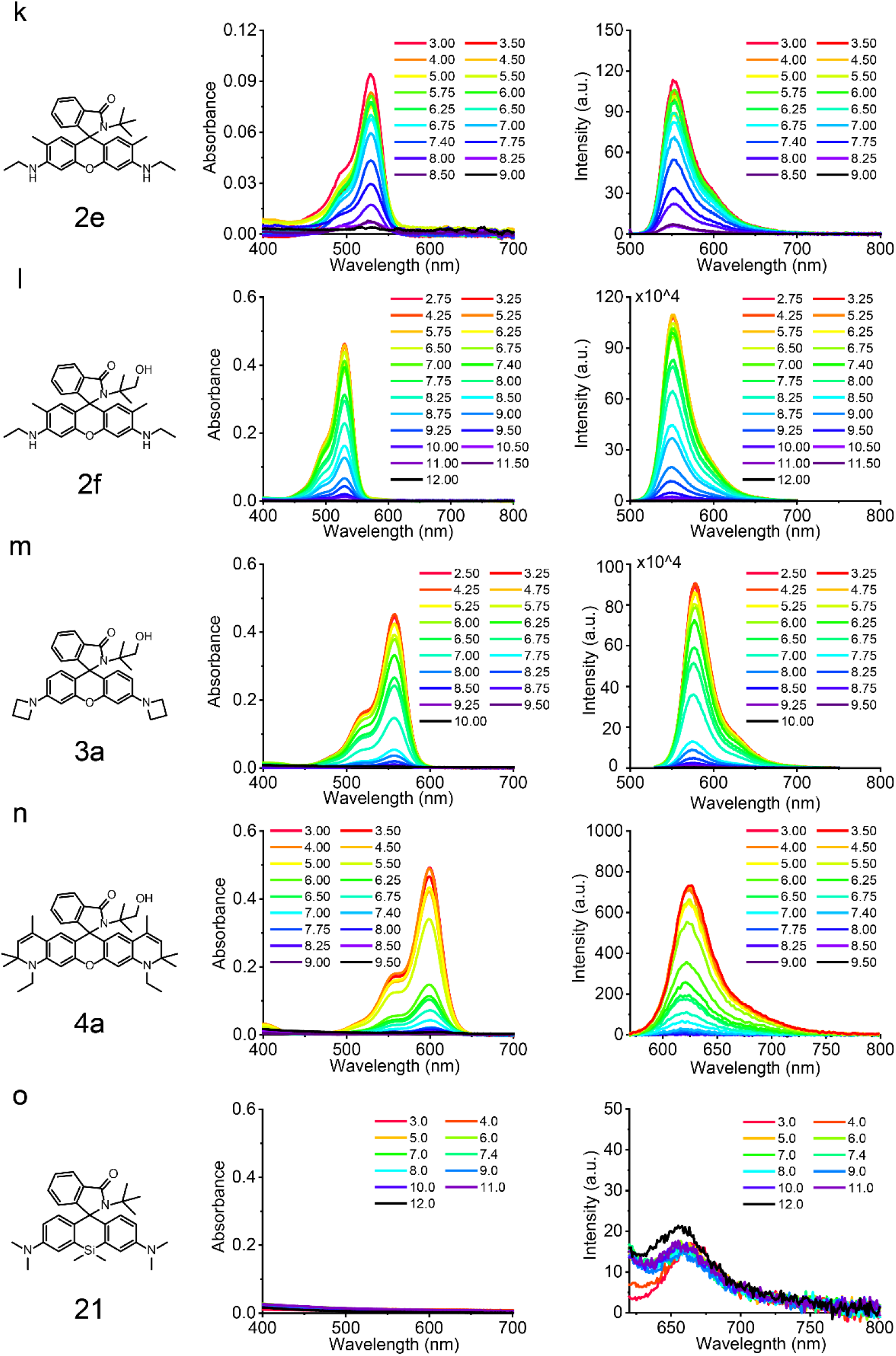
Absorbance (left) and fluorescence (right) spectra of rhodamine derivatives at different pH. Excitation wavelengths for fluorescence spectra were 520 nm, 495 nm, 525 nm, 560 nm, and 600 nm for (a-f), (g-l), (m), (n) and (o).

### 4.2 UV-vis spectra and fluorescence spectra in different dioxane-water mixture

**Figure S2.**
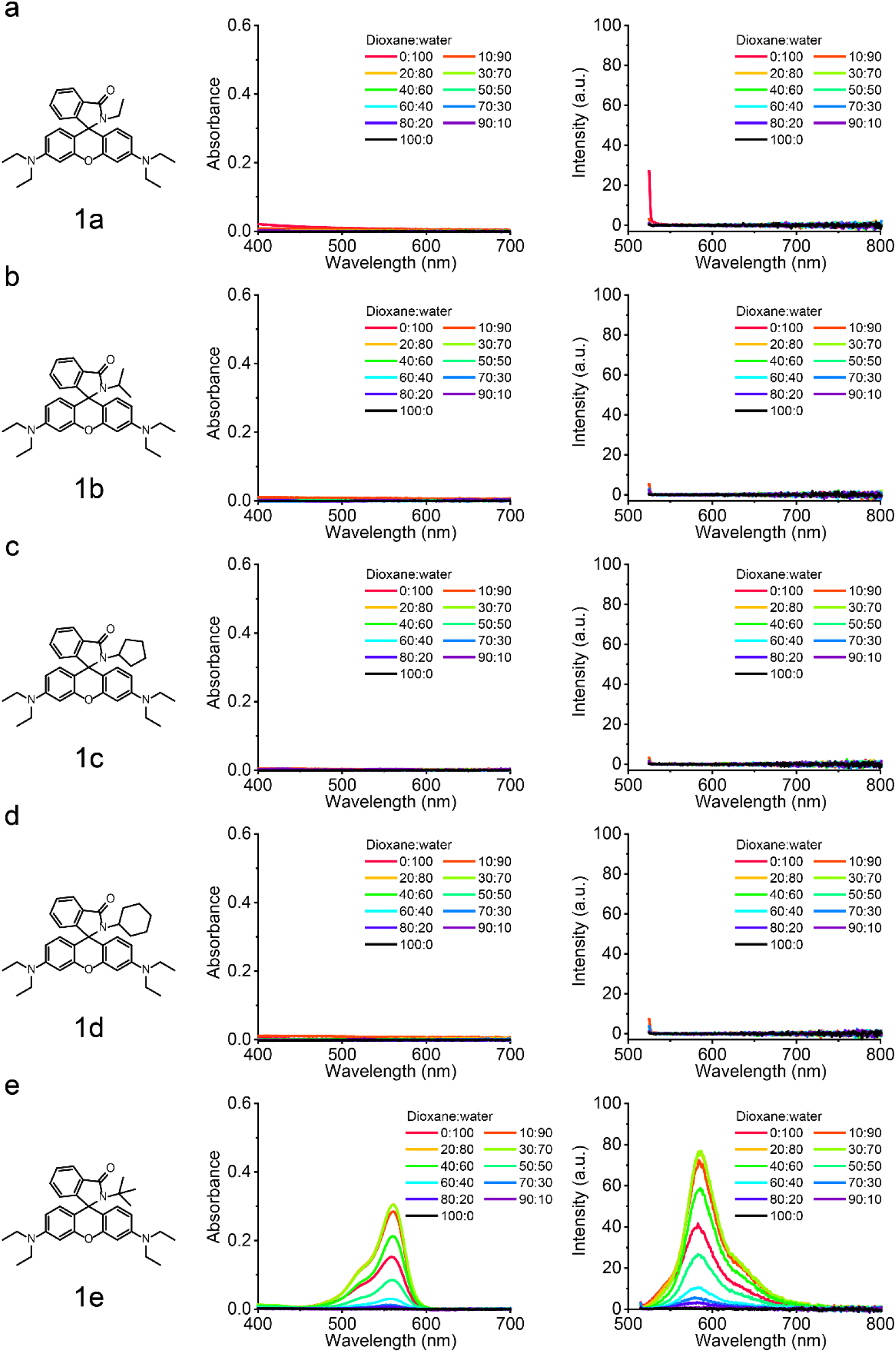
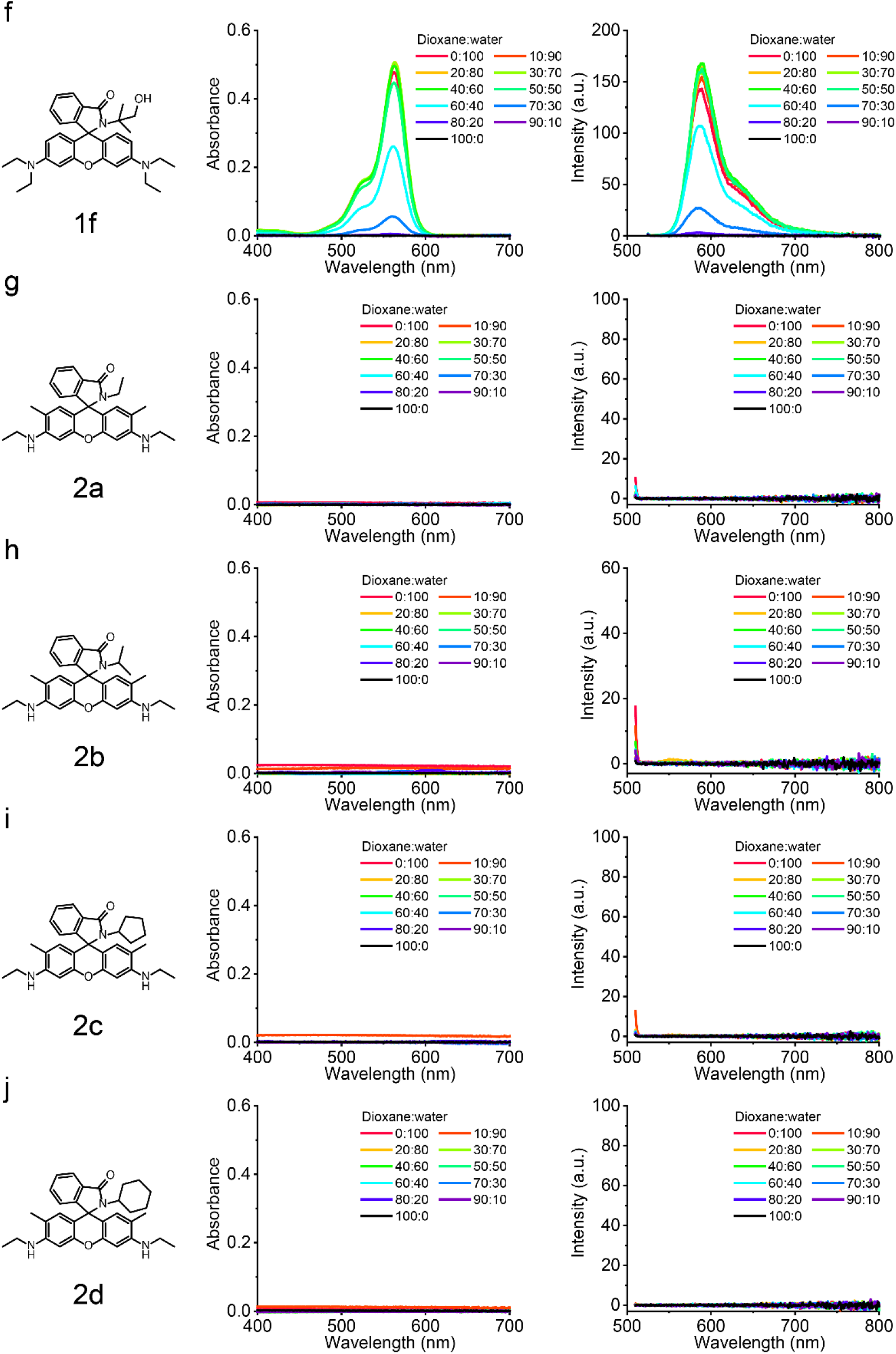
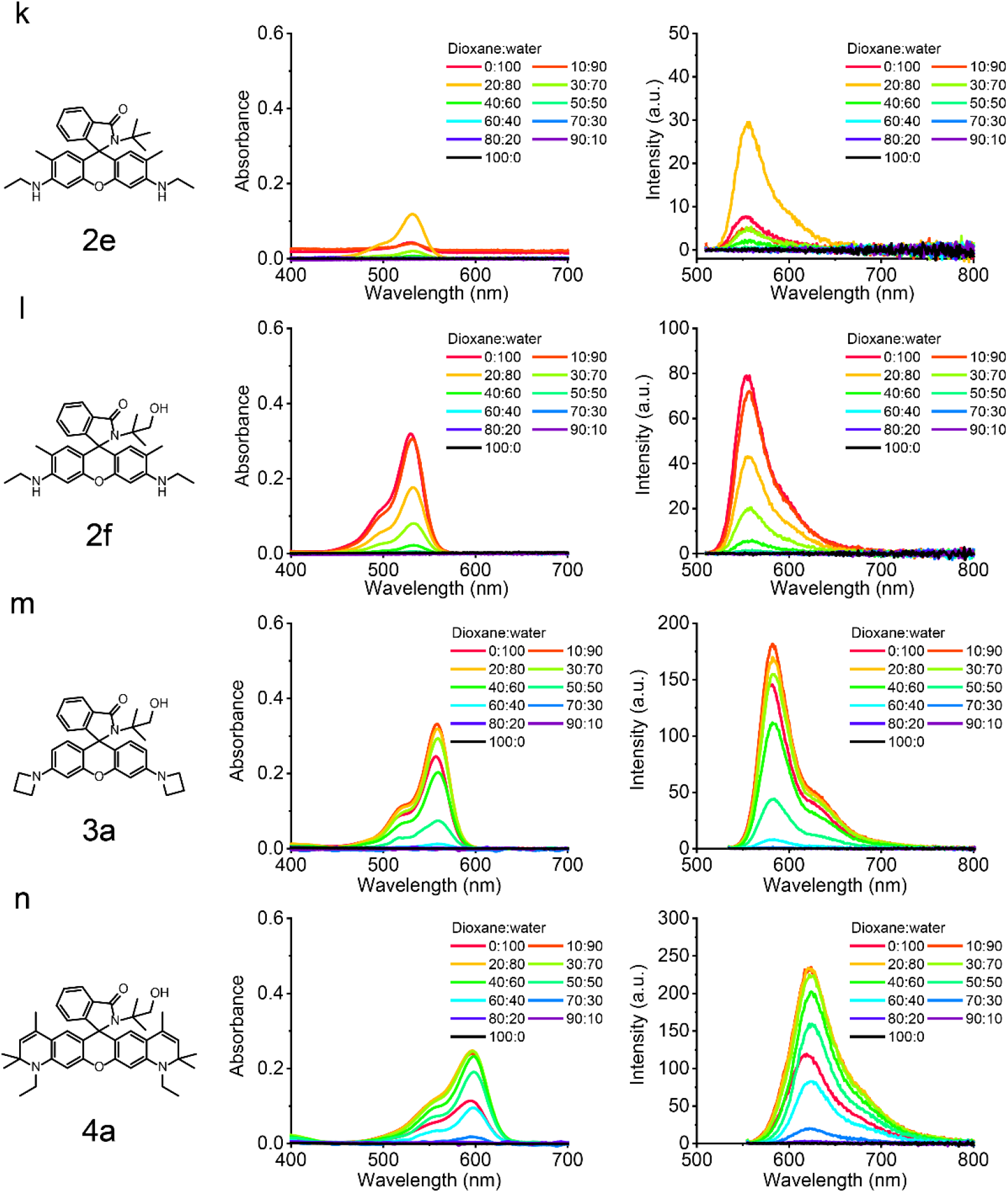
Absorbance (left) and fluorescence (right) spectra of rhodamine derivatives in dioxane-water mixtures. Excitation wavelengths for fluorescence spectra were 520 nm, 510 nm, 505 nm, 530 nm and 550 nm for (a-d), (f), (g-l), (m) and (n).

**Figure S3.**
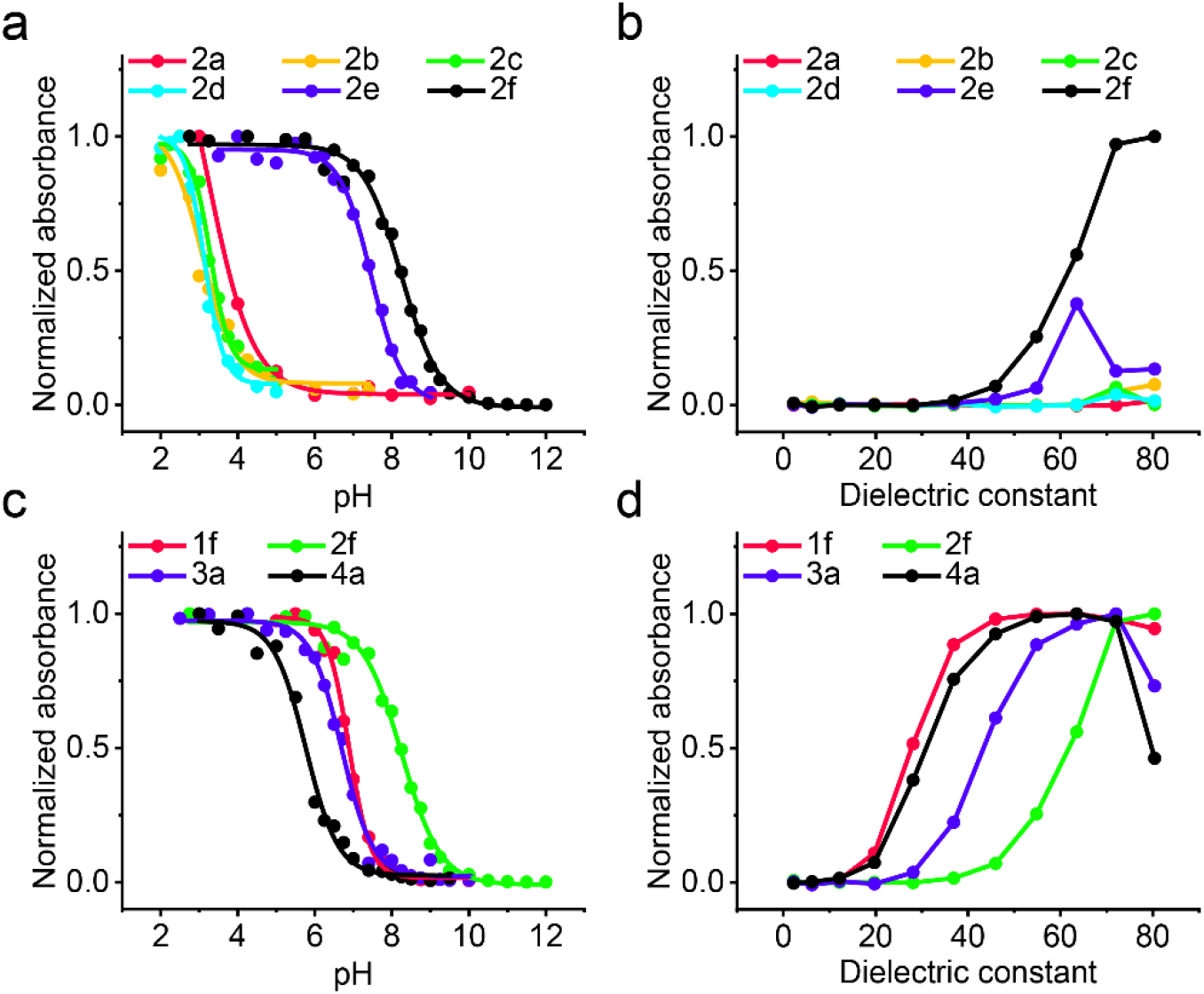
Normalized peak absorbance of rhodamine derivatives. (a, c) Normalized peak absorbance of compound **1f**, **2a**-**2f**, **3a** and **4a** as a function of pH in aqueous solution. (b, d) Normalized peak absorbance of compound **1f**, **2a**-**2f**, **3a** and **4a** as a function of dielectric constant in dioxane-water mixtures.

**Scheme S1.**
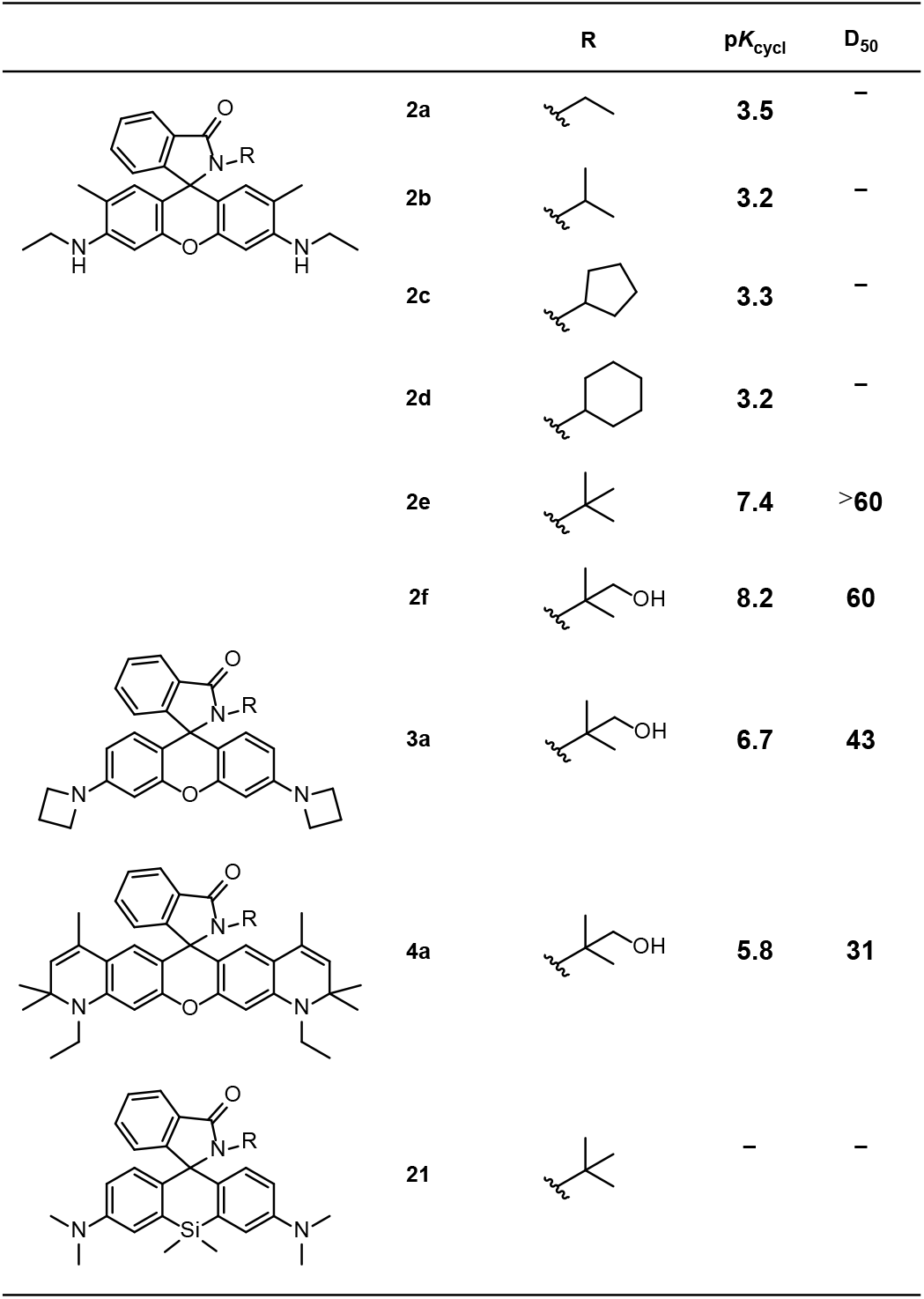
p*K*_cycl_ and D_50_ values of **2a**-**2f**, **3a**, **4a** and **21**.

## 5 Crystallography experiment

### 5.1 Culture of closed form crystals

10 mg **1a/1b/1e/4a** was firstly dissolved in 5 mL DCM. Then petroleum ether was dropwise added to the solution until a slight turbidity appeared. Another 1 mL DCM was next added. The clear solution was further filtered through a 0.22 μm filter. The mixture in 10 mL vial was placed in darkroom with constant temperature of 20 °C.

### 5.2 Culture of open form crystal

10 mg **4a** was firstly dissolved in 5 mL DCM. Then 2 μL trifluoroacetic acid was added to the solution to afford a red solution. Then HPLC acetonitrile was dropwise added to the solution until a slight turbidity appeared. Another 1 mL DCM was next added. The clear solution was further filtered through a 0.22 μm filter. The mixture in 10 mL vial was placed in darkroom with constant temperature of 20 °C.

### 5.3 Summary of crystal data

**Table S1.** Summary of crystal data and intensity collection parameters.

| Crystal data | **1a** | **1b** | **1e** | **4a** closed | **4a** open |
|---|---|---|---|---|---|
| Empirical formula | C<sub>2.4</sub>H<sub>2.64</sub>N<sub>0.24</sub>O<sub>0.16</sub> | C<sub>2.48</sub>H<sub>2.96</sub>N<sub>0.24</sub>O<sub>0.16</sub> | C<sub>2.43</sub>H<sub>2.81</sub>N<sub>0.23</sub>O<sub>0.15</sub> | C<sub>41</sub>H<sub>48.5</sub>N<sub>3.5</sub>O<sub>3</sub> | C<sub>2.67</sub>H<sub>3.05</sub>F<sub>0.19</sub>N<sub>0.19</sub>O<sub>0.32</sub> |
| Formula weight | 37.41 | 38.69 | 37.64 | 638.33 | 46.47 |
| Temperature/K | 293 | 298 | 288.3(6) | 295(1) | 199.98 |
| Crystal system | triclinic | triclinic | monoclinic | triclinic | monoclinic |
| Space group | P-1 | P-1 | P2<sub>1</sub>/c | P-1 | P2<sub>1</sub>/c |
| *a* (Å) | 10.7883(11) | 10.5802(9) | 18.261(2) | 9.4934(2) | 20.147(2) |
| *b* (Å) | 11.9226(12) | 12.0923(9) | 12.1813(13) | 18.7137(4) | 10.8533(9) |
| *c* (Å) | 12.4166(13) | 12.5878(8) | 38.70(4) | 20.4483(5) | 17.792(2) |
| *α* (°) | 82.874(9) | 83.335(6) | 90 | 95.846(2) | 90 |
| *β* (°) | 65.569(10) | 68.195(7) | 97.76(3) | 94.076(2) | 103.435(3) |
| *γ* (°) | 65.336(10) | 64.232(8) | 90 | 90.045(2) | 90 |
| Volume (Å<sup>3</sup>) | 1319.1(3) | 1344.6(2) | 8530(8) | 3604.62(14) | 3784.0(7) |
| *Z* | 25 | 25 | 158 | 4 | 63 |
| *ρ*<sub>calc</sub> (g/cm<sup>3</sup>) | 1.177 | 1.195 | 1.158 | 1.176 | 1.285 |
| *μ* (mm<sup>-1</sup>) | 0.583 | 0.586 | 0.567 | 0.581 | 0.094 |
| F(000) | 500.0 | 520.0 | 3192.0 | 1372.0 | 1552.0 |
| Radiation | CuKα (λ = 1.54184) | CuKα (λ = 1.54184) | CuKα (λ = 1.54184) | CuKα (λ = 1.54178) | MoKα (λ = 0.71073) |
| 2*θ* range for data collection (°) | 7.834 to 134.102 | 7.576 to 134.366 | 7.156 to 136.842 | 6.106 to 134.114 | 4.29 to 50.89 |
| Index ranges | -12 ≤ h ≤ 12, -14 ≤ k ≤ 12, -14 ≤ l ≤ 14 | -12 ≤ h ≤ 12, -14 ≤ k ≤ 14, -14 ≤ l ≤ 15 | -21 ≤ h ≤ 18, -14 ≤ k ≤ 14, -42 ≤ l ≤ 45 | -11 ≤ h ≤ 11, -19 ≤ k ≤ 22, -24 ≤ l ≤ 24 | -24 ≤ h ≤ 24, -12 ≤ k ≤ 13, -21 ≤ l ≤ 21 |
| Reflections collected | 13398 | 13908 | 65967 | 47538 | 34481 |
| Independent reflections | 4653 [R<sub>int</sub> = 0.0484, R<sub>sigma</sub> = 0.0329] | 4772 [R<sub>int</sub> = 0.0332, R<sub>sigma</sub> = 0.0241] | 15084 [R<sub>int</sub> = 0.1129, R<sub>sigma</sub> = 0.0425] | 18360 [R<sub>int</sub> = 0.0575, R<sub>sigma</sub> = 0.0429] | 6963 [R<sub>int</sub> = 0.0798, R<sub>sigma</sub> = 0.0661] |
| Data/restraints/parameters | 4653/0/320 | 4772/0/351 | 15084/0/1021 | 18360/39/1756 | 6963/0/489 |
| Goodness-of-fit on F<sup>2</sup> | 1.048 | 1.019 | 1.508 | 1.055 | 1.111 |
| Final R indexes [I>=2σ (I)] | R<sub>1</sub> = 0.0811, wR<sub>2</sub> = 0.2281 | R<sub>1</sub> = 0.0505, wR<sub>2</sub> = 0.1356 | R1 = 0.1359, wR2 = 0.3918 | R<sub>1</sub> = 0.0611, wR<sub>2</sub> = 0.1610 | R<sub>1</sub> = 0.1190, wR<sub>2</sub> = 0.3449 |
| Final R indexes [all data] | R<sub>1</sub> = 0.1014, wR<sub>2</sub> = 0.2595 | R<sub>1</sub> = 0.0649, wR<sub>2</sub> = 0.1541 | R<sub>1</sub> = 0.1689, wR<sub>2</sub> = 0.4285 | R<sub>1</sub> = 0.0666, wR<sub>2</sub> = 0.1686 | R<sub>1</sub> = 0.1521, wR<sub>2</sub> = 0.3685 |
| Largest diff. peak/hole / e Å<sup>-3</sup> | 0.59/-0.42 | 0.20/-0.19 | 0.67/-0.42 | 0.32/-0.31 | 0.79/-0.58 |

## 6 UV-vis spectra and fluorescence spectra of functionalized rhodamine derivatives

### 6.1 UV-vis spectra and fluorescence spectra at different pH

**Figure. S4.**
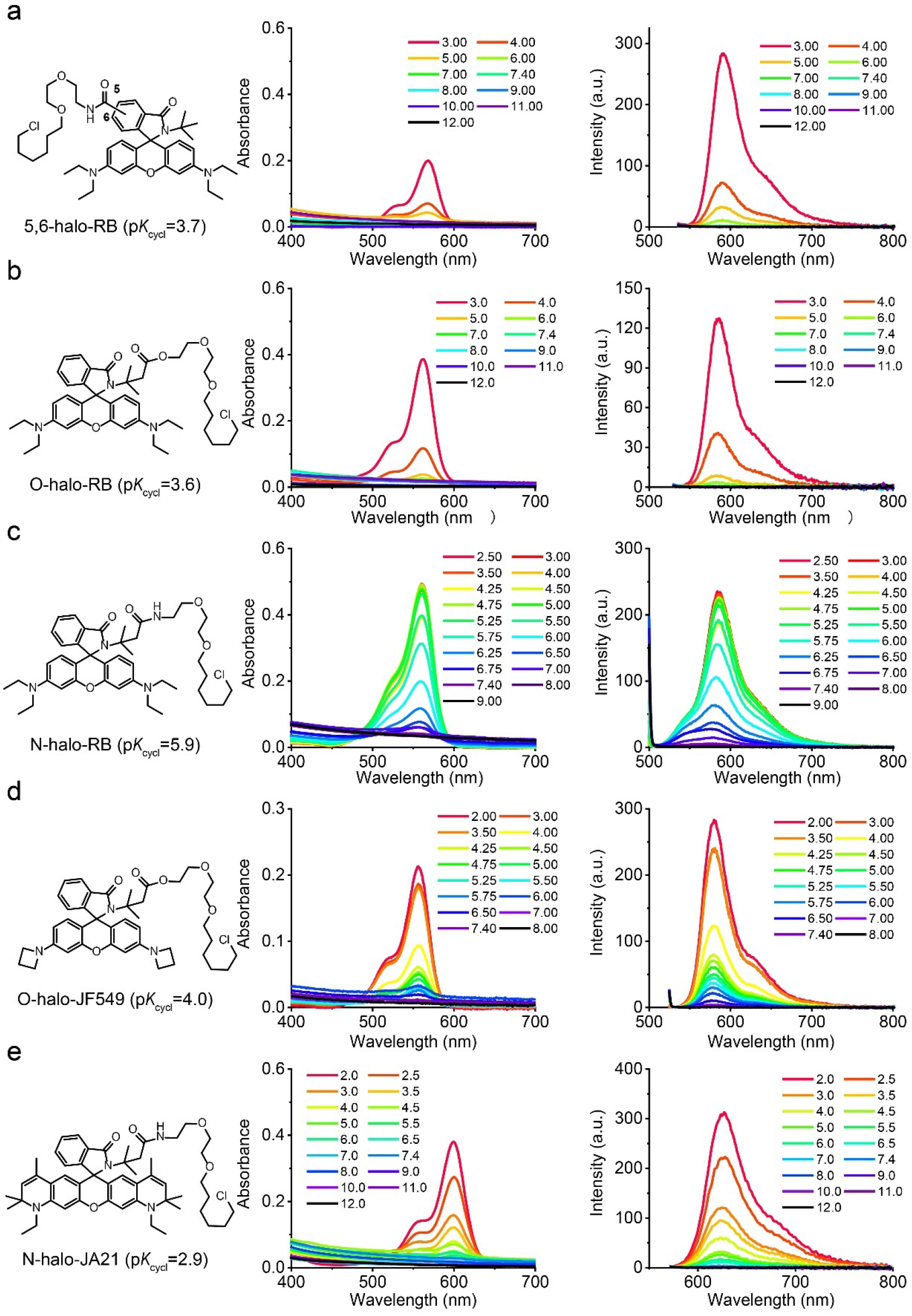
Absorbance (left) and fluorescence (right) spectra of **5,6-halo-RB** (a), **O-halo-RB** (b), **N-halo-RB** (c), **O-halo-JF549** (d) and **N-halo-JA21** (e). at different pH. Excitation wavelengths for fluorescence spectra were 520 nm, 490 nm, 510 nm and 550 nm for (a-b), (c), (d) and (e).

**Scheme S2.**
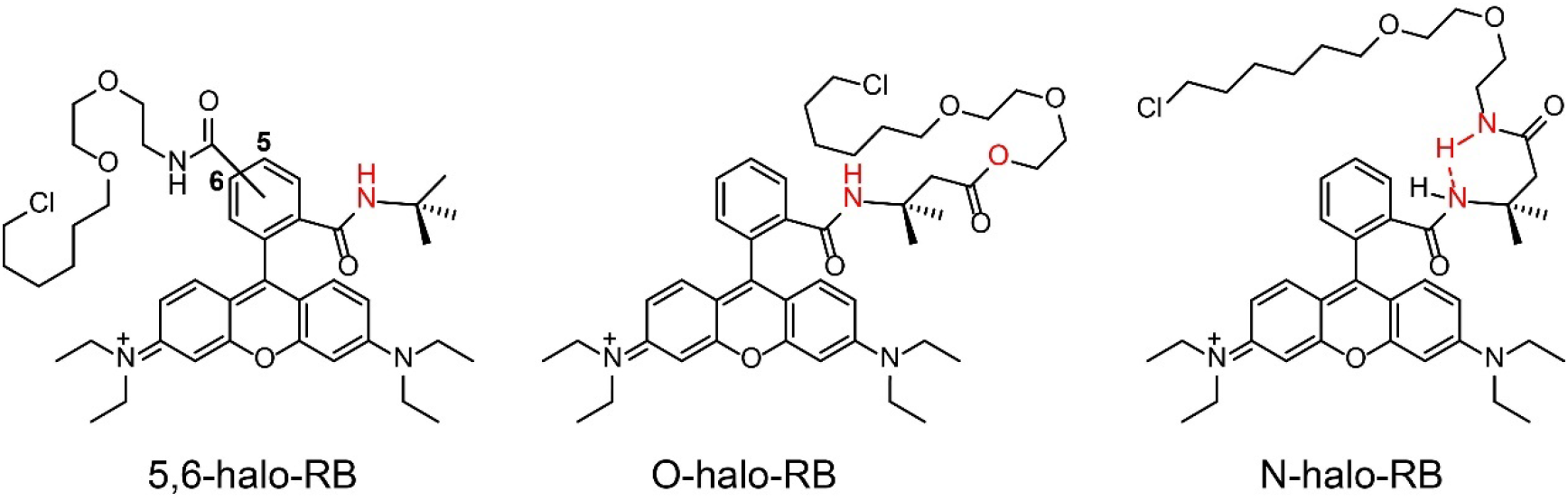
Possible intramolecular interactions with different bonding methods.

**Figure S5.**
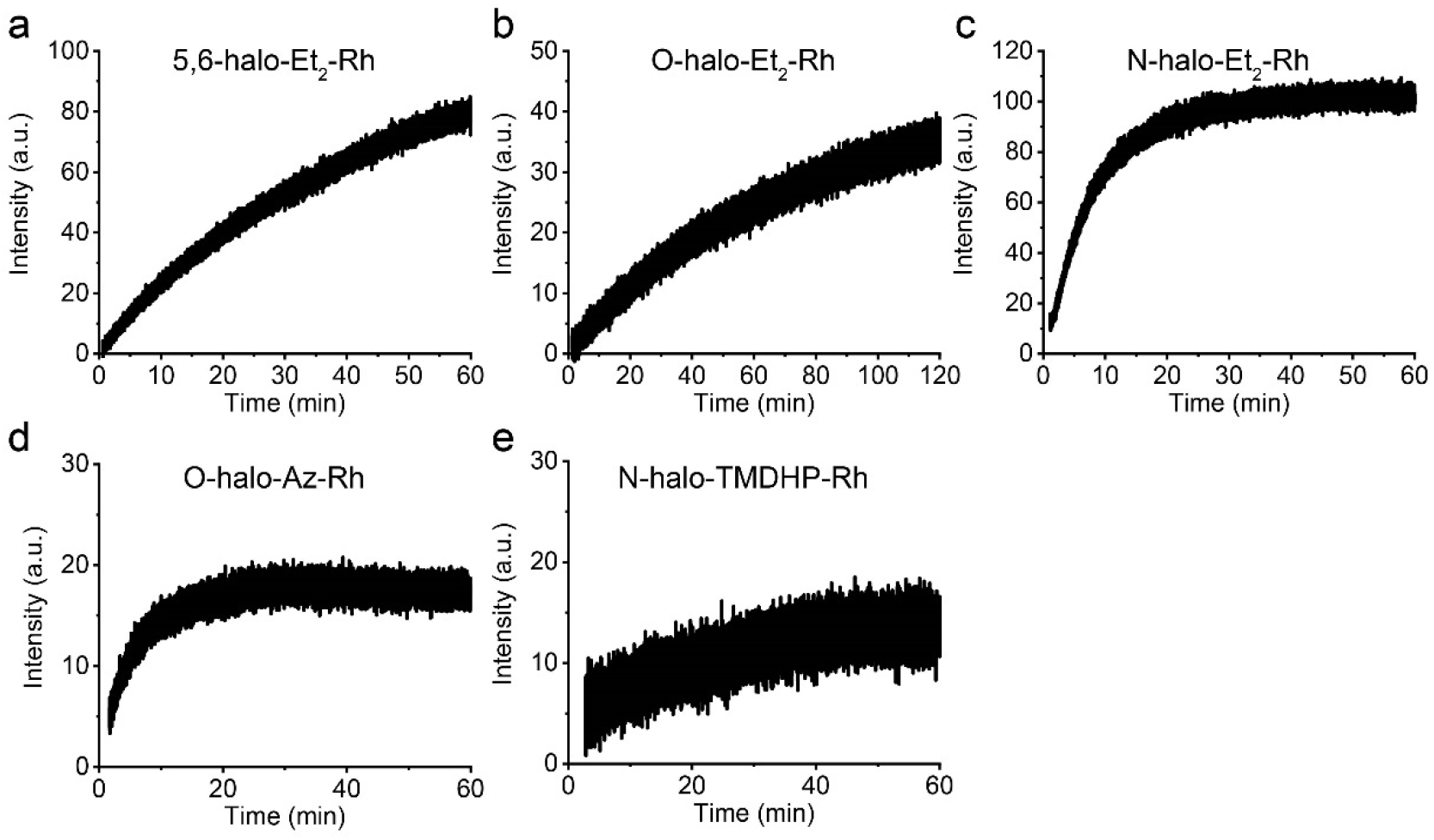
Kinetic study of **5,6-halo-RB** (a), **O-halo-RB** (b), **N-halo-RB** (c), **O-halo-JF549** (d) and **N-halo-JA21** (e) binding to halo protein (0.99 μM). The concentration of the probes were 0.5 μM. Excitation wavelengths for fluorescence spectra were 550 nm, 540 nm and 590 nm for (a-d), (e) and (f).

### 6.2 Binding kinetics of functionalized rhodamine derivatives with halo protein

### 6.3 Single-molecule studies of functionalized rhodamine derivatives with halo protein

**Figure S6.**
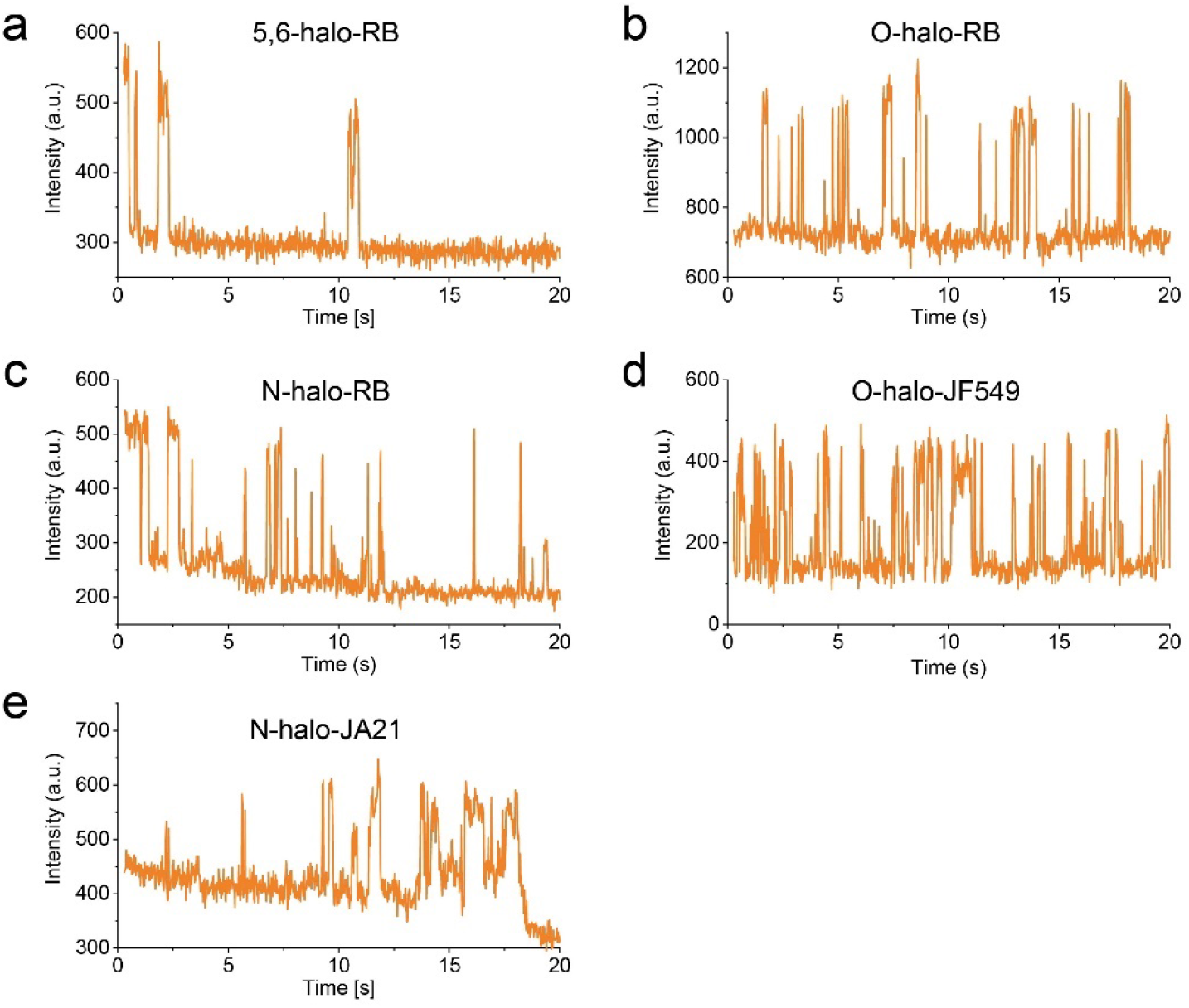
Single-molecule studies of **5,6-halo-RB** (a), **O-halo-RB** (b), **N-halo-RB** (c), **O-halo-JF549** (d) and **N-halo-JA21** (e). The analysis were measured in PMMA. Excitation at 561 nm.

## 7 Imaging details

**Figure S7.**
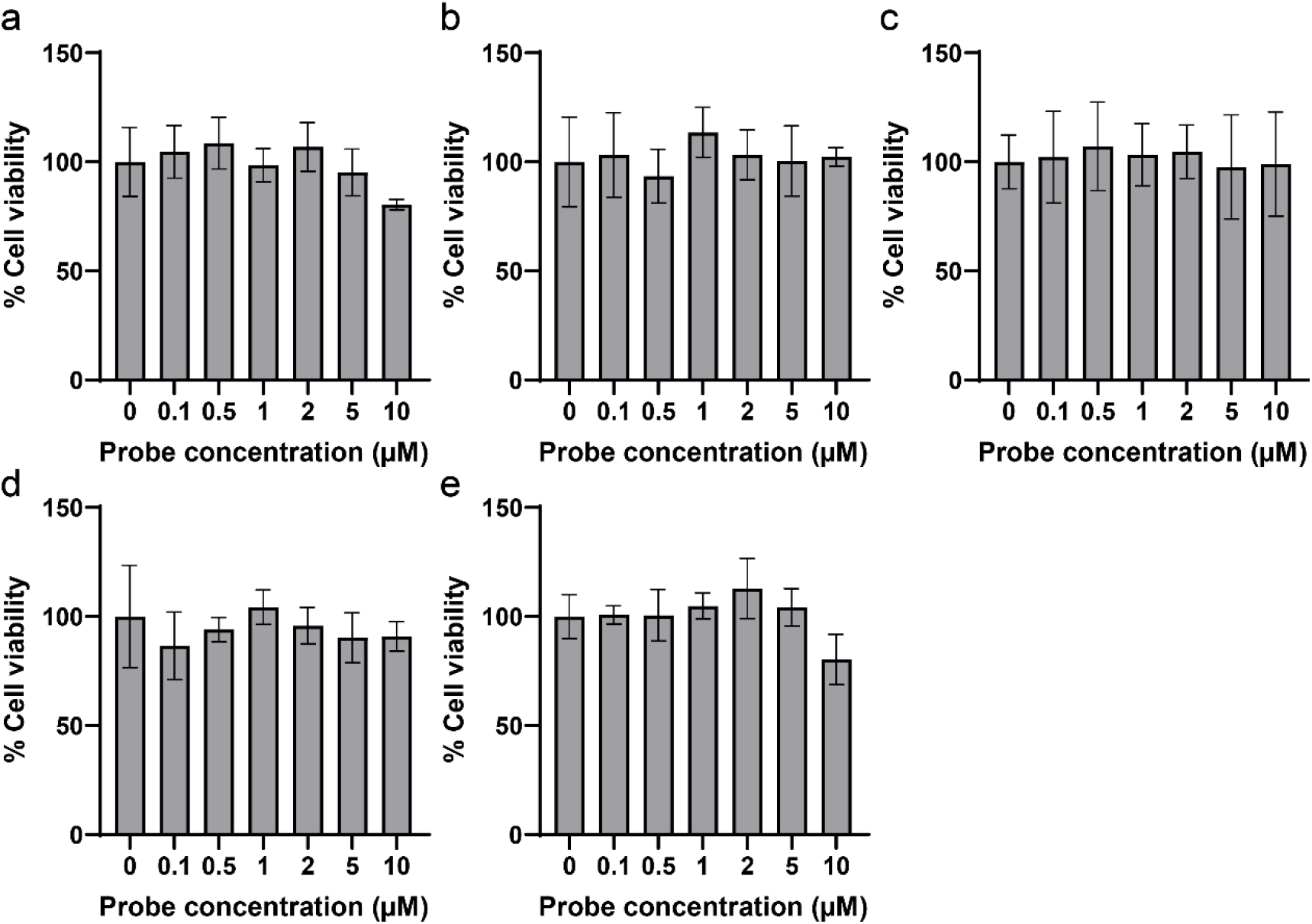
Cytotoxicity of HeLa cells of **5,6-halo-RB** (a), **O-halo-RB** (b), **N-halo-RB** (c), **O-halo-JF549** (d) and **N-halo-JA21** (e) after incubation for 24 h.

### 7.1 Fluorogenic properties of **5,6-halo-RB**, **O-halo-RB** and **N-halo-RB** characterized through confocal imaging

**Figure S8.**
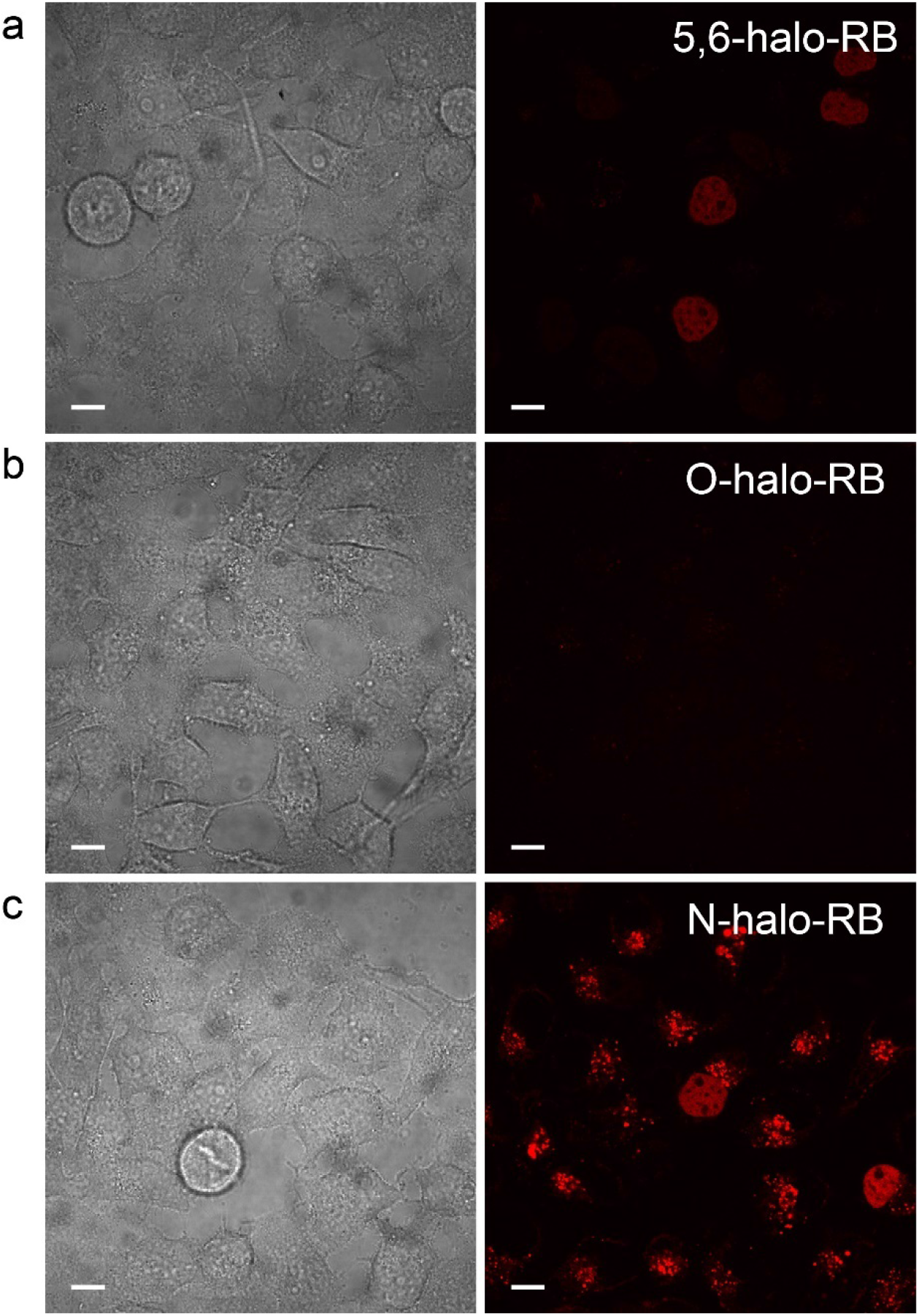
Wide field (left) and 543 channel (right) live cell no wash imaging of hela cells expressing H2B-halo fusion protein labeled with (a) **5,6-halo-RB** (400 nM), (b) **O-halo-RB** (400 nM) and (c) **N-halo-RB** (400 nM) for 1 h. Scale bar: 5 μm

**Figure S9.**
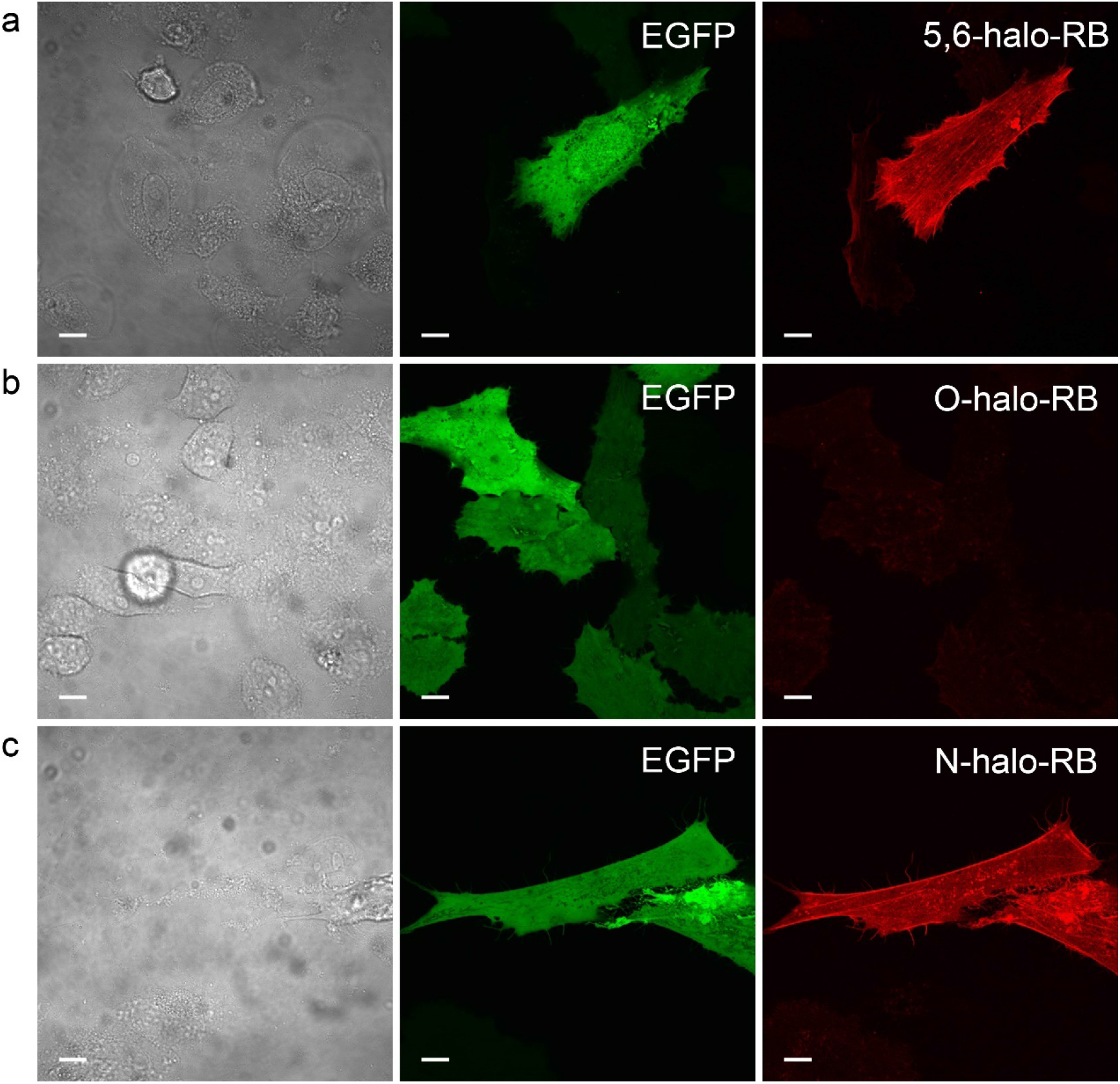
Wide field (left), 488 channel (middle) and 543 channel (right) live cell no wash imaging of hela cells expressing lifeact-halo-EGFP fusion protein labeled with (a) **5,6-halo-RB** (400 nM), (b) **O-halo-RB** (400 nM) and (c) **N-halo-RB** (400 nM) for 1 h. Scale bar: 5 μm

### 7.2 Fluorogenic properties of **5,6-halo-RB** characterized through SIM imaging

**Figure S10.**
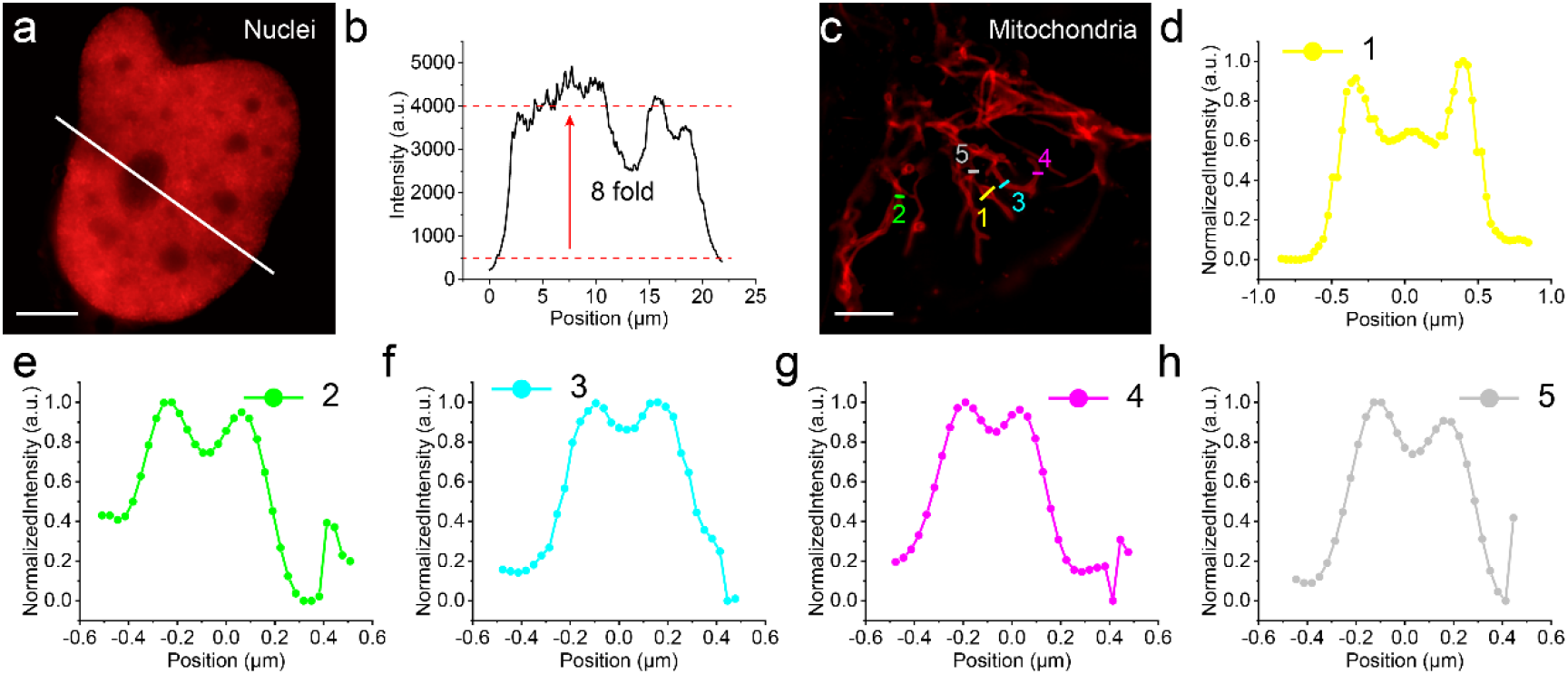
Live cell no wash SIM imaging labeled with **5,6-halo-RB** for 1 h. (a) Hela cells expressing halo-H2B (400 nM). (b) MCF cells expressing halo-EGFP-actin (400 nM). (c) Hela cells expressing halo-EGFP-tom20 (100 nM). (d) Hela cells expressing halo-tubulin (200 nM). Scale bar: 5 μm

### 7.3 SMLM properties of **O-halo-JF549** in STORM imaging

**Figure S11.**
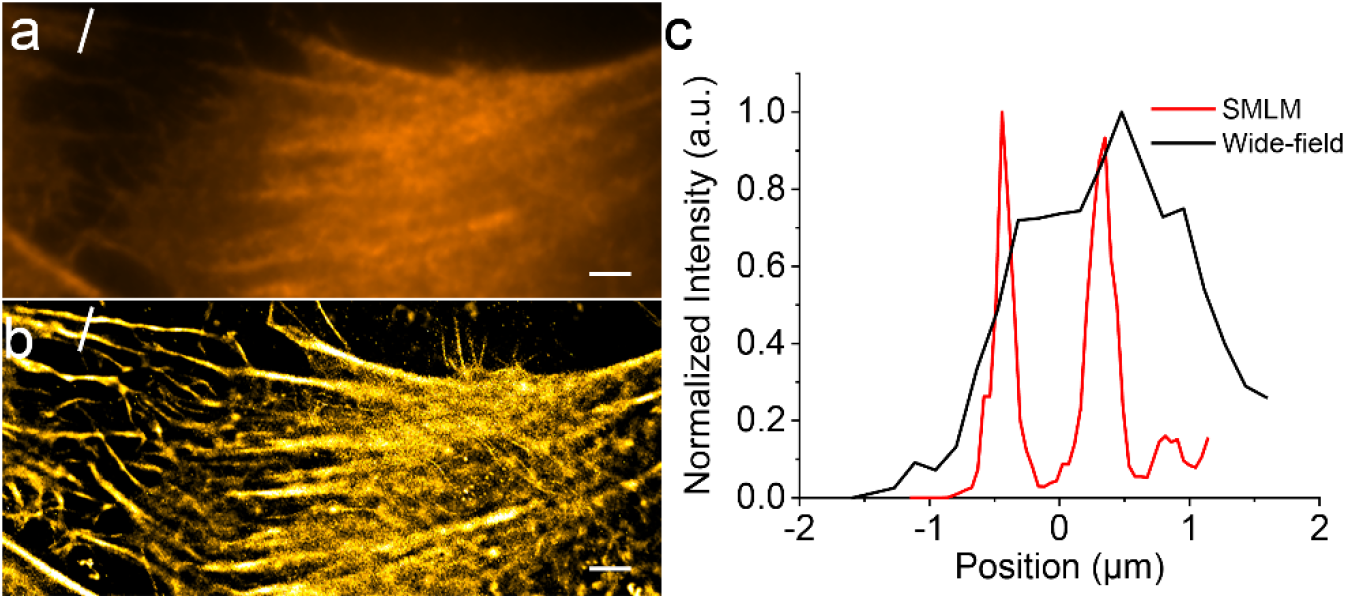
Live cell no was SMLM imaging labeled with **O-halo-JF549** for 1 h. (a) Wide-field imaging. (b) Reconstructed imagine of (a). (c) Normalized intensity across the white line in (a) and (b). Scale bar: 2 μm.

### 7.4 Cell kinetics in STORM imaging

**Figure S12.**
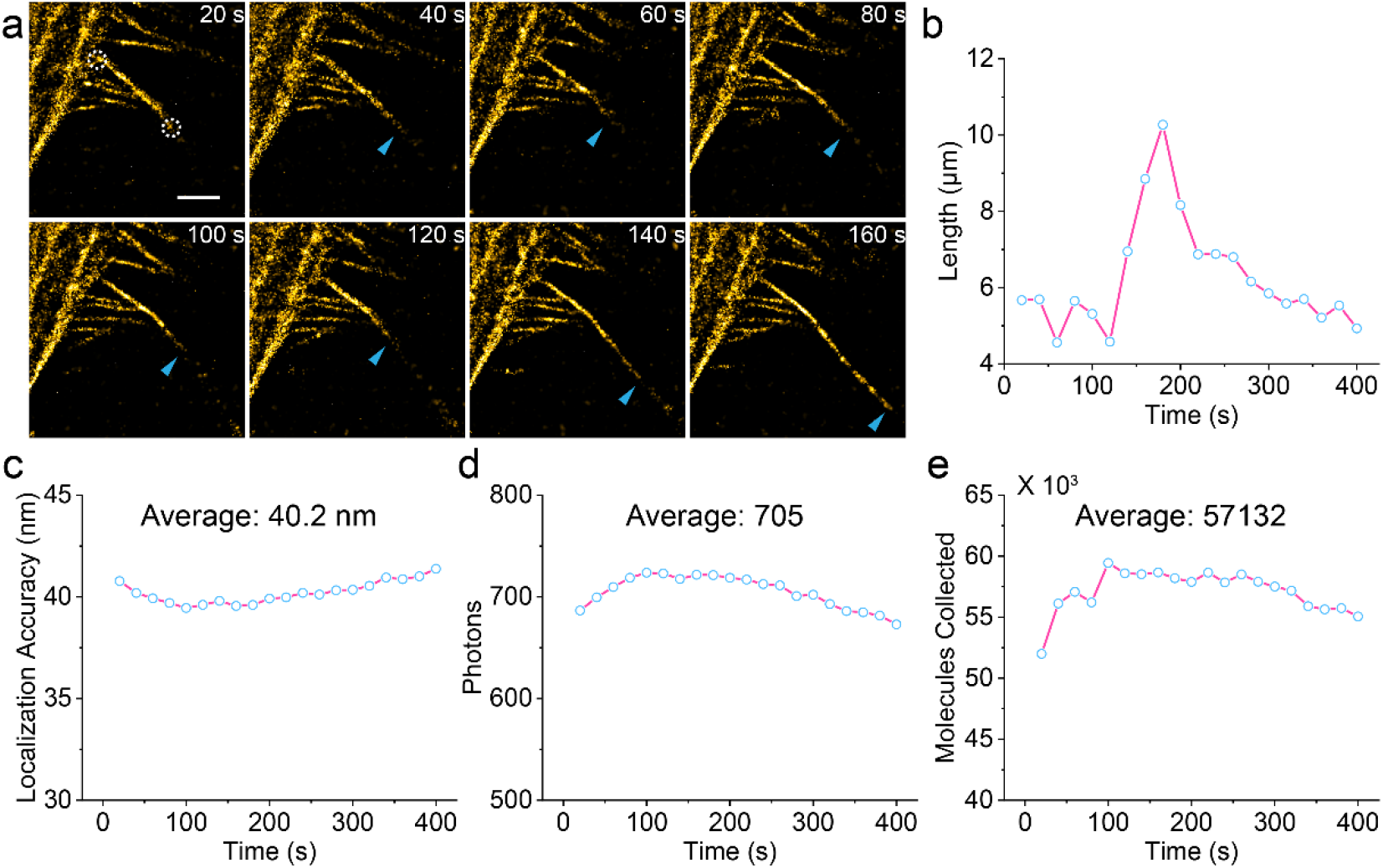
(a) Long-term SMLM imaging of the region of interest of pseudopodia. Scale bar: 2 μm. (b) The length of the selected pseudopodium in (a) as the function of time. (c) Localization accuracy every 1000 frames. (d) Photons collected every 1000 frames. (e) Molecules collected every 1000 frames. The cut off value was 300 photons.

## 8 Spectra characterization

**Figure S13.**
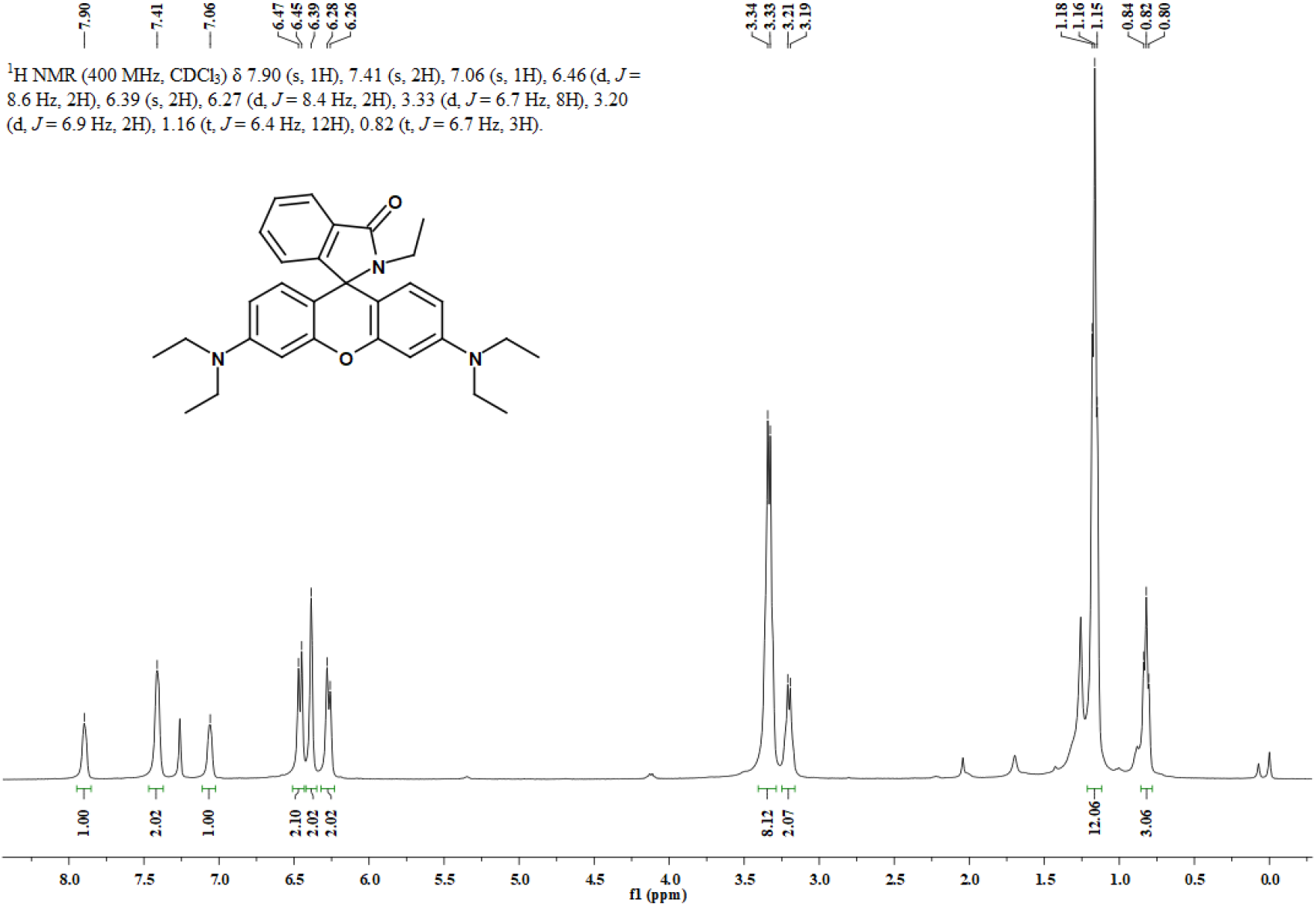
^1^H NMR spectrum of **1a** in CDCl_3_.

**Figure S14.**
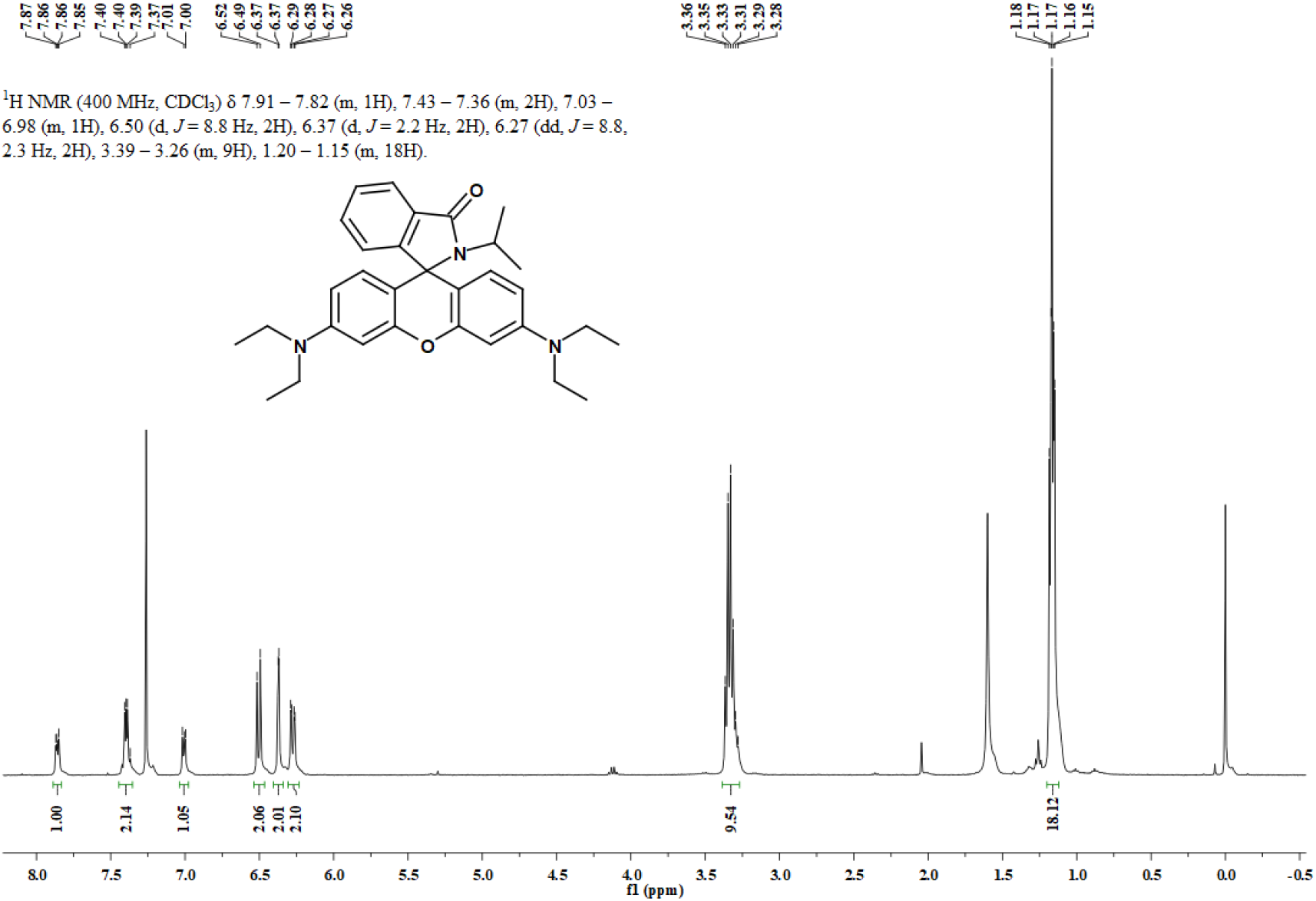
^1^H NMR spectrum of **1b** in CDCl_3_.

**Figure S15.**
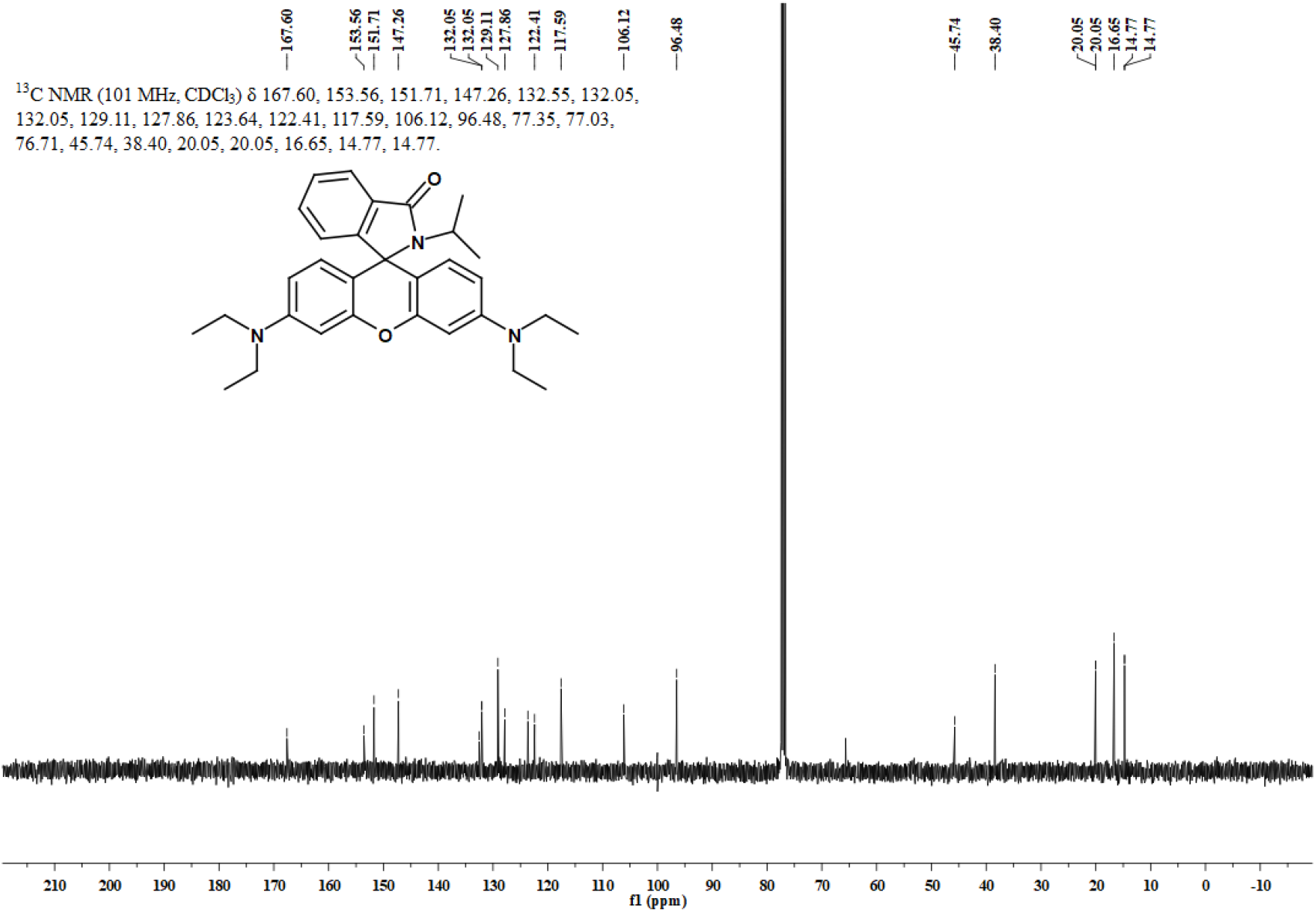
^13^C NMR spectrum of **1b** in CDCl_3_.

**Figure S16.**
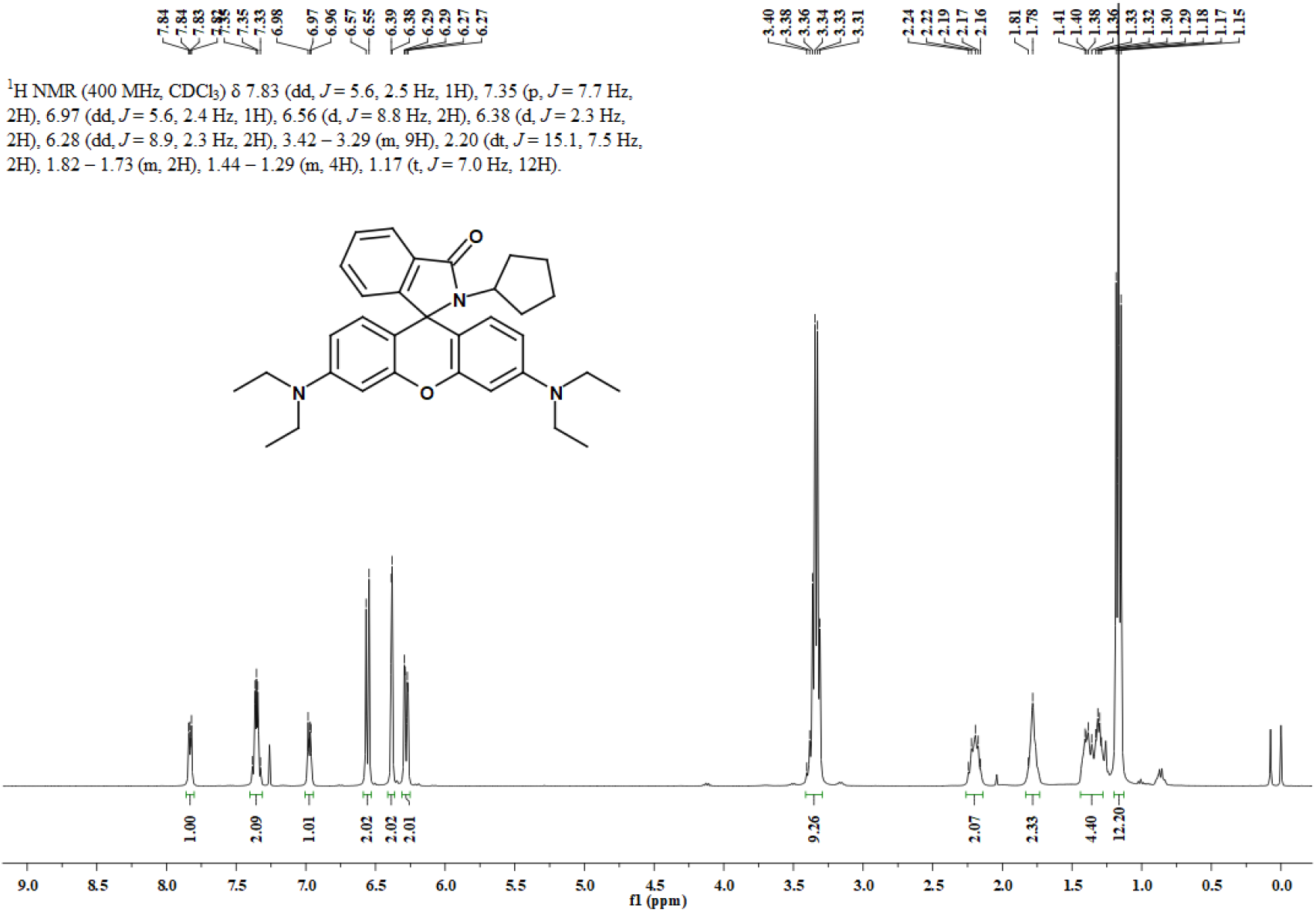
^1^H NMR spectrum of **1c** in CDCl_3_.

**Figure S17.**
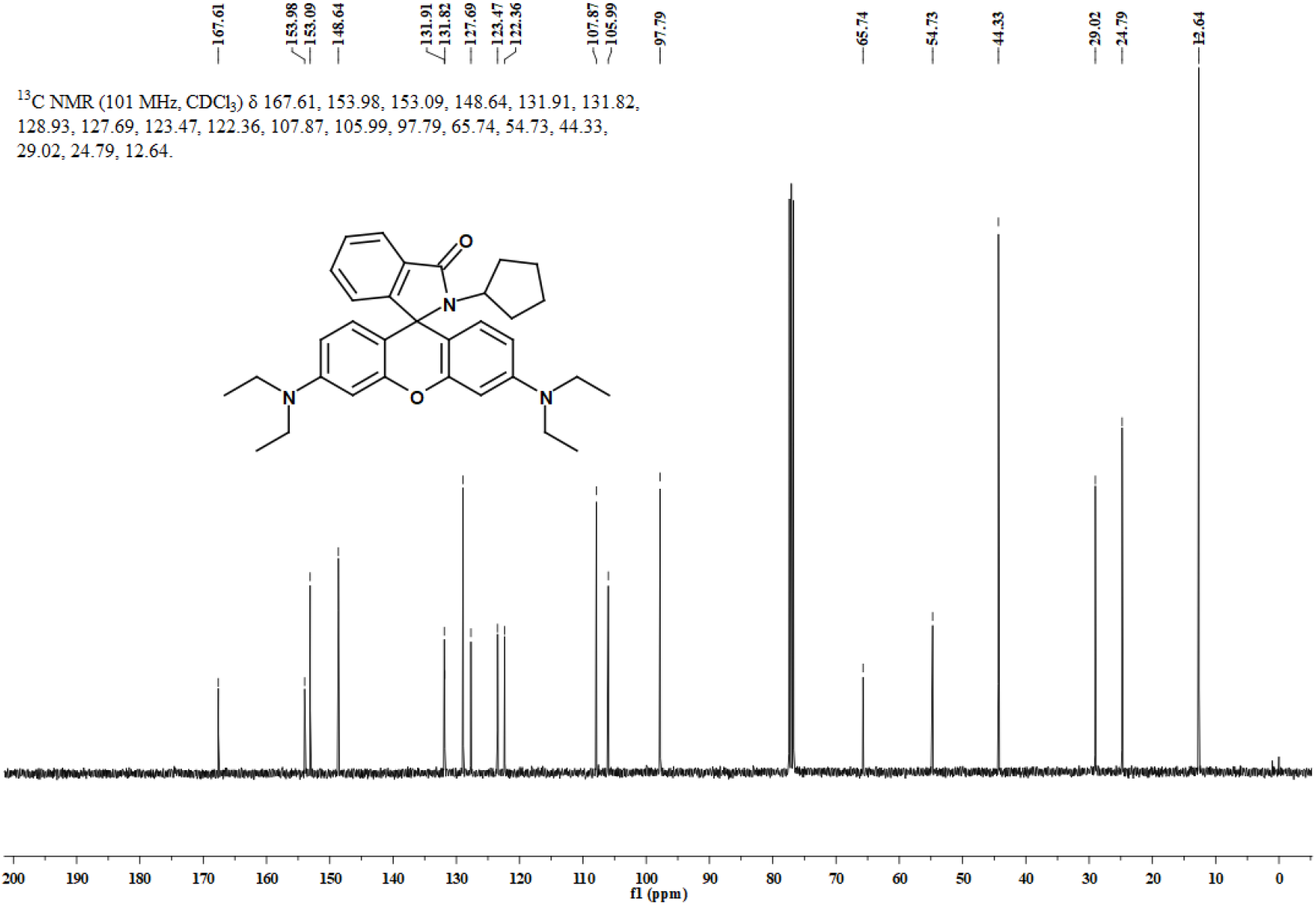
^13^C NMR spectrum of **1c** in CDCl_3_.

**Figure S18.**
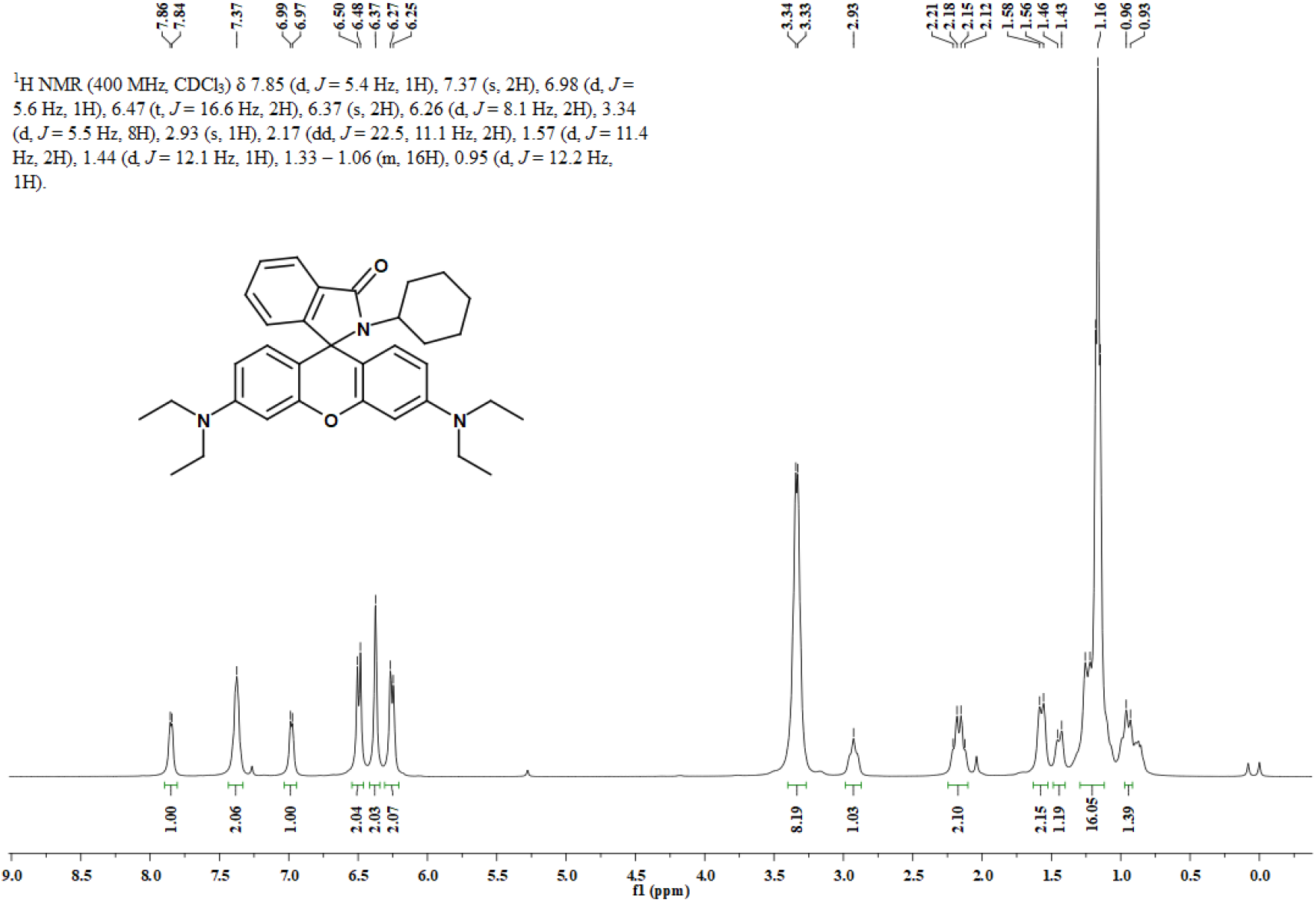
^1^H NMR spectrum of **1d** in CDCl_3_.

**Figure S19.**
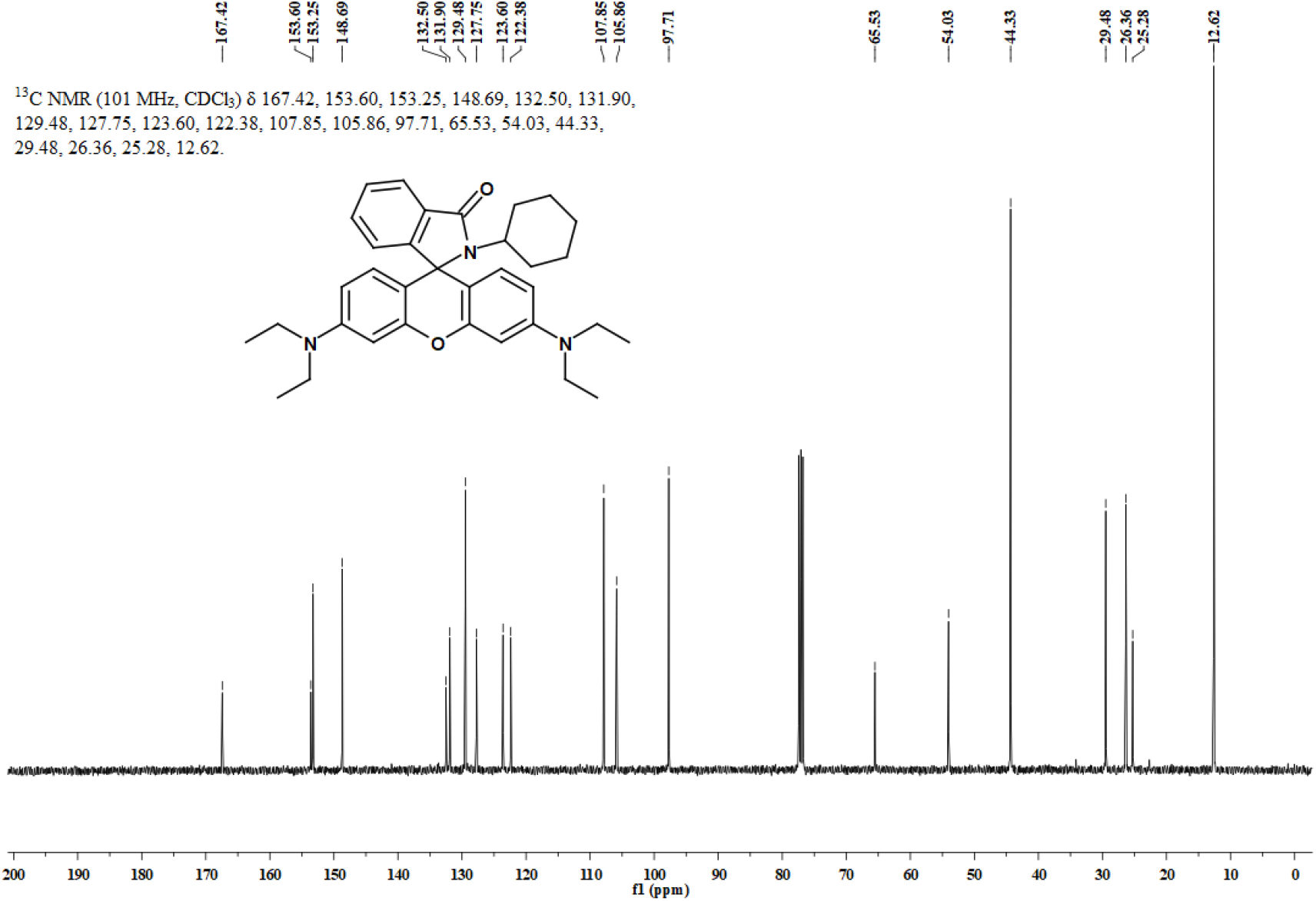
^13^C NMR spectrum of **1d** in CDCl_3_.

**Figure S20.**
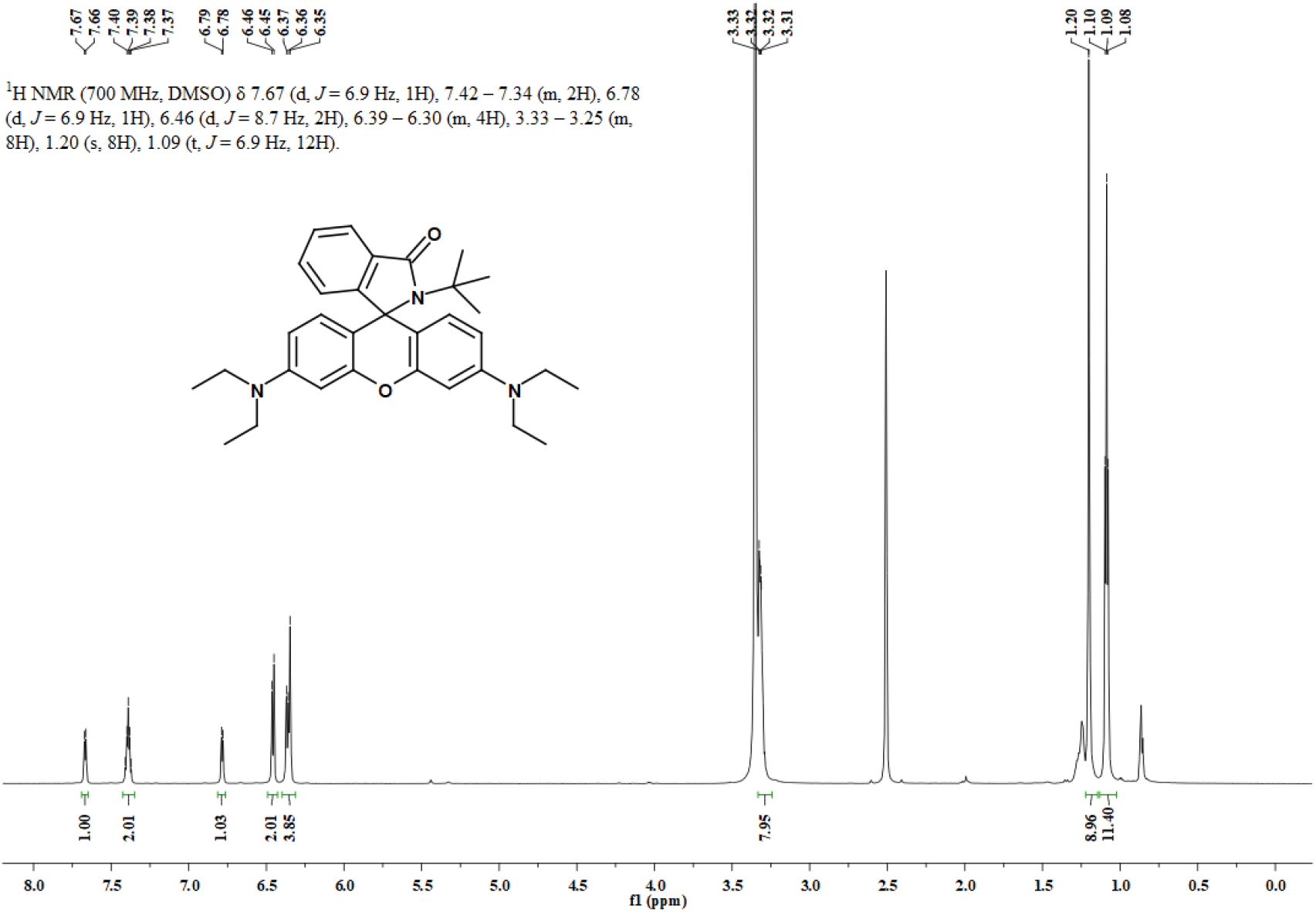
^1^H NMR spectrum of **1e** in CDCl_3_.

**Figure S21.**
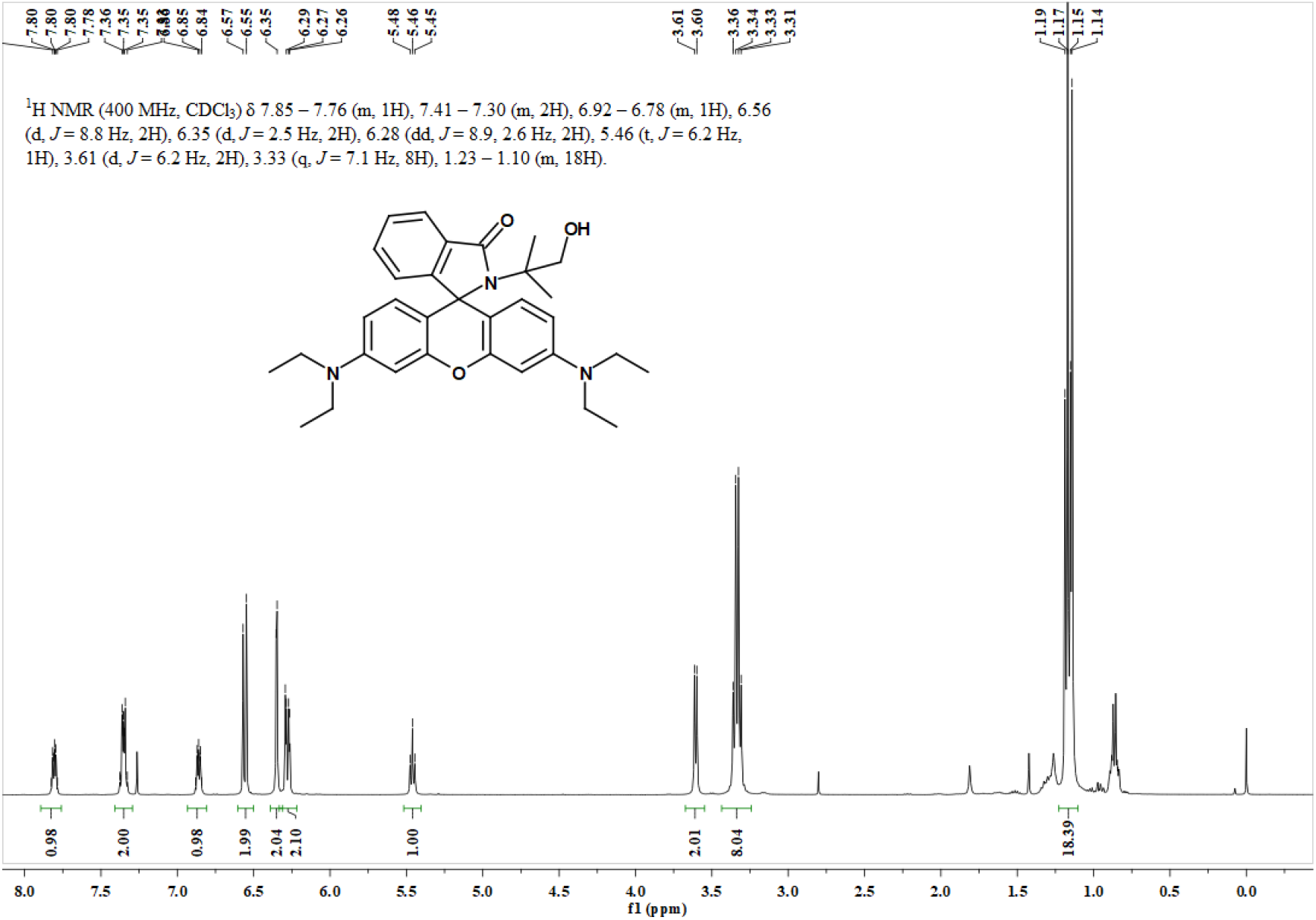
^1^H NMR spectrum of **1f** in CDCl_3_.

**Figure S22.**
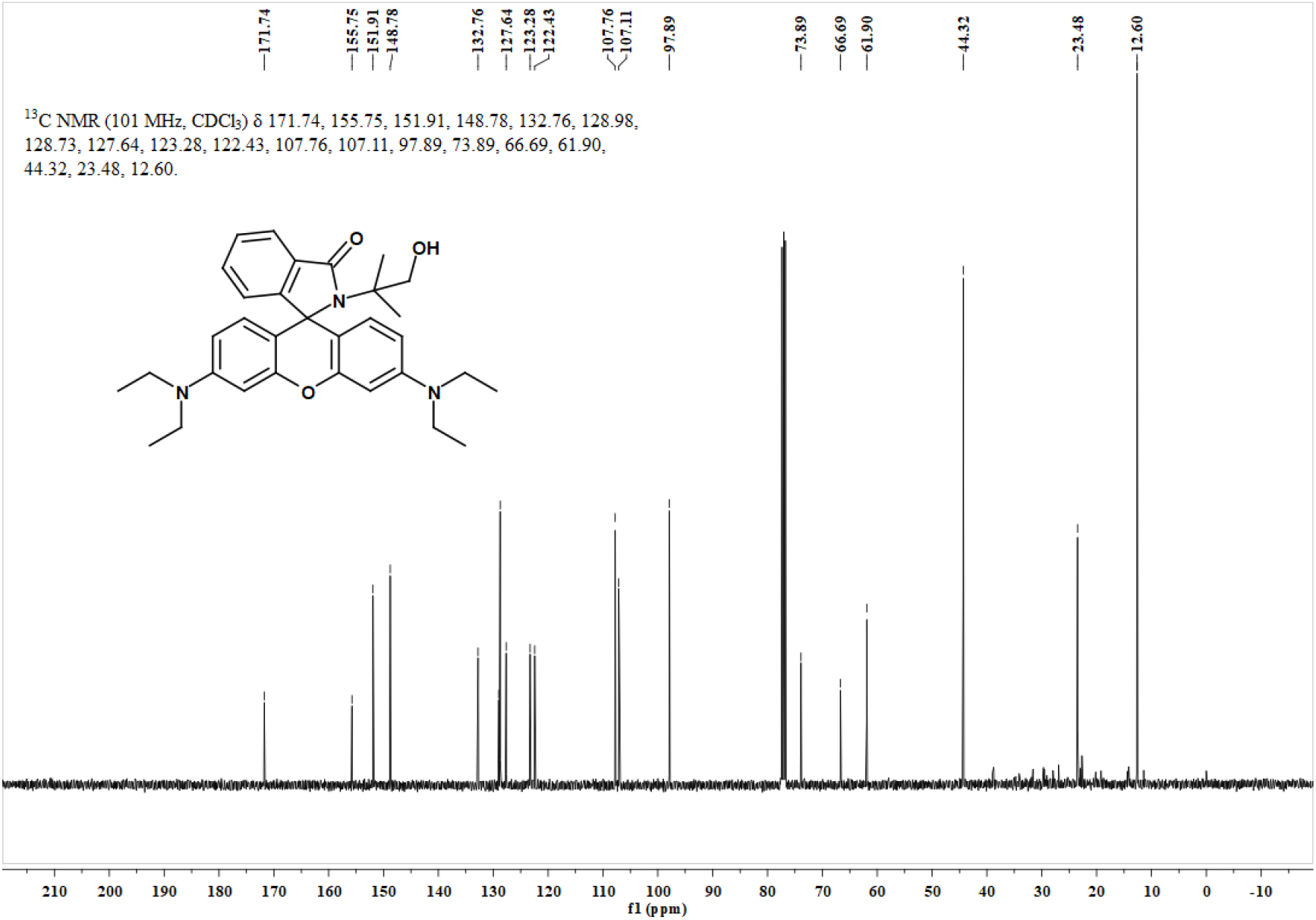
^13^C NMR spectrum of **1f** in CDCl_3_.

**Figure S23.**
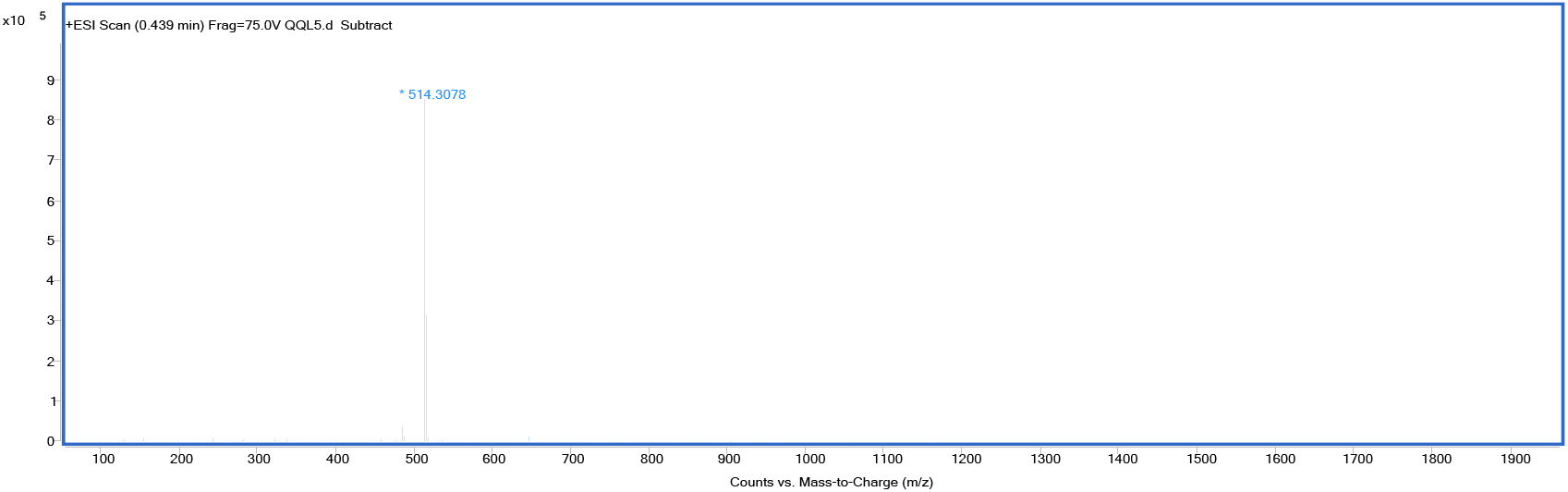
HRMS spectrum of **1f**.

**Figure S24.**
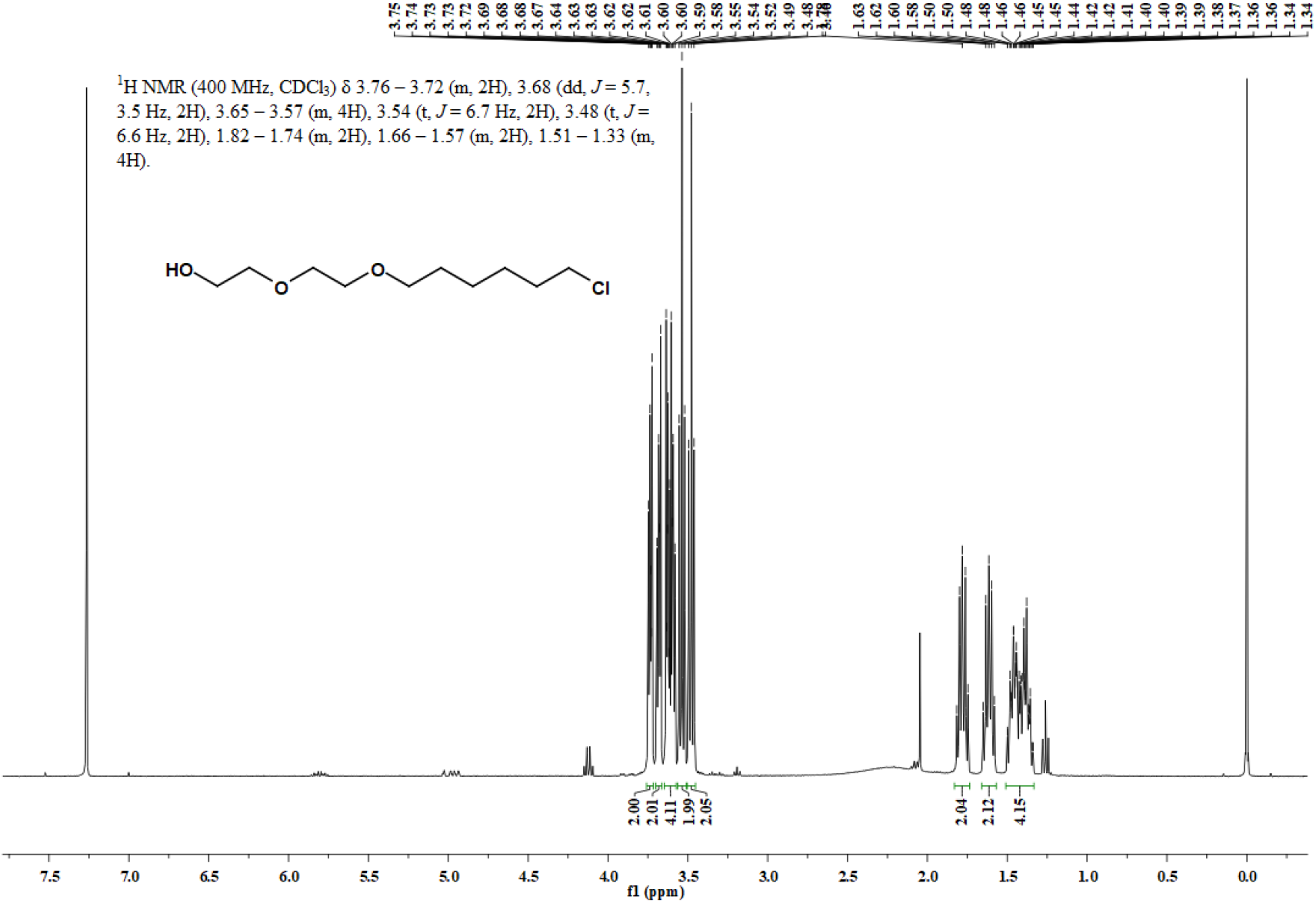
^1^H NMR spectrum of 2-(2-((6-chlorohexyl)oxy)ethoxy)ethan-1-ol in CDCl_3_.

**Figure S25.**
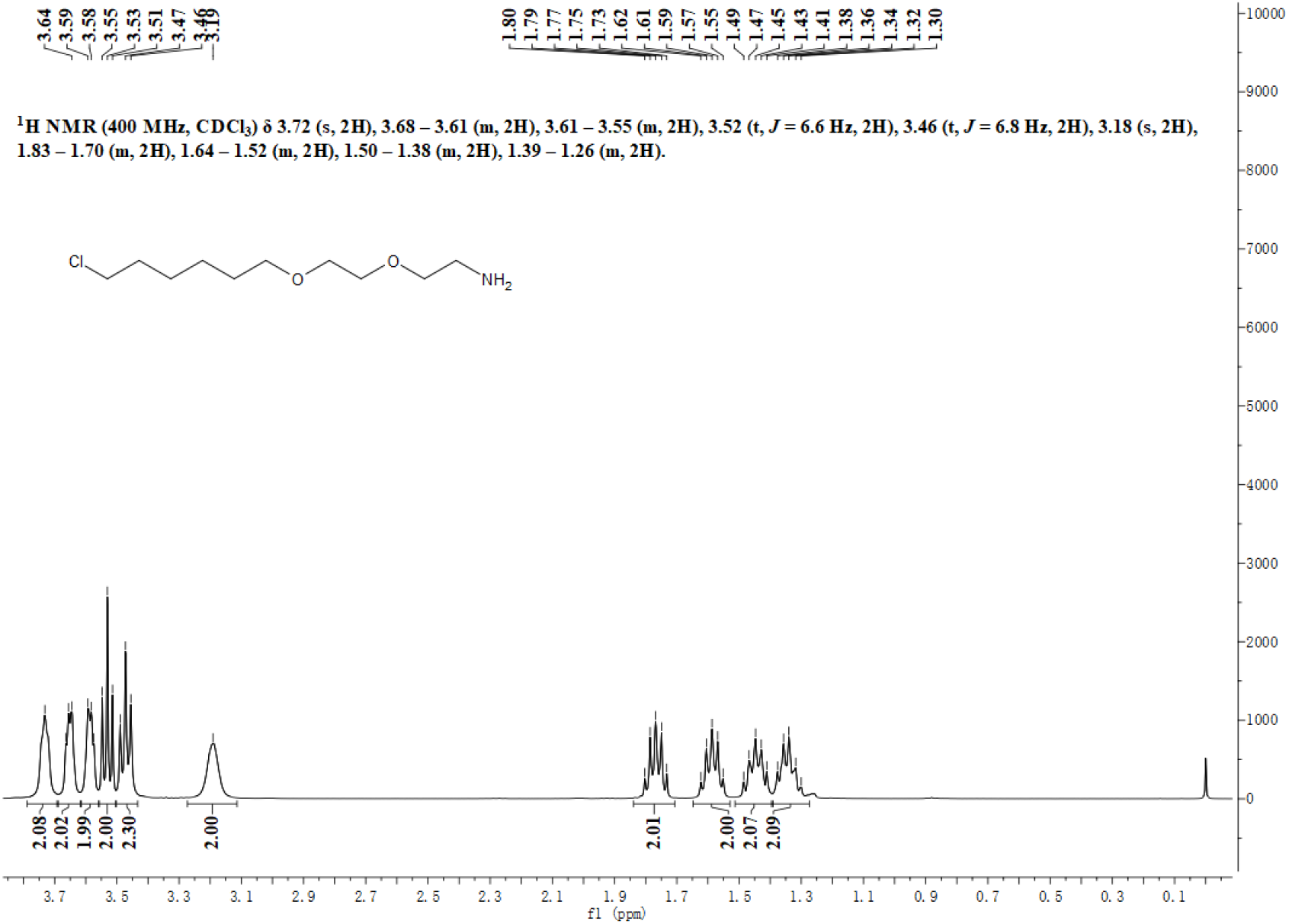
^1^H NMR spectrum of 2-(2-((6-chlorohexyl)oxy)ethoxy)ethan-1-amine in CDCl_3_.

**Figure S26.**
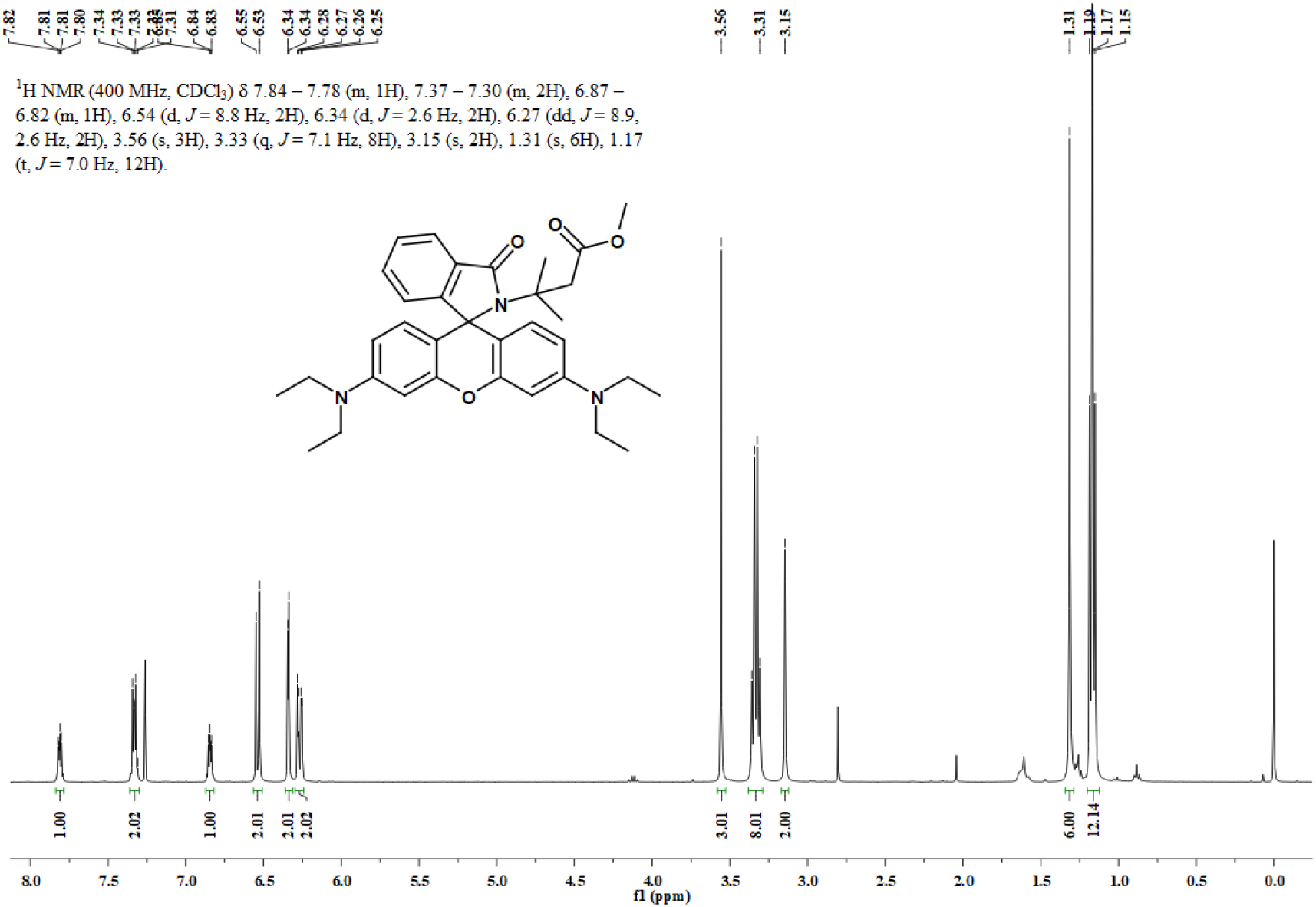
^1^H NMR spectrum of compound **6** in CDCl_3_.

**Figure S27.**
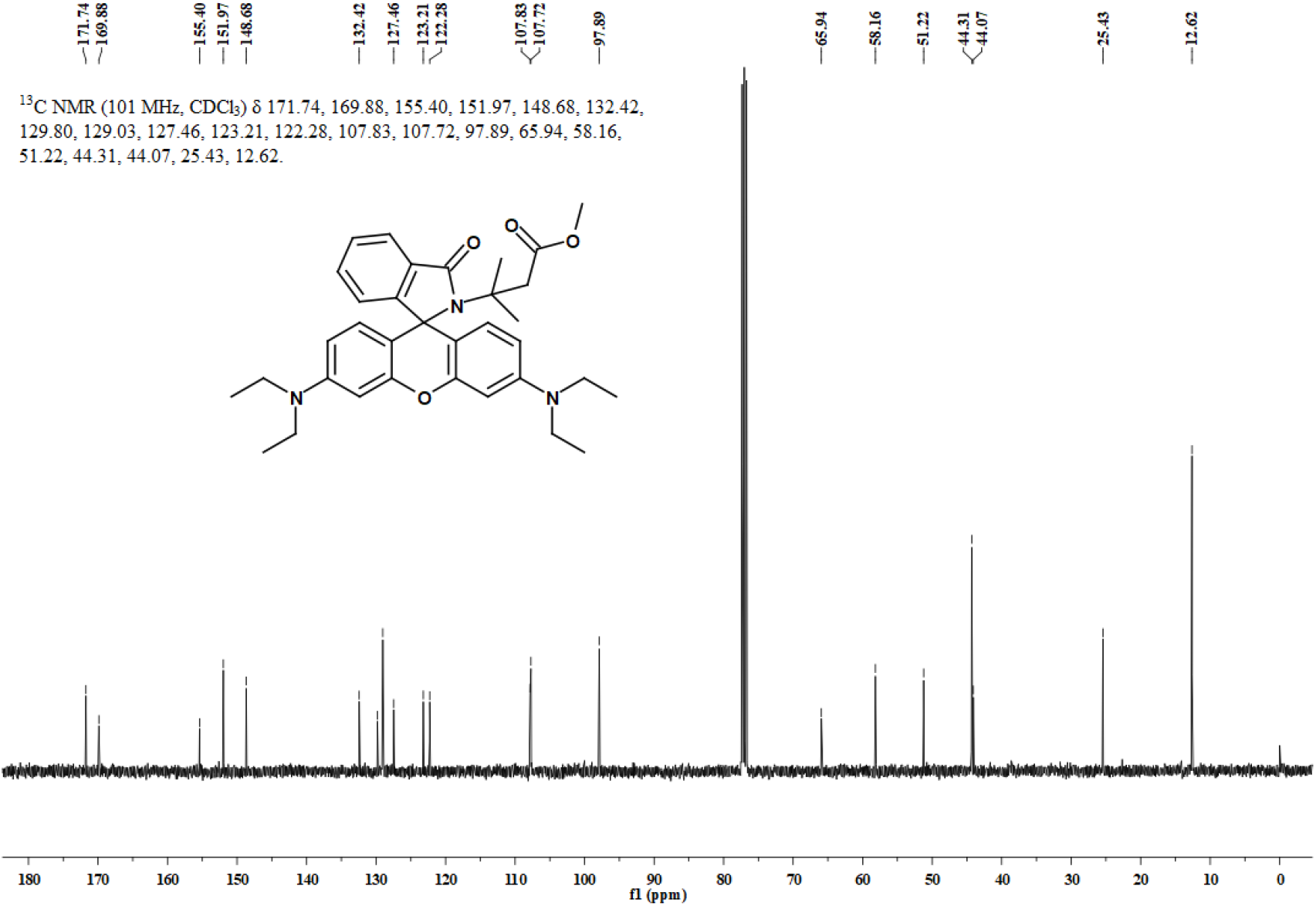
^13^C NMR spectrum of compound **6** in CDCl_3_.

**Figure S28.**
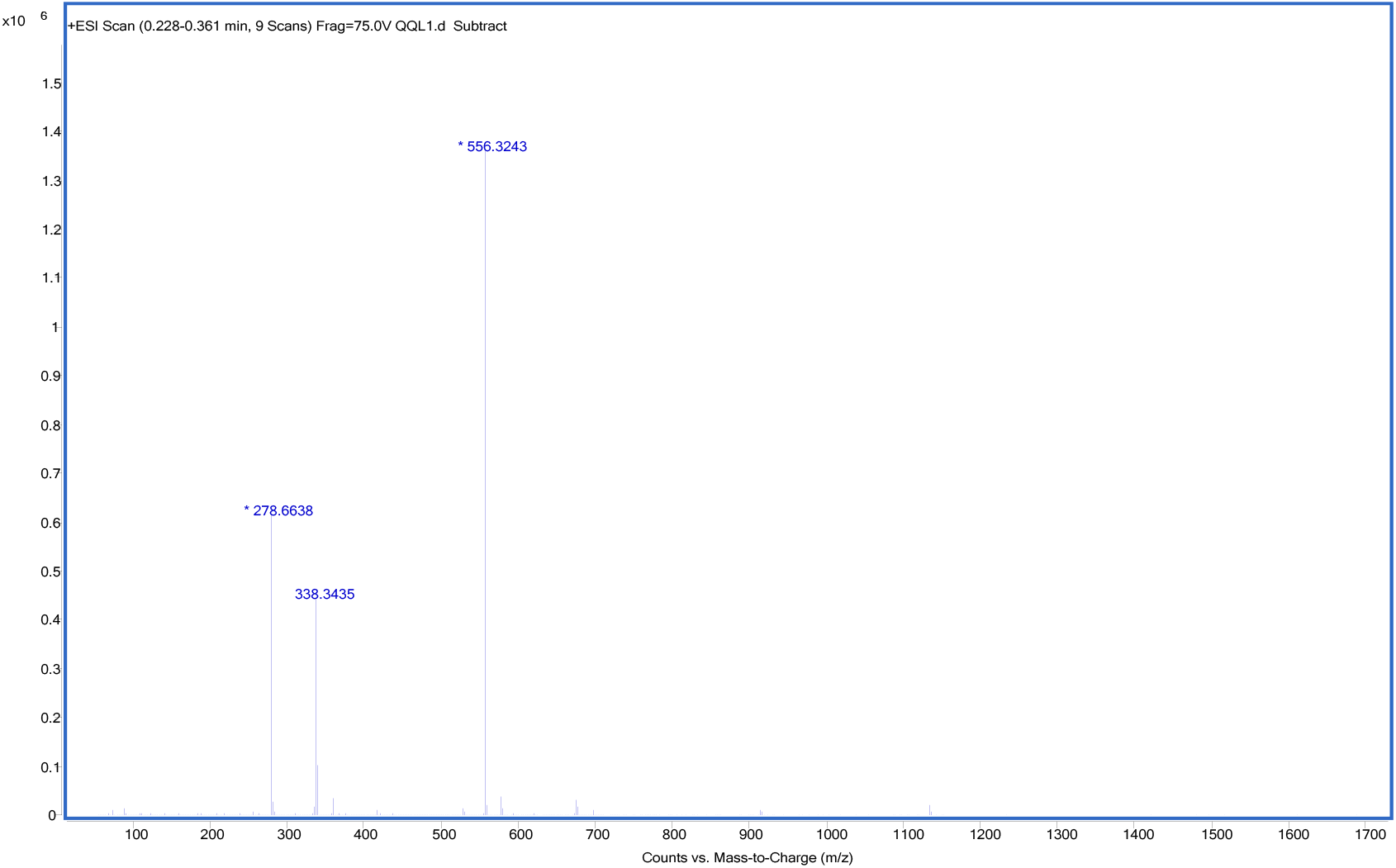
HRMS spectrum of **1f**.

**Figure S29.**
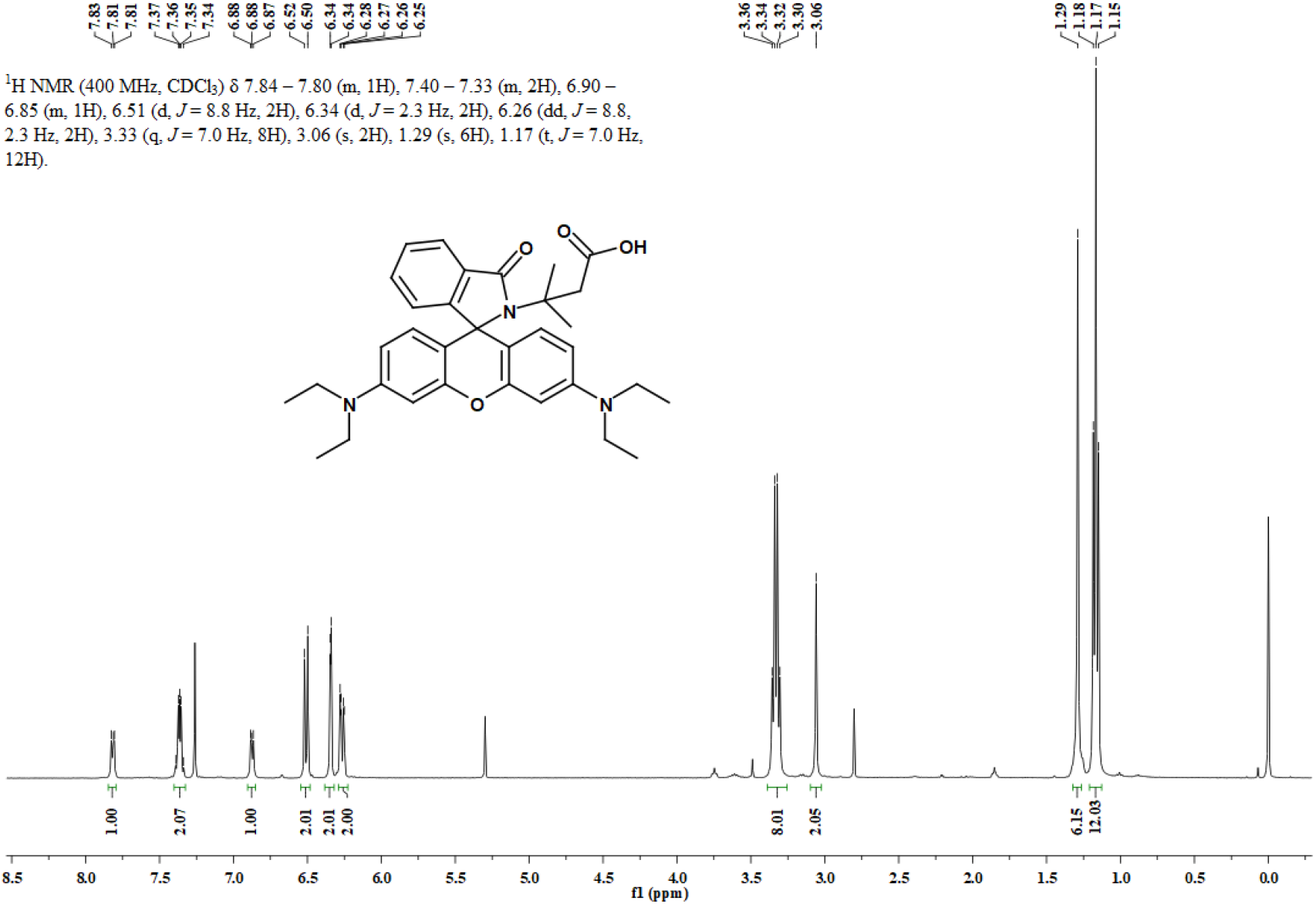
^1^H NMR spectrum of compound **7** in CDCl_3_.

**Figure S30.**
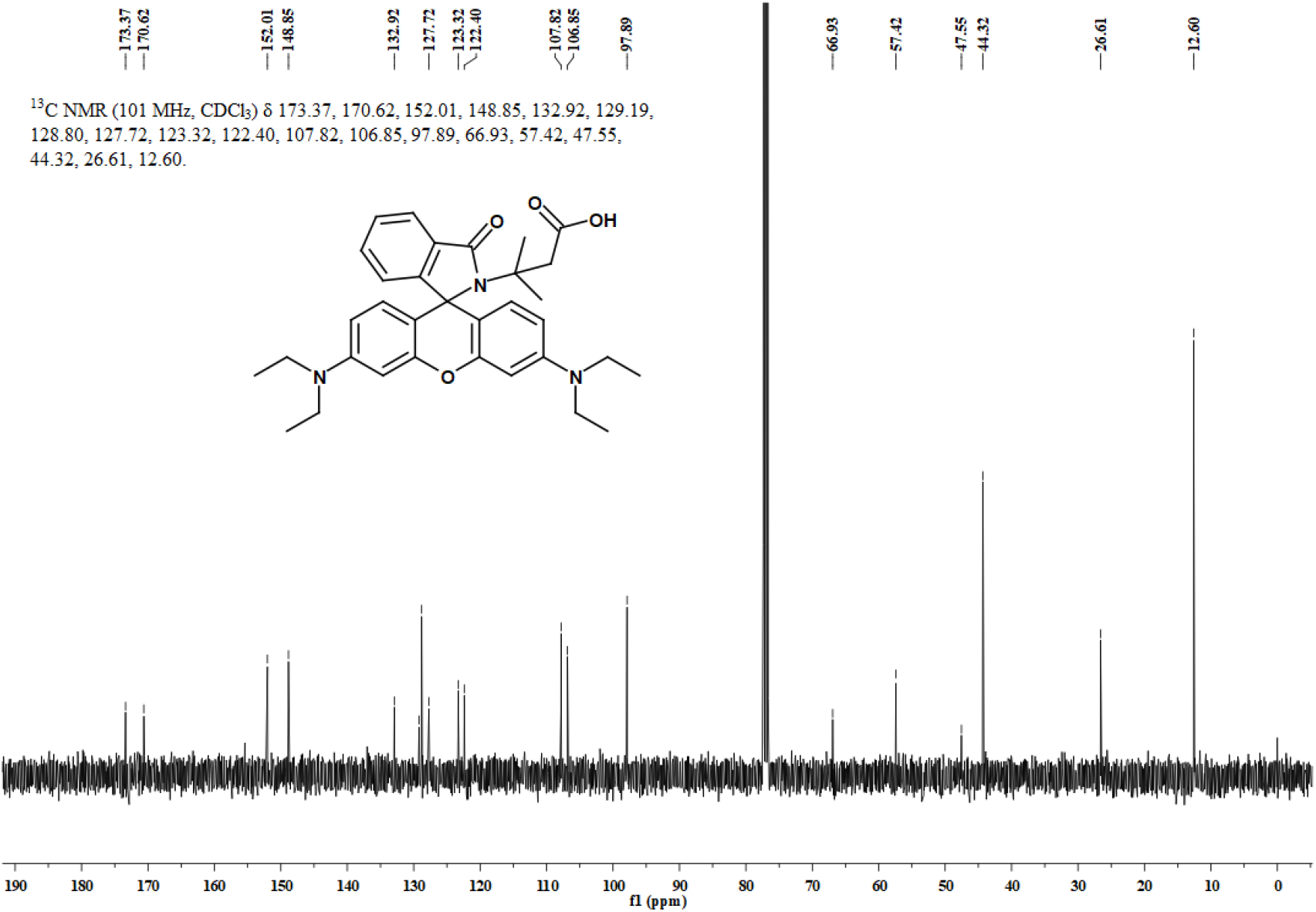
^13^C NMR spectrum of compound **7** in CDCl_3_.

**Figure S31.**
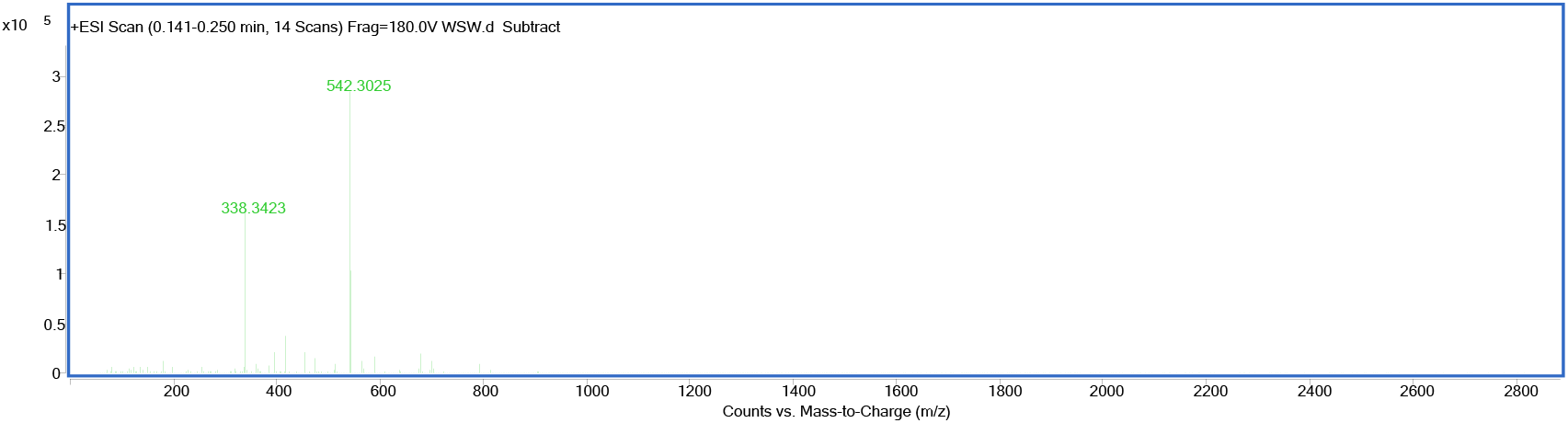
HRMS spectrum of compound **7**.

**Figure S32.**
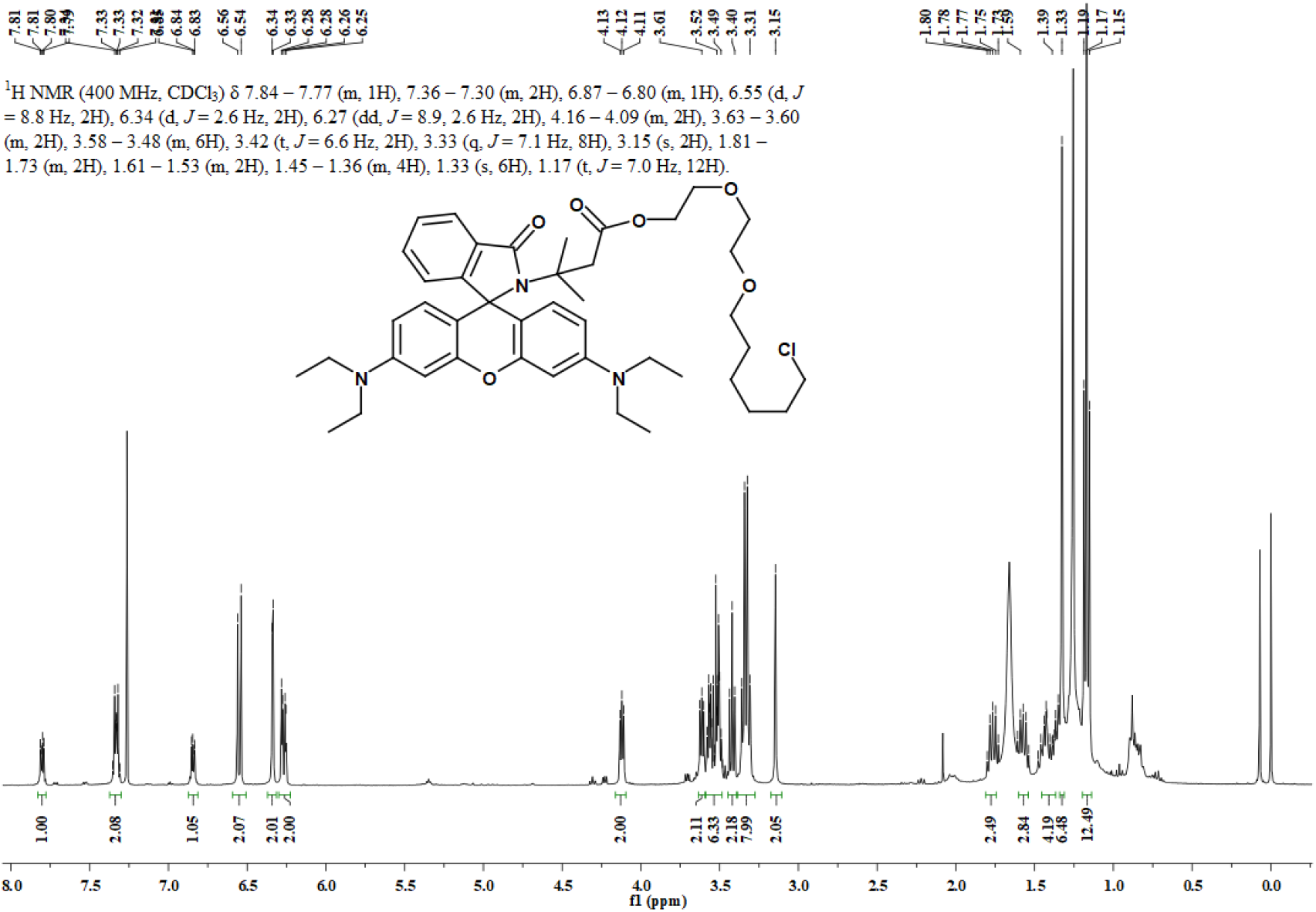
^1^H NMR spectrum of **1h** in CDCl_3_.

**Figure S33.**
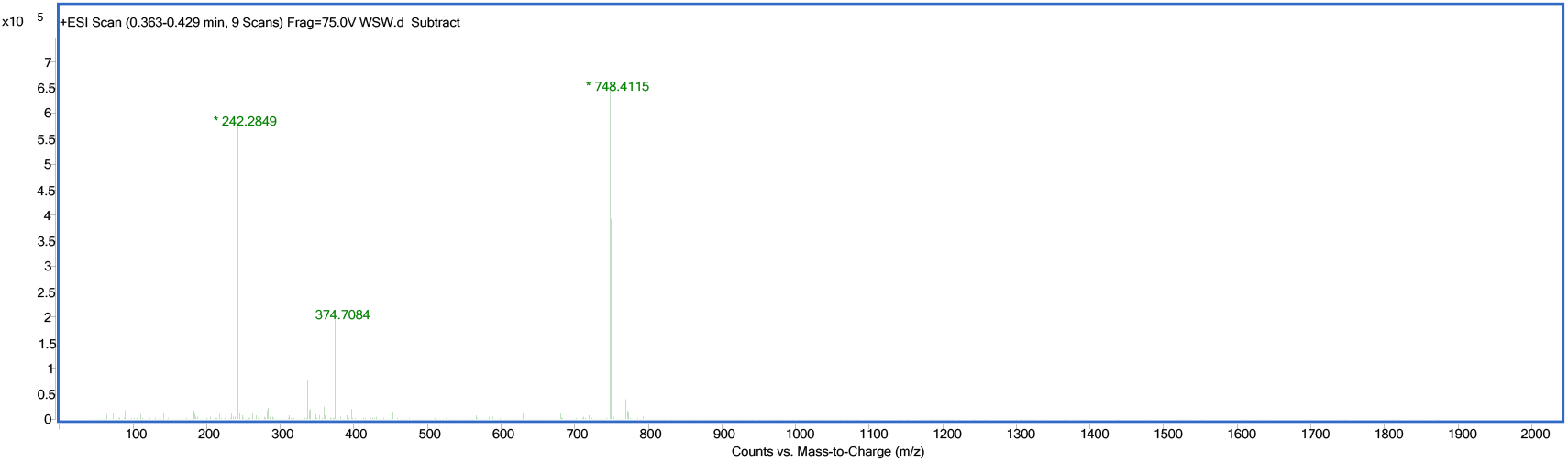
HRMS spectrum of **1h**.

**Figure S34.**
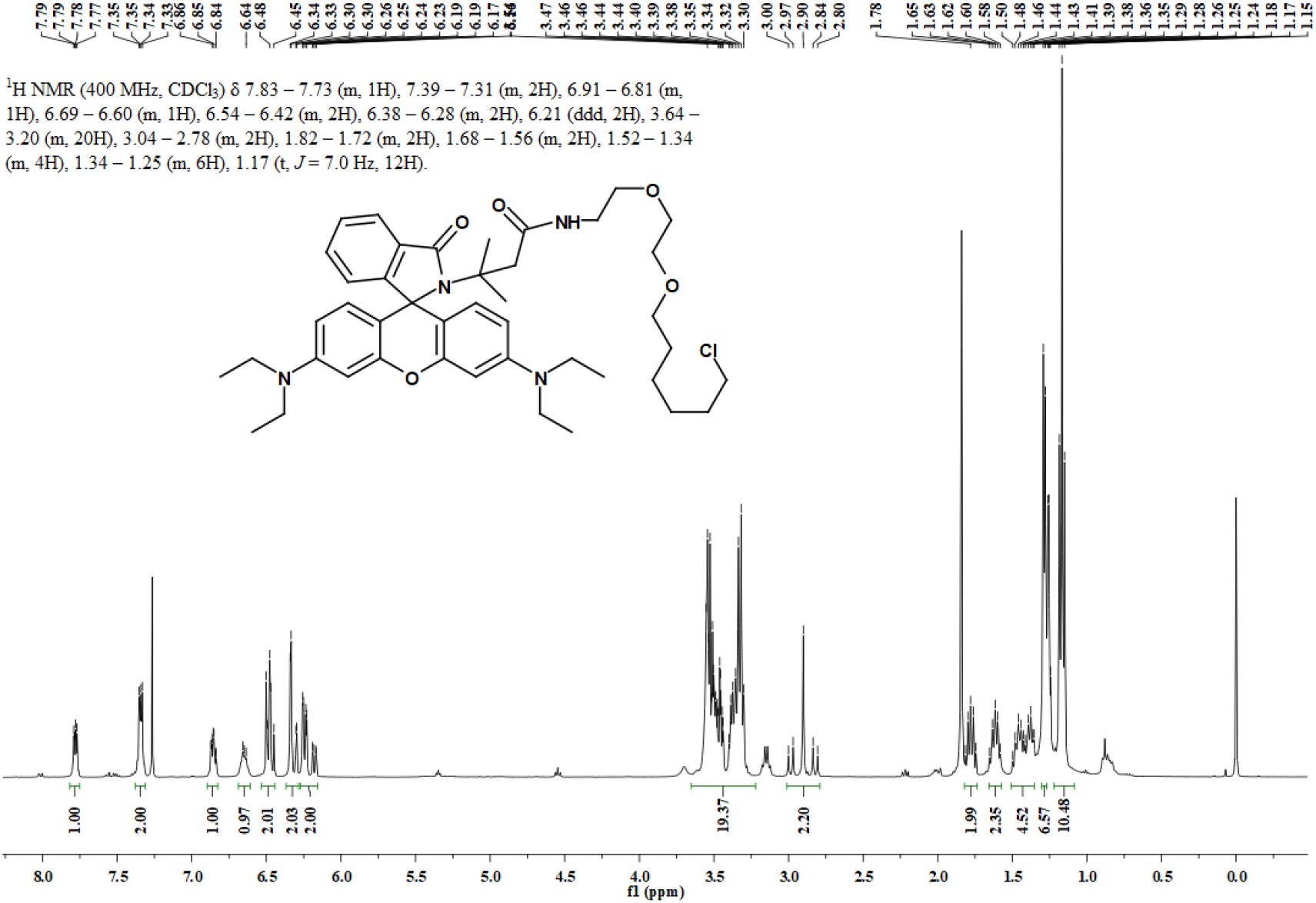
^1^H NMR spectrum of **N-halo-RB** in CDCl_3_.

**Figure S35.**
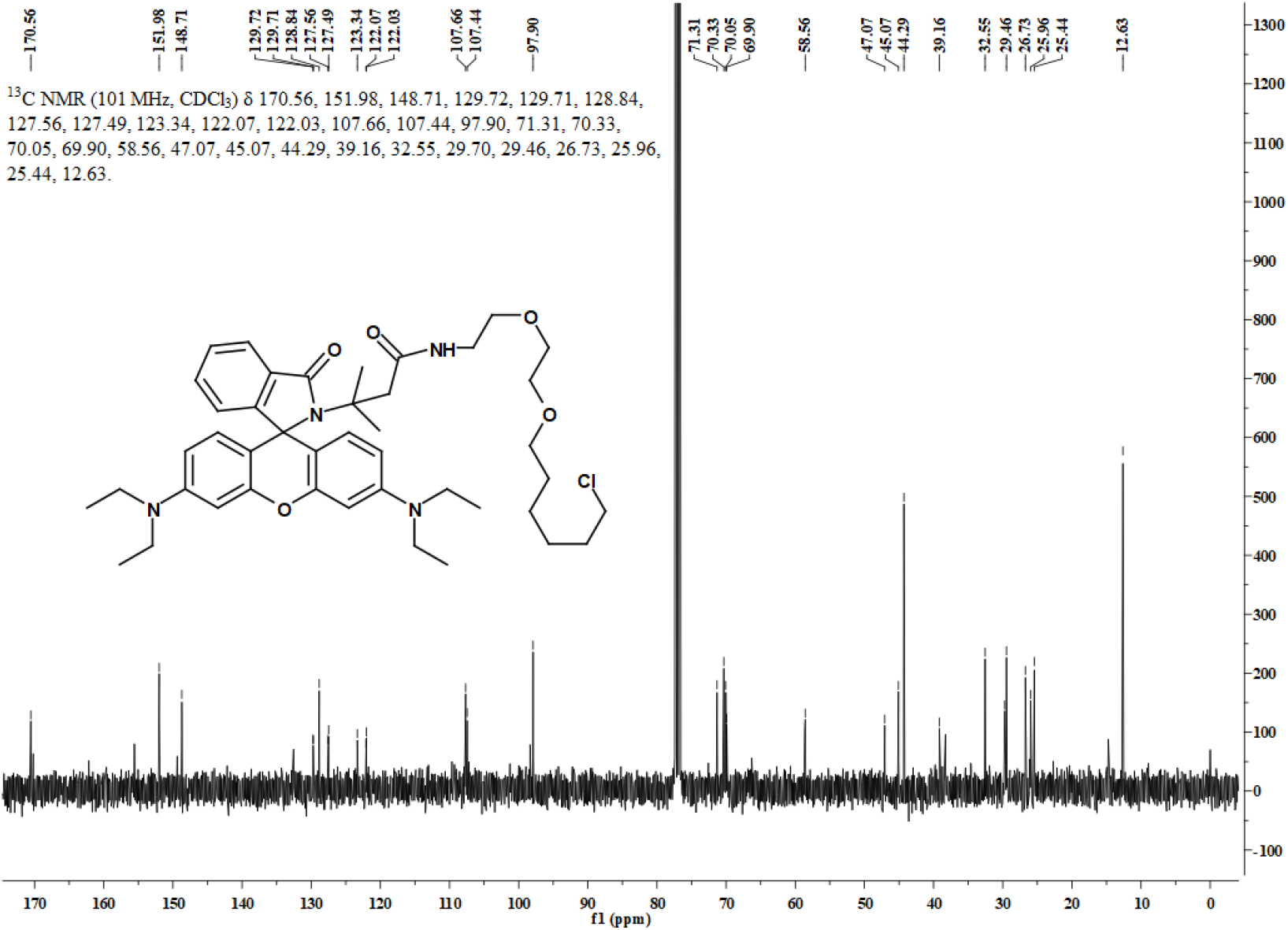
^13^C NMR spectrum of compound **N-halo-RB** in CDCl_3_.

**Figure S36.**
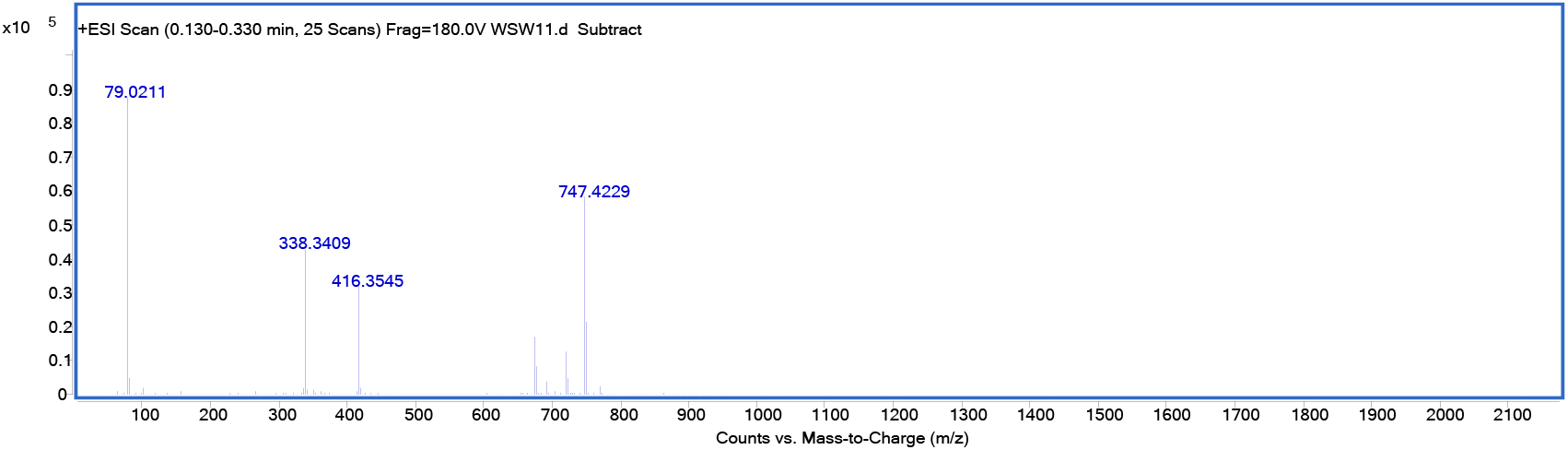
HRMS spectrum of **N-halo-RB**.

**Figure S37.**
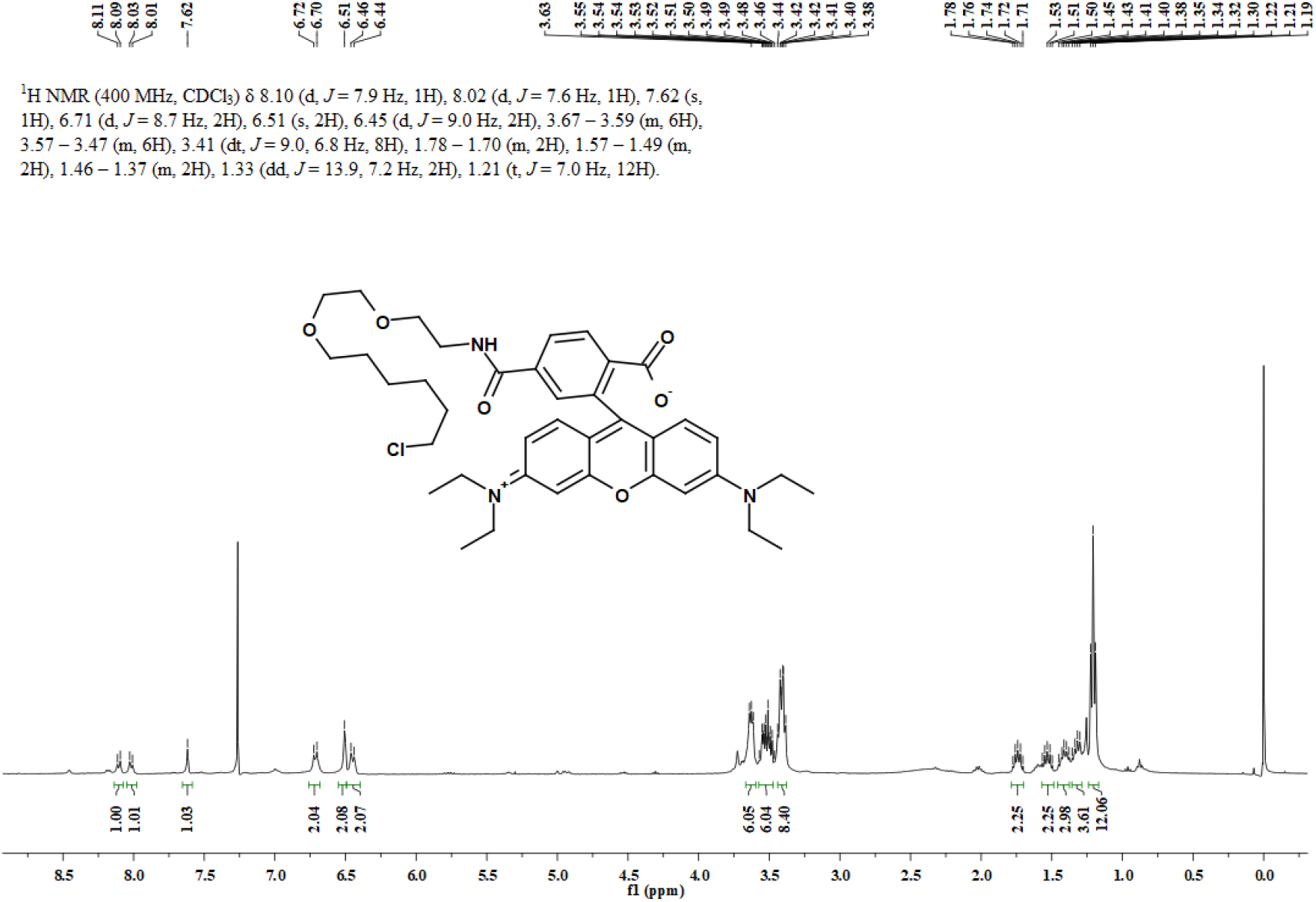
^1^H NMR spectrum of compound **9** in CDCl_3_.

**Figure S38.**
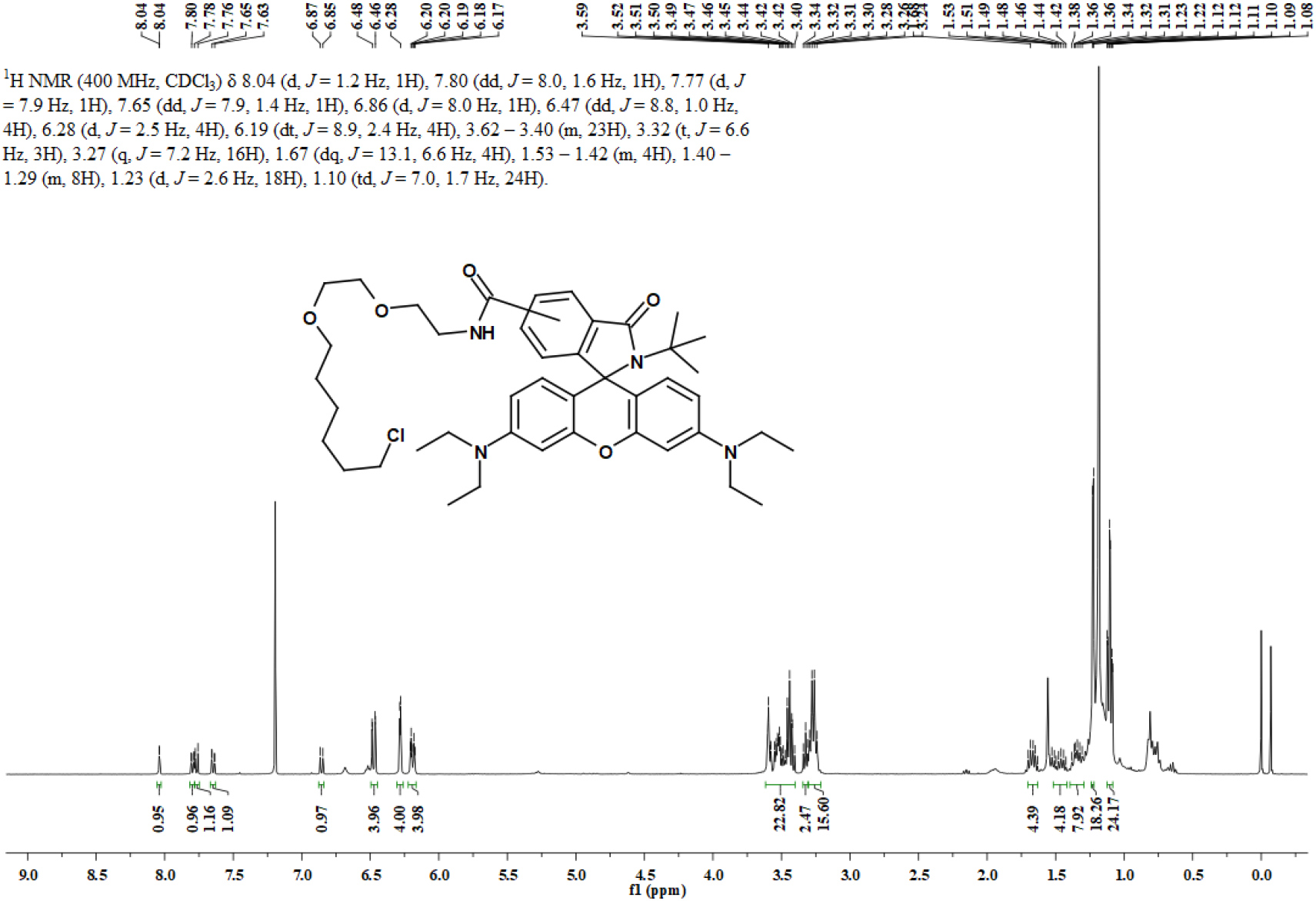
^1^H NMR spectrum of **5,6-halo-RB** in CDCl_3_.

**Figure S39.**
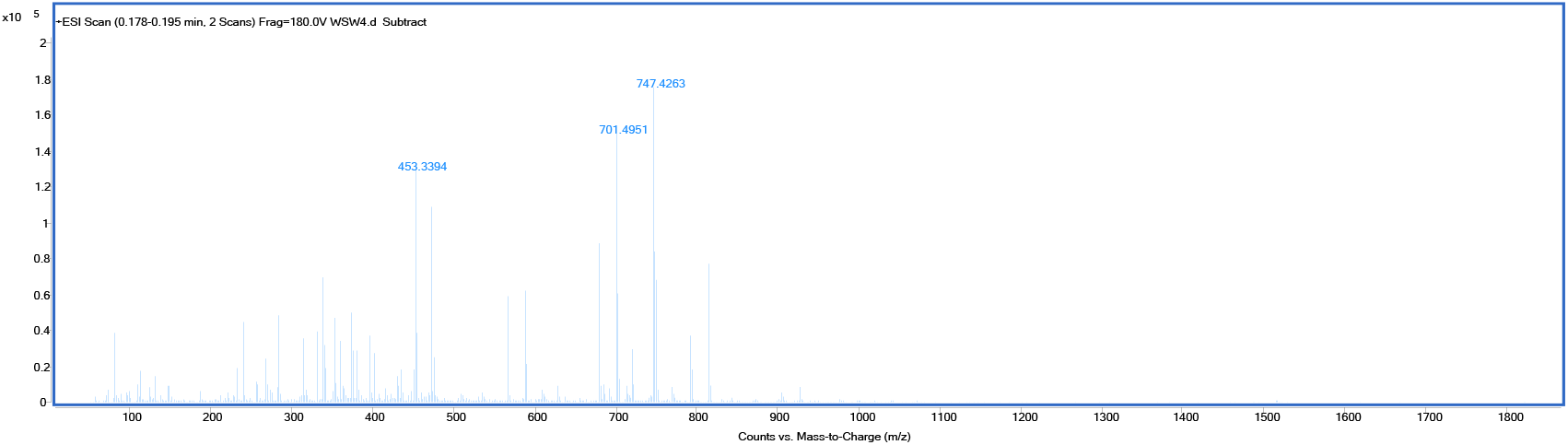
HRMS spectrum of **5,6-halo-RB**.

**Figure S40.**
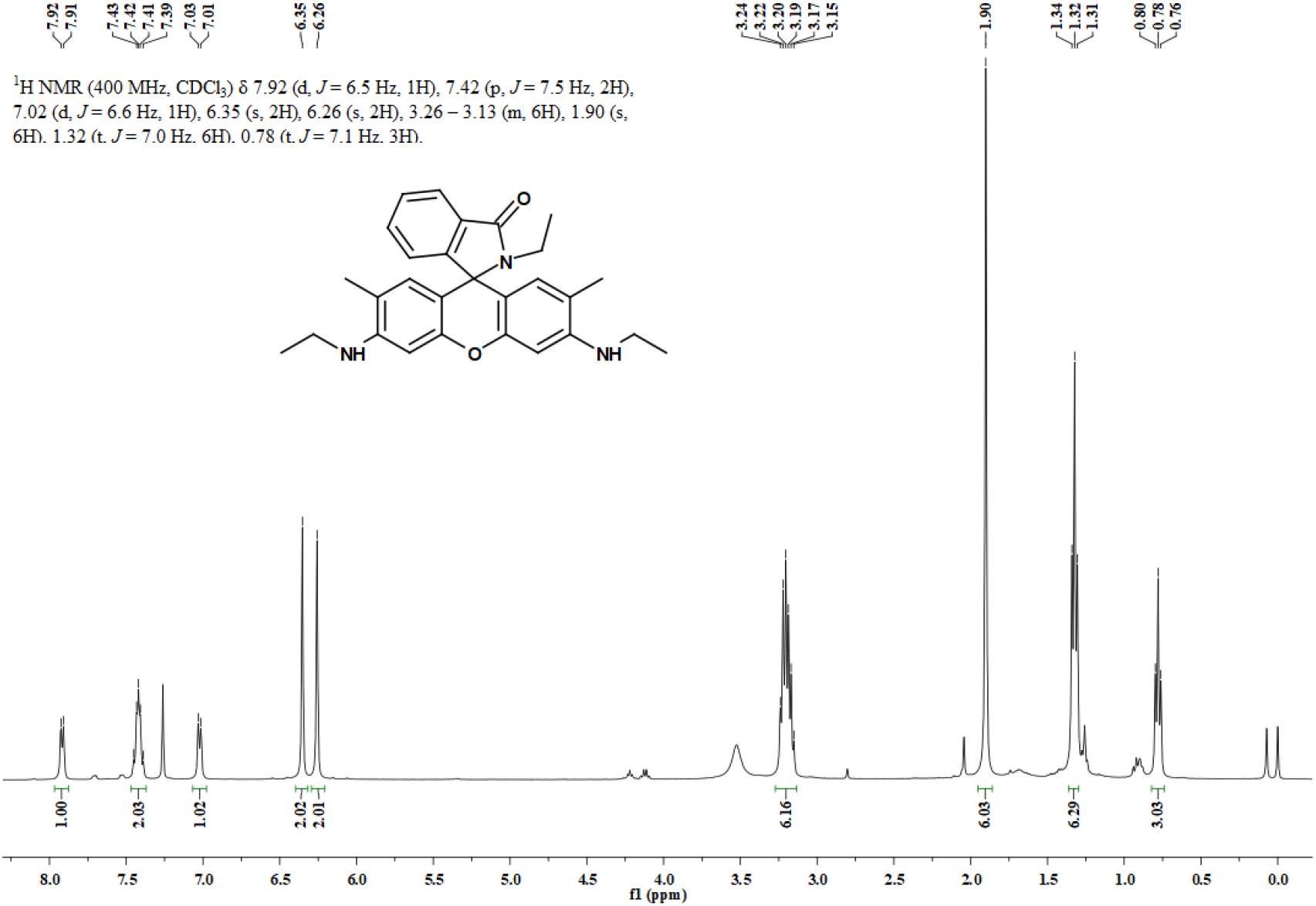
^1^H NMR spectrum of **2a** in CDCl_3_.

**Figure S41.**
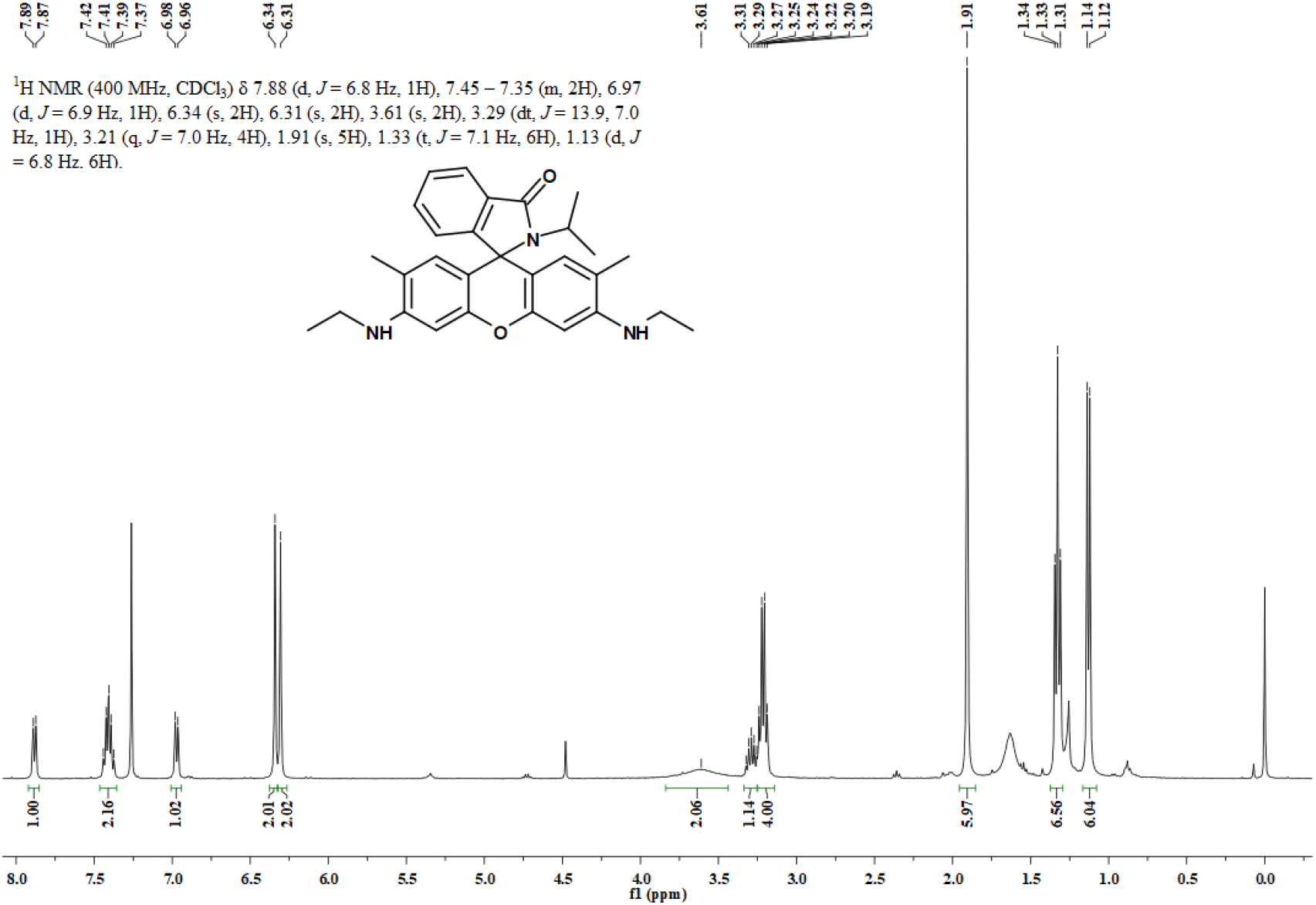
^1^H NMR spectrum of **2b** in CDCl_3_.

**Figure S42.**
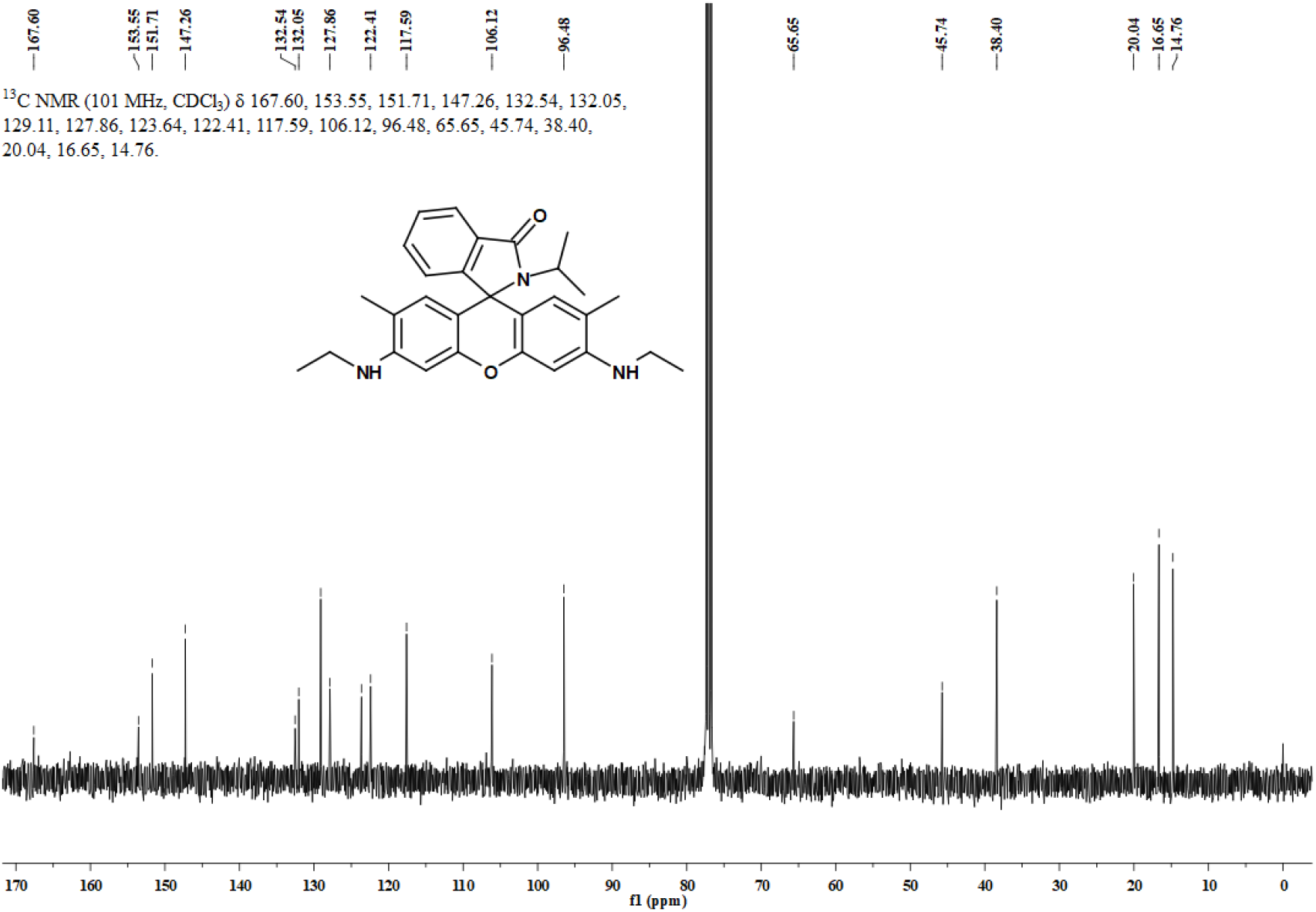
^13^C NMR spectrum of **2b** in CDCl_3_.

**Figure S43.**
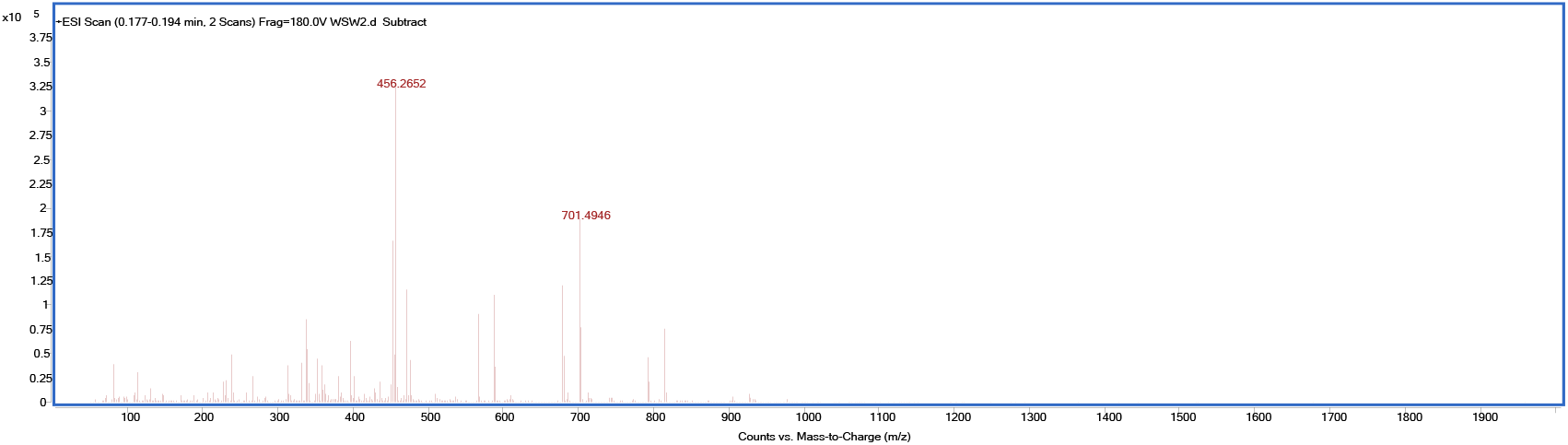
HRMS spectrum of **2b**.

**Figure S44.**
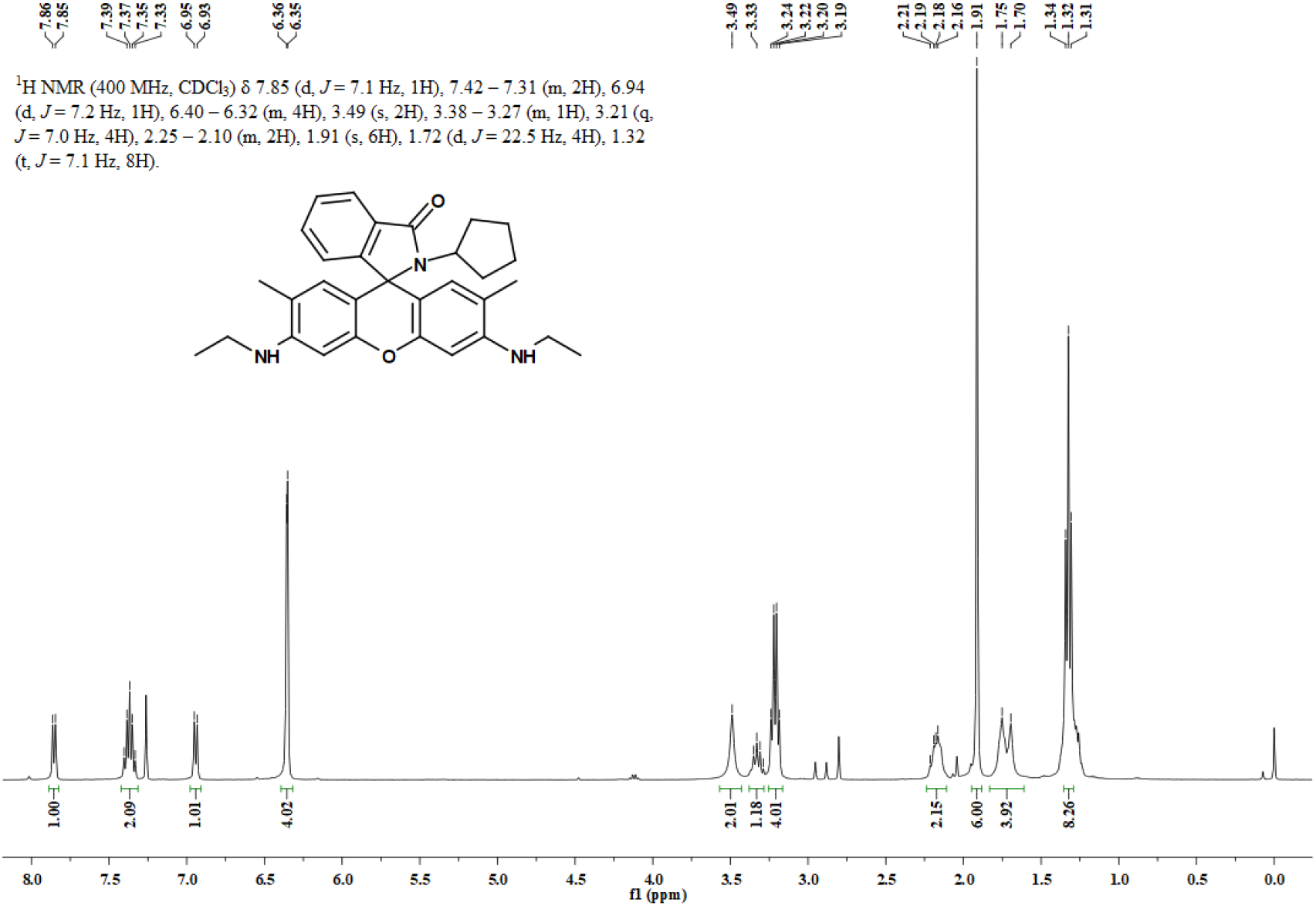
^1^H NMR spectrum of **2c** in CDCl_3_.

**Figure S45.**
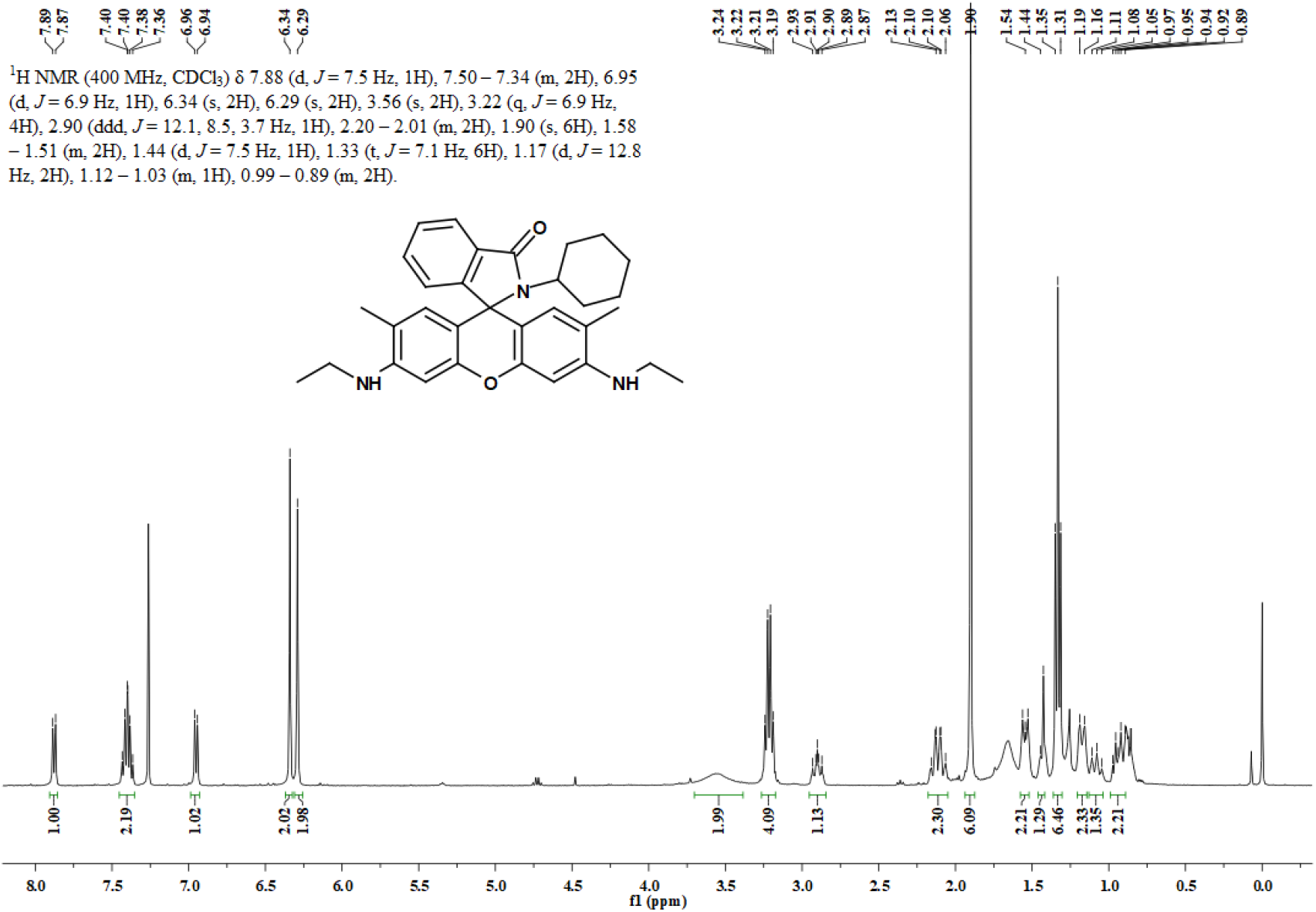
^1^H NMR spectrum of **2d** in CDCl_3_.

**Figure S46.**
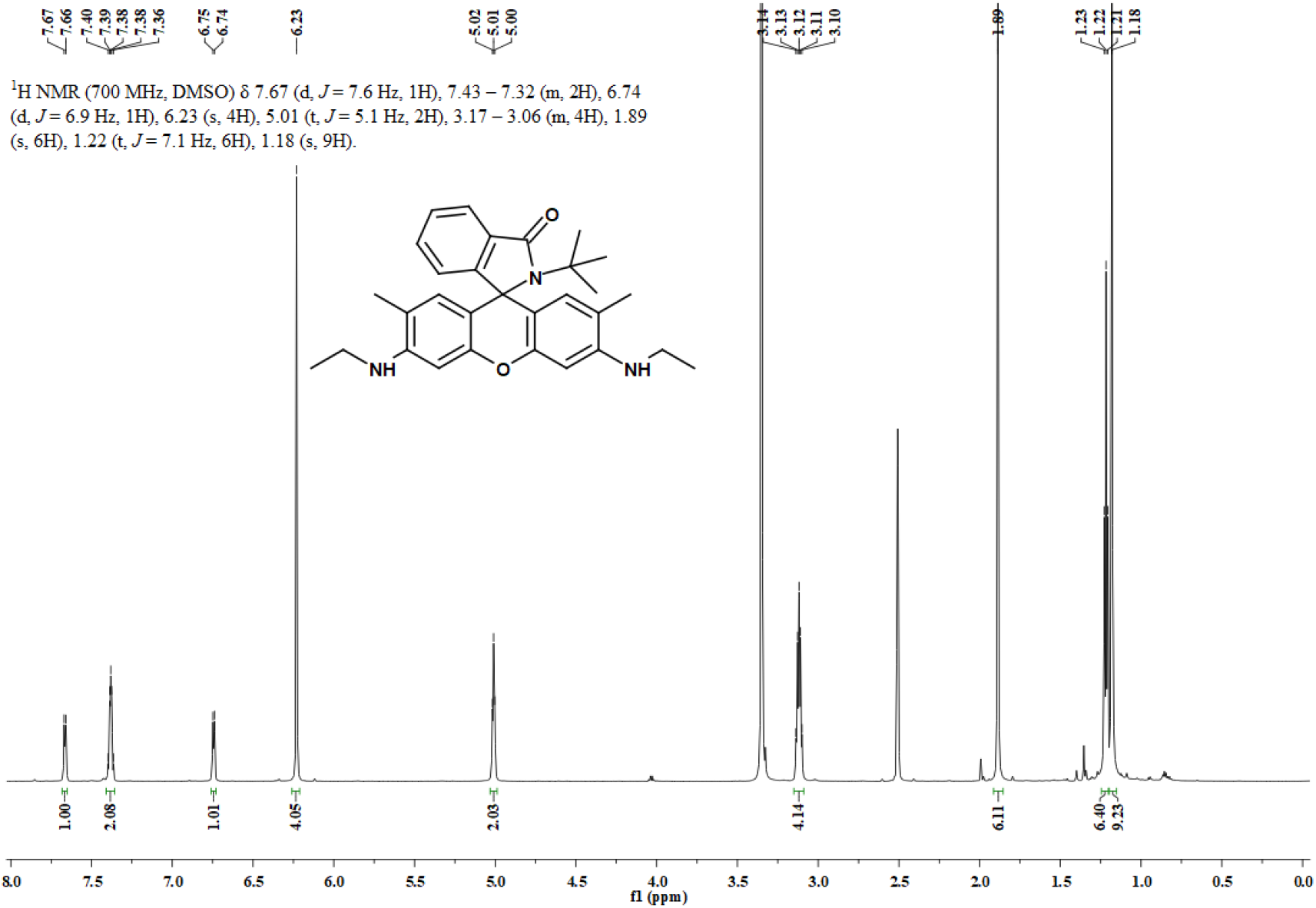
^1^H NMR spectrum of **2e** in DMSO-*d6*.

**Figure S47.**
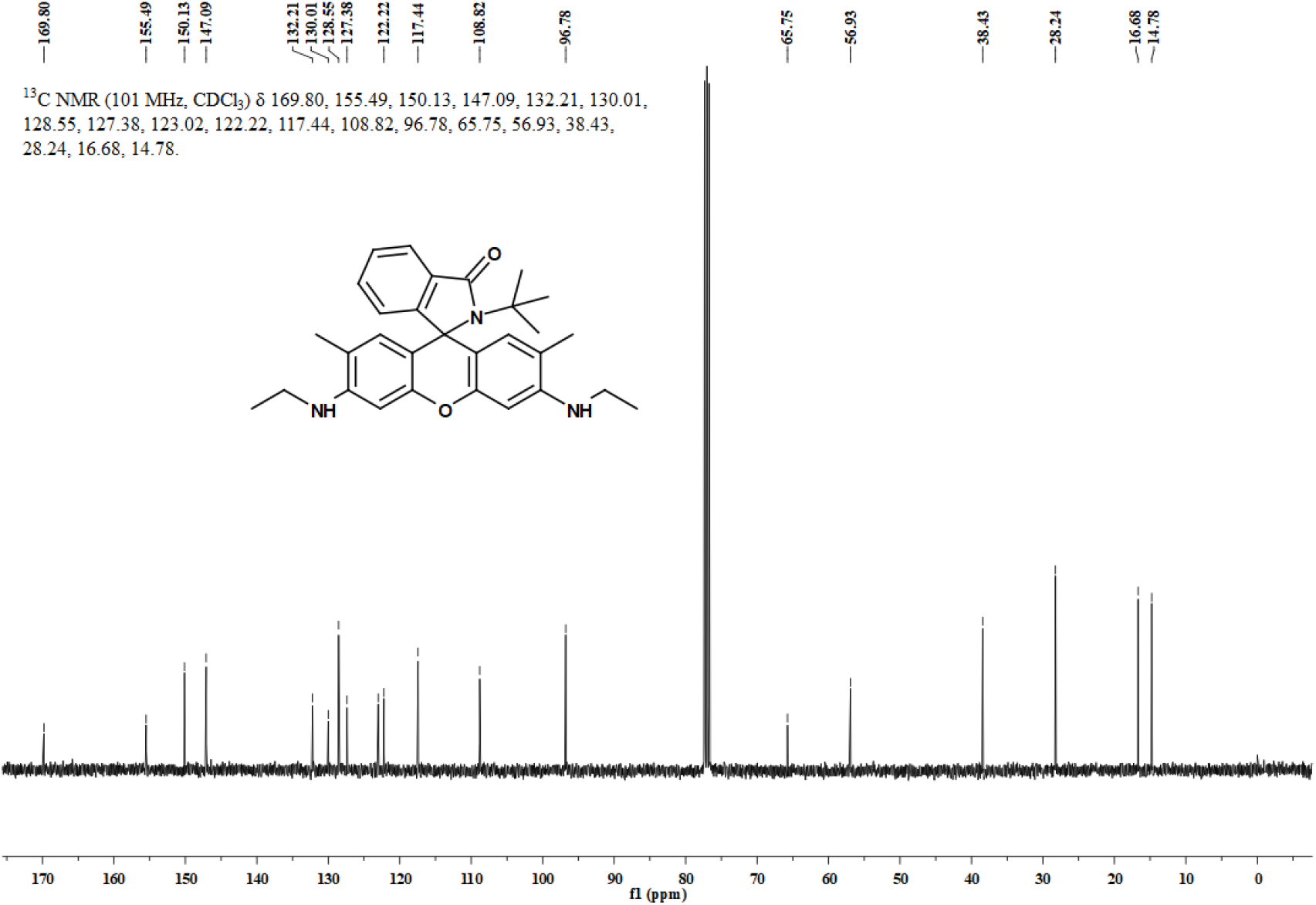
^13^C NMR spectrum of **2e** in CDCl_3_.

**Figure S48.**
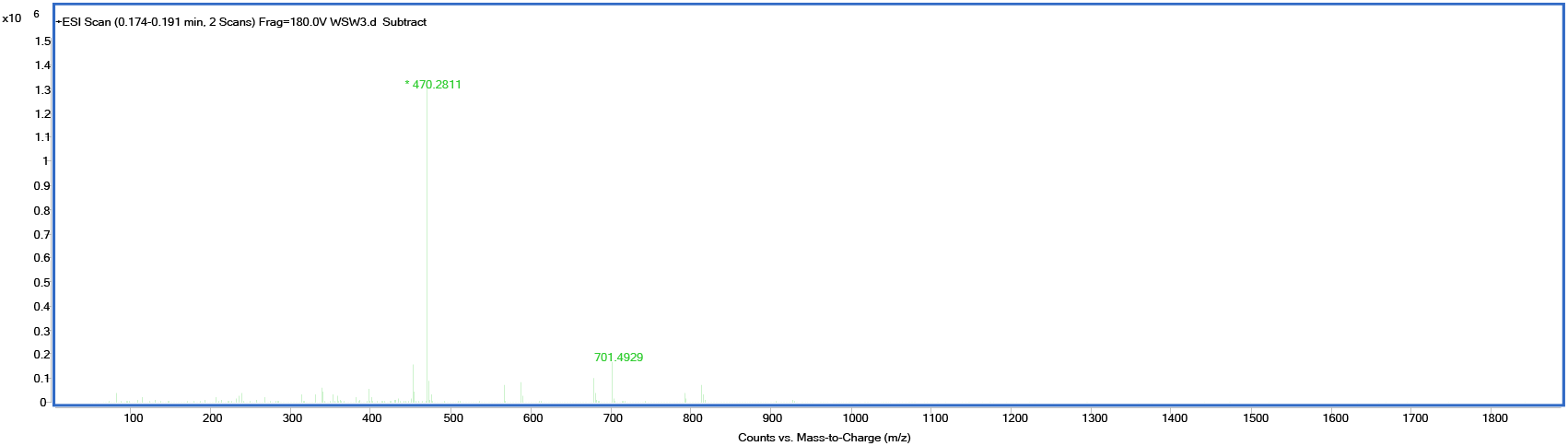
HRMS spectrum of **2e**.

**Figure S49.**
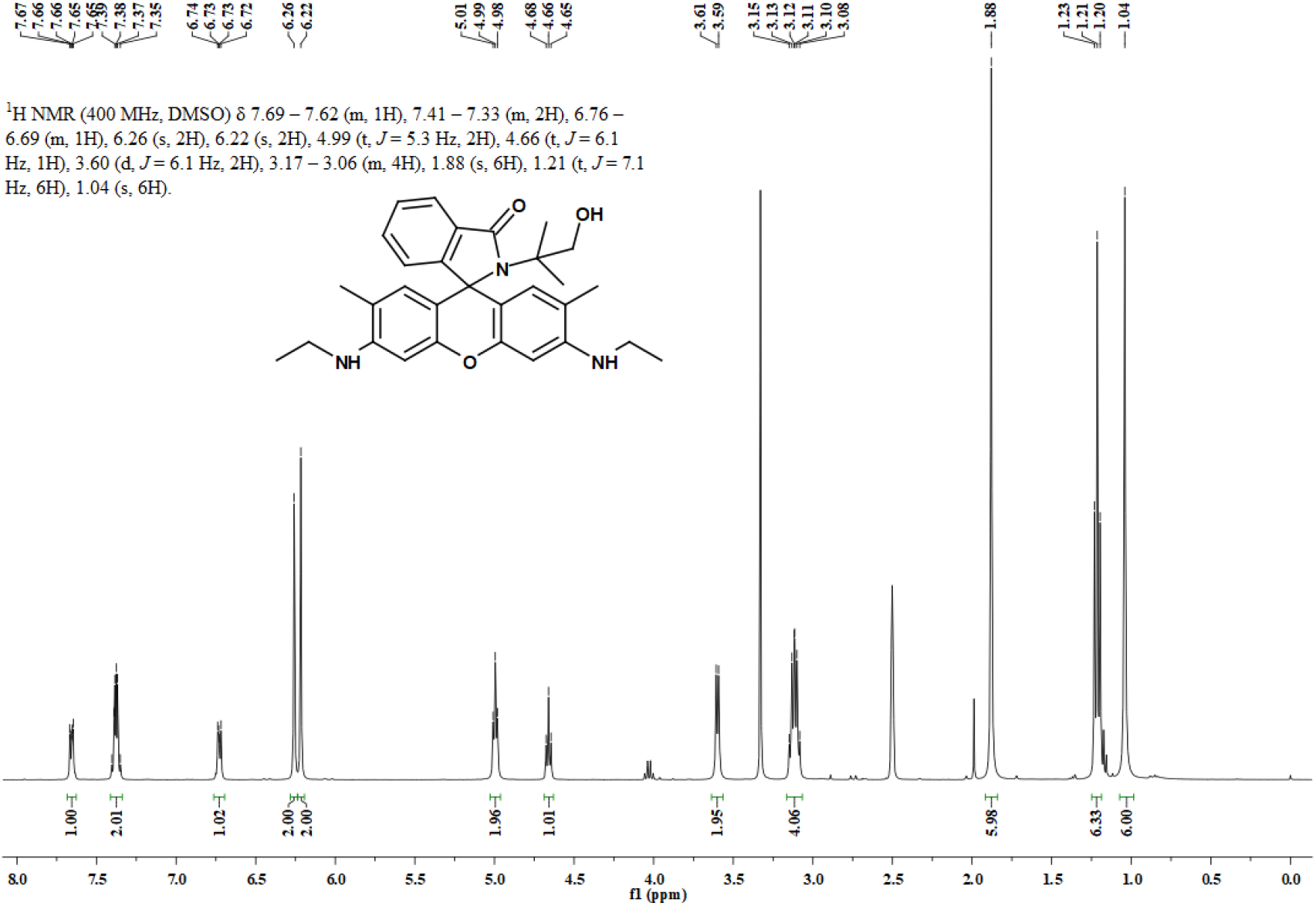
^1^H NMR spectrum of **2f** in DMSO-*d6*.

**Figure S50.**
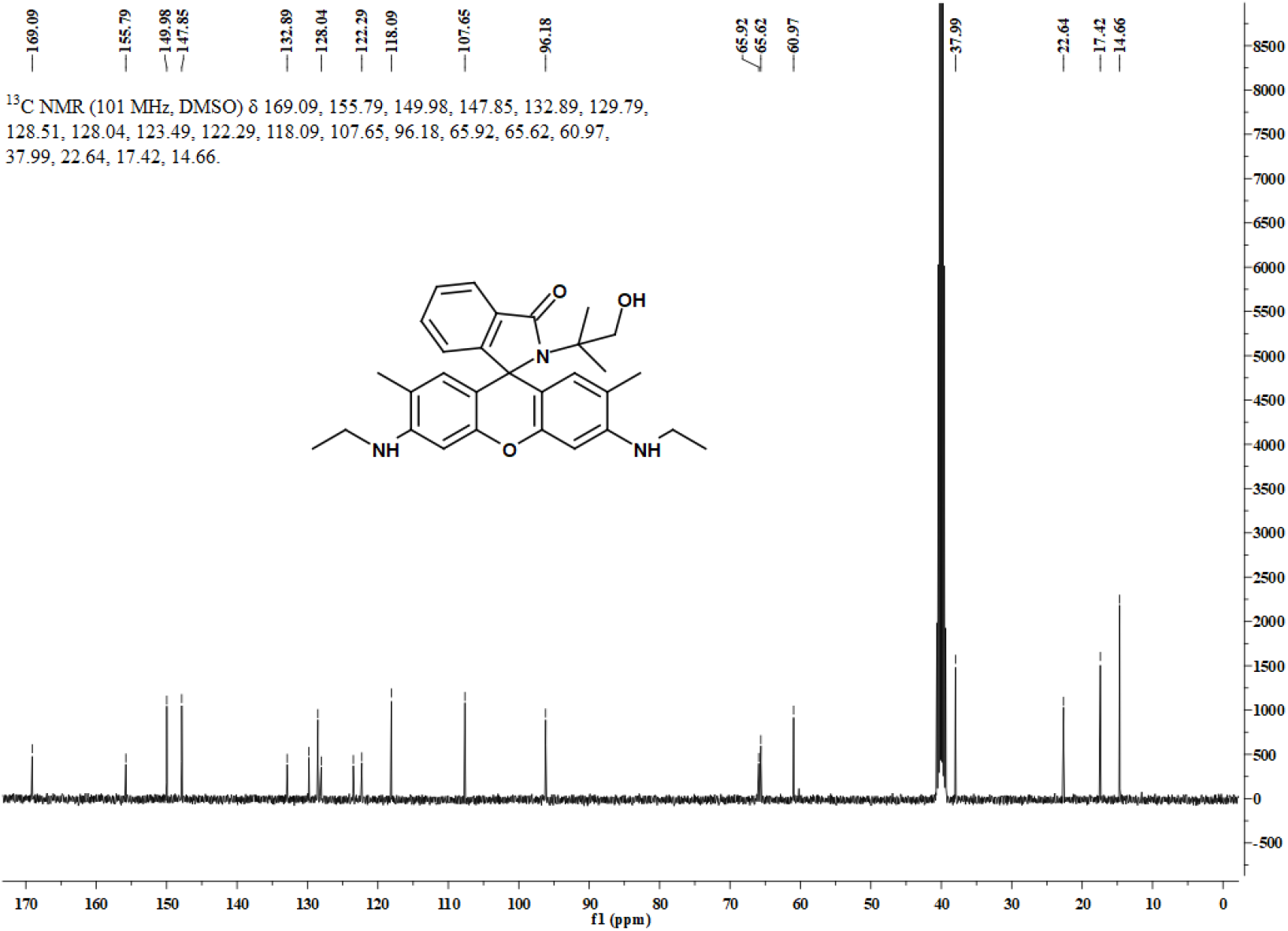
^13^C NMR spectrum of **2e** in DMSO-*d6*.

**Figure S51.**
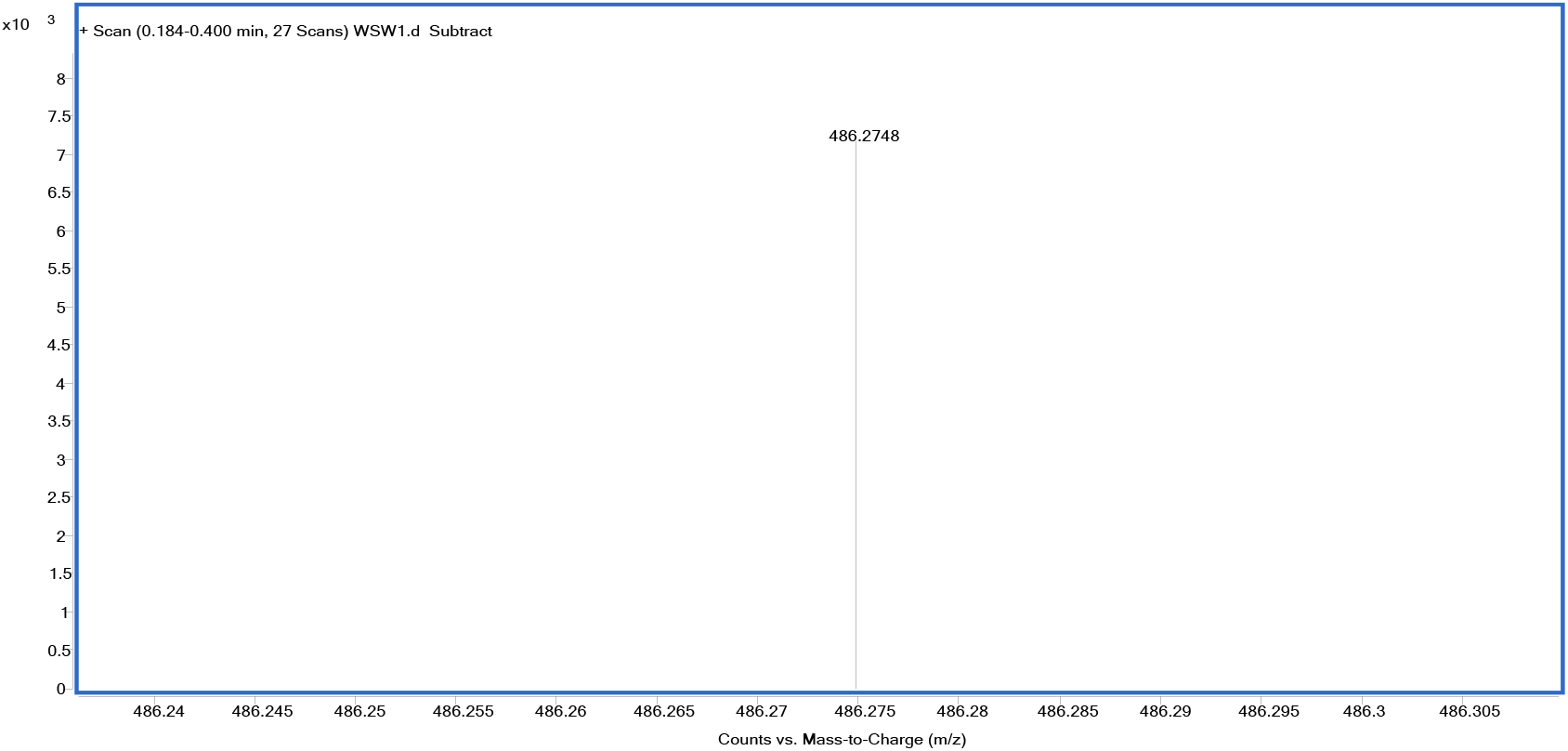
HRMS spectrum of **2e**.

**Figure S52.**
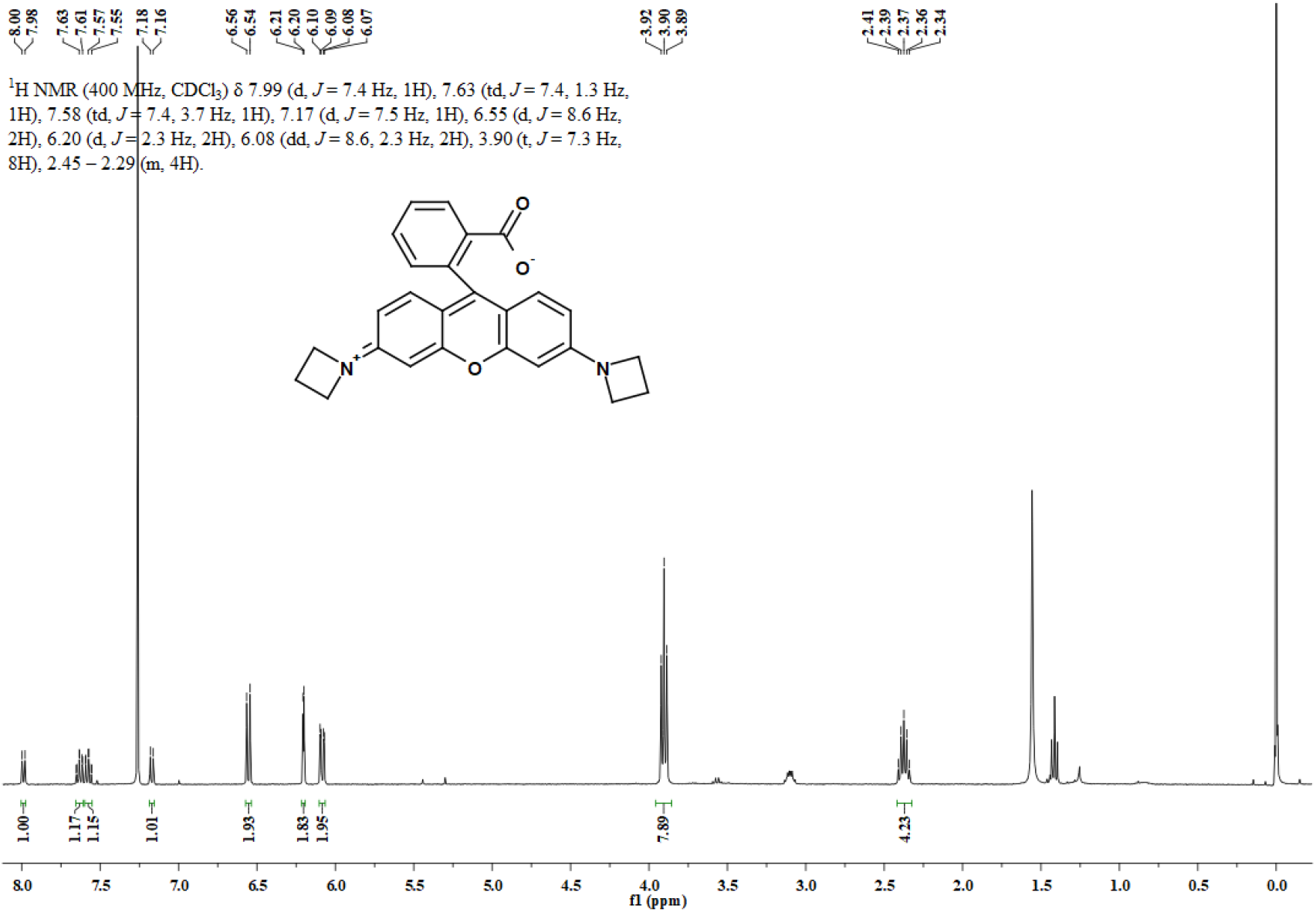
^1^H NMR spectrum of compound **11** in CDCl_3_.

**Figure S53.**
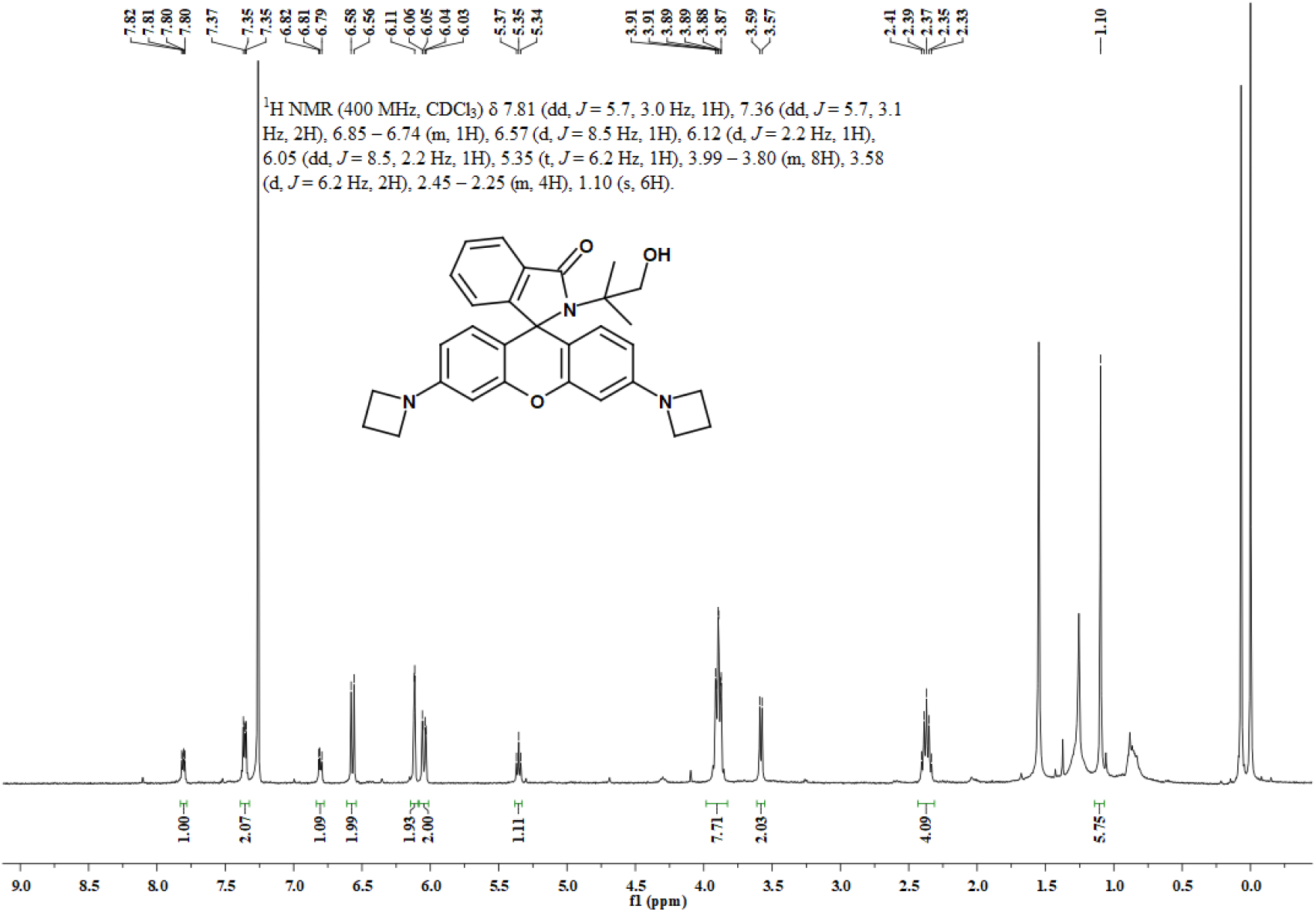
^1^H NMR spectrum of **3a** in CDCl_3_.

**Figure S54.**
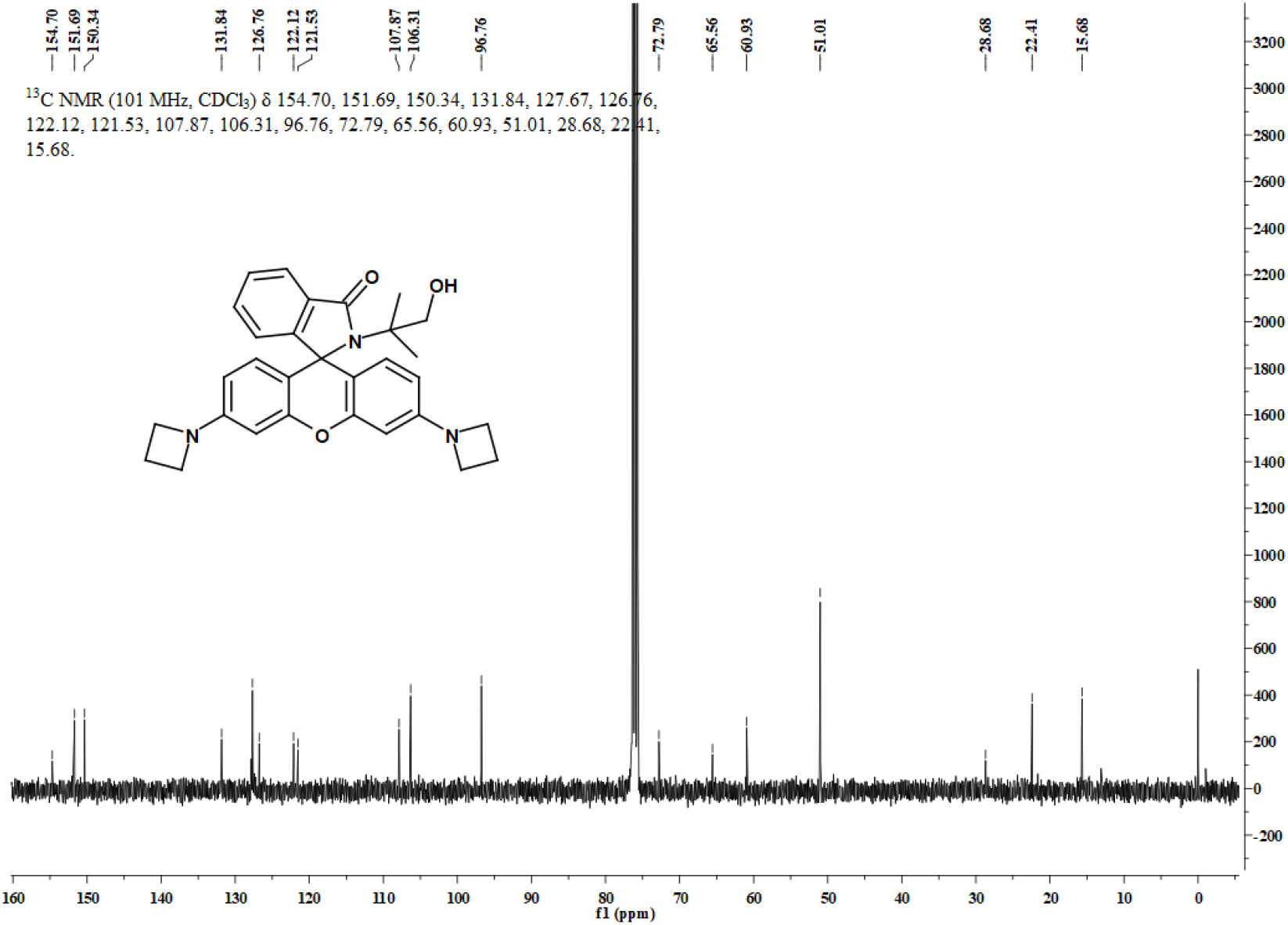
^13^C NMR spectrum of **3a** in CDCl_3_.

**Figure S55.**
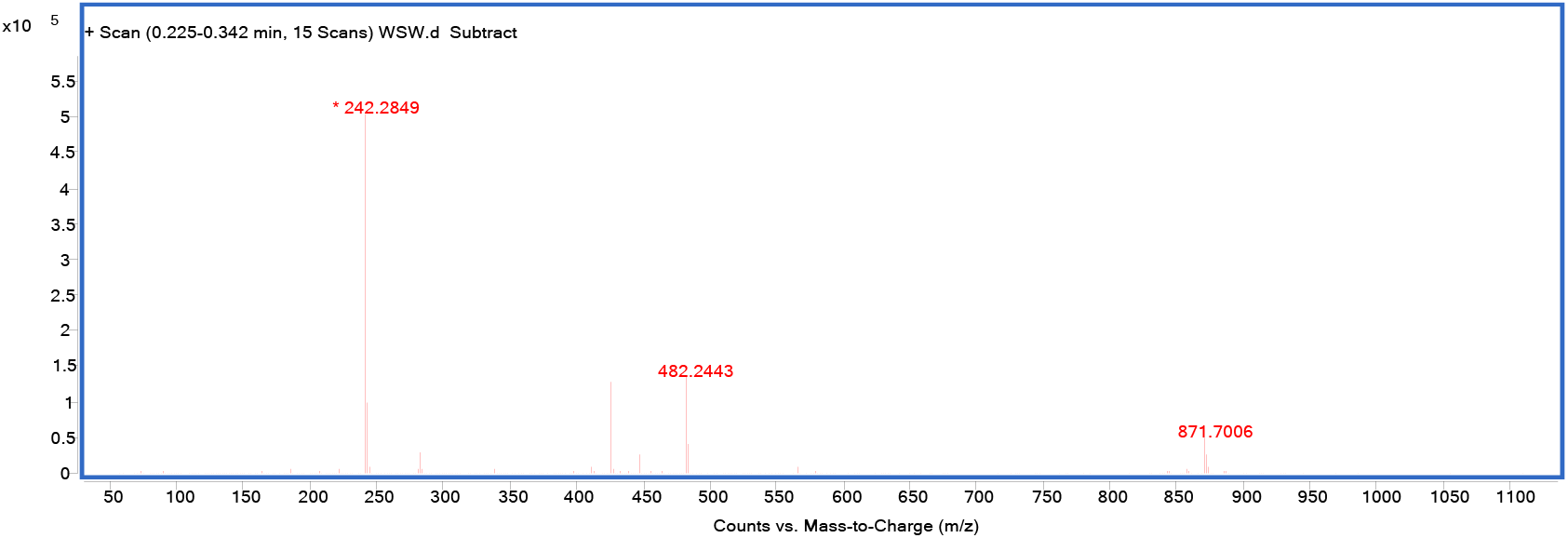
HRMS spectrum of **3a**.

**Figure S56.**
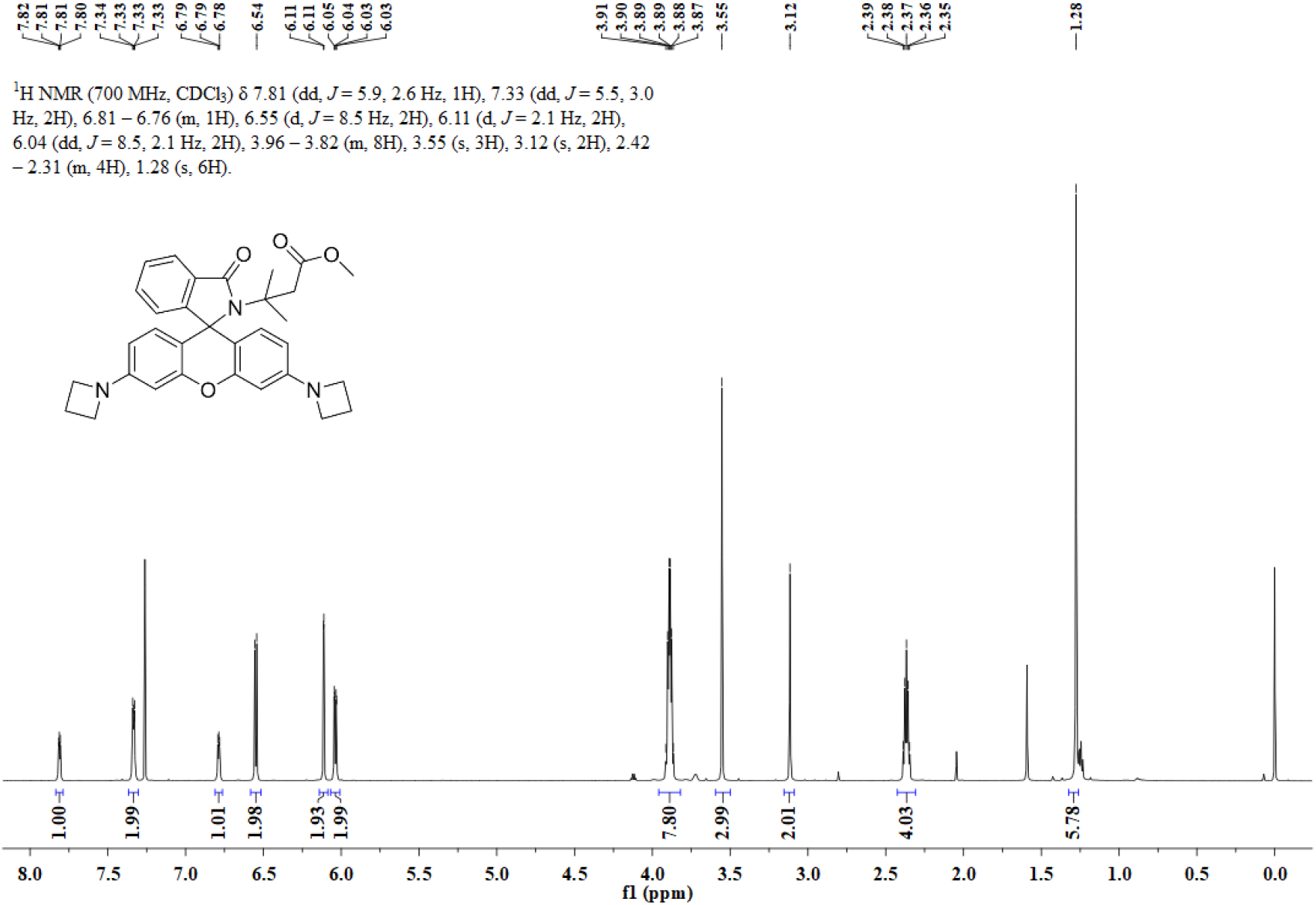
^1^H NMR spectrum of compound **12** in CDCl_3_.

**Figure S57.**
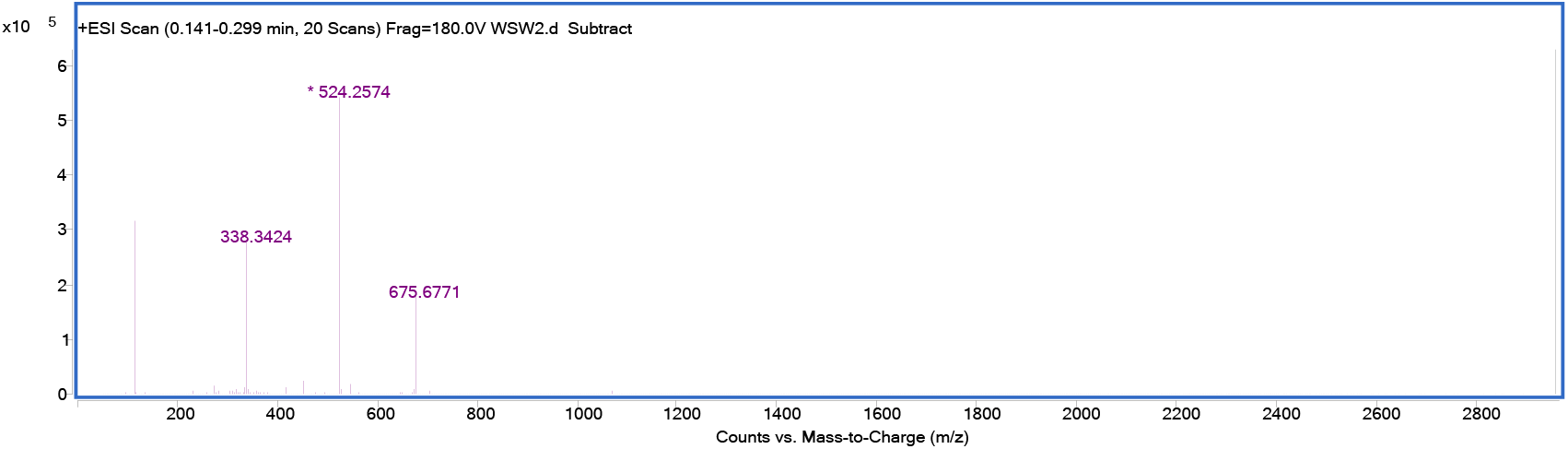
HRMS spectrum of compound **12**.

**Figure S58.**
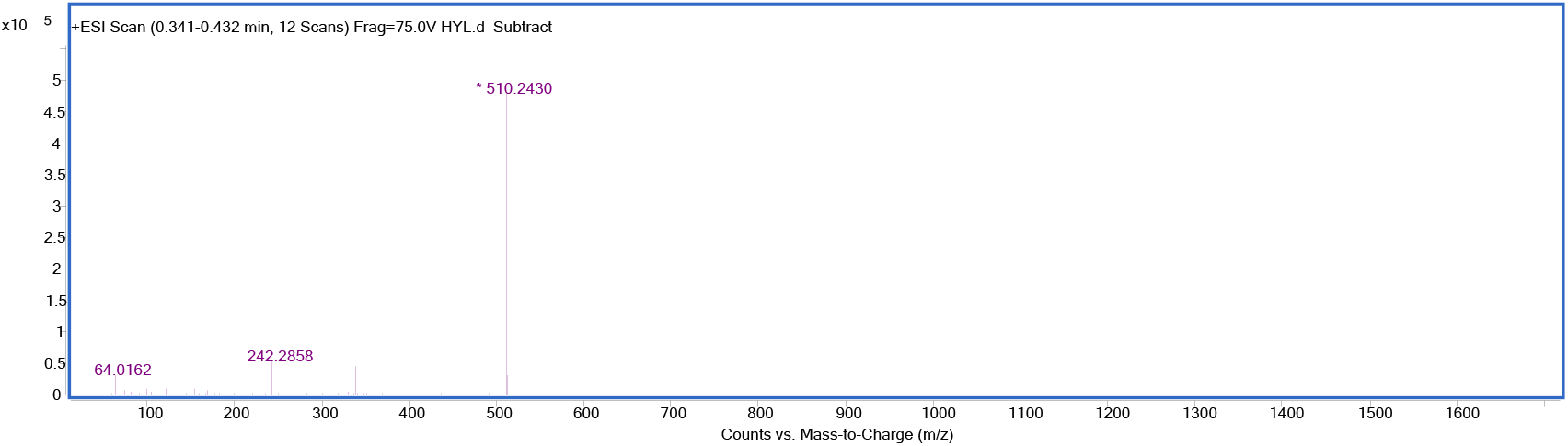
HRMS spectrum of compound **13**.

**Figure S59.**
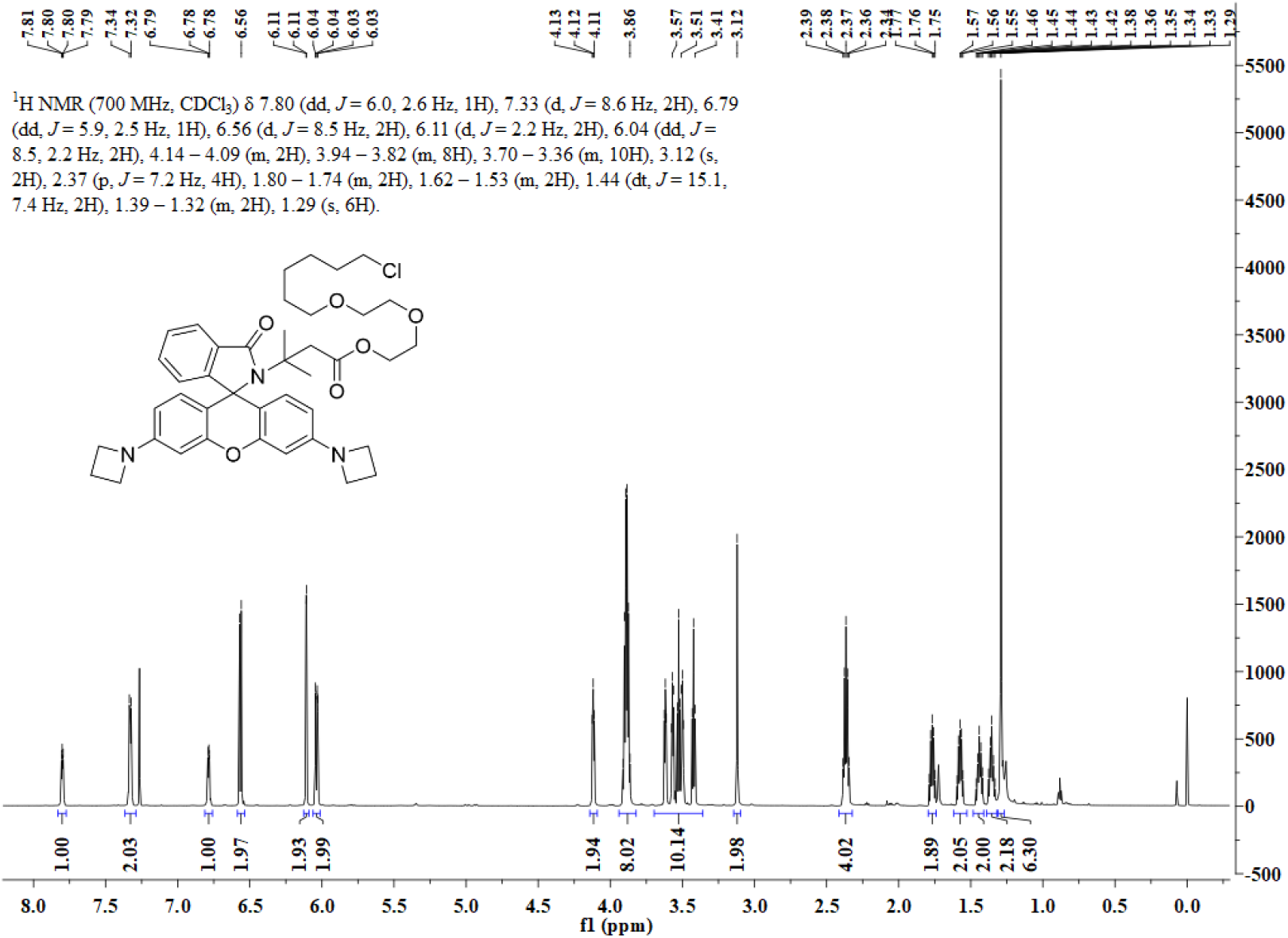
^1^H NMR spectrum of **O-halo-JF549** in CDCl_3_.

**Figure S60.**
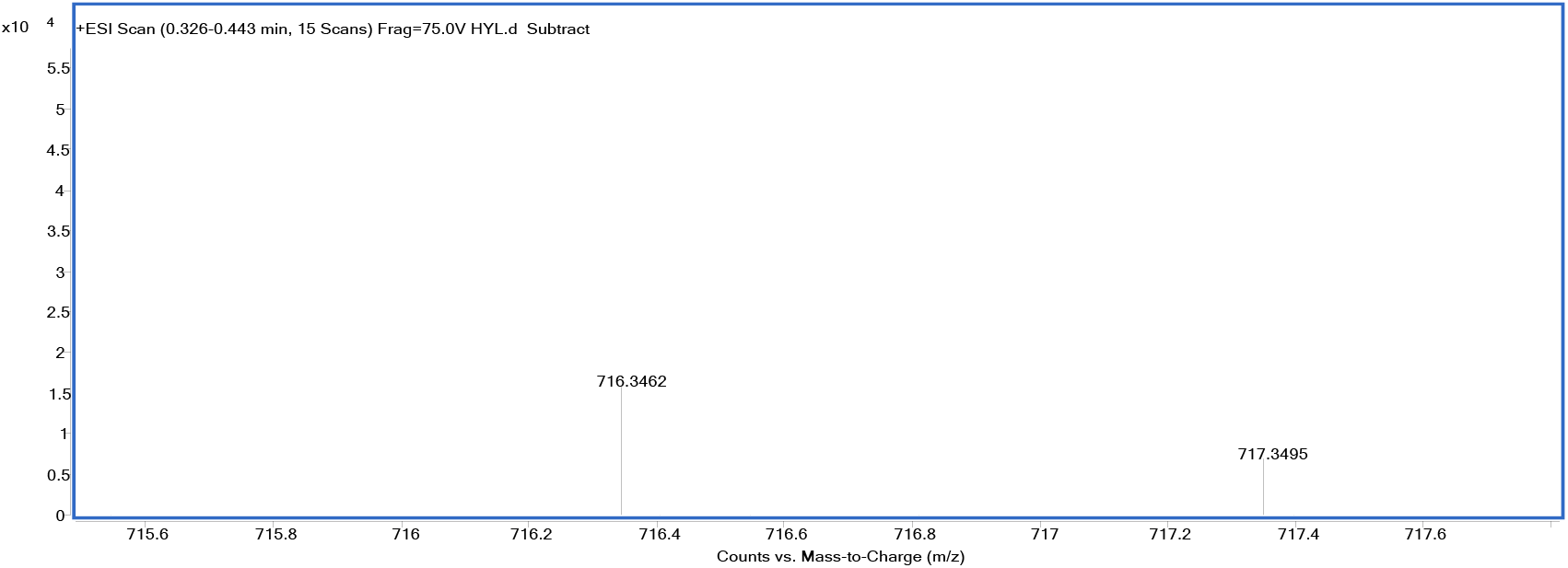
HRMS spectrum of **O-halo-JF549**.

**Figure S61.**
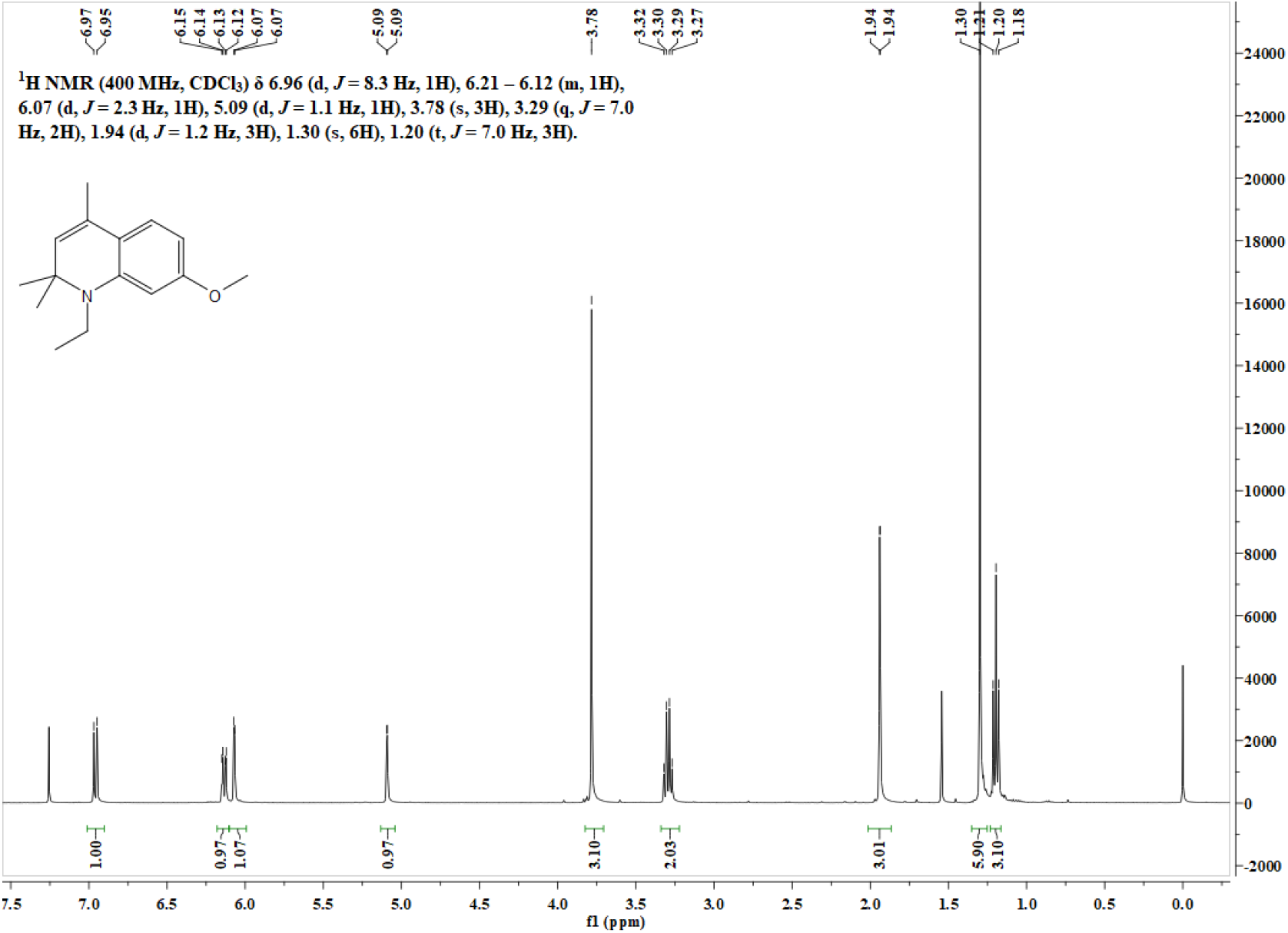
^1^H NMR spectrum of **15** in CDCl_3_.

**Figure S62.**
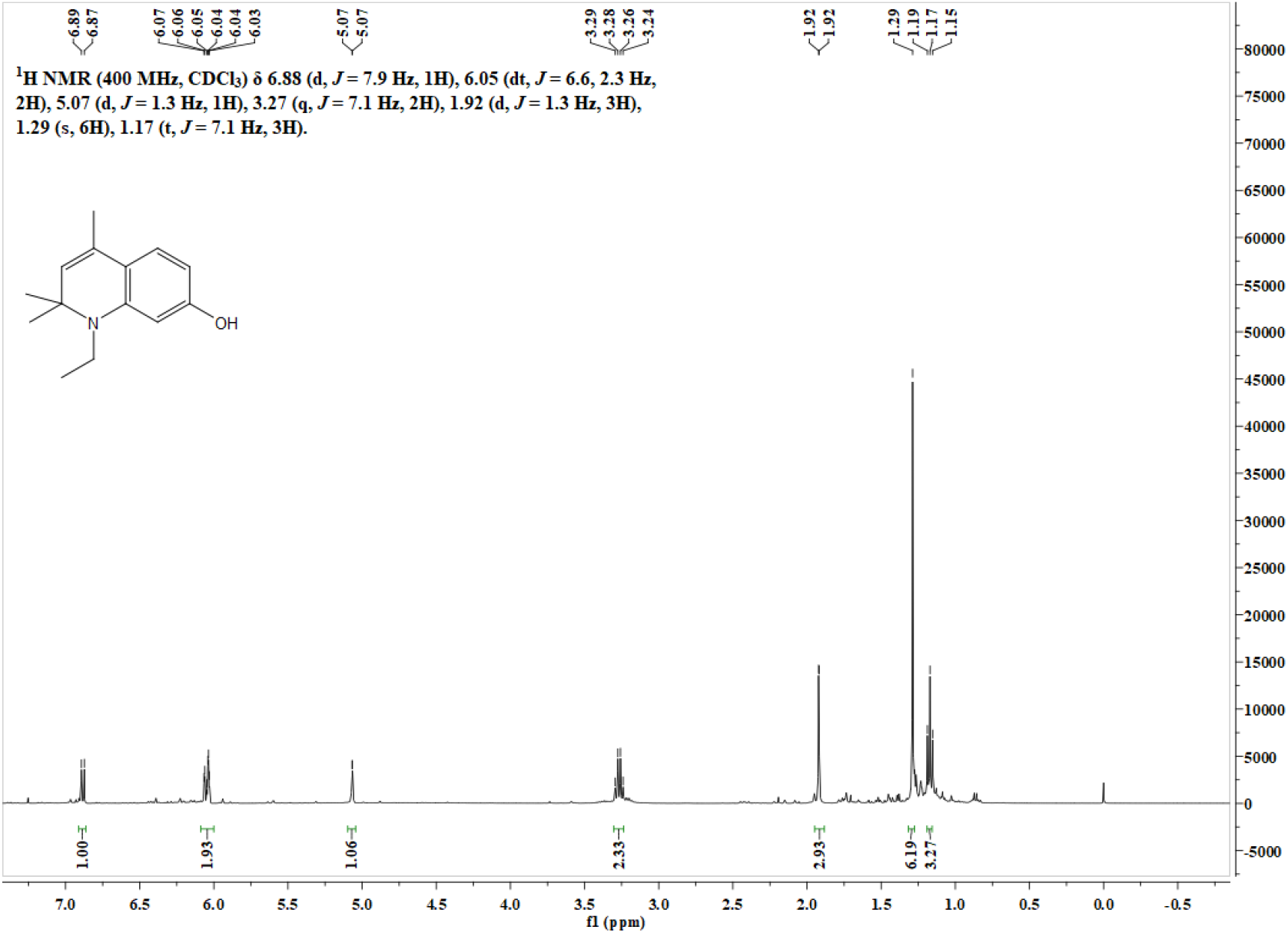
^1^H NMR spectrum of **16** in CDCl_3_.

**Figure S63.**
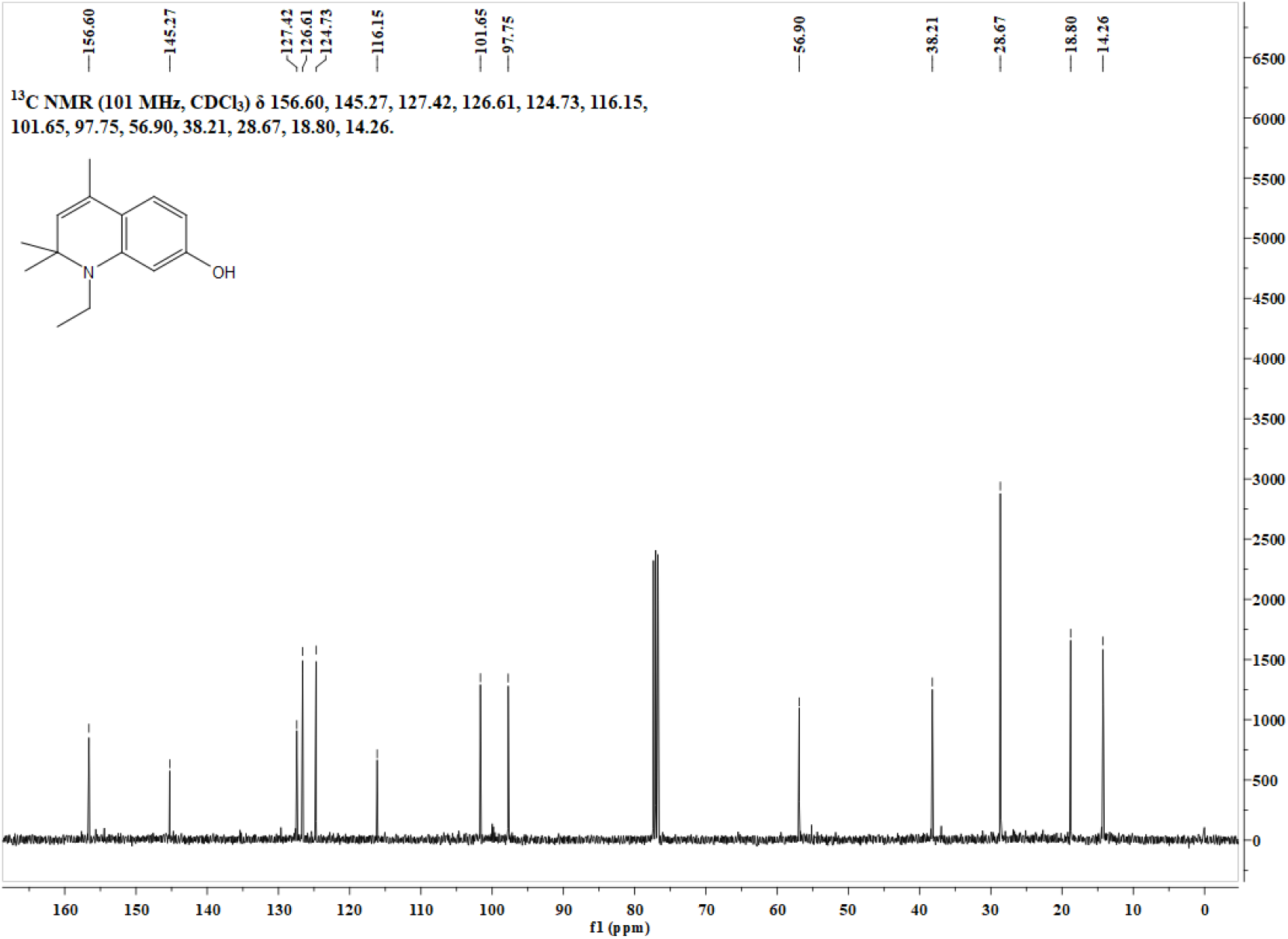
^13^C NMR spectrum of **16** in CDCl_3_.

**Figure S64.**
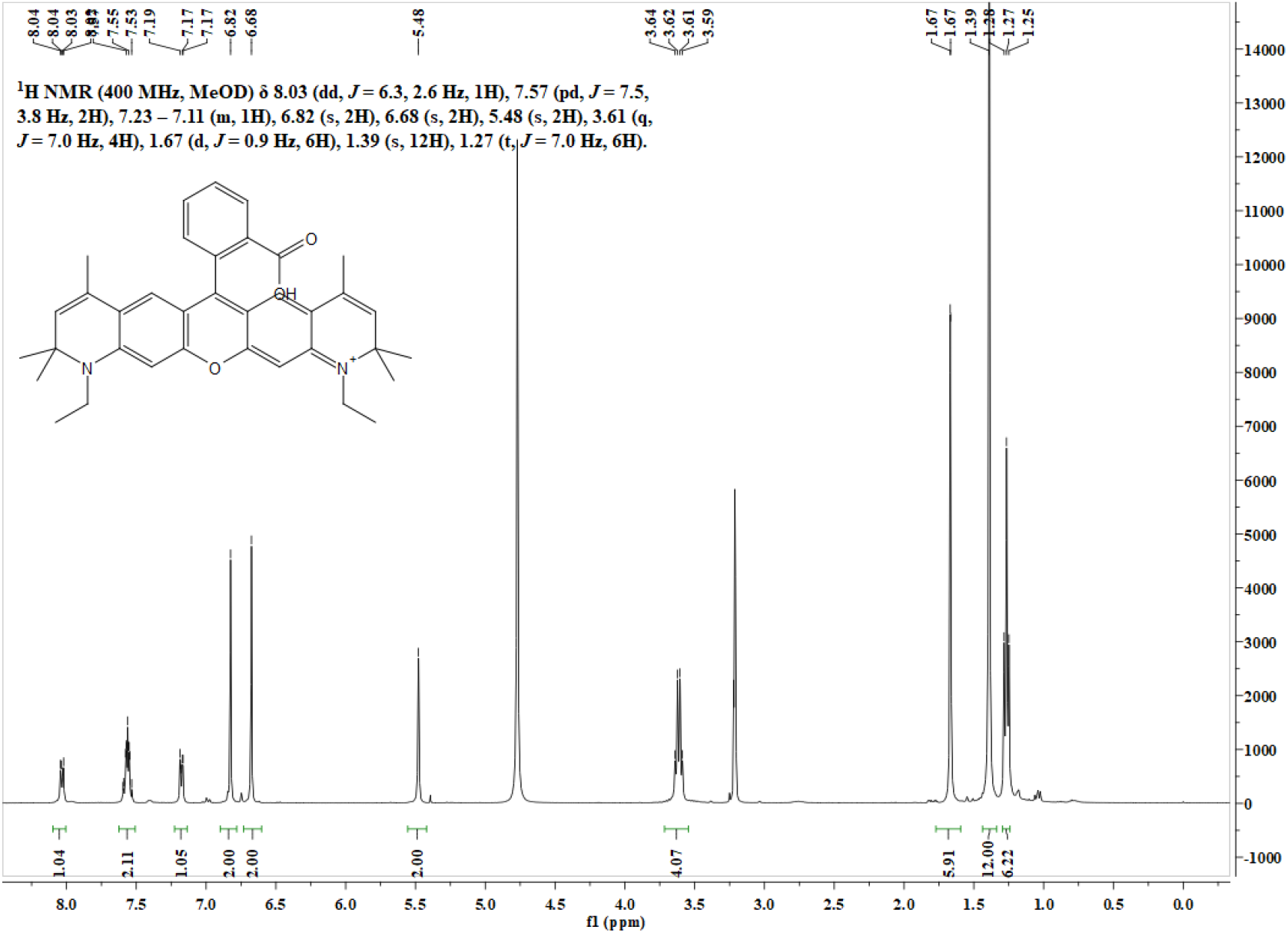
^1^H NMR spectrum of **17** in CD_3_OD.

**Figure S65.**
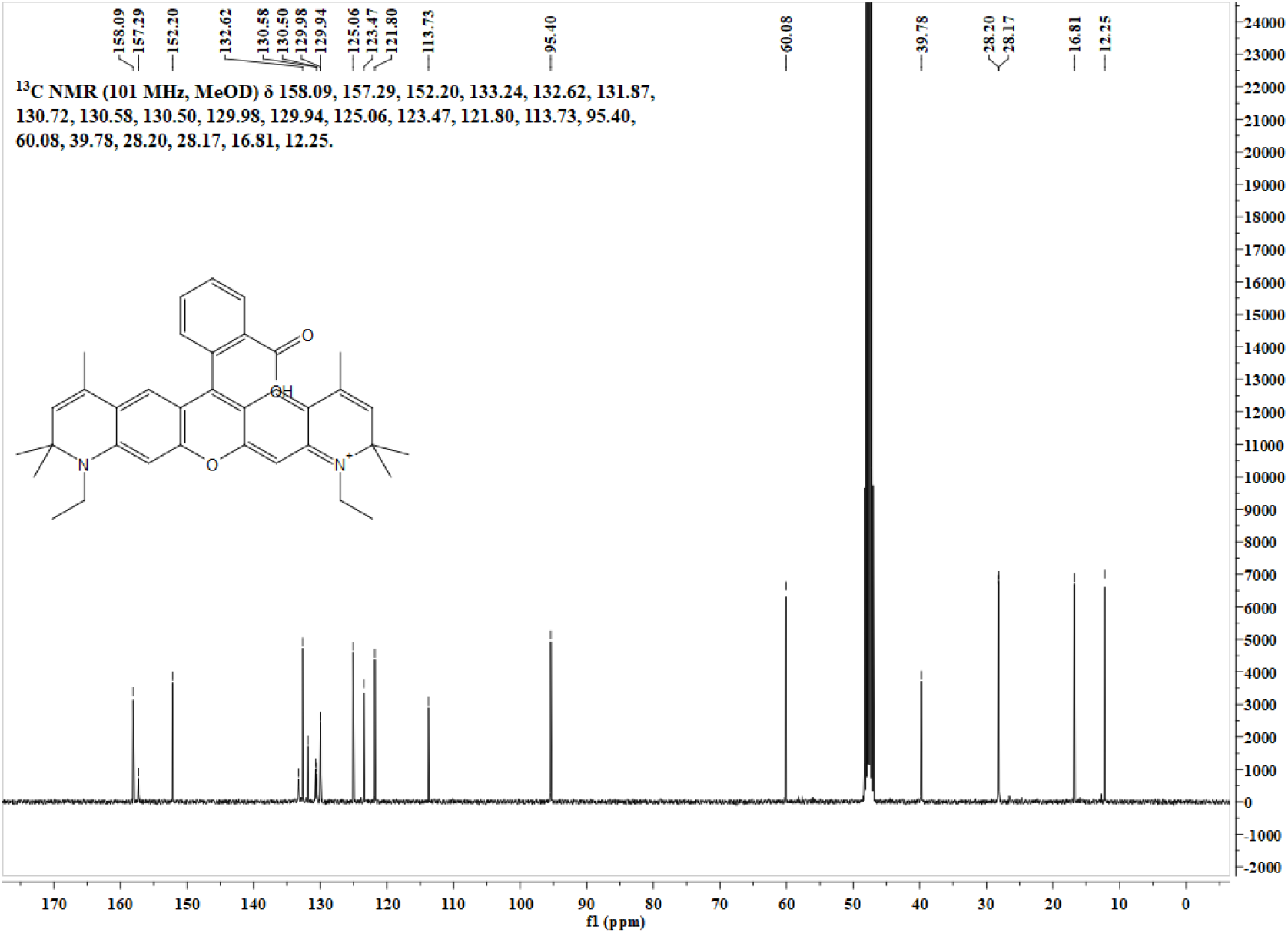
^13^C NMR spectrum of **17** in CD_3_OD.

**Figure S66.**
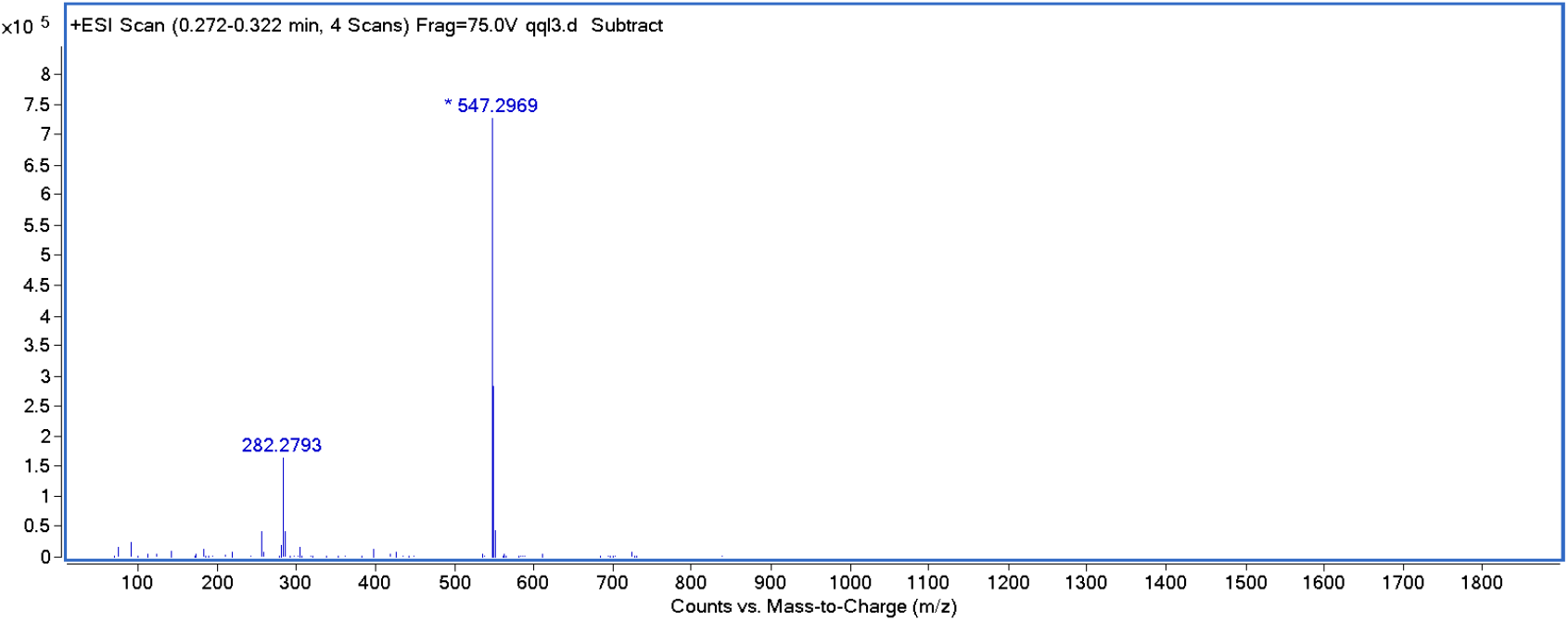
HRMS spectrum of **17**.

**Figure S67.**
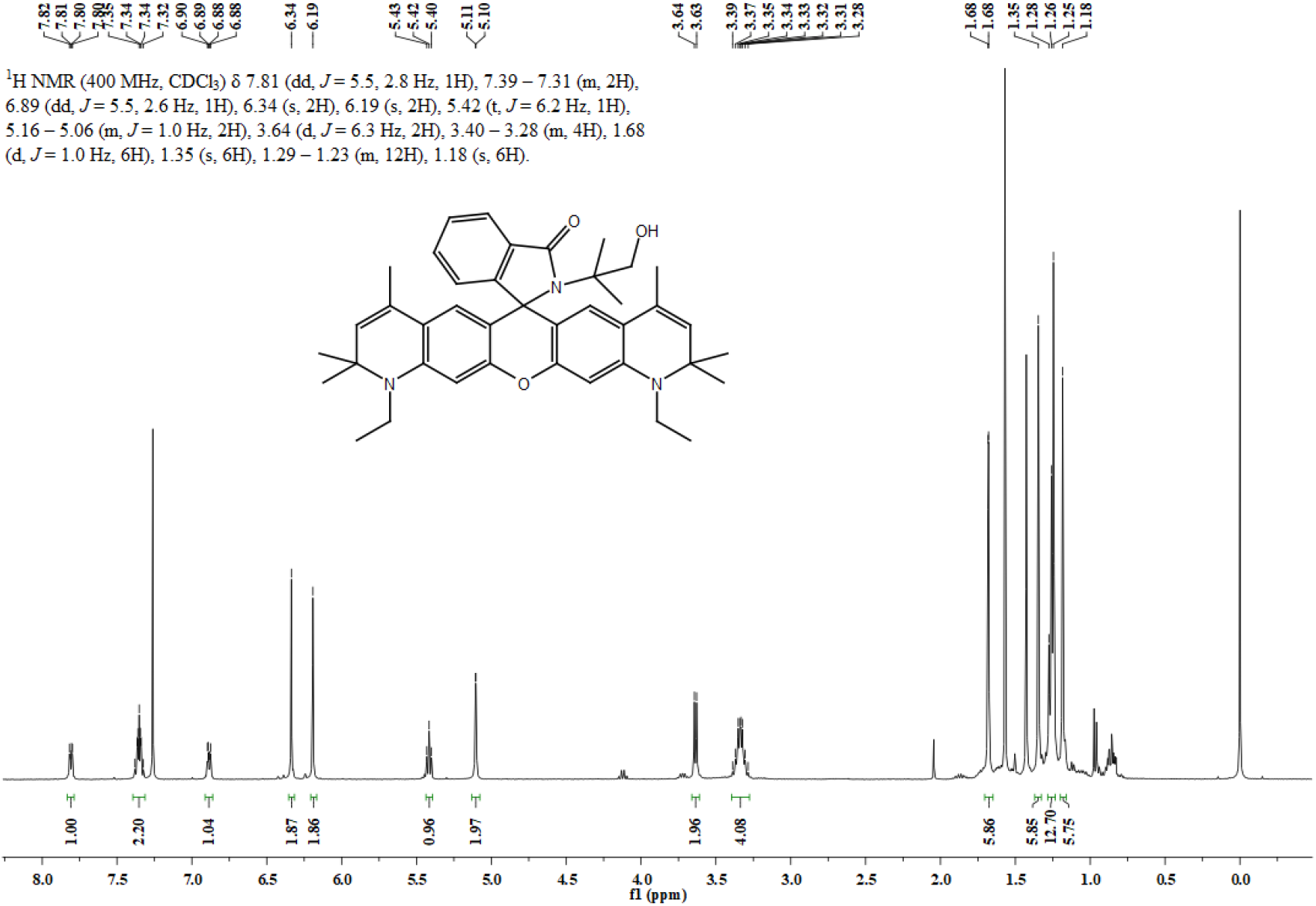
^1^H NMR spectrum of **4a** in CDCl_3_.

**Figure S68.**
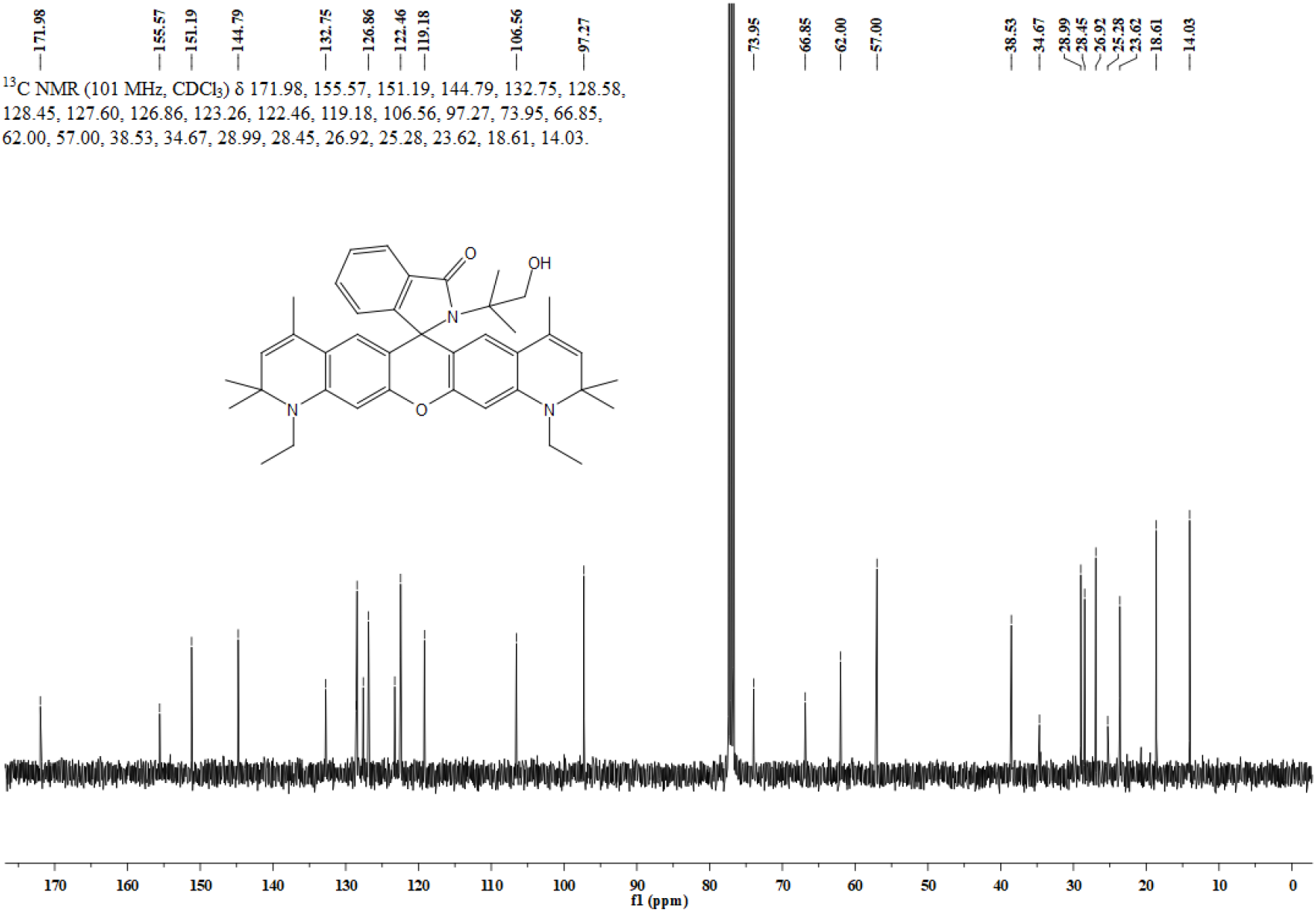
^13^C NMR spectrum of **4a** in CDCl_3_.

**Figure S69.**
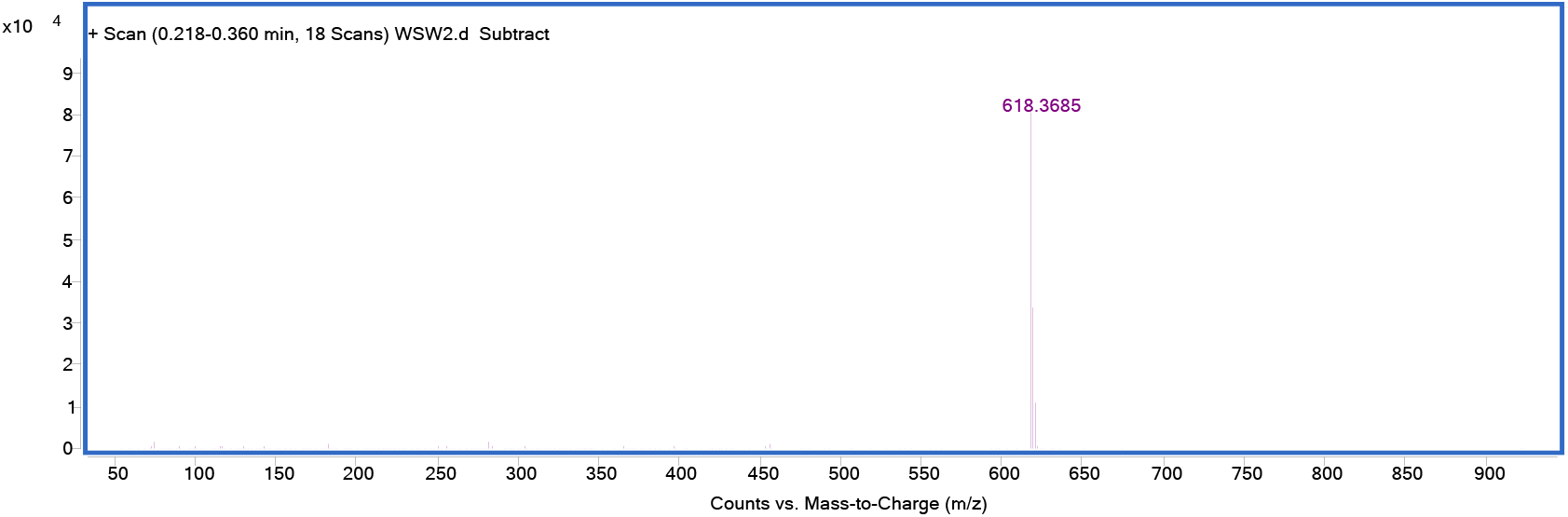
HRMS spectrum of **4a**.

**Figure S70.**
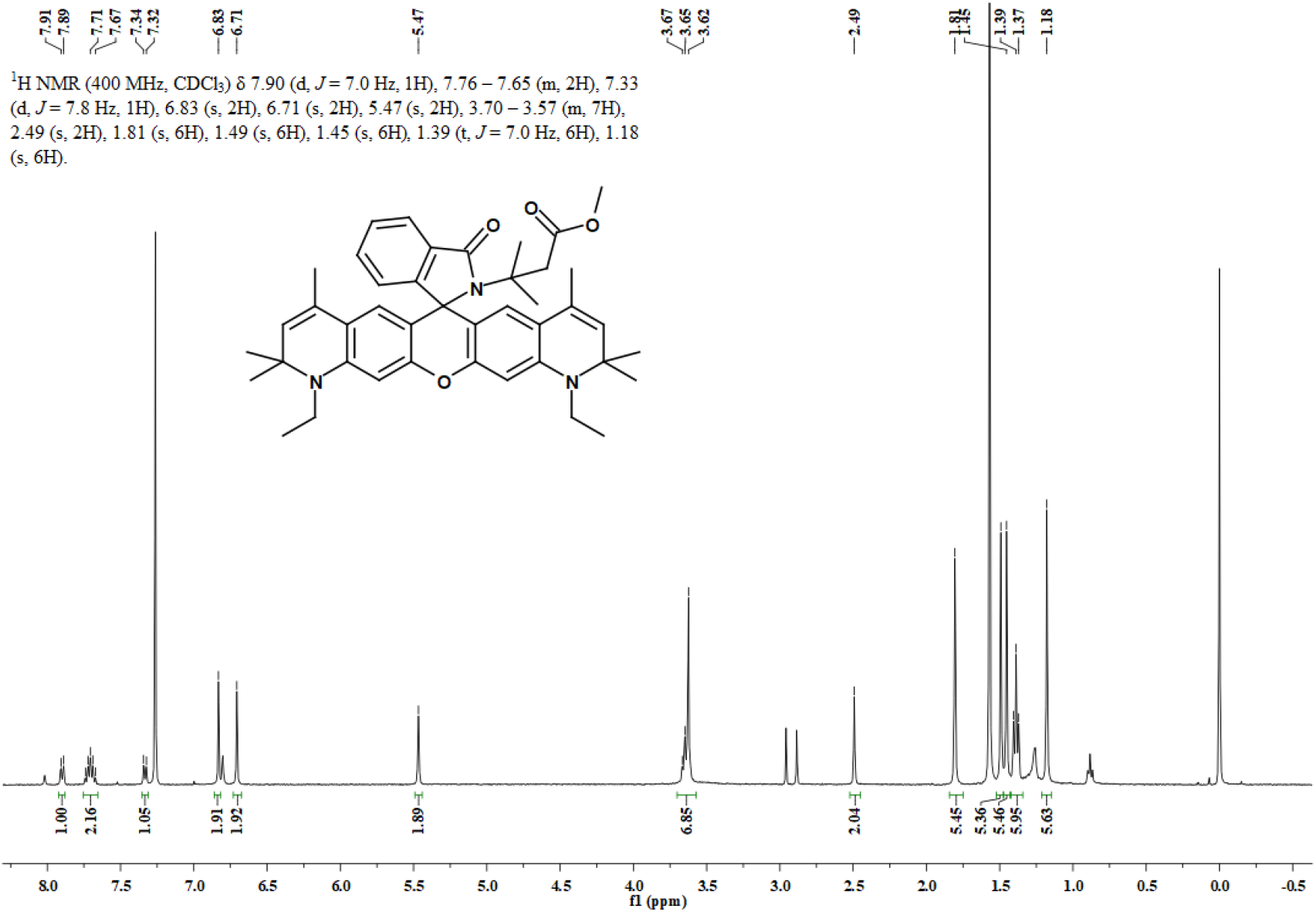
^1^H NMR spectrum of **18** in CDCl_3_.

**Figure S71.**
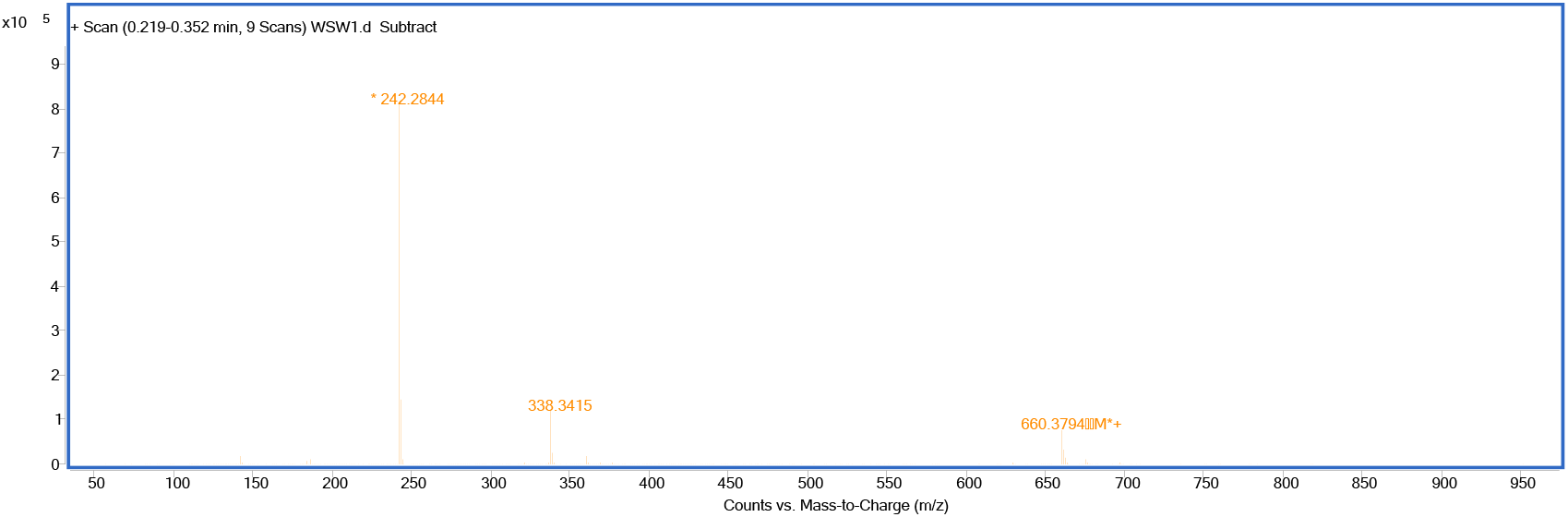
HRMS spectrum of **18**.

**Figure S72.**
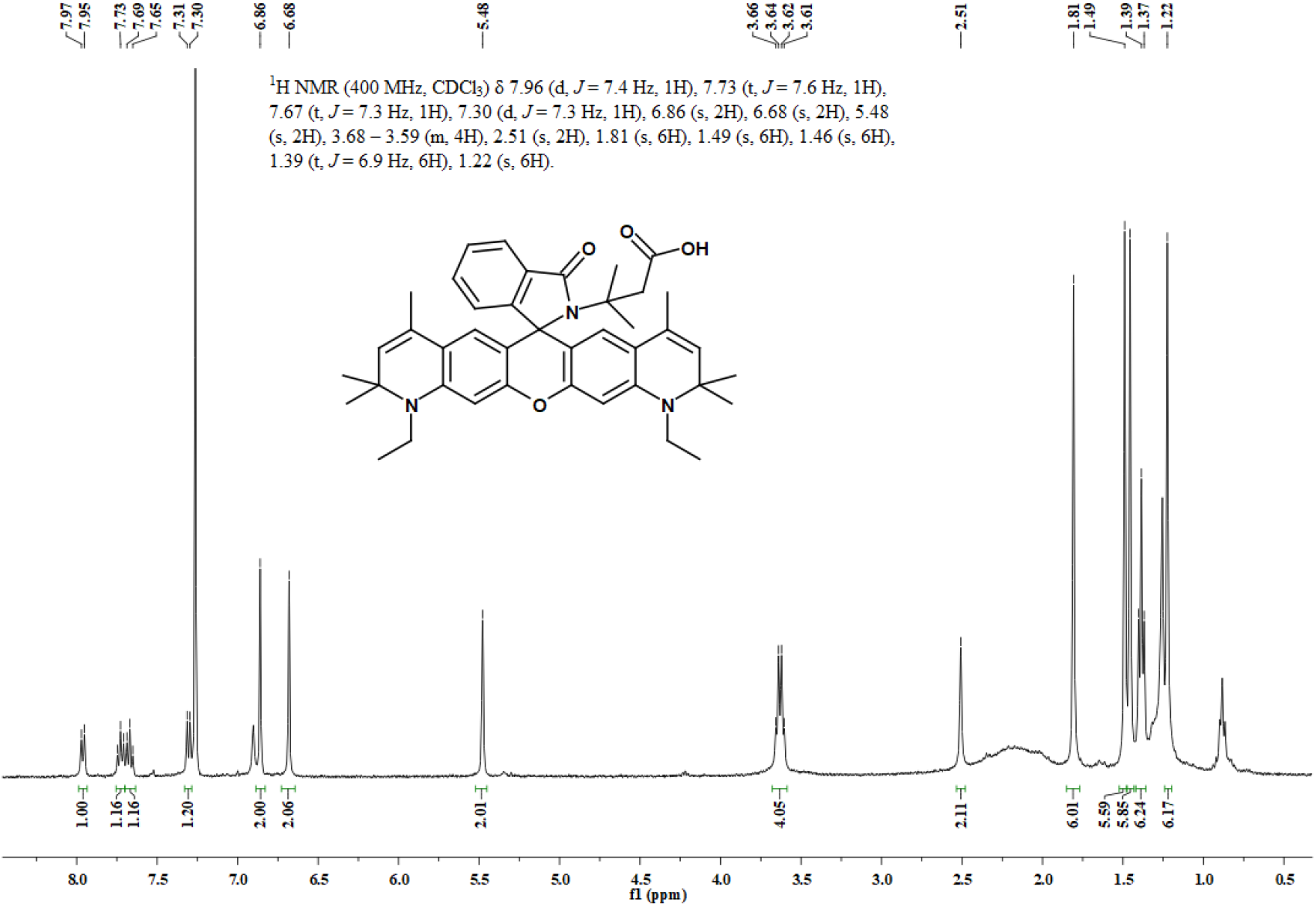
^1^H NMR spectrum of **19** in CDCl_3_.

**Figure S73.**
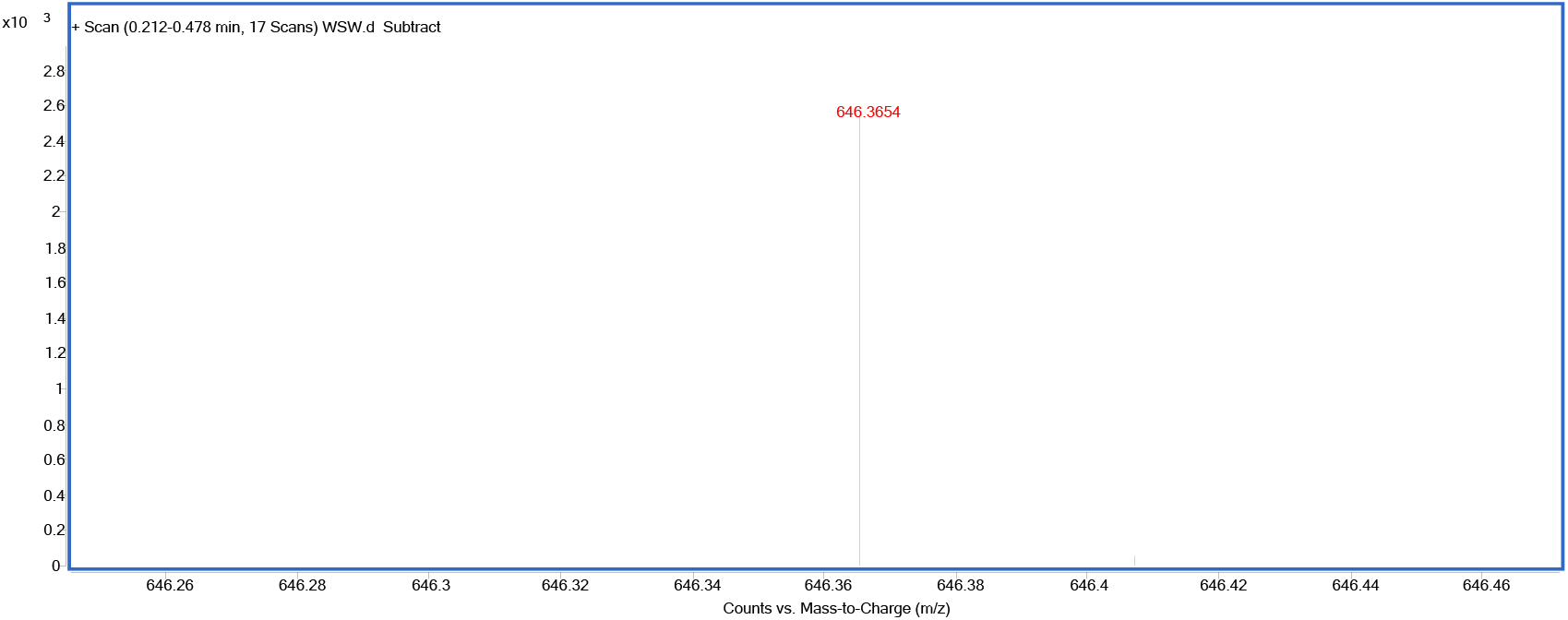
HRMS spectrum of **19**.

**Figure S74.**
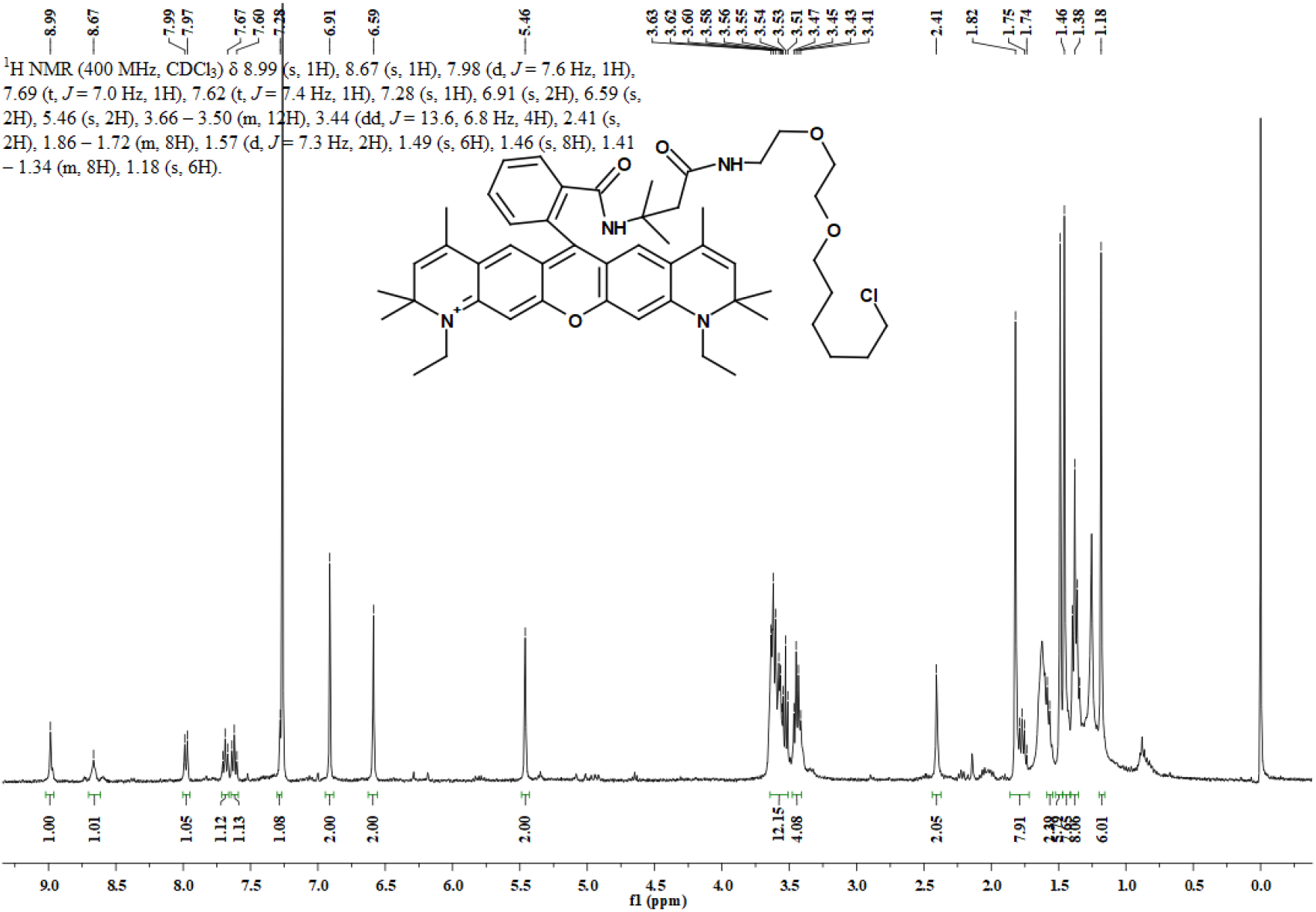
^1^H NMR spectrum of **N-halo-JA21** in CDCl_3_.

**Figure S75.**
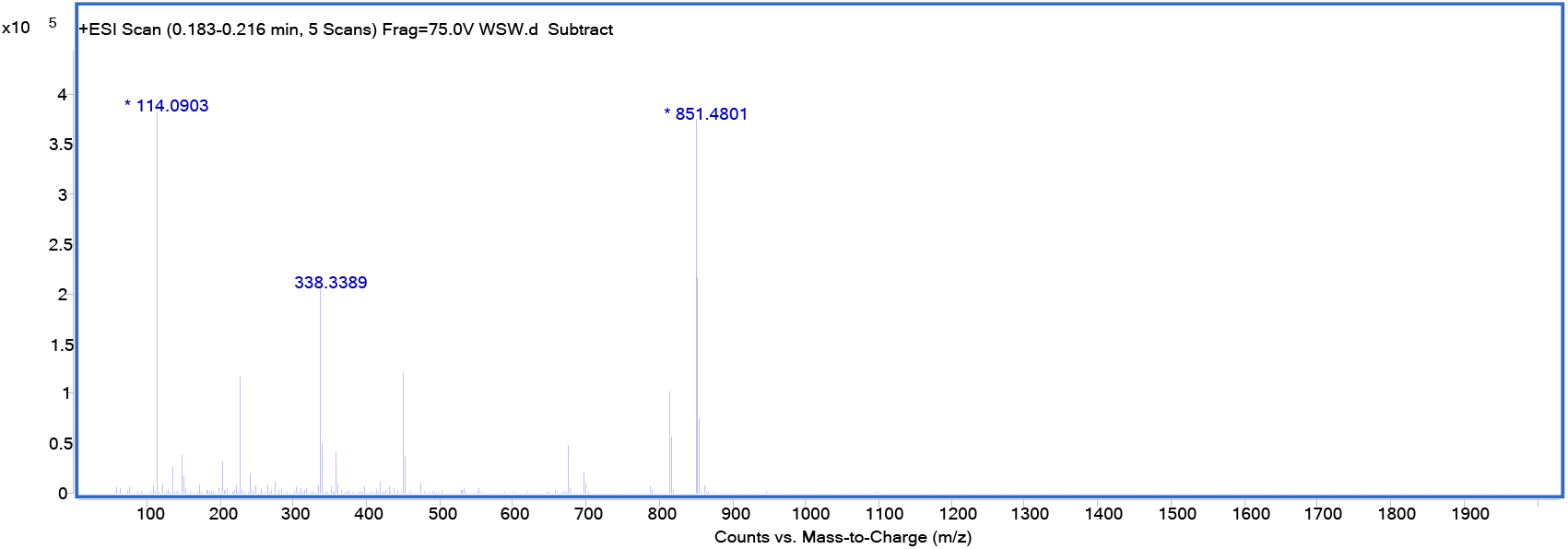
HRMS spectrum of **N-halo-JA21**.

**Figure S76.**
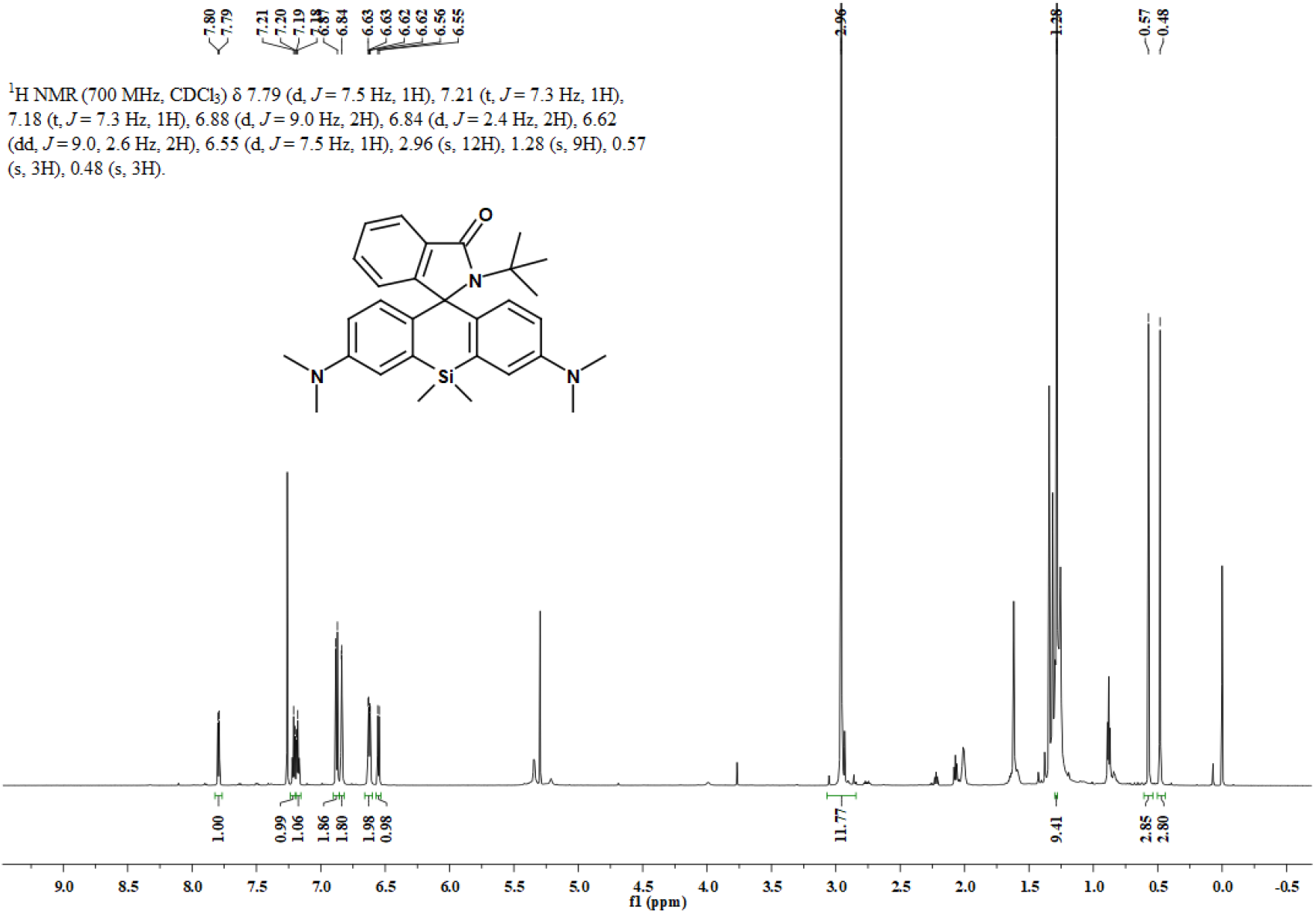
^1^H NMR spectrum of **21** in CDCl_3_.

**Figure S77.**
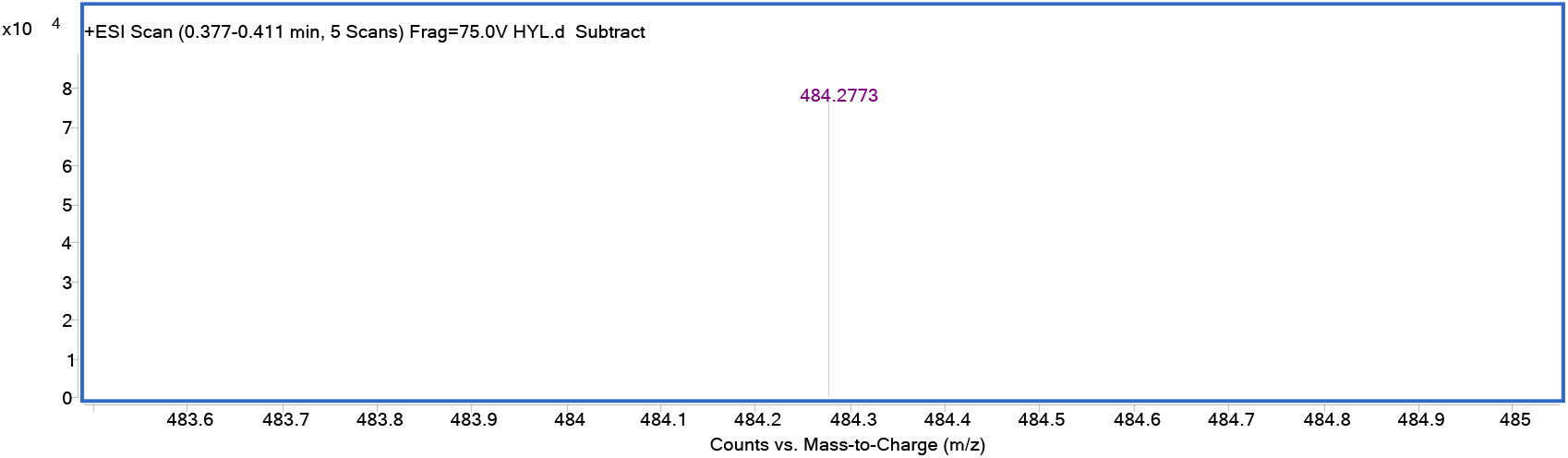
HRMS spectrum of **21**.

